# The Trans-Ancestral Genomic Architecture of Glycaemic Traits

**DOI:** 10.1101/2020.07.23.217646

**Authors:** Ji Chen, Cassandra N. Spracklen, Gaëlle Marenne, Arushi Varshney, Laura J Corbin, Jian’an Luan, Sara Willems, Ying Wu, Xiaoshuai Zhang, Momoko Horikoshi, Thibaud S Boutin, Reedik Mägi, Johannes Waage, Achilleas Pitsilides, Ruifang Li-Gao, Kei Hang, Jie Yao, Mila D Anasanti, Audrey Y Chu, Annique Claringbould, Jani Heikkinen, Jaeyoung Hong, Jouke-Jan Hottenga, Shaofeng Huo, Marika A. Kaakinen, Tin Louie, Winfried März, Hortensia Moreno-Macias, Anne Ndungu, Sarah C. Nelson, Ilja M. Nolte, Kari North, Chelsea K. Raulerson, Debashree Ray, Rebecca Rohde, Denis Rybin, Claudia Schurmann, Xueling Sim, Loz Southam, Isobel Stewart, Carol A. Wang, Yujie Wang, Peitao Wu, Weihua Zhang, Tarunveer S. Ahluwalia, Emil VR Appel, Lawrence F. Bielak, Jennifer A. Brody, Noel P Burtt, Claudia P Cabrera, Brian E Cade, Jin Fang Chai, Xiaoran Chai, Li-Ching Chang, Chien-Hsiun Chen, Brian H Chen, Kumaraswamy N Chitrala, Yen-Feng Chiu, Hugoline G. de Haan, Graciela E Delgado, Ayse Demirkan, Qing Duan, Jorgen Engmann, Segun A Fatumo, Javier Gayán, Franco Giulianini, Jung Ho Gong, Stefan Gustafsson, Yang Hai, Fernando P Hartwig, Jing He, Yoriko Heianza, Tao Huang, Alicia Huerta-Chagoya, Mi Yeong Hwang, Richard A. Jensen, Takahisa Kawaguchi, Katherine A Kentistou, Young Jin Kim, Marcus E Kleber, Ishminder K Kooner, Shuiqing Lai, Leslie A Lange, Carl D Langefeld, Marie Lauzon, Man Li, Symen Ligthart, Jun Liu, Marie Loh, Jirong Long, Valeriya Lyssenko, Massimo Mangino, Carola Marzi, May E Montasser, Abhishek Nag, Masahiro Nakatochi, Damia Noce, Raymond Noordam, Giorgio Pistis, Michael Preuss, Laura Raffield, Laura J. Rasmussen-Torvik, Stephen S Rich, Neil R Robertson, Rico Rueedi, Kathleen Ryan, Serena Sanna, Richa Saxena, Katharina E Schraut, Bengt Sennblad, Kazuya Setoh, Albert V Smith, Lorraine Southam, Thomas Sparsø, Rona J Strawbridge, Fumihiko Takeuchi, Jingyi Tan, Stella Trompet, Erik van den Akker, Peter J Van der Most, Niek Verweij, Mandy Vogel, Heming Wang, Chaolong Wang, Nan Wang, Helen R Warren, Wanqing Wen, Tom Wilsgaard, Andrew Wong, Andrew R Wood, Tian Xie, Mohammad Hadi Zafarmand, Jing-Hua Zhao, Wei Zhao, Najaf Amin, Zorayr Arzumanyan, Arne Astrup, Stephan JL Bakker, Damiano Baldassarre, Marian Beekman, Richard N Bergman, Alain Bertoni, Matthias Blüher, Lori L. Bonnycastle, Stefan R Bornstein, Donald W Bowden, Qiuyin Cai, Archie Campbell, Harry Campbell, Yi Cheng Chang, Eco J.C. de Geus, Abbas Dehghan, Shufa Du, Gudny Eiriksdottir, Aliki Eleni Farmaki, Mattias Frånberg, Christian Fuchsberger, Yutang Gao, Anette P Gjesing, Anuj Goel, Sohee Han, Catharina A Hartman, Christian Herder, Andrew A. Hicks, Chang-Hsun Hsieh, Willa A. Hsueh, Sahoko Ichihara, Michiya Igase, M. Arfan Ikram, W. Craig Johnson, Marit E Jørgensen, Peter K Joshi, Rita R Kalyani, Fouad R. Kandeel, Tomohiro Katsuya, Chiea Chuen Khor, Wieland Kiess, Ivana Kolcic, Teemu Kuulasmaa, Johanna Kuusisto, Kristi Läll, Kelvin Lam, Deborah A Lawlor, Nanette R. Lee, Rozenn N. Lemaitre, Honglan Li, Lifelines Cohort Study, Shih-Yi Lin, Jaana Lindström, Allan Linneberg, Jianjun Liu, Carlos Lorenzo, Tatsuaki Matsubara, Fumihiko Matsuda, Geltrude Mingrone, Simon Mooijaart, Sanghoon Moon, Toru Nabika, Girish N. Nadkarni, Jerry L. Nadler, Mari Nelis, Matthew J Neville, Jill M Norris, Yasumasa Ohyagi, Annette Peters, Patricia A. Peyser, Ozren Polasek, Qibin Qi, Dennis Raven, Dermot F Reilly, Alex Reiner, Fernando Rivideneira, Kathryn Roll, Igor Rudan, Charumathi Sabanayagam, Kevin Sandow, Naveed Sattar, Annette Schürmann, Jinxiu Shi, Heather M Stringham, Kent D. Taylor, Tanya M. Teslovich, Betina Thuesen, Paul RHJ Timmers, Elena Tremoli, Michael Y Tsai, Andre Uitterlinden, Rob M van Dam, Diana van Heemst, Astrid van Hylckama Vlieg, Jana V Van Vliet-Ostaptchouk, Jagadish Vangipurapu, Henrik Vestergaard, Tao Wang, Ko Willems van Dijk, Yongbing Xiang, Tatijana Zemunik, Goncalo R Abecasis, Linda S. Adair, Carlos Alberto Aguilar-Salinas, Marta E Alarcón-Riquelme, Ping An, Larissa Aviles-Santa, Diane M Becker, Lawrence J Beilin, Sven Bergmann, Hans Bisgaard, Corri Black, Michael Boehnke, Eric Boerwinkle, Bernhard O Böhm, Klaus Bønnelykke, D I. Boomsma, Erwin P. Bottinger, Thomas A Buchanan, Mickaël Canouil, Mark J Caulfield, John C. Chambers, Daniel I. Chasman, Yii-Der Ida Chen, Ching-Yu Cheng, Francis S. Collins, Adolfo Correa, Francesco Cucca, H. Janaka de Silva, George Dedoussis, Sölve Elmståhl, Michele K. Evans, Ele Ferranni, Luigi Ferruci, Jose C Florez, Paul Franks, Timothy M Frayling, Philippe Froguel, Bruna Gigante, Mark O. Goodarzi, Penny Gordon-Larsen, Harald Grallert, Niels Grarup, Sameline Grimsgaard, Leif Groop, Vilmundur Gudnason, Xiuqing Guo, Anders Hamsten, Torben Hansen, Caroline Hayward, Susan R. Heckbert, Bernardo L Horta, Wei Huang, Erik Ingelsson, Pankow S James, Jost B Jonas, J. Wouter Jukema, Pontiano Kaleebu, Robert Kaplan, Sharon L.R. Kardia, Norihiro Kato, Sirkka M. Keinanen-Kiukaanniemi, Bong-Jo Kim, Mika Kivimaki, Heikki A. Koistinen, Jaspal S. Kooner, Antje Körner, Peter Kovacs, Diana Kuh, Meena Kumari, Zoltan Kutalik, Markku Laakso, Timo A. Lakka, Lenore J Launer, Karin Leander, Huaixing Li, Xu Lin, Lars Lind, Cecilia Lindgren, Simin Liu, Ruth J.F. Loos, Patrik Magnusson, Anubha Mahajan, Andres Metspalu, Dennis O Mook-Kanamori, Trevor A Mori, Patricia B Munroe, Inger Njølstad, Jeffrey R O’Connell, Albertine J Oldehinkel, Ken K Ong, Sandosh Padmanabhan, Colin N.A. Palmer, Nicholette D Palmer, Oluf Pedersen, Craig E Pennell, David J Porteous, Peter P. Pramstaller, Michael A. Province, Bruce M. Psaty, Lu Qi, Leslie J. Raffel, Rainer Rauramaa, Susan Redline, Paul M Ridker, Frits R. Rosendaal, Timo E. Saaristo, Manjinder Sandhu, Jouko Saramies, Neil Schneiderman, Peter Schwarz, Laura J. Scott, Elizabeth Selvin, Peter Sever, Xiao-ou Shu, P Eline Slagboom, Kerrin S Small, Blair H Smith, Harold Snieder, Tamar Sofer, Thorkild I.A. Sørensen, Tim D Spector, Alice Stanton, Claire J Steves, Michael Stumvoll, Liang Sun, Yasuharu Tabara, E Shyong Tai, Nicholas J Timpson, Anke Tönjes, Jaakko Tuomilehto, Teresa Tusie, Matti Uusitupa, Pim van der Harst, Cornelia van Duijn, Veronique Vitart, Peter Vollenweider, Tanja GM Vrijkotte, Lynne E Wagenknecht, Mark Walker, Ya X Wang, Nick J Wareham, Richard M Watanabe, Hugh Watkins, Wen B Wei, Ananda R Wickremasinghe, Gonneke Willemsen, James F Wilson, Tien-Yin Wong, Jer-Yuarn Wu, Anny H Xiang, Lisa R Yanek, Loïc Yengo, Mitsuhiro Yokota, Eleftheria Zeggini, Wei Zheng, Alan B Zonderman, Jerome I Rotter, Anna L Gloyn, Mark I. McCarthy, Josée Dupuis, James B Meigs, Robert Scott, Inga Prokopenko, Aaron Leong, Ching-Ti Liu, Stephen CJ Parker, Karen L. Mohlke, Claudia Langenberg, Eleanor Wheeler, Andrew P. Morris, Inês Barroso

## Abstract

Glycaemic traits are used to diagnose and monitor type 2 diabetes, and cardiometabolic health. To date, most genetic studies of glycaemic traits have focused on individuals of European ancestry. Here, we aggregated genome-wide association studies in up to 281,416 individuals without diabetes (30% non-European ancestry) with fasting glucose, 2h-glucose post-challenge, glycated haemoglobin, and fasting insulin data. Trans-ancestry and single-ancestry meta-analyses identified 242 loci (99 novel; *P*<5×10^-8^), 80% with no significant evidence of between-ancestry heterogeneity. Analyses restricted to European ancestry individuals with equivalent sample size would have led to 24 fewer new loci. Compared to single-ancestry, equivalent sized trans-ancestry fine-mapping reduced the number of estimated variants in 99% credible sets by a median of 37.5%. Genomic feature, gene-expression and gene-set analyses revealed distinct biological signatures for each trait, highlighting different underlying biological pathways. Our results increase understanding of diabetes pathophysiology by use of trans-ancestry studies for improved power and resolution.

## Introduction

Fasting glucose (FG), 2h-glucose post-challenge (2hGlu), and glycated haemoglobin (HbA1c) are glycaemic traits used to diagnose diabetes^1^. In addition, HbA1c is the most commonly used biomarker to monitor glucose control in patients with diabetes. Fasting insulin (FI) reflects a combination of insulin resistance, a component of type 2 diabetes (T2D), and insulin clearance^2^. Collectively, all four of these glycaemic traits can be useful to better understand T2D pathophysiology^3–5^, are useful measures of cardiometabolic health as they are associated with cardiometabolic outcomes even within the non-diabetic range, albeit modestly so^6^.

To date, genome-wide association studies (GWAS) and analysis of next-generation targeted arrays (Metabochip and exome array) have identified >120 loci associated with glycaemic traits in individuals without diabetes^7–15^. However, despite considerable differences in the prevalence of T2D risk factors across ancestries^16–18^, most glycaemic trait GWAS in individuals without diabetes have insufficient representation of individuals of non-European ancestry and limited resolution for fine-mapping of causal variants and effector transcript identification. Here, we present large-scale trans-ancestry discovery meta-analyses of GWAS for four glycaemic traits (FG, 2hGlu, FI, and HbA1c) in individuals without diabetes with genotype imputation to the 1000 Genomes Project reference panel phase 1 version 3^19^. Our aims were to identify additional glycaemic trait-associated loci; investigate the portability of loci and genetic scores across ancestries; leverage differences in effect allele frequency (EAF), effect size, and linkage disequilibrium (LD) across diverse populations to conduct fine-mapping and aid causal variant/effector transcript identification; and compare and contrast the genetic architecture of these four glycaemic traits to further elucidate their underlying biology and gain insights into pathophysiological pathways implicated in T2D.

## Results

### Study design, lead variant, index variant and trans-ancestry locus definitions

To identify loci associated with glycaemic traits FG, 2hGlu, FI, and HbA1c, we aggregated GWAS in up to 281,416 individuals without diabetes, ~30% of whom were of non-European ancestry [13% East Asian, 7% Hispanic, 6% African-American, 3% South Asian, and 2% sub-Saharan African (Ugandan - data only available for HbA1c)]. Prior to meta-analysis each contributing cohort imputed data to the 1000 Genomes Project reference panel (phase 1 v3, March 2012, or later; **Methods, Supplementary Table 1, Supplementary Figure 1**). In total, up to ~49.3 million variants were directly genotyped or imputed, with between 38.6 million (2hGlu) and 43.5 million variants (HbA1c) available for analysis after exclusions based on minor allele count (MAC < 3) and imputation quality (imputation r^2^ or INFO score <0.40) in each cohort. As we had previously found adjusting for body mass index (BMI) provided similar results for FG and 2hGlu, but aided in new locus discovery for FI^15^, here we conducted analyses for FG, 2hGlu and FI adjusted for BMI, but for simplicity these traits are abbreviated as FG, 2hGlu and FI (**Methods**).

We first performed trait-specific fixed-effect meta-analyses *within* each ancestry using METAL^20^. We defined “single-ancestry lead” variants as the strongest trait-associated variants (*P*<5×10^-8^) within a 1Mb region in a particular ancestry (**Glossary box**). Within each ancestry and each autosome, we used approximate conditional analyses in GCTA^21,22^, to identify distinct “single-ancestry index variants” (*P*<5×10^-8^) that exert conditionally distinct effects on the trait (**Glossary Box, Methods, Supplementary Figure 2**). Overall, this approach identified 124 distinct FG, 15 2hGlu, 48 FI and 139 HbA1c variants that were significant in at least one ancestry (**Supplementary Table 2**).

#### Glossary Box

This study combined analyses of trait-associations across multiple correlated glycaemic traits and across multiple ancestries, which has presented challenges in our ability to apply commonly used terms with clarity. For this reason, we define below terms often used in the field with variable meaning, as well as definitions of new terms used in this study.

**EA** – the effect allele was that defined by METAL based on trans-ancestry FG results and aligned such that the same allele was kept as the effect allele across all ancestries and traits, irrespective of its allele frequency or effect size for that particular ancestry and trait, in this way the effect allele is not necessarily the trait-increasing allele.

**Single-ancestry lead variant** – variant with the smallest p-value amongst all with *P* < 5×10^-8^, within a 1Mb region, based on analysis of a single trait in a single ancestry.

**Single-ancestry index variants** – variants identified by GCTA analysis of each autosome, and that appear to exert conditionally distinct effects on a given trait in a given ancestry (*P* < 5×10^-8^). As defined, these include the single-ancestry lead variant.

**Trans-ancestry lead variant** – variant identified by trans-ethnic meta-analysis of a given trait that has the strongest association for that trait (log_10_BF > 6, which is broadly equivalent to *P* < 5×10^-8^) within a 1Mb region.

**Single-ancestry locus** – a 1Mb region centred on a single-ancestry lead variant which does not contain a lead variant identified in the trans-ancestry meta-analysis (i.e., does not contain a trans-ancestry lead variant).

**Signal** - a conditionally independent association between a trait and a set of variants in LD with each other and which is noted by the corresponding index variant.

**Trans-ancestry locus** – As we expected some genetic variants to influence multiple correlated traits and that functional variants would influence traits across multiple ancestries, we combined results across traits and across ancestries into multi-trait trans-ancestry loci. A **trans-ancestry locus** is a genomic interval that contains trans-ancestry trait-specific lead variants, with/out additional single-ancestry index variants, for one or more trait. This region is defined by starting at the telomere of each chromosome and selecting the first single-ancestry index variant or trans-ancestry lead variant for any trait. If other trans-ancestry lead variants or single-ancestry index variants mapped within 500kb of the first signal, then they were merged into the same locus. This process was repeated until there were no more signals within 500kb of the previous variant. A 500kb interval was added to the beginning of the first signal, and the end of the last signal to establish the final boundary of the trans-ancestry locus. As defined, a trans-ancestry locus may not have a single lead trans-ancestry variant, but may instead contain multiple trans-ancestry lead variants, one for each trait.

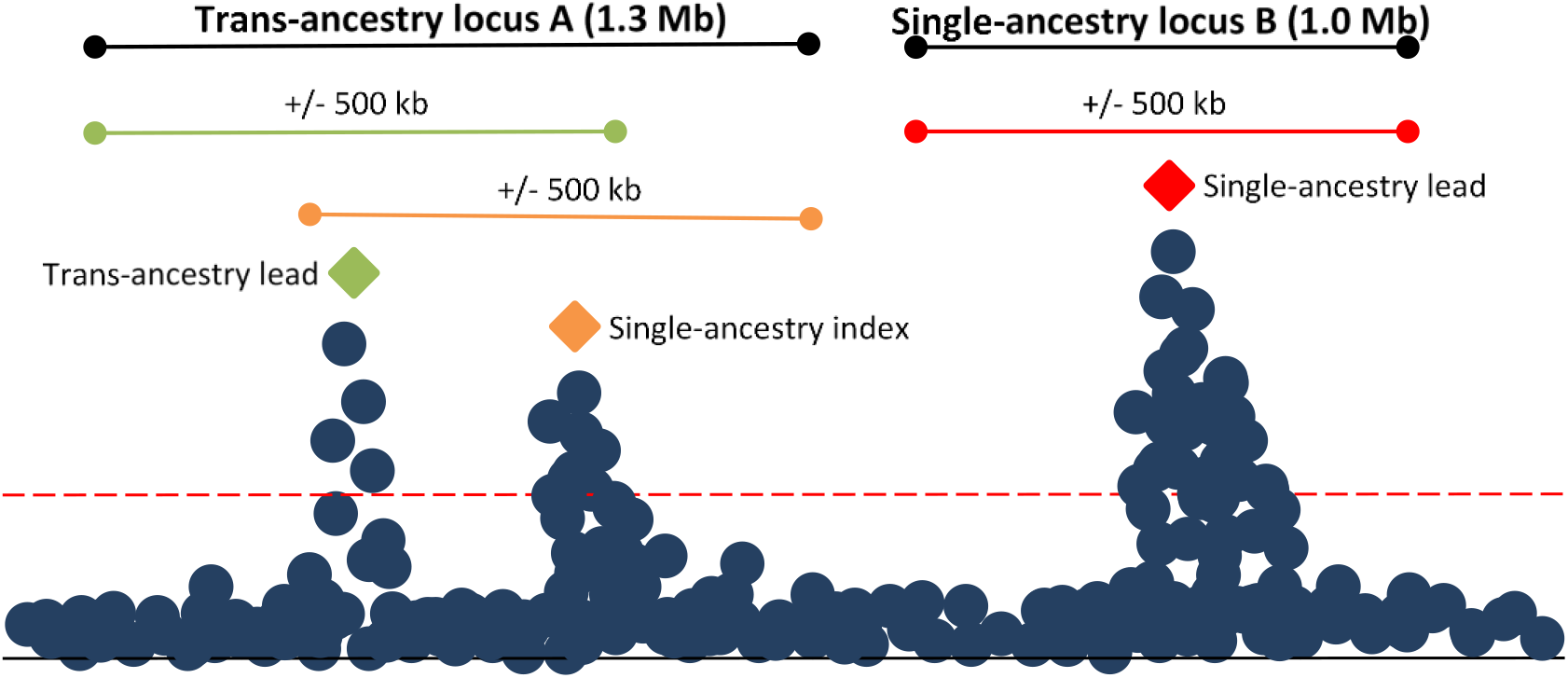

**Locus diagram—** In this diagram, trans-ancestry locus A contains a trans-ancestry lead variant for one glycaemic trait represented by the green diamond, and another single-ancestry index variant for another glycaemic trait represented by the orange triangle. Single-ancestry locus B contains a single-ancestry lead variant represented by the red square. The orange, green and red bars represent a +/- 500Kb window around the orange, green, and red variants, respectively. The black bars indicate the full locus window where trans-ancestry locus A contains trans-ancestry lead and single-ancestry index variants for two traits and single-ancestry locus B has a single-ancestry lead variant for a single trait.

Next, we conducted trait-specific *trans-ancestry* meta-analyses of ancestry-specific results using MANTRA (**Methods, Supplementary Table 1, Supplementary Figures 1 and 3**) to identify genome-wide significant “trans-ancestry lead variants”, defined as the most significant trait-associated variant across all ancestries (log_10_ Bayes Factor [BF] >6, equivalent to *P* < 5×10^-8 23^) (**Glossary box**, **Methods**). Here, we present trans-ancestry results based on data from all participating cohorts as our primary results (**Supplementary Table 2**).

Causal variants are expected to affect multiple related glycaemic traits and may be shared across ancestries. Therefore, we combined all single-ancestry lead variants, single-ancestry index variants, and/or trans-ancestry lead variants (for any trait) mapping within 500Kb of each other, into a single “trans-ancestry locus” that was bounded by a 500Kb flanking sequence (**Glossary Box**). As defined, a trans-ancestry locus may contain multiple causal variants affecting one or more glycaemic traits, exerting their effect in one or more ancestry.

### Glycaemic trait locus discovery

In the trans-ancestry meta-analyses, we observed genome-wide significant associations at 235 trans-ancestry loci, of which 59 contained trans-ancestry lead variants for more than one trait. In addition, we identified seven “single-ancestry loci” that did not contain any trans-ancestry lead variants (**Glossary box, Supplementary Table 2**). Of the 242 trans-ancestry and single-ancestry loci, 99 (including 6 of the 7 single-ancestry) had not been previously associated with any of the four glycaemic traits or with T2D, at the time of analysis (**Figure 1, Supplementary Figures 1 and 3, Supplementary Table 3, Supplementary note**). Based on the largest European and East Asia ancestry T2D GWAS meta-analyses^23,24^, the lead variants at 19 novel glycaemic trait loci have strong evidence of association with T2D (*P*<10^-4^; six loci with *P*<5×10^-8^), suggesting some of the novel loci are also important in diabetes pathophysiology (**Supplementary Tables 2 and 4**).

**Figure 1.**
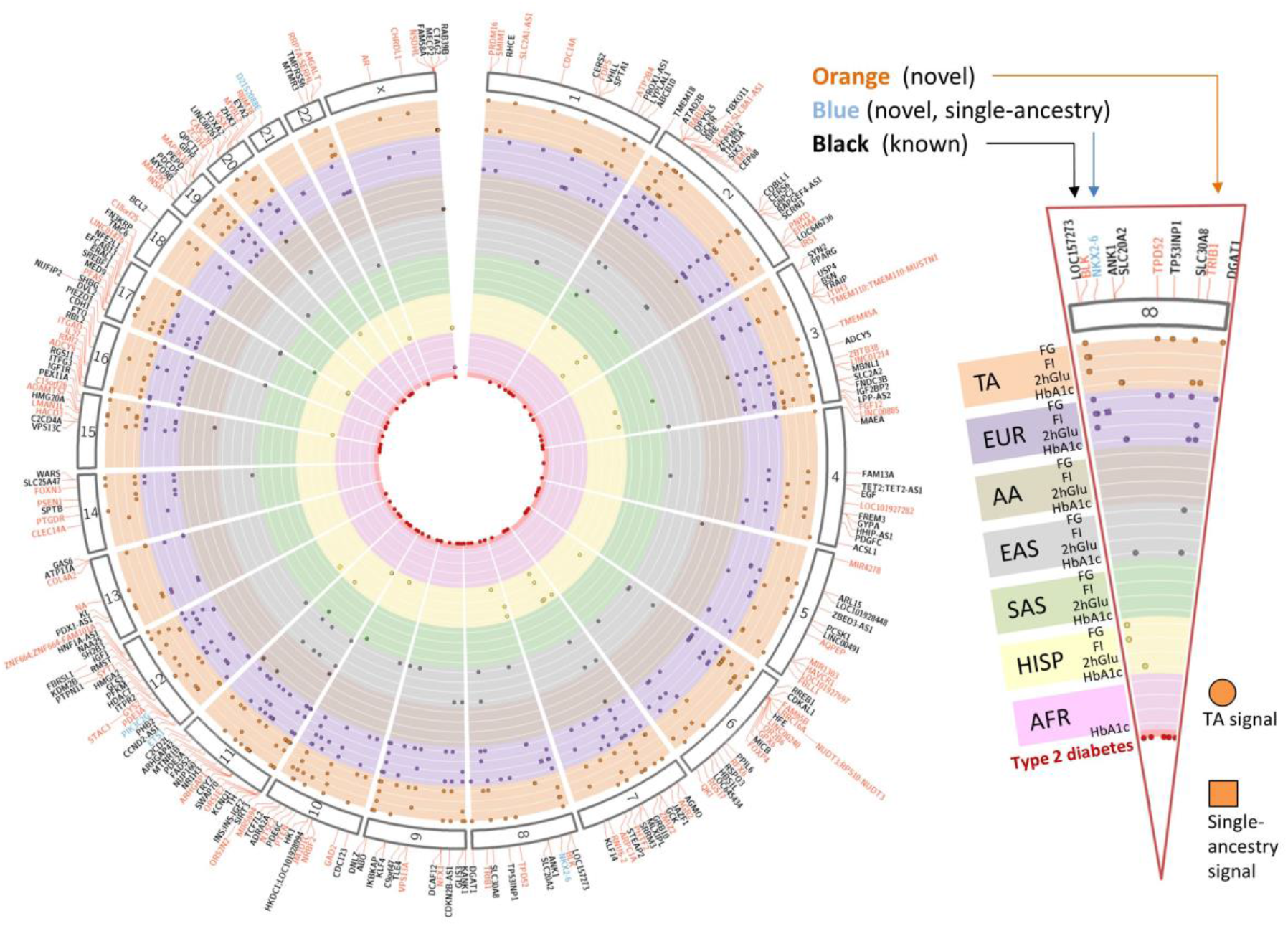
Summary of all 242 loci identified in this study. 235 trans-ancestry loci are shown in orange (novel) or black (established) along with seven single-ancestry loci (blue) represented by nearest gene. Each locus is mapped to corresponding chromosome (outer segment). Each set of rows shows the results from the trans-ancestry analysis (orange) and each of the ancestries: European (purple), African American (tan), East Asian (grey), South Asian (green), Hispanic (yellow), sub-Saharan African (Ugandan-pink). Loci with a corresponding type 2 diabetes signal are represented by red circles in the middle of the plot.

Of the 99 novel loci, six were identified in a single ancestry and did not overlap a trans-ancestry locus (**Supplementary Table 3**). Three single-ancestry loci were associated in individuals of non-European ancestry: (i) an African American association for FG (lead variant rs61909476) near the gene *ETS1*, (ii) an African American association for FI (lead variant rs12056334) near the gene *LOC100128993* (an uncharacterised RNA gene; **Supplementary Note**), and (iii) a Hispanic association for FG (lead variant rs12315677) within the gene *PIK3C2G* (**Supplementary Table 3**). The associations of rs61909476 and rs12315677 with FG are noteworthy. The variant rs61909476 has a similar EAF in both European (~10%) and African American (~7%) ancestry populations (**Supplementary Table 2**), but the effect on FG is only detectable in African American individuals (β=0.0812 mmol/l, SE=0.01 mmol/l, *P*=3.9×10^-8^, European individuals β=0.0015 mmol/l, SE=0.0031 mmol/l, *P*=0.44) (**Supplementary Figure 4, Supplementary note**). The nearest gene, *ETS1*, encodes a transcription factor which has been shown to localize to insulin-positive cells in mouse islets, and its overexpression was shown to decrease glucose-stimulated insulin secretion in mouse islets^25^. Located within the *PIK3C2G* gene, rs12315677 has a similar EAF in both Hispanic and European ancestry populations (84% and 86%, respectively), but is significantly associated with FG only in our Hispanic GWAS (β=0.0387 mmol/l, SE=0.0075 mmol/l, *P*=4.0×10^-8^) compared with European ancestry (β=-0.0029 mmol/l, SE=0.0029 mmol/l, *P*=0.39) (**Supplementary Figure 5, Supplementary note**). *PIK3C2G* has been shown to be a Rab5 effector which, when deleted in *Pik3c2g^-/-^* mice, selectively inhibits *Akt2* activation and leads to a phenotype characterised by reduced glycogen storage in the liver, hyperlipidaemia, adiposity, and insulin resistance with increasing age, or after a high fat diet^26^. Instances where the EAFs are similar between populations, but the effect sizes differ, could be due to specific genotype-by-environment effects that differ across ancestries, or lower imputation accuracy in ancestries with smaller sample sizes, although this would likely lead to deflated effect sizes and imputation quality is good for these variants (average r^2^=0.81). It is also possible that the variants detected here are not themselves causal, but are in LD with ancestry-specific causal variants that are not directly interrogated in our meta-analysis and that differ in frequency across ancestries. To try and investigate this hypothesis, we looked at data from 1000G in the cognate populations for evidence of rarer alleles in those ancestries that may themselves be driving the association signals (**Supplementary Table 5).** We could not detect evidence for other rarer alleles driving these associations, but this does not preclude the possibility that other rarer variants exist which are not represented in the 1000G populations. The final three single-ancestry loci were identified in individuals of European ancestry, but without any evidence of association in the other ancestries despite similar MAF, although this may be due to differences in power given the much smaller sample sizes in non-European ancestries (**Supplementary Figures 6-8**).

Next, we investigated the contribution of non-European ancestry data to novel trans-ancestry locus discovery, independent of the total sample size in the trans-ancestry meta-analysis. To do this, we artificially boosted the sample size of the European meta-analysis to match that of trans-ancestry meta-analysis by rescaling the standard errors of allelic effect sizes (**Supplementary note**). Using this approach, we determined that 21 of the novel trans-ancestry loci would not have been discovered if the sample size obtained in the trans-ancestry analyses was comprised exclusively of European ancestry individuals (**Supplementary note**). Instead, their discovery was due to the higher EAF and/or larger effect size in non-European ancestry populations. In particular, two loci (nearest genes *LINC00885* and *MIR4278*) contain East Asian and African American single-ancestry lead variants, respectively, suggesting that these specific ancestries may be driving the trans-ancestry discovery (**Supplementary Tables 2-3**). Combined with the three single-ancestry non-European loci described above, our results show that 24% (24/99) of novel loci were discovered due to the contribution of non-European ancestry participants, strengthening the argument for extending genetic studies to larger samples sizes in diverse populations.

### Allelic architecture of glycaemic traits

Trans-ancestry and single-ancestry loci comprised a range of association patterns, with most loci harbouring one single-ancestry signal for any given trait (**Supplementary note**). However, 29 loci contained multiple distinct index variants that did not fully overlap between ancestries. The most complex locus we observed was in the region spanning *G6PC2*, which contained 14 distinct FG index variants in the European single-ancestry meta-analysis. Of these, four are shared (*P*<5×10^-8^) with South Asian ancestry, two with East Asian ancestry, and two with Hispanic ancestry (**Supplementary Figure 9**). The complexity of association signals at this locus is consistent with previous work that also reported common variant (MAF>5%) association signals and multiple rare variant (MAF≤1%) associations at this locus that influenced protein function by multiple mechanisms^27^.

Combined, single-ancestry lead, single-ancestry index, and trans-ancestry lead variants increase the number of established loci for FG to 102 (182 signals, 53 novel loci), FI to 66 (95 signals, 49 novel loci), 2hGlu to 21 (28 signals, 11 novel loci), and HbA1c to 127 (218 signals, 62 novel loci) (**Supplementary Table 2**) and demonstrate significant overlap across glycaemic traits (**Supplementary Figure 10**). We also detected (*P*<0.05 or log_10_BF>0) the vast majority (~90%) of previously established glycaemic trait association signals in our data, 70-88% of which attained genome-wide significance in the current analyses (see further details in the **Supplementary Note**). Given that analyses for FG, FI, and 2hGlu were performed adjusted for BMI, we also confirmed that collider bias was not influencing discovery for more than 98% of our results (**Supplementary note**)^28^.

Finally, as expected, given the greater power due to increased sample sizes, new association signals tended to have smaller effect sizes and/or EAFs in European ancestry individuals (in whom this analysis was conducted) compared to previously established signals (**Supplementary Figure 11**).

### Characterisation of trans-ancestry lead variants and European index variants across ancestries

We next employed a series of complementary analyses to better understand the transferability of trans-ancestry lead variants across all ancestries. For each trans-ancestry lead variant, we investigated the pairwise EAF correlation between ancestries, as well as the pairwise summarised heterogeneity of effect sizes between ancestries^29^ (**Methods** and **Supplementary Note**). In agreement with population history and evolution, these results demonstrated considerable EAF correlation (ρ^2^>0.70) between European and Hispanic populations, European and South Asian populations, and Hispanic and South Asian populations, consistent across all four traits, and between African Americans and Ugandans for HbA1c (**Supplementary Figure 12**). Despite significant EAF correlations, some pairwise comparisons exhibited strong evidence for effect size heterogeneity between ancestries that was less consistent between traits (**Supplementary Figure 12).** However, sensitivity analyses demonstrated that, across all comparisons, the evidence for heterogeneity is driven by a small number of variants, with between 81.5% (for HbA1c) and 85.7% of trans-ancestry lead variants (for FG) showing no evidence for trans-ancestry heterogeneity (*P*>0.05) (**Supplementary Note**).

We also took LD pruned European single-ancestry index variants and compared the direction of effect of these variants in European ancestry individuals with that in other ancestries (**Supplementary Note**). Consistent with the lack of heterogeneity in effect sizes, we saw >70% concordance in the direction of effect for all traits into all ancestries, with the exception of HbA1c into African Americans and Ugandans (**Supplementary Table 6**). Imperfect concordance between ancestries could reflect lower power in non-European ancestry groups due to sample size or variation in allele frequency, or could be explained by LD differences between index SNPs and causal variants. For HbA1c, we hypothesized that lower concordance might also be a reflection of the different pathways (glycaemic and non-glycaemic) through which variants can affect HbA1c levels, particularly effects mediated via the red blood cell (RBC) where balancing selection can lead to different associations in individuals of African ancestry^7^ (**Supplementary Note** and below).

To further investigate the potential utility of trans-ancestry analyses, and to evaluate whether larger sample sizes might yield additional European ancestry signals that would be transferable across ancestries, we extended these concordance analyses to the entire genome, clumping variants mapping >1Mb apart (to eradicate the effect of LD in all ancestries) in different bins of association p-values obtained from the European ancestry meta-analysis (**Methods**). Aside from the bins with the weakest evidence for association in Europeans (i.e. in all bins with *P*≤0.05), we observed nominally significant concordance in the direction of effects between European and other ancestries for all traits except for 2hGlu, in which analyses were underpowered (**Supplementary Table 6**).

### Transferability of genetic scores (GS) across ancestries

To investigate the portability of GS across ancestries (the equivalent of genetic risk scores used for disease studies but instead for quantitative traits), we built a GS on the basis of effect sizes at European single-ancestry index variants (*P*<5×10^-8^), after LD pruning (r^2^<0.1), and assessed its utility for predicting trait variance explained in other ancestries (**Methods, Supplementary Note, Supplementary Table 7**). As a benchmark, we first assessed the predictive power (trait variance explained, as assessed by R^2^) of the GS into each cohort contributing to the European meta-analysis and three additional European cohorts that were not part of the meta-analysis. We then assessed the trait variance explained by the GS into the other ancestries and observed that the R2 fell within the range of values observed across European cohorts (**Figure 2A**).

**Figure 2.**
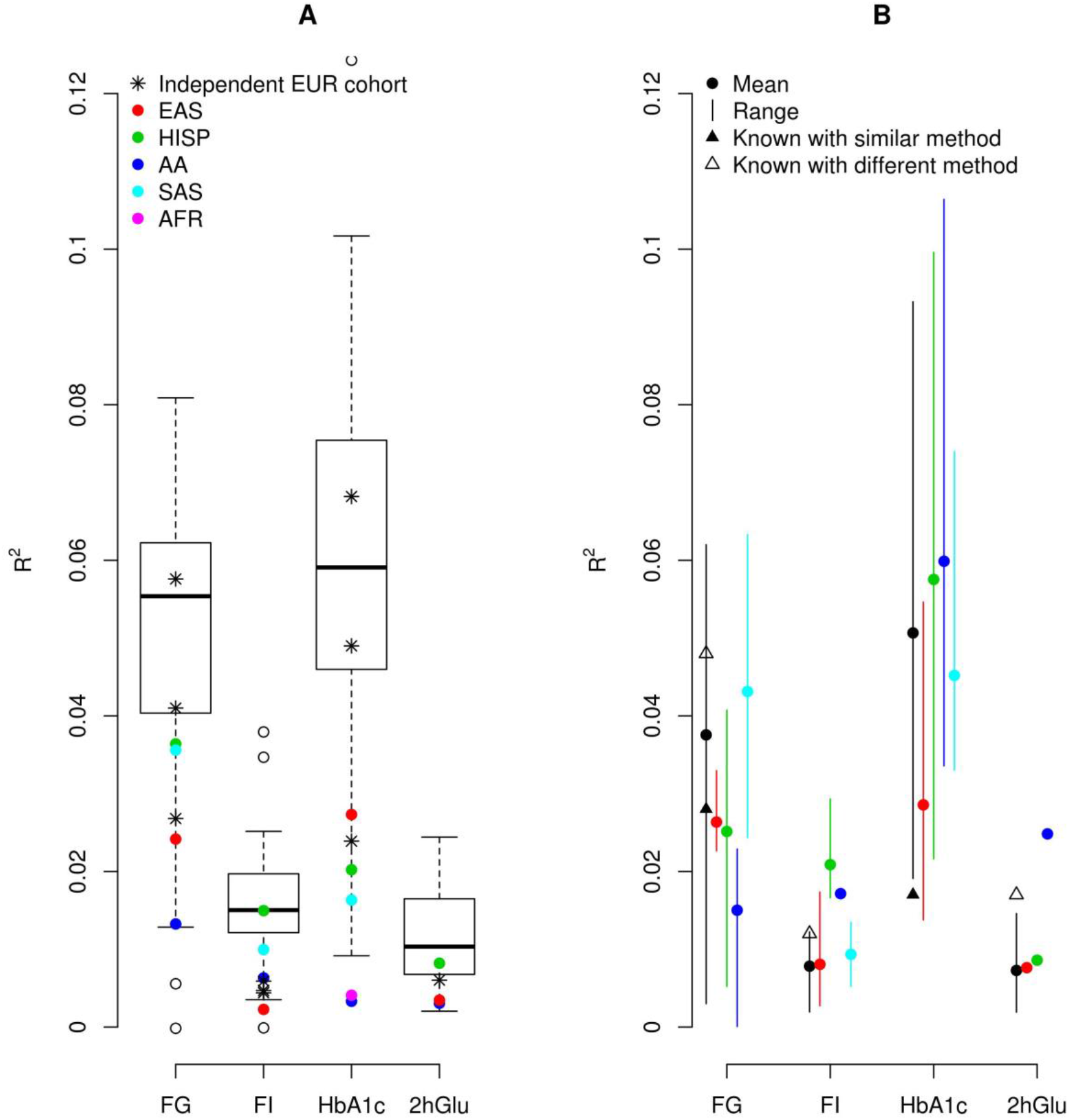
Transferability of GS across ancestries. Coloured dots represent data from the different ancestries: EUR in black, EAS in red, HISP in green, AA in blue and SAS in light blue. A) represents trait variance explained (FG, FI, HbA1c and 2hGlu) in each ancestry based on a GS build on the basis of effect sizes at European single-ancestry lead and index variants, after LD pruning (r^2^<0.1). The boxplot shows the maximum, first quantile, median, third quantile and minimum of variance explained in the EUR ancestry cohorts included in the study. The black asterisks show additional EUR cohorts that were not part of the original meta-analysis, while the dots represent variance explained in each of the other ancestries. B) represents trait variance explained when using a GS with a combination of individual trait trans-ancestry lead variants and single-ancestry lead and index variants, within each ancestry. Variance explained (mean and range of R^2^) for each trait (FG, FI, HbA1c, and 2hGlu) in each ancestry is shown. R^2^ was estimated in 1 to 11 cohorts with sample sizes ranging from 489 to 9,758 (**Supplementary Tables 9-12**). Closed and open triangles display previous known results using a similar method^30^ or a different method^15^.

We next expanded the GS to include all distance-based clumped variants across the genome with nominal evidence of association (*P*<1×10^-5^) in European ancestry individuals. This expansion improved the trait variance explained (greater *R*^2^) of the GS into European ancestry individuals compared to the GS built from LD pruned single-ancestry European lead and index variants (*P*<1×10^-8^) but substantially worsened performance into other ancestries (**Supplementary Table 8**).

Finally, using GS with a combination of individual trait trans-ancestry lead variants and single-ancestry lead and index variants within each ancestry, we were able to demonstrate that these explained, on average, between 0.7% (2hGlu in EUR) and 6% (HbA1c in AA) of the variance in trait distribution (**Methods, Figure 2B, Supplementary Tables 9-12**). In Europeans, these estimates represent an improvement (i.e. more variance explained) relative to previous estimates, derived using similar methodology, of 2.8% for FG and 1.7% for HbA1c^30^. Whilst variance explained estimates of 4.8% (FG), 1.2% (FI) and 1.7% (2hGlu) reported by Scott et al^15^ are in excess of our estimates, we hypothesise this is likely to be at least partly attributable to a difference in statistical approaches (see further discussion in **Supplementary Note**).

### Fine-mapping

Of the 242 identified loci, 231 were autosomal trans-ancestry loci and six were autosomal single-ancestry loci, which we took forward for fine-mapping (**Supplementary Table 2**). Due to the absence of LD maps from adequately sized populations, fine-mapping was not attempted for the 5 loci (4 trans-ancestry and 1 single-ancestry) mapping to the X chromosome. Using FINEMAP with ancestry-specific LD and an average LD matrix across ancestries, we conducted fine-mapping both within single-ancestries (all 237 autosomal loci) and across ancestries (231 autosomal trans-ancestry loci) for each trait (**Methods**). Because 59 of the 231 trans-ancestry loci were associated with more than one trait, we conducted trans-ancestry fine-mapping for a total of 305 locus-trait associations. Of these 305 locus-trait combinations, FINEMAP estimated the presence of a single causal variant responsible for the association at 186 loci (61%), while multiple distinct causal variants were implicated at 126 loci (39%), for a total of 464 causal variants (**Figure 3A**).

**Figure 3.**
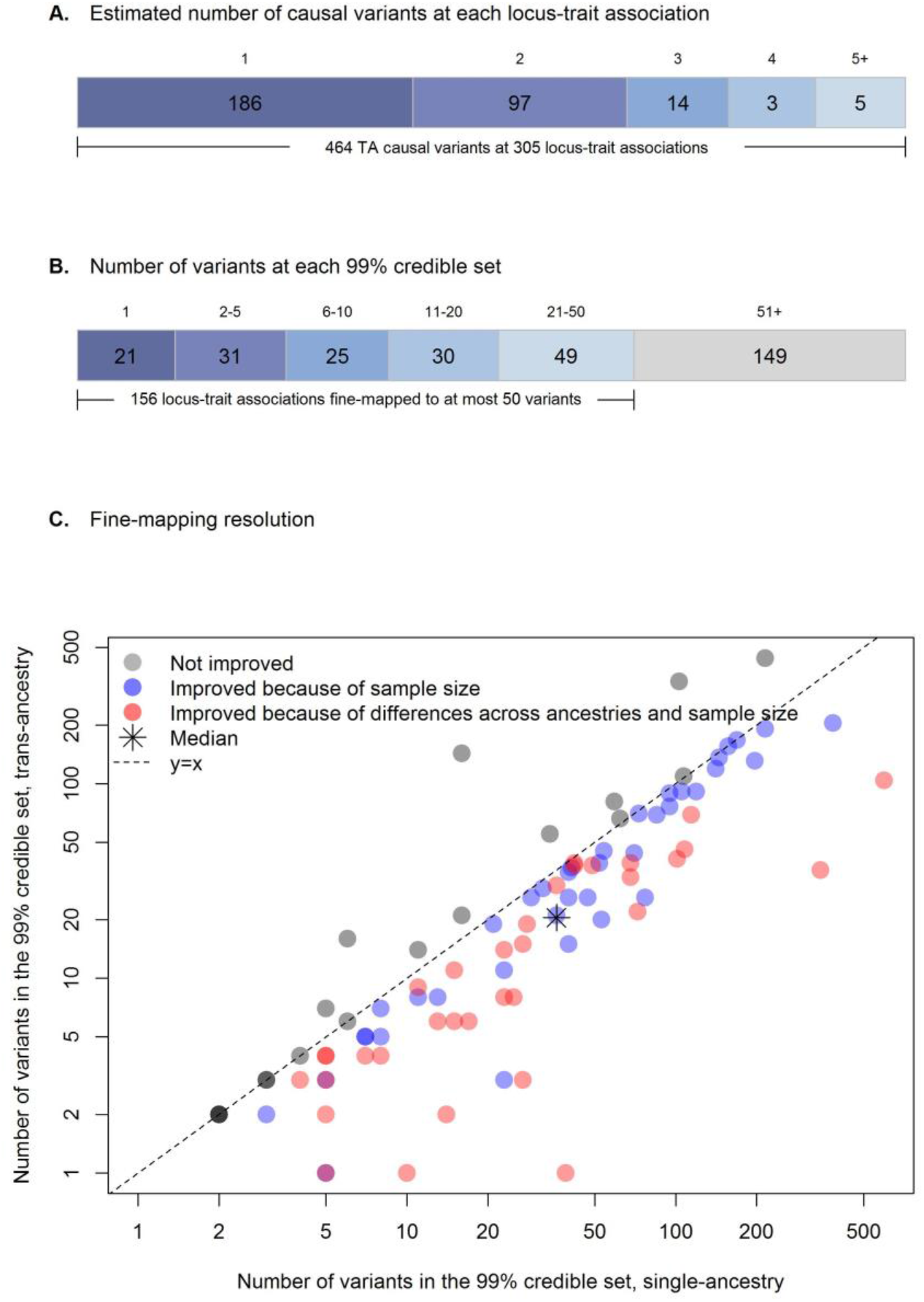
Trans-ancestry fine-mapping. A) Number of plausible causal variants at each locus-trait association derived from FINEMAP. B) Number of variants within each 99% credible set. Twenty-one locus-trait associations at 19 loci were mapped to a single variant in the 99% credible set. C) Fine-mapping resolution. For each of the 98 locus-trait associations with a predicted single causal variant in both trans-ancestry and single-ancestry analyses, the number of variants included in the 99% credible set in the single-ancestry fine-mapping (x axis; logarithmic scale) is plotted against those in the trans-ancestry fine-mapping (y axis; logarithmic scale). Trans-ancestry and single-ancestry fine-mapping were based on the same set of variants. After removing eight locus-trait associations with one variant in the 99% credible sets in both trans-ancestry and single-ancestry analyses, there were 18 locus-trait associations (in grey) where trans-ancestry fine-mapping did not improve the resolution of fine-mapping results (i.e. number of variants in the 99% credible set did not decrease). Of the 72 locus-trait associations with improved trans-ancestry fine-mapping resolution (blue and red) further analyses in European fine-mapping emulating the total sample size in trans-ancestry fine-mapping demonstrated that 34 locus-trait associations (in red) were improved because of both total sample size and differences across ancestries, while 38 locus-trait associations (in blue) were only improved due to increased sample size in the original trans-ancestry fine-mapping analysis.

#### Credible sets for causal variants

At each locus, we next constructed credible sets (CS) for each causal variant that account for >=99% of the posterior probability of association (PPA). We identified 21 locus-trait associations (at 19 loci) for which the 99% CS included a single variant, and we highlight five examples below. (**Methods, Supplementary Note, Figure 3B, Supplementary Table 13**).

First, we highlight two positive controls which provide confidence in the results. At one locus near *MTNR1B*, rs10830963 (PPA>0.999, for both HbA1c and FG), located in an *MTNR1B* intron, has shown allelic differences in enhancer activity and transcription factor binding^31^. At an additional FG-associated locus near *SIX3*, rs12712928 (PPA=0.997) has shown allelic differences in transcriptional activity, transcription factor binding, and association with islet expression levels of nearby genes *SIX3* and *SIX2*^32,33^. The EAF and effect size of this variant is larger in EAS than in other ancestries (heterogeneity p-value=7.2×10^-8^), which is driving the association at this locus.

Second, we highlight three novel findings. At a locus near *PFKM* associated with HbA1c, trans-ancestry fine-mapping identified rs12819124 (PPA>0.999) as the likely causal variant. This variant has been previously associated with mean corpuscular haemoglobin^34^, suggesting an effect of this locus on HbA1c is via the RBC. We note that this locus also harbours an association with FI in European and trans-ancestry meta-analyses, although it appears to be distinct from the HbA1c signal based on distance and LD. Fine-mapping of the nearby FI signal in European ancestry populations identified rs111264094 (PPA=0.994) as the likely causal variant (**Supplementary Figures 13-14**). rs111264094 is a low frequency variant in Europeans (EAF=0.025) that is monomorphic or rare in other ancestries, is located >600 kb from HbA1c-associated variant rs12819124, and is in low LD with rs12819124 in European ancestry populations (r^2^<0.1), which supports the hypothesis of two distinct signals (one for FI and one HbA1c) at this locus.

At the *HBB* locus, we also identify rs334 (PPA>0.999; Glu7Val) as the likely causal variant associated with HbA1c. rs334 is a causal variant of sickle cell anaemia^35^, with previously reported associations with urinary albumin-to-creatinine ratio in Caribbean Hispanic individuals^36^, severe malaria in a Tanzanian study population^37^, haematocrit and mean corpuscular volume in Hispanic/Latino populations^38^, and more recently with RBC distribution in Ugandan individuals^39^, all of which point to an effect of this variant on HbA1c via non-glycaemic pathways.

Lastly, our credible set analysis identified rs1799815 (PPA=0.993) as the likely causal variant at the *INSR* locus associated with FI. rs1799815 is a synonymous variant (Tyr3033Tyr) within *INSR*, the well-known insulin receptor gene that regulates the insulin signalling pathway. *INSR* as a target gene for this locus is further supported by our finding that rs1799815 colocalizes as an eQTL for *INSR* expression in adipose tissue (details shown below). The remaining locus-trait associations with a single variant in the 99% CS (**Supplementary Table 13**) point to variants that could be prioritised for downstream functional follow-up to further elucidate their impact on glycaemic trait physiology.

In addition to identifying 99% CS with a single variant, trans-ancestry fine-mapping identified 99% CS with 50 or fewer variants at 156 locus-trait associations (**Figure 3B, Supplementary Table 13**). Overall, 74 locus-trait associations contained 87 variants with PPA>0.90; that is, some locus-trait associations contain more than one variant with a high predicted probability of being causal as there can be more than one causal variant in a locus (**Supplementary Table 14**). In addition to those already described above, the identified variants are strong candidate causal variants that merit prioritisation for future functional validation. For example, among the 87 variants, 10 are coding variants including several missense such as the *HBB* Glu7Val mentioned above, *GCKR* Leu446Pro, *RREB1* Asp1771Asn, *G6PC2* Pro324Ser, *GLP1R* Ala316Thr, and *TMPRSS6* Val736Ala, each of which have been proposed or shown to affect gene function^12,40–44^. We also additionally identify *AMPD3* Val311Leu (PPA=0.989) and *TMC6* Trp125Arg (PPA>0.999) variants associated with HbA1c which were previously detected in an exome array analysis but had not been fine-mapped with certainty due to the absence of backbone GWAS data^27^. Our current fine-mapping data now suggest these variants are likely to be causal and identify the cognate genes as the effector transcripts driving these associations.

Finally, we evaluated the resolution obtained in the trans-ancestry versus single-ancestry fine-mapping (**Methods, Supplementary Note**). To do this, we compared the number of variants in 99% CS across 98 locus-trait associations which, as suggested by FINEMAP, had a single causal variant in both trans-ancestry and single-ancestry analyses. Fine-mapping within and across ancestries was conducted using the same set of variants. At 8 of 98 locus-trait associations single-ancestry fine-mapping identified a single variant in the CS. In addition, at 72 of the 98 locus-trait associations, the number of variants in the 99% CS was smaller in trans-ancestry fine-mapping than in single-ancestry analyses (**Figure 3C**), which likely reflects the larger sample size and differences in LD structure, EAFs, and effect sizes across diverse populations. To quantify the estimated improvement in fine-mapping resolution attributable to the multi-ancestry GWAS, we then compared 99% CS sizes from the trans-ancestry fine-mapping to single-ancestry-specific data emulating the same total sample size by rescaling the standard errors (**Methods**). Of the 72 locus-trait associations with estimated improved fine-mapping in trans-ancestry analysis, resolution at 38 (53%) was improved because of the larger sample size in the trans-ancestry fine-mapping analysis (**Figure 3C**), and this estimated improved resolution would likely have been obtained in a European-only fine-mapping effort with equivalent sample size. However, at 34 (47%) loci, the inclusion of samples from multiple diverse populations yielded estimated improved resolution. On average, ancestry differences led to a reduction in the median number of variants in the 99% CS from 24 to 15 variants (37.5% median reduction; **Figure 3C**), demonstrating the value of conducting fine-mapping across ancestries.

### HbA1c Signal Classification

We, and others, have previously suggested that HbA1c-associated variants appear to exert their effects on HbA1c levels through both glycaemic and non-glycaemic pathways^7,45^ Classification of loci into these pathways can have important implications for T2D diagnostic accuracy^7,46^. To further elucidate the biology of HbA1c-associated variants, we took advantage of prior association results for other glycaemic, RBC, and iron traits, and used a fuzzy clustering approach to classify variants into their most likely mode of action (**Methods, Supplementary note**). Of the 202 autosomal HbA1c-associated trans-ancestry lead variants and single-ancestry index variants, 16 (8%) could not be characterized due to missing summary statistics in the other datasets and 17 (8%) could not be classified into a “known” class (**Supplementary note**). The remaining signals were classified as principally: a) glycaemic (n=51; 25%), b) affecting iron levels/metabolism (n=12; 6%), or c) RBC traits (n=106; 53%). We found a genetic risk score (GRS) composed of all HbA1c-associated signals was strongly associated with T2D risk (OR=2.5, 95% CI 2.5-2.6, *P*=2.4×10^-301^). However, when we tested partitioned GRSs composed of these different classes of variants (**Methods**), we found the T2D association was mainly driven by those variants influencing HbA1c through glycaemic pathways (OR=2.8, 95% CI 2.7-2.9, *P*=1.1×10^-251^), with weaker evidence of association (despite the larger number of variants in the GRS) and a more modest risk (OR=1.4, 95% CI 1.3-1.5, *P*=6.9×10^-4^) imparted by signals in the mature RBC cluster that were not glycaemic (i.e. where those specific variants had *P*>0.05 for FI, 2hGlu and FG) (**Supplementary Figure 15, Supplementary note**). This contrasts our previous finding where we found no significant association between a risk score of non-glycaemic variants and T2D^7^. Our current results could be partly driven by T2D cases being diagnosed based on HbA1c levels that may be influenced by the non-glycaemic signals, or by glycaemic effects not captured by FI, 2hGlu or FG measures.

### Biological signatures of glycaemic trait associated loci

To better understand distinct and shared biological signatures underlying variant-trait associations, we conducted genomic feature enrichment, eQTL co-localisation, and tissue and gene-set enrichment analyses across all four traits.

#### Epigenomic landscape of trait-associated variants

We next explored the genomic context underlying glycaemic trait loci by computing overlap enrichment for static annotations such as coding, conserved regions, histone modification ChIP-seq peaks, and super enhancers, merged across various cell types^47–49^ using the GREGOR tool^50^. We observed that FG, FI and HbA1c signals (**Supplementary Table 7**) were significantly (*P*<8.4×10^-4^, Bonferroni threshold correcting for 59 total annotations) enriched in evolutionarily conserved regions, whereas 2hGlu signals were only nominally enriched (**Fig 4A, Supplementary Figure 16, Supplementary Table 15**).

**Figure 4.**
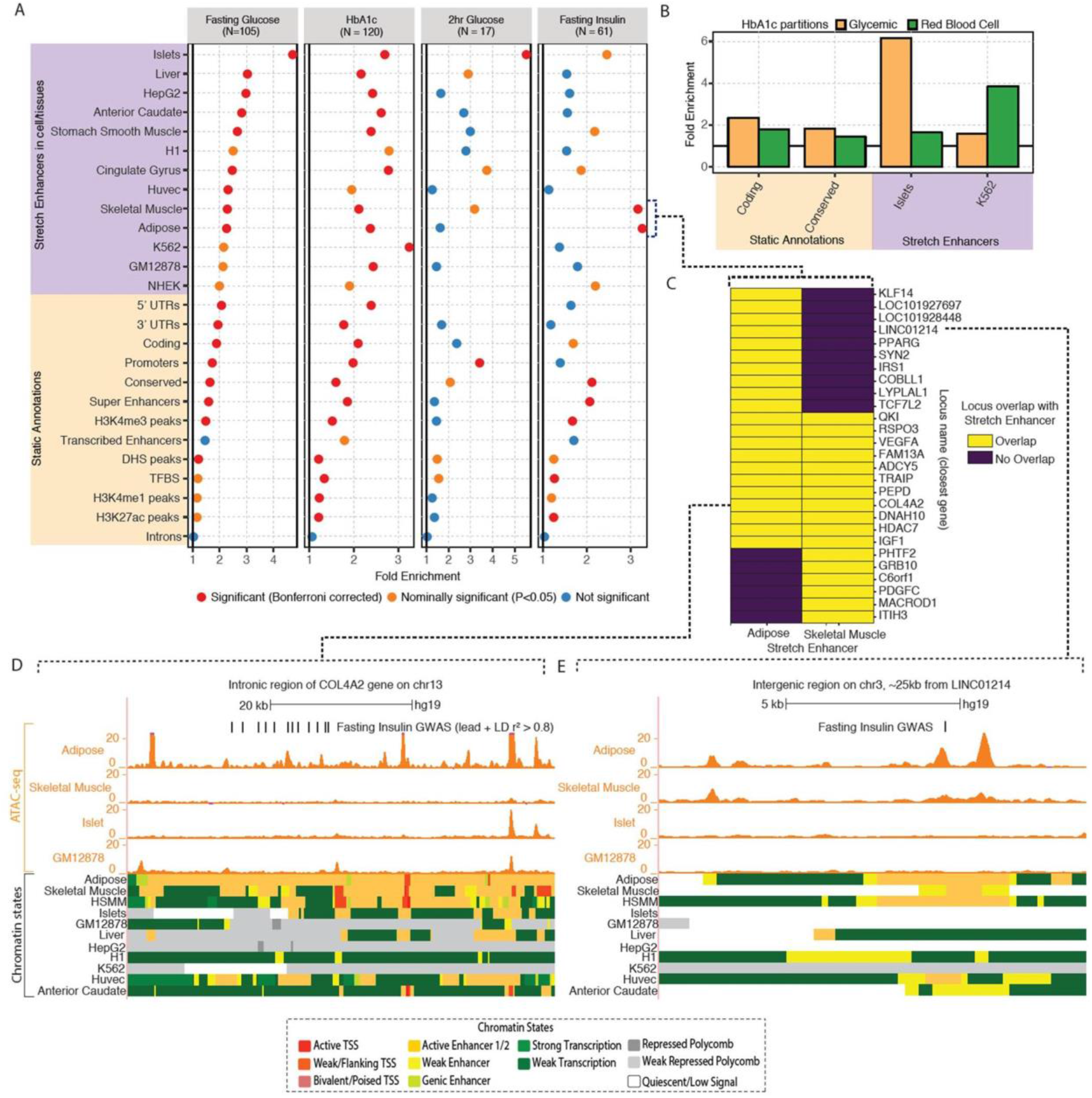
Epigenomic landscape of trait-associated variants. A: Enrichment of GWAS variants to overlap genomic regions including ‘Static Annotations’ which are common or ‘static’ across cell types and ‘Stretch Enhancers’ which are identified in each tissue/cell type. The numbers of signals for each trait are indicated in parentheses. Enrichment was calculated using GREGOR^50^. Significance (red) is determined after Bonferroni correction to account for 59 total annotations tested for each trait; nominal significance (*P*<0.05) is indicated in yellow. B: Enrichment for HbA1c GWAS signals partitioned into “hard” Glycaemic and Red Blood Cell cluster (signals from “hard” mature Red Blood Cell and reticulocyte clusters together) to overlap annotations including stretch enhancers in Islets and the blood-derived leukemia cell line K562, respectively (additional partitioned results in **Supplementary Table 17**). C: Individual FI GWAS signals that drive enrichment in Adipose and Skeletal Muscle stretch enhancers. D, E: Genome browser shots of FI GWAS signals – intronic region of the *COL4A2* gene (D) and an inter-genic region ~25kb from *LINC01214* gene (E) showing GWAS SNPs (lead and LD r^2^>0.8 proxies), ATAC-seq signal tracks and chromatin state annotations in different tissues/cell types.

We then focussed on the epigenomic landscapes defined in individual cell/tissue types. Previously, stretch enhancers (enhancer chromatin states ≥3kb in length) in pancreatic islets were shown to be highly cell-specific and strongly enriched with T2D risk signals^51^. We therefore calculated the enrichment of glycaemic trait-associated signals (**Supplementary Table 7**) in previously defined stretch enhancers^33^ across a diverse panel of cell types and tissues most relevant to the traits of interest: pancreatic islets, skeletal muscle, adipose, and liver (**Methods**). These analyses strongly suggest that variants associated with these glycaemic traits influence the function of tissue specific enhancers. Namely, FG- and 2hGlu-associated signals have the highest enrichment in islet stretch enhancers (FG: fold enrichment=4.70, *P*=2.7×10^-24^; 2hGlu: fold enrichment=5.51, *P*=3.6×10^-4^ **Figure 4A, Supplementary Table 16**), which highlights the relevance of pancreatic islet tissue for the regulation of FG and 2hGlu. Interestingly, FI-associated variants are strongly enriched for overlap with stretch enhancers in skeletal muscle (fold enrichment=3.17, *P*=7.8×10^-6^) and adipose tissue (fold enrichment=3.27, *P*=1.8×10^-7^), which is consistent with these tissues being key targets of insulin action and their involvement in the insulin resistance phenotype (**Figure 4A**). We note that the high enrichment of stretch enhancers in individual cell types (see upper “stretch enhancer” labelled portion of **Figure 4A**) as compared to super enhancers merged across cell types (see lower “static annotations” labelled portion of **Figure 4A**) highlights the importance of using cell-specific annotations in enrichment analyses. HbA1c-associated signals are enriched in stretch enhancers of multiple cell types and tissues likely because of the complex nature of this trait, but have the strongest enrichment in stretch enhancers from the blood-derived leukaemia cell line K562 (fold enrichment=3.24, *P*=1.21×10^-7^, **Figure 4A**). We next sought to identify potential cell specific epigenomic enrichments that are associated with the classified HbA1c-associated variants corresponding to the “hard” glycaemic and red blood cell clusters, the latter being the joint group of mature red blood cell and reticulocyte clusters. We found that these partitioned variants display expected cell type-specific enrichment trends with the HbA1c glycaemic variants significantly enriched in islet stretch enhancers (fold enrichment=6.25, *P*=4.02×10^-10^ **Figure 4B, Supplementary Table 17**) and not in K562. Conversely, the HbA1c red blood cell variants are significantly enriched in K562 stretch enhancers (fold enrichment=3.85, *P*=3.32×10^-8^, **Figure 4B, Supplementary Table 17**) and not in islets.

To complement the overlap enrichment results from GREGOR, we also computed enrichment with two additional approaches: fGWAS^52^ and GARFIELD^53^. These independent analyses yielded consistent results (**Supplementary Figures 17-18, Supplementary Tables 15 and 18**), demonstrating reproducibility across different approaches.

Given the observed enrichment of FI loci with stretch enhancers from adipose and skeletal muscle tissue, we sought to explore these loci in more detail. We found that 11 of the 27 loci driving these enrichment signals include variants that overlap stretch enhancers in both adipose and skeletal muscle (**Figure 4C**). At the *COL4A2* locus, variants within an intronic region of the gene overlap stretch enhancer chromatin states in adipose tissue, skeletal muscle, and a human skeletal muscle myoblast (HSMM) cell line that are not shared across other cell types and tissues; among these variants, rs9555695 (in the 99% CS) also overlaps accessible chromatin regions in adipose (**Figure 4D**). At a narrow signal (no proxy variants with LD r^2^>0.7 in Europeans, for the lead trans-ancestry rs62271373 variant), rs62271373 (PPA = 0.94) located in an intergenic region ~25kb from the *LINC01214* gene overlaps stretch enhancer chromatin states in adipose and HSMM and active enhancer chromatin states in skeletal muscle, but does not overlap any enhancer states in other tissues (**Figure 4E**). The lead rs62271373 variant also overlaps an ATAC-seq peak in adipose tissue. Collectively, the tissue-specific stretch enhancer epigenomic signatures at GWAS signals provide an opportunity to nominate tissues where these variants are likely to be active. Such a map will be helpful in future efforts to deconvolute GWAS signals into tissue-specific disease pathology.

#### Co-localisation of GWAS and eQTLs

Among the 99 novel glycaemic trait loci identified by this study, we identified co-localised eQTLs at 34 loci in blood, pancreatic islets, subcutaneous or visceral adipose, skeletal muscle, or liver, providing suggestive evidence of causal genes (**Supplementary Table 19**). The co-localised eQTLs include several genes previously reported at glycaemic trait loci: *ADCY5, CAMK1D, IRS1, JAZF1*, and *KLF14*^54–56^. For some additional loci, the co-localised genes have prior evidence for a role in glycaemic regulation. For example, the lead trans-ancestry variant and likely causal variant, rs1799815 (PPA=0.993, mentioned above), associated with FI is the strongest variant associated with expression of *INSR*, encoding the insulin receptor, in subcutaneous adipose from METSIM (*P*=2×10^-9^) and GTEx (*P*=5×10^-6^). The A allele at rs1799815 is associated with higher FI and lower expression of *INSR*, which is consistent with the well-established relationship in humans and model organisms between insulin resistance and reduced function of INSR protein^57^. In a second example, rs841572, the trans-ancestry lead variant associated with FG, is the variant with the highest PPA (PPA=0.535) among the 20 variants in the 99% CS and is in strong LD (r^2^=0.87) with the lead eQTL variant (rs841576, also in the 99% CS) associated with expression of *SLC2A1* in blood from eQTLGen (*P*=1×10^-8^). *SLC2A1*, also known as *GLUT1*, encodes the major glucose transporter in brain, placenta, and erythrocytes, and is responsible for glucose entry into the brain^58^. The A allele at rs841572 is associated with lower FG and lower *SLC2A1* expression. While rare missense variants in *SLC2A1* are an established cause of seizures and epilepsy^59^, our data suggest that *SLC2A1* variants also affect plasma glucose levels within a healthy physiological range. At both loci, the novel associations and co-localised eQTLs provide strong human genetic support for early glycaemia candidate genes.

The co-localised eQTLs also provide new insights into the mechanisms at glycaemic trait loci. For example, rs9884482 (a variant in the 99% CS) is associated with FI and expression of *TET2* in subcutaneous adipose (*P*=2×10^-20^); rs9884482 is in high LD (r^2^=0.96 in Europeans) with the lead *TET2* eQTL variant (rs974801). *TET2* encodes a DNA-demethylase through which *TET2* can affect transcriptional repression^60^. Adipose Tet2 expression is reduced in diet-induced insulin resistance in mice^61^, and knockdown of Tet2 blocked adipogenesis by repressing *Pparg* expression^61,62^. Consistently, in human adipose tissue, rs9884482-C was associated with lower expression of *TET2* and higher FI. In a second example, HbA1c-associated variant rs617948 (a variant in the 99% CS) is the lead variant associated with expression of *C2CD2L* in blood from eQTLGen (*P*=3×10^-96^). *C2CD2L*, also known as *TMEM24*, has been shown to regulate pulsatile insulin secretion and facilitate release of insulin pool reserves^63,64^. The G allele at rs617948 was associated with higher HbA1c and lower *C2CD2L*, providing evidence for a role of this insulin secretion protein in glucose homeostasis. Our HbA1c “soft” clustering classification assigns this signal to both the “unknown” (0.51 probability) and “reticulocyte” (0.42 probability) clusters, and this variant has no evidence for association with FG, FI or 2hGlu (*P*>0.05), but is strongly associated with HbA1c (*P*<6.8×10^-8^), reticulocytes (RET; P<5×10^-7^) and HbA1c adjusted for FG (*P*<6.12×10^-7^; **Supplementary Table 20, Supplementary Note**). Together, these results would suggest a possible effect of this variant on reticulocyte biology, and an effect on insulin secretion (mediated through *C2CD2L*) which is not captured by any of our traits, both of which potentially influencing HbA1c levels through different tissues, and providing a plausible explanation for the classification as “unknown”.

#### Tissue Expression

Consistent with results based on effector transcripts and expression analysis based on GTEx data^27^, we found significant differences in tissue expression across the glycaemic trait-associated variants. FG-associated variants were enriched for genes expressed in the pancreas (at FDR<0.05), while there was insufficient power (insufficient number of genome-wide significant associations) in 2hGlu analysis to identify enrichment for any tissues or cell types at a more relaxed FDR<0.2 threshold. FI-associated variants were enriched for connective tissue and cells (which includes adipose tissue), endocrine glands, blood cells, and muscles (at FDR<0.2) and HbA1c-associated variants were significantly enriched for genes expressed in the pancreas, hemic, and immune system (at FDR<0.05) (**Figure 5, Supplementary Table 21**). Consistent with our previous analysis^27^, FI-enrichment for connective tissue was driven by adipose tissue (subcutaneous and visceral), while the newly described enrichment with endocrine glands was driven by the adrenal glands and cortex (**Supplementary Table 21**). Beyond enrichment for genes expressed in glycaemic-related tissues, the association of HbA1c-associated variants with genes expressed in blood is consistent with the role of RBC in this glycaemic measure and our previous results^27^.

**Figure 5.**
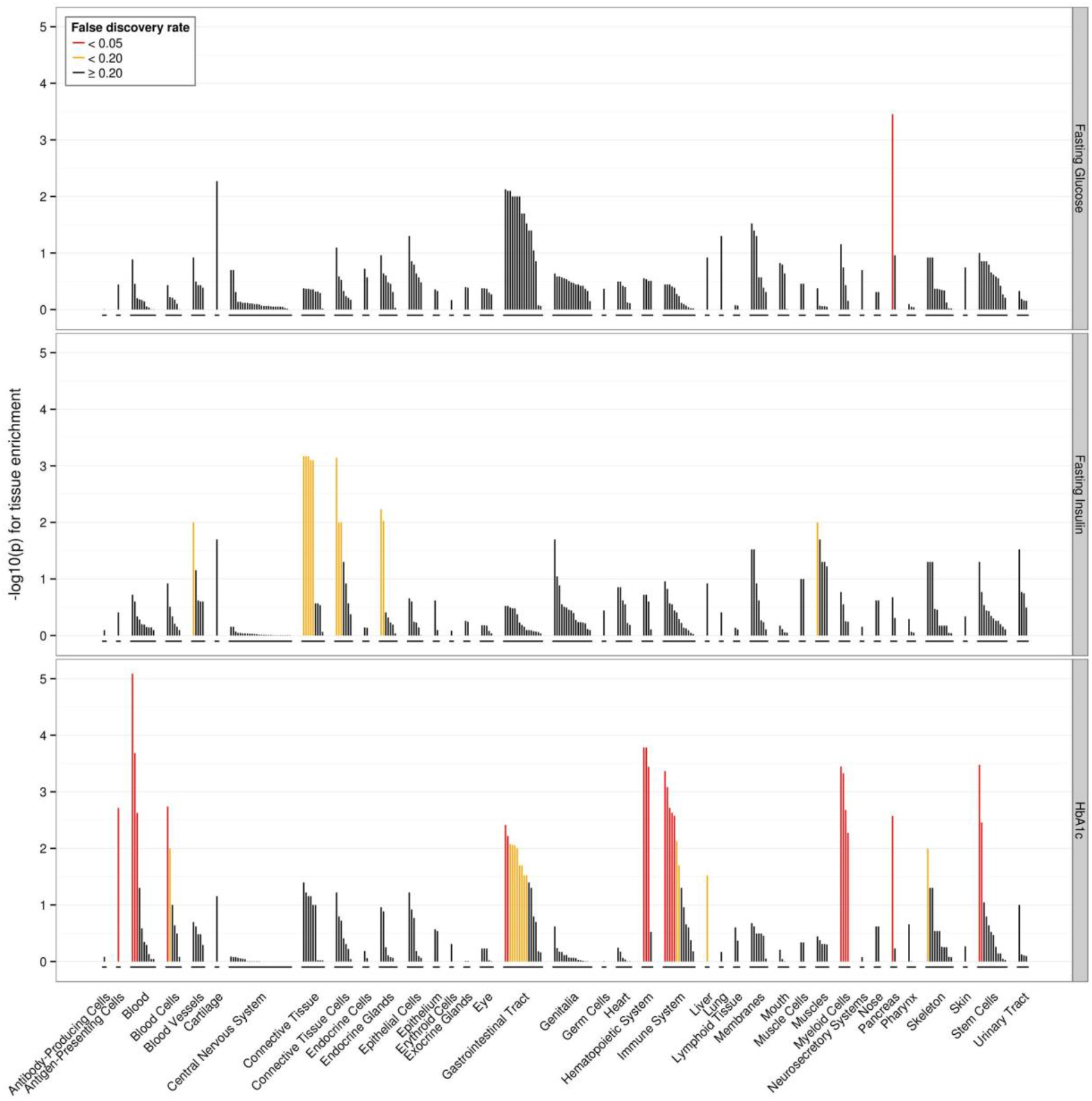
Tissues and cell types significantly enriched for genes within glycaemic-associated loci. Top panel FG-associated loci, middle panel FI-associated loci, bottom panel Hba1c-associated loci. FRD thresholds are shown in red (q<0.05), orange (q<0.2), grey (q≥0.2).

The association between FI-associated variants and genes expressed in adrenal glands is notable. Incidental adrenal masses (those detected through routine use of radiological imaging but for which patients have not yet shown signs of adrenal hormone excess) have often been associated with hypertension, dyslipidaemia, glucose intolerance, and obesity, all hallmarks of insulin resistance^65^. However, it has not been clear whether incidental adrenal masses are a cause or consequence of the associated insulin resistance^66,67^. Our results would suggest that FI-associated variants (a surrogate for insulin resistance) are enriched in genes expressed in the adrenal glands, suggesting a possible direct role for these in insulin resistance. One hypothesis is that these genes might influence cortisol levels, which could subsequently contribute to insulin resistance and FI levels through impairment of the insulin receptor signalling pathway in peripheral tissues, as well as influencing body fat distribution, stimulate lipolysis, and other indirect mechanisms^67,68^.

#### Gene-set Analyses

Next, we performed gene-set analysis using DEPICT (**Methods**). In keeping with previous results^27^, we found distinct gene-sets enriched (FDR<0.05) for each glycaemic trait (except 2hGlu, for which genome-wide associations were insufficient to have power in this analysis). FG-associated variants highlighted gene-sets involved in metabolism in addition to gene-sets involved in more general cellular function such as “cytoplasmic vesicle membrane” and “circadian clock”” (**Figure 6A**). In contrast, in addition to metabolism related gene-sets FI-associated variants highlighted pathways related to growth, cancer and reproduction (**Figure 6B**). This is consistent with the role of insulin as a mitogenic hormone, and with epidemiological links between insulin and certain types of cancer^69^ and reproductive disorders such as polycystic ovary syndrome^70^. HbA1c-associated variants highlighted a wide network of gene-sets (**Figure 6C**), including those linked to metabolism, as well as those linked to haematopoiesis, again recapitulating our postulated effects of variants on glucose and RBC biology. Additional pathways highlighted from HbA1c-associated variants also highlighted previous “CREBP PPi” and lipid biology related to T2D^71^ and HbA1c^72^, respectively, and potential new biology through which variants may influence HbA1c.

**Figure 6.**
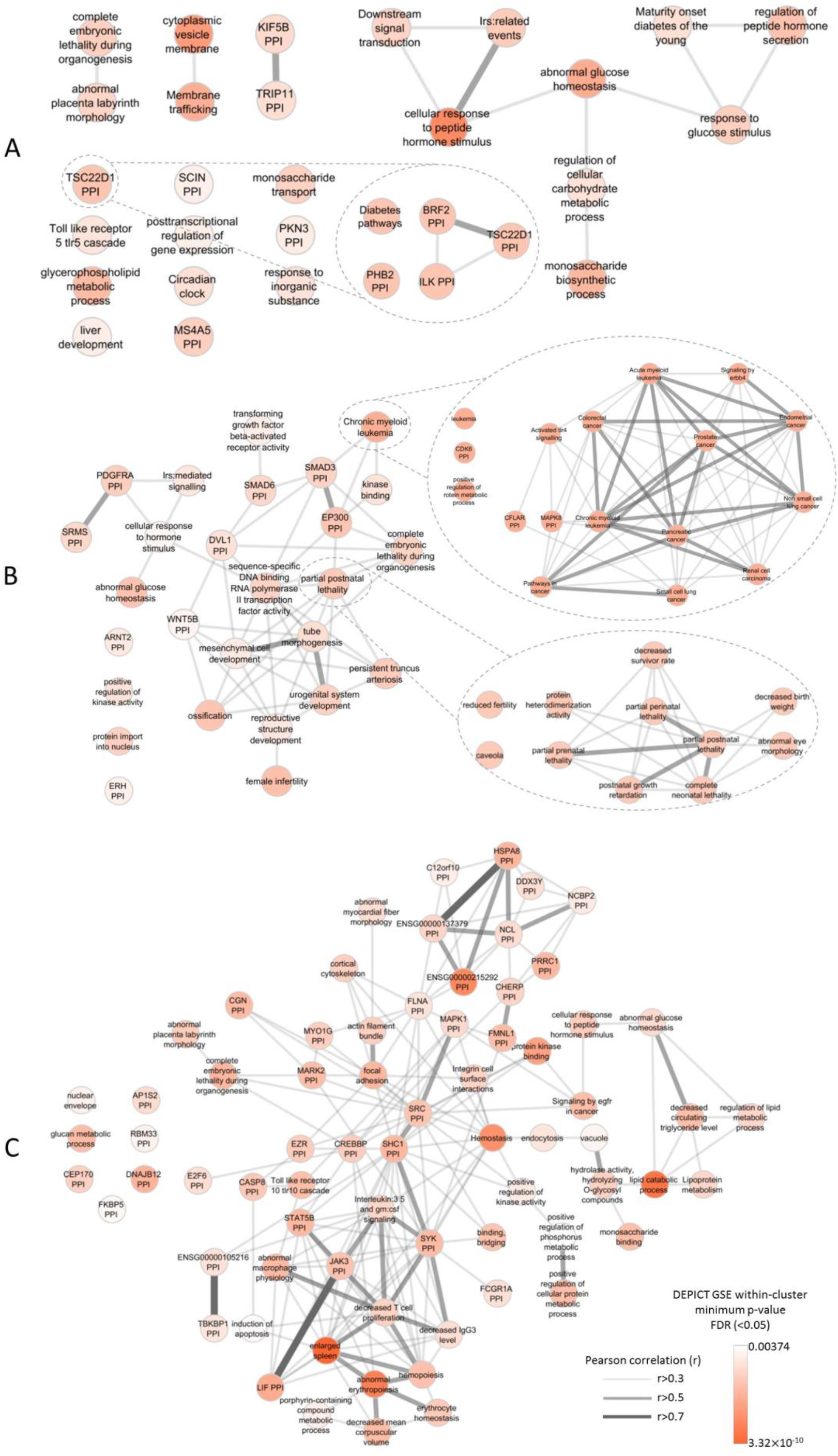
Gene-set enrichment analyses. Results from affinity-propagation clustering of significantly enriched gene sets (FDR<0.05) identified by DEPICT for A) FG, B) FI, and C) HbA1c. Each node is a cluster of gene-sets represented by an exemplar gene-set with similarities between the clusters represented by the Pearson correlation coefficients (r>0.3). The nodes are coloured according to the minimum gene-set enrichment p-value for gene-sets in that cluster. Example clusters are expanded to show the contributing gene-sets.

## Discussion

Here we describe a large meta-analysis of GWAS of glycaemic traits for which 30% of the population was composed of East Asian, Hispanic, African-American, South Asian and sub-Saharan African participants, in addition to the European ancestry participants. Overall, this effort identified 242 loci (235 trans-ancestry and seven single-ancestry), which jointly explain between 0.7% (2hGlu in European ancestry individuals, SE=0.85% for 2hGlu) and 6% (HbA1c in African American ancestry, SE=1.2% for HbA1c) of the variance in glycaemic traits in any given ancestry.

A key aim of our study was to evaluate the added advantage of including population diversity into genetic discovery and fine-mapping efforts. Beyond the overall larger sample size included in the trans-ancestry meta-analysis, we were able to estimate the contribution of non-European ancestry data in locus discovery and fine-mapping resolution. We found that 24 of the 99 newly discovered loci owe their discovery to the inclusion of East Asian, Hispanic, African-American, South Asian and sub-Saharan African participant data, due to differences in EAF and effect sizes across ancestries.

Comparison of 295 trans-ancestry lead variants (315 locus-trait associations) across ancestries demonstrated that between 81.5% (for HbA1c) and 85.7% (for FG) of the trans-ancestry lead variants had no evidence of trans-ancestry heterogeneity in allelic effects (*P*>0.05). Expanded analyses including variants across the whole genome, demonstrated at least nominal concordance in the direction of effects between populations of European ancestry and other ancestries for all but the least significant association signals observed in European ancestry GWAS. These observations are consistent with a tail of variants with modest but homogenous effects on glycaemic traits across ancestries that would be amenable to discovery with even larger sample sizes in trans-ancestry meta-analysis.

Recently, there has been ongoing debate regarding the utility of GS for trait prediction across different studies, and in particular, across diverse populations^73–76^. Here, we attempted to more precisely estimate the utility of weighted GS constructed from European ancestry effect sizes, to predict trait variance explained in studies independent from those used in genetic discovery, and across ancestries. We show that GSs built from index variants in Europeans (P<5×10^-8^) explain similar proportions of the trait variance across populations, though much of the trait variance remains unexplained even in European ancestry individuals. Also, these European participant-based scores will fail to detect ancestry-specific variant effects which can have large effect sizes on specific traits^7^. Consequently, when these analyses are extended to variants with weaker associations, we observe that while (as expected) variance explained is improved in European ancestry participants performance is worsened into other ancestries. We suggest that for the less stringent thresholds, the association signals will be less “peaked” in European ancestry GWAS. As a result, the SNP selected for the GS is less likely to be causal, meaning that differences in LD structure compared with other ancestry groups adds noise to the prediction.

We further demonstrate that fine-mapping resolution is improved in trans-ancestry, compared to single-ancestry fine-mapping efforts. In ~50% of our loci, we were able to demonstrate the improvement is due to differences in EAF, effect size, or LD structure between ancestries, and not just due to the overall increased sample size available for trans-ancestry fine-mapping. By performing trans-ancestry fine-mapping, and co-localising GWAS signals with eQTL signals and coding variants, we identify new candidate causal genes. Altogether, these results provide additional strong motivation for continued expansion of genetic and genomic efforts in diverse populations, not least to improve understanding of these traits in diverse ancestries in whom individuals are often disproportionally affected by T2D.

Given data on four different glycaemic traits, and their utility to diagnose and monitor T2D and metabolic health, we also sought to characterise biological features underlying these traits. We show that despite significant sharing of genetic loci across the four glycaemic traits, each trait is also characterised by a unique set of features based on stretch enhancer, gene expression and gene-set signatures. Combining genetic data from these traits with T2D data will further elucidate pathways driving normal physiology and pathophysiology, and help further develop useful predictive scores for disease classification and management^4,5^

## Acknowledgments

A full list of acknowledgments and funding appears in the **Supplementary Note**.

## Methods

### Study design and participants

This study included trait data from four glycaemic traits: fasting glucose (FG), fasting insulin (FI), 2hr post-challenge glucose (2hGlu), and glycated haemoglobin (HbA1c). The total number of contributing cohorts ranged from 41 (2hGlu) to 131 (FG), and the maximum sample size for each trait ranged from 85,916 (2hGlu) to 281,416 (FG) (**Supplementary Table 1**). Overall, European ancestry (EUR) participants dominated the sample size for all traits, representing between 68.0% (HbA1c) to 73.8% (2hGlu) of the overall sample size. African Americans (AA) represented between 1.7% (2hGlu) to 5.9% (FG) of participants; individuals of Hispanic ancestry (HISP) represented between 6.8% (FG) to 14.6% (2hGlu) of participants; individuals of East-Asian ancestry (EAS) represented between 9.9% (2hGlu) to 15.4% (HbA1c) of participants; and South-Asian ancestry (SAS) individuals represented between 0% (no contribution to 2hGlu) to 4.4% (HbA1c) of participants. Data from Ugandan participants were only available for the HbA1c analysis and represented 2% of participants.

### Phenotypes

Analyses included data for FG and 2hGlu measured in mmol/l, FI measured in pmol/l, and HbA1c in % [where possible, studies reported HbA1c as a National Glycohemoglobin Standardization Program (NGSP) percent]. Similar to previous MAGIC efforts^7^, individuals were excluded if they had type 1 or type 2 diabetes (defined by physician diagnosis); reported use of diabetes-relevant medication(s); or had a FG ≥7 mmol/L, 2hGlu ≥11.1mmol/L, or HbA1c ≥ 6.5%, as detailed in **Supplementary Table 1**. 2hGlu measures were obtained 120 minutes after a glucose challenge in an oral glucose tolerance test (OGTT). Measures for FG and FI taken from whole blood were corrected to plasma level using the correction factor 1.13^77^.

### Genotyping, quality control, and imputation

Each participating cohort performed study-level quality control, imputation, and association analyses following a shared analysis plan. Cohorts were genotyped using commercially available genome-wide arrays or the Illumina CardioMetabochip (Metabochip) array (**Supplementary Table 1**)^78^. Prior to imputation, each cohort performed stringent sample and variant quality control (QC) to ensure only high-quality variants were kept in the genotype scaffold for imputation. Sample quality control checks included removing samples with low call rate < 95%, extreme heterozygosity, sex mismatch with X chromosome variants, duplicates, first- or second-degree relatives (unless by design), or ancestry outliers. Following sample QC, cohorts applied variant QC thresholds for call rate (< 95%), Hardy-Weinberg Equilibrium (HWE) *P* < 1×10^-6^, and minor allele frequency (MAF). Full details of QC thresholds and exclusions by participating cohort are available in **Supplementary Table 1**.

Imputation was performed up to the 1000 Genomes Project phase 1 (v3) cosmopolitan reference panel^79^, with a small number of cohorts imputing up to the 1000 Genomes phase 3 panel^19^ or population-specific reference panels (**Supplementary Table 1**).

### Study level association analyses

Each of the glycaemic traits (FG, natural log FI, and 2hGlu) were regressed on BMI (except HbA1c), study-specific covariates, and principal components (unless implementing a linear mixed model). Analyses for FG, FI, and 2hGlu were adjusted for BMI as we had previously shown this did not materially affect results for FG and 2hGlu but improved our ability to detect FI-associated loci^15^. For simplicity, we refer to the traits as FG, FI and 2hGlu. For a discussion on collider bias see **Supplementary Note section 2c**. Both the raw and rank-based inverse normal transformed residuals from the regression were tested for association with genetic variants using SNPTEST^23^ or Mach2Qtl^80,81^. Poorly imputed variants, defined as imputation r^2^ < 0.4 or INFO score < 0.4, were excluded from downstream analyses (**Supplementary Table 1**). Following study level QC, approximately 12,229,036 variants (GWAS cohorts) and 1,999,204 variants (Metabochip cohorts) were available for analysis (**Supplementary Table 1**).

### Centralised quality control

Each contributing cohort shared their summary statistic results with the central analysis group who performed additional QC using EasyQC^82^. Allele frequency estimates were compared to estimates from 1000Gp1 reference panel^79^, and variants were excluded from downstream analyses if there was a minor allele frequency difference > 0.2 for AA, EUR, HISP, and EAS populations against AFR, EUR, MXL, and ASN populations from 1000 Genomes Phase 1, respectively, or a minor allele frequency difference > 0.4 for SAS against EUR populations. At this stage, additional variants were excluded from each cohort file if they met one of the following criteria: were tri-allelic; had a minor allele count (MAC) < 3; demonstrated a standard error of the effect size ≥ 10; or were missing an effect estimate, standard error, or imputation quality. All data that survived QC (approximately 12,186,053 variants from GWAS cohorts and 1,998,657 variants from Metabochip cohorts) were available for downstream meta-analyses.

### Single-ancestry meta-analyses

Single-ancestry meta-analyses were performed within each ancestry group using the fixed-effects inverse variance meta-analysis implemented in METAL^20^. We applied a double-genomic control (GC) correction^15,83^ to both the study-specific GWAS results and the single-ancestry meta-analysis results. Study-specific Metabochip results were GC-corrected using 4,973 SNPs included on the Metabochip array for replication of associations with QT-interval, a phenotype not correlated with our glycaemic traits^15^.

### Identification of single-ancestry index variants

To identify distinct association index variants across each chromosome within each ancestry (**Glossary box**), we performed approximate conditional analyses implemented in GCTA^21^ using the -- cojo-slct option (autosomes) and distance-based clumping (X chromosome). Linkage disequilibrium (LD) correlations for GCTA were estimated from a representative cohort from each ancestry: WGHS (EUR); CHNS (EAS); SINDI (SAS); BioMe (AA); SOL (HISP) and Uganda (for itself). The results from GCTA were comparable when using alternative cohorts for the LD reference. For any index variant with a QC flag which caused reason for concern, we performed manual inspection of forest plots to decide whether the signal was likely to be real (**Supplementary note**). Among 335 single-ancestry index variants across all traits, this manual inspection was done for 40 signals of which 32 passed and 8 failed after inspection. Thus, a total of 327 single-ancestry index variants passed and 8 failed.

### Trans-ancestry meta-analyses

To leverage power across all ancestries, we also conducted trait-specific trans-ancestry meta-analysis by combining the single-ancestry meta-analysis results using MANTRA (**Supplementary Figure 3**)^84^.We defined log_10_Bayes’ Factor (BF) > 6 as genome-wide significant, approximately comparable to *P* < 5×10^-8^.

### Manual curation of trans-ancestry lead variants

To ensure trans-ancestry lead variants were robust, we performed manual inspection of forest plots by at least two authors, for any variants with flags indicating possible QC issues (**Supplementary Note**). Of 463 trans-ancestry lead variants across all traits, 184 passed without inspection, 131 passed after inspection, and 148 failed after inspection.

### Correlation in EAF and heterogeneity in effect sizes of TA lead variants across ancestries

For each pair of ancestries, we calculated Pearson’s correlation in EAFs for each trans-ancestry lead variant. The pairwise summarised heterogeneity of effect sizes between ancestries was then tested using the joint F-test of heterogeneity^29^. The test statistic is the sum of Cochran Q-statistics for heterogeneity across all trans-ancestry signals. Under the null hypothesis, the statistics follows the χ^2^ distribution with n degrees of freedom, where n is the number of the trans-ancestry lead variants.

### Concordance analyses of LD pruned European single-ancestry index variants into other ancestries

We compared the direction of effect of variants on each trait separately. For each trait, we identified variants reported in the European ancestry meta-analysis and each non-European ancestry meta-analysis, in turn. These variants were assigned to *P*-value bins, according to the strength of the association with the trait in the European ancestry meta-analysis: *P* < 5×10^-8^; 5×10^-8^ ≤*P* <5×10^-6^; 5×10^-6^ ≤*P* <5×10^-4^; 5×10^-4^ ≤*P* <0.05; and *P* ≥ 0.05. Within each *P*-value bin, we selected a set of “independent” variants that were separated by 1 Mb. We defined independence using a distance-based threshold because of differences in patterns of LD between ancestry groups. For each *P*-value bin, the proportion of variants with the same direction of effect on the trait between the two ancestries was calculated along with a *P*-value from the binomial test to determine if the proportion of variants with the same direction of effect was greater than that expected by chance (50%, one sided).

### LD-pruned variant lists

Several downstream analyses (for example, genomic feature enrichment, genetic scores, and estimation of variance explained by associated variants) require independent LD-pruned variants (r^2^ < 0.1) to avoid double-counting variants which might otherwise be in LD with each other and that do not provide additional “independent” evidence. Therefore, for these analyses we generated different lists of either TA or single-ancestry LD pruned (r^2^ < 0.1) variants, keeping in each case the variant with the strongest evidence of association (**Supplementary Table 7**). Subsequently, we combined TA and single-ancestry variant lists and conducted further LD pruning. For some analyses, we took the TA pruned variant list and added single-ancestry signals if the LD r^2^ < 0.1, while for others we started with the single-ancestry pruned lists and supplemented with TA lead variants if the LD r^2^ < 0.1. One exception was the list used for eQTL co-localisations, which included all single-ancestry European signals (without LD pruning) and supplemented with any additional TA lead variants (starting from the variants with the most significant P-values) in EUR LD r^2^ <0.1 with any of the variants already in list, and that reached at least *P* < 1×10^-5^ in the European ancestry meta-analysis.

### Transferability of genetic scores (GSs) across ancestries

To determine the power of a European-based genetic score (GS) to predict trait values within non-European populations, we began with the list of European LD-pruned index variants (**Supplementary Table 7**) and their effect sizes. We first tested the GS in four European cohorts with individual level data that did not contribute to this meta-analysis: WHITEHALL II, Oxford Biobank, VIKING and UKHLS (**Supplementary Table 1**). We used individual level genotype data to build an effect-size weighted GS for each individual, and then obtained the trait variance explained via linear regression. We then tested the European GS in each European ancestry cohort contributing to the meta-analysis with > 1,000 samples by: (i) adjusting the effect size estimate of each variant to remove the contribution of the cohort^30^; and (ii) obtaining the proportion of the trait explained by the GS (R^2^) on the basis of cohort-level summary statistics using the R package “gtx”. Finally, we obtained the proportion of the trait explained by the European GS in other ancestry groups on the basis of single-ancestry meta-analysis summary statistics using the R package “gtx”. Variants reported in < 50% of the total sample size in each ancestry group for each trait were excluded as they can yield unstable estimates of R^2^. Standard errors of effect size estimates were adjusted to account for differences in the sample size reported for each variant. Additional sensitivity analyses were also performed using: (i) variants that exhibited only modest evidence of heterogeneity (*P* > 1×10^-6^) between ancestries from the trans-ancestry analysis; (ii) variants with no evidence of statistically significant between-ancestry heterogeneity (*P* > 0.05) (iii) variants with *P* < 1×10^-5^ in the European meta-analysis (1Mb distance clumped); and (iv) variants with *P* < 0.05 in the European meta-analysis (1Mb distance clumped).

### Trait variance explained

To determine how much of the phenotypic variance of each trait could be explained by the corresponding trait-associated loci, variants were combined in a series of weighted GS. The analysis was performed in a subset of the cohorts included in the discovery GWAS (with representation from each ancestry) and in a smaller number of independent cohorts (European ancestry only). Up to three GS were generated per trait (and for each ancestry), representing single-ancestry signals, single-ancestry signals plus trans-ancestry signals, and trans-ancestry signals plus single-ancestry signals (**Supplementary Table 7**). In the case of the European ancestry cohorts that contributed to the GWAS, we employed the method of Nolte *et al*.^30^ to adjust the effect sizes (betas) from the GWAS for the contribution of that cohort, providing sets of cohort-specific effect sizes that were then used to generate the GS. The association between each GS and its corresponding trait was tested by linear regression and the adjusted R^2^ from the model extracted as an estimate of the variance explained.

### Fine-mapping

Of the 242 loci identified in this study, 237 were autosomal loci which we took forward for fine-mapping (**Supplementary Table 2**). We used the Bayesian fine-mapping method FINEMAP^85^ (version 1.1) to refine association signals and attempt to identify likely causal variants at each locus. FINEMAP estimates the maximum number of causal variants at each locus, calculates the posterior probability of each variant being causal, and proposes the most likely configuration of causal variants. The posterior probabilities of the configurations in each locus were used to construct 99% credible sets.

We performed both single-ancestry and trans-ancestry fine-mapping. In both analyses, only data from cohorts genotyped on GWAS arrays were used, and analyses were limited to trans-ancestry lead variants and other single-ancestry lead variants present in at least 90% of the samples for each trait. For the single-ancestry fine-mapping, FINEMAP estimates the number of causal variants in a region up to a maximum number, which we set to be two plus the number of distinct signals identified from the GCTA signal selection. FINEMAP uses single-ancestry and trait-specific z-scores from the fixed-effect meta-analysis in METAL^20^ and an ancestry-specific LD reference, which we created from a subset of cohorts (combined sample size > 30% of the sample size for that ancestry), weighting each cohort by sample size. In the trans-ancestry fine-mapping, FINEMAP was similarly used to estimate the number of causal variants starting with two, and trait-specific z-scores and LD maps were generated from the sample size weighted average of those used in the single-ancestry fine-mapping. The maximum number of causal variants was iteratively increased by one until it was larger than the number of causal variants supported by data (Bayes factor), which was the estimated maximum number of causal variants used in the final run of fine-mapping analysis.

To compare fine-mapping results obtained from the single-ancestry and trans-ancestry efforts, analyses were limited to fine-mapping regions with evidence for a single likely causal variant in both, enabling a straightforward comparison of credible sets (**Supplementary note**). To ensure any difference in the fine-mapping results was not driven by different sets of variants being present in the different analyses, we repeated the single-ancestry fine-mapping limited to the same set of variants used in the trans-ancestry fine-mapping. The fine-mapping resolution was assessed based on comparisons of the 99% credible sets in terms of number of variants included in the set, and length of the region. To assess whether the improvement in the trans-ancestry fine-mapping was due to differences in LD, increased sample size, or both, we repeated the trans-ancestry fine-mapping mimicking the sample size present in the single-ancestry fine-mapping by dividing the standard errors by the square root of the sample size ratio and compared the results with those from the single-ancestry fine-mapping.

### Functional Annotation of trait-associated variants

#### HbA1c signal classification

There were 202 autosomal HbA1c-associated signals from either the single-ancestry (i.e. all GCTA-signals from any ancestry) or trans-ancestry meta-analyses. To classify these signals in terms of their likely mode of action (i.e., glycaemic, erythrocytic, or other^7^), we examined association summary statistics for the lead variants at the 202 signals in other large European datasets for 19 additional traits: three glycaemic traits from this study (FG, 2hGlu and FI); seven mature red blood cell (RBC) traits^86^ (red blood cell count, mean corpuscular volume, haematocrit, mean corpuscular haemoglobin, mean corpuscular haemoglobin concentration, haemoglobin concentration and red cell distribution width); five reticulocyte traits (reticulocyte count, reticulocyte fraction of red cells, immature fraction of reticulocytes, high light scatter reticulocyte count and high light scatter percentage of red cells)^86^, and four iron traits (serum iron, transferrin, transferrin saturation and ferritin)^87^. Of the 202 autosomal HbA1c signals, data were available for the lead (n=177) or proxy (European LD r^2^ > 0.8, n = 9) variants at 186 signals.

The additional traits were clustered using hierarchical clustering to ensure biologically related traits would cluster together (**Supplementary note**). We then used a non-negative matrix factorization (NMF)^88^ process to cluster the HbA1c signals. Each cluster was labelled as glycaemic, reticulocyte, mature RBC, or iron related based on the strength of association of signals in the cluster to the glycaemic, reticulocyte, mature RBC and iron traits (**Supplementary note**). To verify that our cluster naming was correct, we used HbA1c association results conditioned on either FG or iron traits, or type 2 diabetes association results (**Supplementary note**).

#### HbA1c genetic risk scores (GRSs) and type 2 diabetes (T2D) risk

We constructed GRS for each cluster of HbA1c-associated signals (based on hard clustering) and tested the association of each cluster with T2D risk using samples from the UK Biobank. Pairs of HbA1c signals in LD (EUR r^2^ > 0.10) were LD pruned by removing the signal with the less significant *P*-value of association with HbA1c. The GRS for each cluster was calculated based on the logarithm of odds ratios from the latest T2D study summary statistics^89^ and UK Biobank genotypes imputed to the Haplotype Reference Consortium^19^. From 487,409 UK Biobank samples, we excluded participants for the following reasons: 373 with mismatched sex; 9 not used in the kinship calculation; 78,365 non-European ancestry individuals; and 138,504 with missing T2D status, age, or sex information. We further removed 26,896 related participants (kinship > 0.088, preferentially removing individuals with the largest number of relatives and controls where a T2D case was related to a control). T2D cases were defined by: (i) a history of diabetes without metformin or insulin treatment, (ii) self-reported diagnosis of T2D, or (iii) diagnosis of T2D in a national registry (N = 17,022). Controls were participants without a history of T2D (N = 226,240). We tested for association between each GRS and T2D using logistic regression including covariates for age, sex, and the first five principle components. Significance of association was evaluated by a bootstrap approach to incorporate the variance of each HbA1c associated signal in the T2D summary data. To do this, we generated the GRS of each cluster 200 times by resampling the logarithm of odds ratio of each signal with T2D. For each non-glycaemic class that had a GRS significantly associated with T2D, we performed sensitivity analyses to evaluate whether the association was driven from variants that also belonged to a glycaemic cluster when using a soft clustering approach (the signals were classified as also glycaemic in the soft clustering or had an association *P* ≤ 0.05 with any of the three glycaemic traits).

#### Chromatin states

To identify genetic variants within association signals that overlapped predicted chromatin states, we used a previously published, 13 chromatin state model that included 31 diverse tissues, including pancreatic islets, skeletal muscle, adipose, and liver^33^. Briefly, this model was generated from cell/tissue ChIP-seq data for H3K27ac, H3K27me3, H3K36me3, H3K4me1, and H3K4me3, and input control from a diverse set of publicly available data^47,51,90,91^ using the ChromHMM program^92^. As reported previously^33^, stretch enhancers were defined as contiguous enhancer chromatin state (Active Enhancer 1 and 2, Genic Enhancer and Weak Enhancer) segments longer than 3kb^51^.

#### Enrichment of genetic variants in genomic features

We used GREGOR (version 1.2.1) to calculate the enrichment of GWAS variants overlapping static and stretch enhancers^50^. For calculating the enrichment of glycaemic trait-associated variants in these annotations, we used the filtered list of trait-associated variants as described above (**Supplementary Table 7**) as input. For calculating the enrichment of sub-classified HbA1c variants, we included the list of loci characterized as Glycaemic, another list of loci characterized as Reticulocyte or mature Red Blood Cell, collectively representing the red blood cell fraction, along with lists of iron related or unclassified loci (**Supplementary Table 17**). We used the following parameters in GREGOR enrichment analyses: European r^2^ threshold (for inclusion of variants in LD with the lead variant) = 0.8, LD window size = 1 Mb, and minimum neighbour number = 500.

We used fGWAS (version 0.3.6)^52^ to calculate enrichment of glycaemic trait-associated variants in static and stretch enhancer annotations using summary level GWAS results. We used the default fGWAS parameters for enrichment analyses for individual annotations for each trait. For each annotation, the model provided the natural log of maximum likelihood estimate of the enrichment parameter. Annotations were considered as significantly enriched if the log2 (parameter estimate) and respective 95% confidence intervals were above zero or significantly depleted if the log2 (parameter estimate) and respective 95% confidence intervals were below zero.

We tested enrichment of trait-associated variants in static and stretch enhancer annotations with GARFIELD (v2)^53^. We formatted annotation overlap files as required by the tool; prepared input data at two GWAS thresholds - of 1×10^-5^ and a more stringent 1×10^-8^ by pruning and clumping with default parameters (garfield-prep-chr script). We calculated enrichment in each individual annotation using garfield-test.R with –c option set to 0. We also calculated the effective number of annotations using the garfield-Meff-Padj.R script. We used the effective number of annotations for each trait to obtain Bonferroni corrected significance thresholds for enrichment for each trait.

### eQTL analyses

To aid in the identification of candidate casual genes at the European-only and trans-ancestry association signals, we examined whether any of the lead variants associated with glycaemic traits (**Supplementary Table 7**) were also associated with expression level (FDR < 5%) of nearby transcripts located within 1 Mb in existing eQTL data sets of blood, subcutaneous adipose, visceral adipose, skeletal muscle, and pancreatic islet samples^54,55,93–96^. LD was estimated from the collected cohort pairwise LD information, where available, else from the European samples in 1000G phase 3. GWAS and eQTL signals likely co-localise when the GWAS variant and the variant most strongly associated with the expression level of the corresponding transcript (eSNP) exhibit high pairwise LD (r^2^ > 0.8; 1000 Genomes Phase 3, EUR). At these signals, we conducted reciprocal conditional analyses to test association between the GWAS variant and transcript level when the eSNP was also included in the model, and vice versa. We report GWAS and eQTL signals as co-localised if the association for the eSNP was not significant (FDR ≥ 5%) when conditioned on the GWAS variant; we also report signals from the eQTLGen whole blood meta-analysis data that meet only the LD threshold because conditional analysis was not possible.

#### Tissue and gene-set analysis

We performed enrichment analysis using DEPICT (Data-driven Expression-Prioritized Integration for Complex Traits) version 3, specifically developed for 1000 Genomes Project imputed meta-analysis data^97^ to identify cell types and tissues in which genes at trait-associated variants were strongly expressed, and to detect enrichment of gene-sets or pathways. DEPICT data included human gene expression data for 19,987 genes in 10,968 reconstituted gene sets, and 209 tissues/cell types. Because gene expression data in DEPICT is based on European samples and LD, we selected trait-associated variants with *P* < 10^-5^ in the European meta-analysis and tested for enrichment of signals in each reconstituted gene-set, and each tissue or cell type. Enrichment results with a false discovery rate (FDR) < 0.05 were considered significant. We ran DEPICT based on association results for all traits among: (i) cohorts with genome-wide data, or (ii) all cohorts (genome-wide and Metabochip cohorts). Because results were broadly consistent between the two approaches, we present results from the analysis that contained all cohorts as it had greater statistical power.

## References

1 in Use of Glycated Haemoglobin (HbA1c) in the Diagnosis of Diabetes Mellitus: Abbreviated Report of a WHO Consultation (World Health Organization Copyright © World Health Organization 2011., 2011).

2 Goodarzi, M. O. et al. Fasting insulin reflects heterogeneous physiological processes: role of insulin clearance. American journal of physiology. Endocrinology and metabolism 301, E402–408, doi:10.1152/ajpendo.00013.2011 (2011).

3 Dimas, A. S. et al. Impact of type 2 diabetes susceptibility variants on quantitative glycemic traits reveals mechanistic heterogeneity. Diabetes 63, 2158–2171, doi:10.2337/db13-0949 (2014).

4 Udler, M. S. et al. Type 2 diabetes genetic loci informed by multi-trait associations point to disease mechanisms and subtypes: A soft clustering analysis. PLoS medicine 15, e1002654, doi:10.1371/journal.pmed.1002654 (2018).

5 Udler, M. S., McCarthy, M. I., Florez, J. C. & Mahajan, A. Genetic Risk Scores for Diabetes Diagnosis and Precision Medicine. Endocrine reviews 40, 1500–1520, doi:10.1210/er.2019-00088 (2019).

6 Sarwar, N. et al. Diabetes mellitus, fasting blood glucose concentration, and risk of vascular disease: a collaborative meta-analysis of 102 prospective studies. Lancet 375, 2215–2222, doi:10.1016/s0140-6736(10)60484-9 (2010).

7 Wheeler, E. et al. Impact of common genetic determinants of Hemoglobin A1c on type 2 diabetes risk and diagnosis in ancestrally diverse populations: A transethnic genome-wide meta-analysis. PLoS medicine 14, e1002383, doi:10.1371/journal.pmed.1002383 (2017).

8 Dupuis, J. et al. New genetic loci implicated in fasting glucose homeostasis and their impact on type 2 diabetes risk. Nature genetics 42, 105–116, doi:10.1038/ng.520 (2010).

9 Manning, A. K. et al. A genome-wide approach accounting for body mass index identifies genetic variants influencing fasting glycemic traits and insulin resistance. Nature genetics 44, 659–669, doi:10.1038/ng.2274 (2012).

10 Walford, G. A. et al. Genome-Wide Association Study of the Modified Stumvoll Insulin Sensitivity Index Identifies BCL2 and FAM19A2 as Novel Insulin Sensitivity Loci. Diabetes 65, 3200–3211, doi:10.2337/db16-0199 (2016).

11 Horikoshi, M. et al. Discovery and Fine-Mapping of Glycaemic and Obesity-Related Trait Loci Using High-Density Imputation. PLoS genetics 11, e1005230, doi:10.1371/journal.pgen.1005230 (2015).

12 Mahajan, A. et al. Identification and functional characterization of G6PC2 coding variants influencing glycemic traits define an effector transcript at the G6PC2-ABCB11 locus. PLoS genetics 11, e1004876, doi:10.1371/journal.pgen.1004876 (2015).

13 Hwang, J. Y. et al. Genome-wide association meta-analysis identifies novel variants associated with fasting plasma glucose in East Asians. Diabetes 64, 291–298, doi:10.2337/db14-0563 (2015).

14 Chen, P. et al. Multiple nonglycemic genomic loci are newly associated with blood level of glycated hemoglobin in East Asians. Diabetes 63, 2551–2562, doi:10.2337/db13-1815 (2014).

15 Scott, R. A. et al. Large-scale association analyses identify new loci influencing glycemic traits and provide insight into the underlying biological pathways. Nature genetics 44, 991–1005, doi:10.1038/ng.2385 (2012).

16 Spanakis, E. K. & Golden, S. H. Race/ethnic difference in diabetes and diabetic complications. Current diabetes reports 13, 814–823, doi:10.1007/s11892-013-0421-9 (2013).

17 Tillin, T. et al. Insulin resistance and truncal obesity as important determinants of the greater incidence of diabetes in Indian Asians and African Caribbeans compared with Europeans: the Southall And Brent REvisited (SABRE) cohort. Diabetes care 36, 383–393, doi:10.2337/dc12-0544 (2013).

18 Whincup, P. H. et al. Early emergence of ethnic differences in type 2 diabetes precursors in the UK: the Child Heart and Health Study in England (CHASE Study). PLoS medicine 7, e1000263, doi:10.1371/journal.pmed.1000263 (2010).

19 Auton, A. et al. A global reference for human genetic variation. Nature 526, 68–74, doi:10.1038/nature15393 (2015).

20 Willer, C. J., Li, Y. & Abecasis, G. R. METAL: fast and efficient meta-analysis of genomewide association scans. Bioinformatics (Oxford, England) 26, 2190–2191, doi:10.1093/bioinformatics/btq340 (2010).

21 Yang, J., Lee, S. H., Goddard, M. E. & Visscher, P. M. GCTA: a tool for genome-wide complex trait analysis. American journal of human genetics 88, 76–82, doi:10.1016/j.ajhg.2010.11.011 (2011).

22 Yang, J. et al. Conditional and joint multiple-SNP analysis of GWAS summary statistics identifies additional variants influencing complex traits. Nature genetics 44, 369–375, s361–363, doi:10.1038/ng.2213 (2012).

23 Genome-wide association study of 14,000 cases of seven common diseases and 3,000 shared controls. Nature 447, 661–678, doi:10.1038/nature05911 (2007).

24 Mahajan, A. et al. Fine-mapping type 2 diabetes loci to single-variant resolution using high-density imputation and islet-specific epigenome maps. Nature genetics 50, 1505–1513, doi:10.1038/s41588-018-0241-6 (2018).

25 Luo, Y. et al. Transcription factor Ets1 regulates expression of thioredoxin-interacting protein and inhibits insulin secretion in pancreatic beta-cells. PloS one 9, e99049, doi:10.1371/journal.pone.0099049 (2014).

26 Braccini, L. et al. PI3K-C2gamma is a Rab5 effector selectively controlling endosomal Akt2 activation downstream of insulin signalling. Nature communications 6, 7400, doi:10.1038/ncomms8400 (2015).

27 Ng, N. H. J. et al. Tissue-Specific Alteration of Metabolic Pathways Influences Glycemic Regulation. bioRxiv, 790618, doi:10.1101/790618 (2019).

28 Aschard, H., Vilhjalmsson, B. J., Joshi, A. D., Price, A. L. & Kraft, P. Adjusting for heritable covariates can bias effect estimates in genome-wide association studies. American journal of human genetics 96, 329–339, doi:10.1016/j.ajhg.2014.12.021 (2015).

29 Lee, J. J. et al. Gene discovery and polygenic prediction from a genome-wide association study of educational attainment in 1.1 million individuals. Nature genetics 50, 1112–1121, doi:10.1038/s41588-018-0147-3 (2018).

30 Nolte, I. M. et al. Missing heritability: is the gap closing? An analysis of 32 complex traits in the Lifelines Cohort Study. European journal of human genetics: EJHG 25, 877–885, doi:10.1038/ejhg.2017.50 (2017).

31 Gaulton, K. J. et al. Genetic fine mapping and genomic annotation defines causal mechanisms at type 2 diabetes susceptibility loci. Nature genetics 47, 1415–1425, doi:10.1038/ng.3437 (2015).

32 Spracklen, C. N. et al. Identification and functional analysis of glycemic trait loci in the China Health and Nutrition Survey. PLoS genetics 14, e1007275, doi:10.1371/journal.pgen.1007275 (2018).

33 Varshney, A. et al. Genetic regulatory signatures underlying islet gene expression and type 2 diabetes. Proceedings of the National Academy of Sciences of the United States of America 114, 2301–2306, doi:10.1073/pnas.1621192114 (2017).

34 Kichaev, G. et al. Leveraging Polygenic Functional Enrichment to Improve GWAS Power. American journal of human genetics 104, 65–75, doi:10.1016/j.ajhg.2018.11.008 (2019).

35 Shriner, D. & Rotimi, C. N. Whole-Genome-Sequence-Based Haplotypes Reveal Single Origin of the Sickle Allele during the Holocene Wet Phase. American journal of human genetics 102, 547–556, doi:10.1016/j.ajhg.2018.02.003 (2018).

36 Kramer, H. J. et al. African Ancestry-Specific Alleles and Kidney Disease Risk in Hispanics/Latinos. Journal of the American Society of Nephrology: JASN 28, 915–922, doi:10.1681/asn.2016030357 (2017).

37 Ravenhall, M. et al. Novel genetic polymorphisms associated with severe malaria and under selective pressure in North-eastern Tanzania. PLoS genetics 14, e1007172, doi:10.1371/journal.pgen.1007172 (2018).

38 Hodonsky, C. J. et al. Genome-wide association study of red blood cell traits in Hispanics/Latinos: The Hispanic Community Health Study/Study of Latinos. PLoS genetics 13, e1006760, doi:10.1371/journal.pgen.1006760 (2017).

39 Gurdasani, D. et al. Uganda Genome Resource Enables Insights into Population History and Genomic Discovery in Africa. Cell 179, 984–1002.e1036, doi:10.1016/j.cell.2019.10.004 (2019).

40 Rees, M. G. et al. Cellular characterisation of the GCKR P446L variant associated with type 2 diabetes risk. Diabetologia 55, 114–122, doi:10.1007/s00125-011-2348-5 (2012).

41 Bonomo, J. A. et al. The ras responsive transcription factor RREB1 is a novel candidate gene for type 2 diabetes associated end-stage kidney disease. Human molecular genetics 23, 6441–6447, doi:10.1093/hmg/ddu362 (2014).

42 Wessel, J. et al. Low-frequency and rare exome chip variants associate with fasting glucose and type 2 diabetes susceptibility. Nature communications 6, 5897, doi:10.1038/ncomms6897 (2015).

43 Scott, R. A. et al. A genomic approach to therapeutic target validation identifies a glucose-lowering GLP1R variant protective for coronary heart disease. Science translational medicine 8, 341ra376, doi:10.1126/scitranslmed.aad3744 (2016).

44 Nai, A. et al. TMPRSS6 rs855791 modulates hepcidin transcription in vitro and serum hepcidin levels in normal individuals. Blood 118, 4459–4462, doi:10.1182/blood-2011-06-364034 (2011).

45 Soranzo, N. et al. Common variants at 10 genomic loci influence hemoglobin A(1)(C) levels via glycemic and nonglycemic pathways. Diabetes 59, 3229–3239, doi:10.2337/db10-0502 (2010).

46 Sarnowski, C. et al. Impact of Rare and Common Genetic Variants on Diabetes Diagnosis by Hemoglobin A1c in Multi-Ancestry Cohorts: The Trans-Omics for Precision Medicine Program. American journal of human genetics 105, 706–718, doi:10.1016/j.ajhg.2019.08.010 (2019).

47 Kundaje, A. et al. Integrative analysis of 111 reference human epigenomes. Nature 518, 317–330, doi:10.1038/nature14248 (2015).

48 Nagel, M. et al. Meta-analysis of genome-wide association studies for neuroticism in 449,484 individuals identifies novel genetic loci and pathways. Nature genetics 50, 920–927, doi:10.1038/s41588-018-0151-7 (2018).

49 Savage, J. E. et al. Genome-wide association meta-analysis in 269,867 individuals identifies new genetic and functional links to intelligence. Nature genetics 50, 912–919, doi:10.1038/s41588-018-0152-6 (2018).

50 Schmidt, E. M. et al. GREGOR: evaluating global enrichment of trait-associated variants in epigenomic features using a systematic, data-driven approach. Bioinformatics (Oxford, England) 31, 2601–2606, doi:10.1093/bioinformatics/btv201 (2015).

51 Parker, S. C. et al. Chromatin stretch enhancer states drive cell-specific gene regulation and harbor human disease risk variants. Proceedings of the National Academy of Sciences of the United States of America 110, 17921–17926, doi:10.1073/pnas.1317023110 (2013).

52 Pickrell, J. K. Joint analysis of functional genomic data and genome-wide association studies of 18 human traits. American journal of human genetics 94, 559–573, doi:10.1016/j.ajhg.2014.03.004 (2014).

53 Iotchkova, V. et al. GARFIELD classifies disease-relevant genomic features through integration of functional annotations with association signals. Nature genetics 51, 343–353, doi:10.1038/s41588-018-0322-6 (2019).

54 van de Bunt, M. et al. Transcript Expression Data from Human Islets Links Regulatory Signals from Genome-Wide Association Studies for Type 2 Diabetes and Glycemic Traits to Their Downstream Effectors. PLoS genetics 11, e1005694, doi:10.1371/journal.pgen.1005694 (2015).

55 Civelek, M. et al. Genetic Regulation of Adipose Gene Expression and Cardio-Metabolic Traits. American journal of human genetics 100, 428–443, doi:10.1016/j.ajhg.2017.01.027 (2017).

56 Scott, L. J. et al. The genetic regulatory signature of type 2 diabetes in human skeletal muscle. Nature communications 7, 11764, doi:10.1038/ncomms11764 (2016).

57 Ben Harouch, S., Klar, A. & Falik Zaccai, T. C. in GeneReviews((R)) (eds M. P. Adam et al.) (University of Washington, Seattle University of Washington, Seattle. GeneReviews is a registered trademark of the University of Washington, Seattle. All rights reserved., 1993).

58 Agus, D. B. et al. Vitamin C crosses the blood-brain barrier in the oxidized form through the glucose transporters. The Journal of clinical investigation 100, 2842–2848, doi:10.1172/jci119832 (1997).

59 Wolking, S. et al. Focal epilepsy in glucose transporter type 1 (Glut1) defects: case reports and a review of literature. Journal of neurology 261, 1881–1886, doi:10.1007/s00415-014-7433-5 (2014).

60 Guallar, D. et al. RNA-dependent chromatin targeting of TET2 for endogenous retrovirus control in pluripotent stem cells. Nature genetics 50, 443–451, doi:10.1038/s41588-018-0060-9 (2018).

61 Bian, F. et al. TET2 facilitates PPARgamma agonist-mediated gene regulation and insulin sensitization in adipocytes. Metabolism: clinical and experimental 89, 39–47, doi:10.1016/j.metabol.2018.08.006 (2018).

62 Yoo, Y. et al. TET-mediated hydroxymethylcytosine at the Ppargamma locus is required for initiation of adipogenic differentiation. International journal of obesity (2005) 41, 652–659, doi:10.1038/ijo.2017.8 (2017).

63 Lees, J. A. et al. Lipid transport by TMEM24 at ER-plasma membrane contacts regulates pulsatile insulin secretion. Science (New York, N.Y.) 355, doi:10.1126/science.aah6171 (2017).

64 Pottekat, A. et al. Insulin biosynthetic interaction network component, TMEM24, facilitates insulin reserve pool release. Cell reports 4, 921–930, doi:10.1016/j.celrep.2013.07.050 (2013).

65 Terzolo, M. et al. Adrenal incidentaloma: a new cause of the metabolic syndrome? The Journal of clinical endocrinology and metabolism 87, 998–1003, doi:10.1210/jcem.87.3.8277 (2002).

66 Muscogiuri, G., Colao, A. & Orio, F. Insulin-Mediated Diseases: Adrenal Mass and Polycystic Ovary Syndrome. Trends in endocrinology and metabolism: TEM 26, 512–514, doi:10.1016/j.tem.2015.07.010 (2015).

67 Altieri, B. et al. Adrenocortical tumors and insulin resistance: What is the first step? International journal of cancer 138, 2785–2794, doi:10.1002/ijc.29950 (2016).

68 Androulakis, II et al. Patients with apparently nonfunctioning adrenal incidentalomas may be at increased cardiovascular risk due to excessive cortisol secretion. The Journal of clinical endocrinology and metabolism 99, 2754–2762, doi:10.1210/jc.2013-4064 (2014).

69 Johansson, M. et al. The influence of obesity-related factors in the etiology of renal cell carcinoma-A mendelian randomization study. PLoS medicine 16, e1002724, doi:10.1371/journal.pmed.1002724 (2019).

70 Diamanti-Kandarakis, E. & Dunaif, A. Insulin resistance and the polycystic ovary syndrome revisited: an update on mechanisms and implications. Endocrine reviews 33, 981–1030, doi:10.1210/er.2011-1034 (2012).

71 Morris, A. P. et al. Large-scale association analysis provides insights into the genetic architecture and pathophysiology of type 2 diabetes. Nature genetics 44, 981–990, doi:10.1038/ng.2383 (2012).

72 Leong, A. et al. Mendelian Randomization Analysis of Hemoglobin A(1c) as a Risk Factor for Coronary Artery Disease. Diabetes care 42, 1202–1208, doi:10.2337/dc18-1712 (2019).

73 Lambert, S. A., Abraham, G. & Inouye, M. Towards clinical utility of polygenic risk scores. Human molecular genetics 28, R133–r142, doi:10.1093/hmg/ddz187 (2019).

74 Duncan, L. et al. Analysis of polygenic risk score usage and performance in diverse human populations. Nature communications 10, 3328, doi:10.1038/s41467-019-11112-0 (2019).

75 Khera, A. V. et al. Polygenic Prediction of Weight and Obesity Trajectories from Birth to Adulthood. Cell 177, 587–596.e589, doi:10.1016/j.cell.2019.03.028 (2019).

76 Mostafavi, H. et al. Variable prediction accuracy of polygenic scores within an ancestry group. eLife 9, doi:10.7554/eLife.48376 (2020).

77 D’Orazio, P. et al. Approved IFCC recommendation on reporting results for blood glucose (abbreviated). Clinical chemistry 51, 1573–1576, doi:10.1373/clinchem.2005.051979 (2005).

78 Voight, B. F. et al. The metabochip, a custom genotyping array for genetic studies of metabolic, cardiovascular, and anthropometric traits. PLoS genetics 8, e1002793, doi:10.1371/journal.pgen.1002793 (2012).

79 Abecasis, G. R. et al. An integrated map of genetic variation from 1,092 human genomes. Nature 491, 56–65, doi:10.1038/nature11632 (2012).

80 Li, Y., Willer, C. J., Ding, J., Scheet, P. & Abecasis, G. R. MaCH: using sequence and genotype data to estimate haplotypes and unobserved genotypes. Genetic epidemiology 34, 816–834, doi:10.1002/gepi.20533 (2010).

81 Pei, Y. F., Zhang, L., Li, J. & Deng, H. W. Analyses and comparison of imputation-based association methods. PloS one 5, e10827, doi:10.1371/journal.pone.0010827 (2010).

82 Winkler, T. W. et al. Quality control and conduct of genome-wide association meta-analyses. Nature protocols 9, 1192–1212, doi:10.1038/nprot.2014.071 (2014).

83 Devlin, B. & Roeder, K. Genomic control for association studies. Biometrics 55, 997–1004 (1999).

84 Morris, A. P. Transethnic meta-analysis of genomewide association studies. Genetic epidemiology 35, 809–822, doi:10.1002/gepi.20630 (2011).

85 Benner, C. et al. FINEMAP: efficient variable selection using summary data from genome-wide association studies. Bioinformatics (Oxford, England) 32, 1493–1501, doi:10.1093/bioinformatics/btw018 (2016).

86 Astle, W. J. et al. The Allelic Landscape of Human Blood Cell Trait Variation and Links to Common Complex Disease. Cell 167, 1415–1429.e1419, doi:10.1016/j.cell.2016.10.042 (2016).

87 Benyamin, B. et al. Novel loci affecting iron homeostasis and their effects in individuals at risk for hemochromatosis. Nature communications 5, 4926, doi:10.1038/ncomms5926 (2014).

88 Binesh, N. & Rezghi, M. Fuzzy clustering in community detection based on nonnegative matrix factoriztion with two novel evaluation criteria. Applied Soft Computing 69, 689–703 (2018).

89 Scott, R. A. et al. An Expanded Genome-Wide Association Study of Type 2 Diabetes in Europeans. Diabetes 66, 2888–2902, doi:10.2337/db16-1253 (2017).

90 Ernst, J. et al. Mapping and analysis of chromatin state dynamics in nine human cell types. Nature 473, 43–49, doi:10.1038/nature09906 (2011).

91 Mikkelsen, T. S. et al. Comparative epigenomic analysis of murine and human adipogenesis. Cell 143, 156–169, doi:10.1016/j.cell.2010.09.006 (2010).

92 Ernst, J. & Kellis, M. ChromHMM: automating chromatin-state discovery and characterization. Nature methods 9, 215–216, doi:10.1038/nmeth.1906 (2012).

93 Battle, A., Brown, C. D., Engelhardt, B. E. & Montgomery, S. B. Genetic effects on gene expression across human tissues. Nature 550, 204–213, doi:10.1038/nature24277 (2017).

94 Zhernakova, D. V. et al. Identification of context-dependent expression quantitative trait loci in whole blood. Nature genetics 49, 139–145, doi:10.1038/ng.3737 (2017).

95 Westra, H. J. et al. Systematic identification of trans eQTLs as putative drivers of known disease associations. Nature genetics 45, 1238–1243, doi:10.1038/ng.2756 (2013).

96 Joehanes, R. et al. Integrated genome-wide analysis of expression quantitative trait loci aids interpretation of genomic association studies. Genome biology 18, 16, doi:10.1186/s13059-016-1142-6 (2017).

97 Pers, T. H. et al. Biological interpretation of genome-wide association studies using predicted gene functions. Nature communications 6, 5890, doi:10.1038/ncomms6890 (2015).

