## Supplementary Figure for "The Trans-Ancestral Genomic Architecture of Glycaemic Traits"

***<u>Supplemental Figure Legends</u>***

**Supplementary Figure 1**. Flow diagram of this study. The figure shows the data, key methods and main analyses included in this effort.

**Supplementary Figure 2**. Manhattan plot and QQ plot of the single-ancestry meta-analysis of each trait. Figures show the quantile-quantile plot and Manhattan plot for the single-ancestry meta-analysis of each trait in each available ancestry. Traits are labelled by uppercase letters A (FG), B (FI), C (HbA1c) and D (2hGlu). Ancestries are labelled by roman numerals i (EUR), ii (EAS), iii (HISP), iv (AA), v (SAS) and vi (AFR).

**Supplementary Figure 3**. Manhattan plot of the trans-ancestry meta-analysis of each trait. Figures show the Manhattan plot for the trans-ancestry meta-analysis of each trait. Traits are labelled by uppercase letters A (FG), B (FI), C (2hGlu) ad D (HbA1c).

**Supplementary Figure 4.** Forest plot of FG-associated variant rs6190947. Novel FG locus identified near *ETS1* in African Americans. Results were not significant in other ancestry populations. Among the African American cohorts, sample sizes ranged from 319 (CFS) to 6,519 (WHI) with a minimum imputation score of $r^2$=0.56 and $P_{het}$=0.40.

**Supplementary Figure 5**. Forest plot of FG-associated variant rs12315677. Novel FG locus identified near *PIK3C2G* in Hispanics. Results were not significant in other ancestry populations. Among the Hispanic cohorts, sample sizes ranged from 130 (TRIPOD) to 10,065 (SOL) with a minimum imputation score of $r^2$=0.69 and $P_{het}$=0.43.

**Supplementary Figure 6.** Forest plot of FI-associated variant rs13258890. Novel FI locus identified near *NKX2-6* in Europeans. Results were not significant in other ancestry populations. Among the European cohorts, sample sizes ranged from 155 (HELICPomak) to 8,518 (METSIM) with a minimum imputation score of $r^2$=0.76 and $P_{het}$=0.53.

**Supplementary Figure 7**. Forest plot of FI-associated variant rs200678953. Novel FI locus identified near *D21S2088E* in Europeans. Results were not significant in other ancestry populations. Among the European cohorts, sample sizes ranged from 155 (HELICPomak) to 8,518 (METSIM) with a minimum imputation score of $r^2$=0.44 and $P_{het}$=0.96.

**Supplementary Figure 8**. Forest plot of HbA1c-associated variant rs184506746. Novel HbA1c locus identified near CD99L2 in Europeans. Results were not significant in other ancestry populations. Among the European cohorts, sample sizes ranged from 496 (BioMe) to 4,289 (SardiNIA) with a minimum imputation score of $r^2$=0.47 and $P_{het}$=0.37.

**Supplementary Figure 9**. Locus zoom plot of FG-associated locus G6PC2. Figure includes top five panels to show the associations in five ancestries and one bottom panel to show the genes and MAFs. On each of top five panels, points present the -$\log_{10}$(p-value) and are coloured by their LD level with the trans-ancestry lead variant in purple diamond. The colourful par labelled by $R^2$ shows and maximum LD level of each variant with the single-ancestry signals in the black circles. The colourful par labelled by LDscore shows the summation of $R^2$ between each variant and all the other variants divided by the maximum of the summations.

**Supplementary Figure 10.** Venn diagram. Figure shows the overlaps of TA loci between traits.

**Supplementary Figure 11.** Frequency versus effect. These plots display the allele frequency versus effect size for all signals detected through the trans-ancestry meta-analyses, for each of the four traits. Frequency and effect are from the European meta-analyses. The power curves were computed based on the European sample size for each trait, and the mean (m) and standard deviation (sd) computed on the FENLAND study - FG : m=4.83 mmol/l, sd=0.68 - FI : m=3.69 mmol/l, sd=0.60 - 2hGlu : m=5.30 mmol/l, sd=1.74 - HbA1c : m=5.55%, sd=0.48.

**Supplementary Figure 12.** EAF correlation and heterogeneity test. Figure shows the Pearson correlation of EAF on the lower tri-angle and p-value of heterogeneity test on the upper tri-angle of the trans-ancestry lead variants associated with each trait between ancestries. Correlations > 0.7 are in bold.

**Supplementary Figure 13.** Locus zoom plot of FI-associated locus *HDAC7*. Figure includes top five panels to show the associations in five ancestries and one bottom panel to show the genes and MAFs. On each of top five panels, points present the $-\log_{10}$(p-value) and are coloured by their LD level with the trans-ancestry lead variant in purple diamond. The colourful par labelled by $R^2$ shows and maximum LD level of each variant with the single-ancestry signals in the black circles. The colourful par labelled by LDscore shows the summation of $R^2$ between each variant and all the other variants divided by the maximum of the summations.

**Supplementary Figure 14**. Forest plot of FI-associated variant rs111264094. Established FI locus identified near *HDAC7* in Europeans. Results were not significant in other ancestry populations. Among the European cohorts, sample sizes ranged from 155 (HELICPomak) to 8,518 (METSIM) with a minimum imputation score of $r^2$=0.42 and $P_{het}$=0.78.

**Supplementary Figure 15**. Forest plot of T2D GRS. Figure shows the association results between the T2D GRS from the DIAGRAM consortium (by HbA1c cluster categories) and risk for T2D in the UK Biobank.

**Supplementary Figure 16.** Enrichment of glycemic trait associated GWAS variants to overlap genomic annotations using GREGOR. Figure shows enrichment for 59 total static and stretch enhancer annotations considered. Significance (red) is determined after Bonferroni correction to account for 59 total annotations tested for each trait; nominal significance (P<0.05) is indicated in yellow.

**Supplementary Figure 17**. Enrichment of glycemic trait associated GWAS variants to overlap genomic annotations using fGWAS. Figure shows log2(Fold Enrichment) of GWAS variants to overlap 59 static and stretch enhancer annotations calculated. Significant enrichment (red) is considered if the 95% confidence intervals do not overlap 0.

**Supplementary Figure 18.** Enrichment of glycemic trait associated GWAS variants to overlap genomic annotations using GARFIELD. Figure shows the beta or effect size (log odds ratio) for GWAS variants to overlap 59 static and stretch enhancer annotations. GWAS variants were included at two significance thresholds, 1e-05 (A) and 1e-08 (B). Significance (red) is determined after Bonferroni correction to account for effective annotations tested for each trait reported by GARFIELD (see supplementary note); nominal significance (P<0.05) is indicated in yellow.

*Supplementary Figure 1*

### Figure FlowDiag

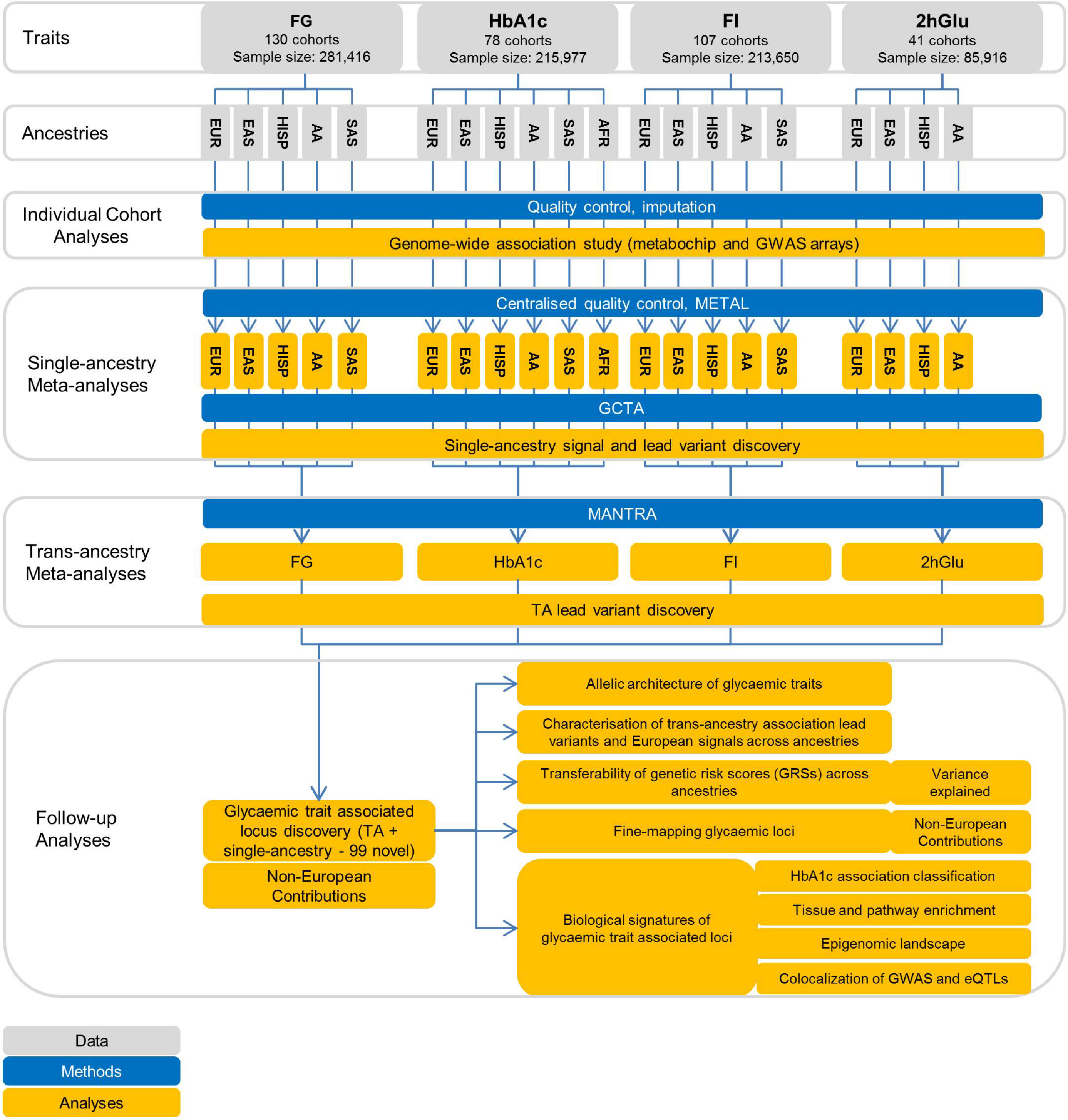

### A. Fasting Glucose

#### i. European

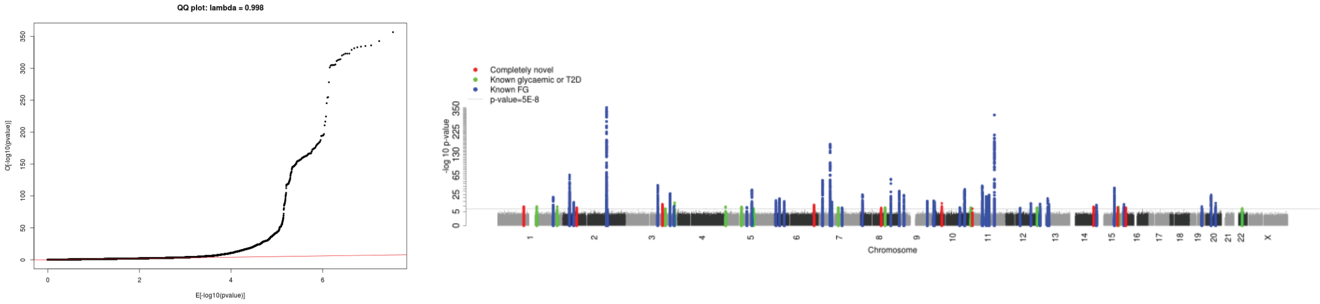


#### ii. East Asian

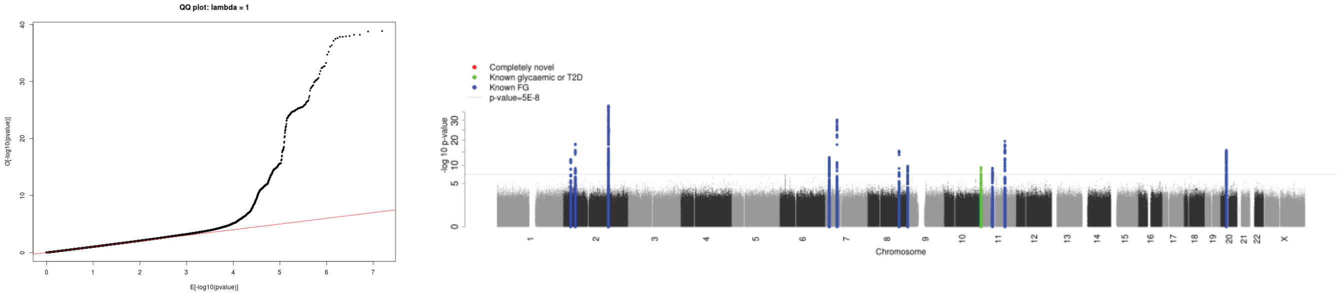


#### iii. Hispanic

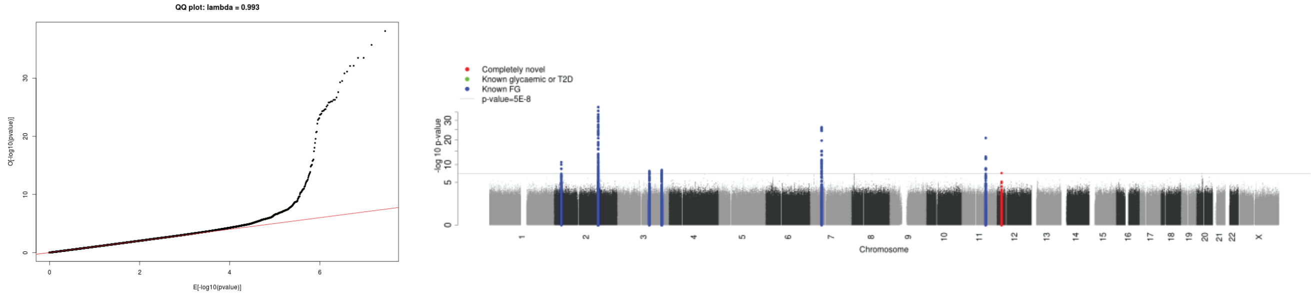


#### iv. African American

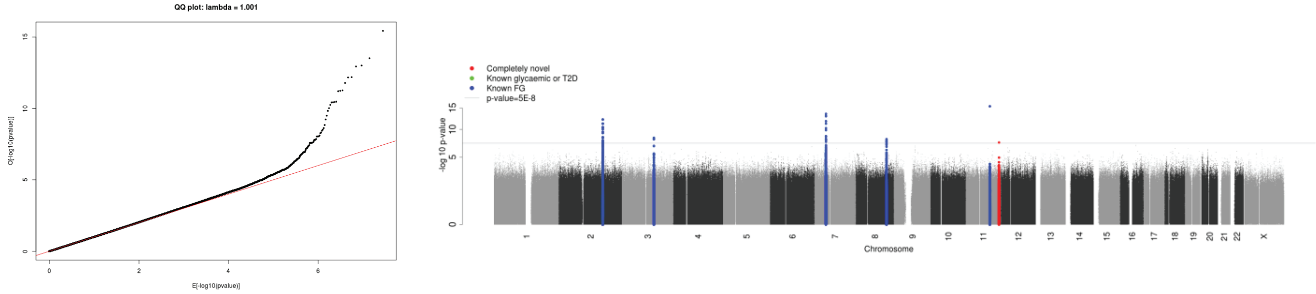


#### v. South Asian

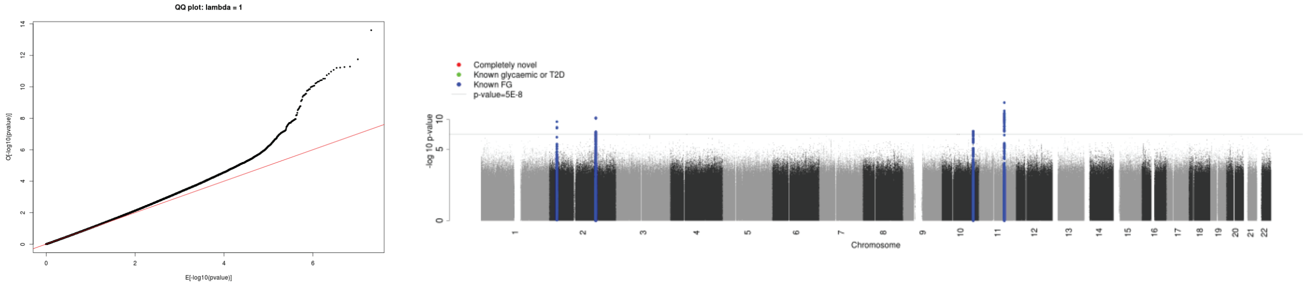

#### B. Fasting Insulin

##### i. European

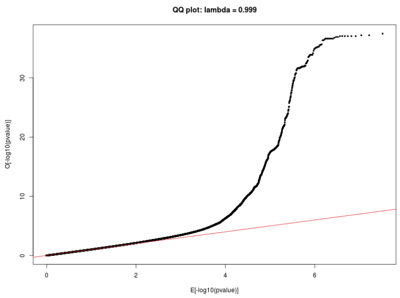


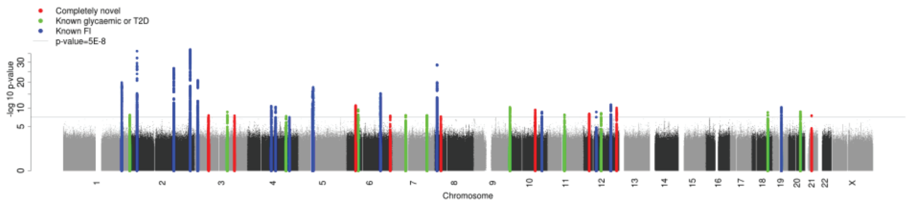


##### ii. East Asian

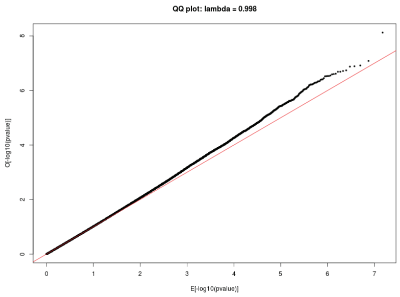


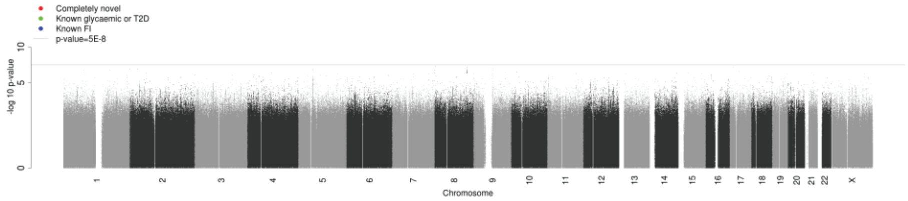


##### iii. Hispanic

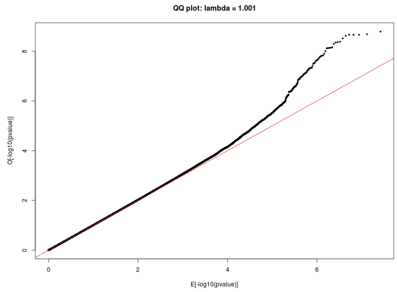


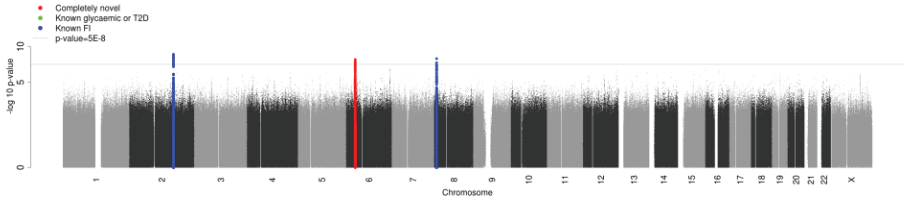


##### iv. African American

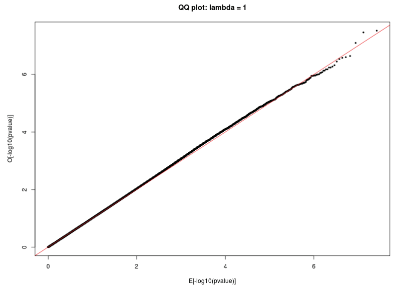


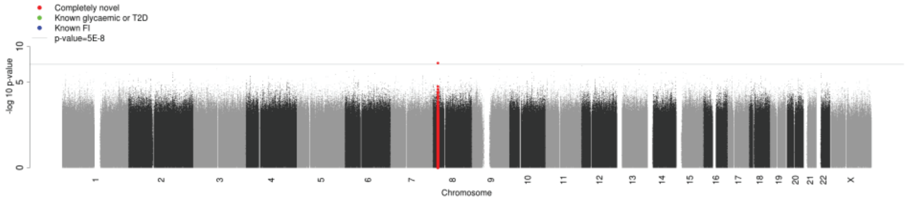


##### v. South Asian

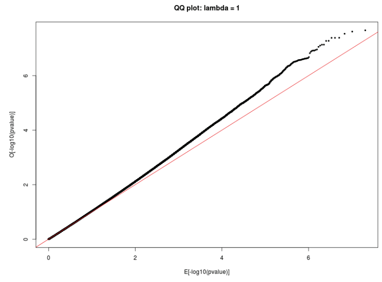


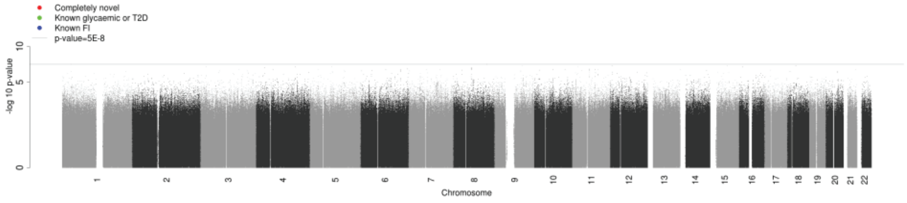

#### C. HbA1c

##### i. European

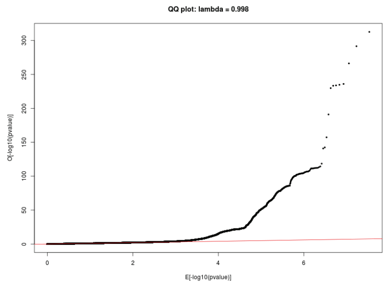


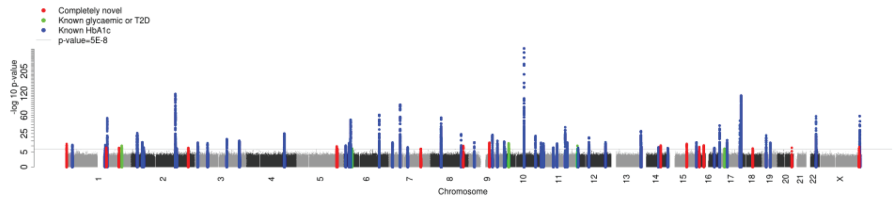


##### ii. East Asian

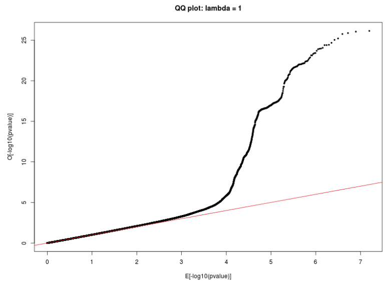


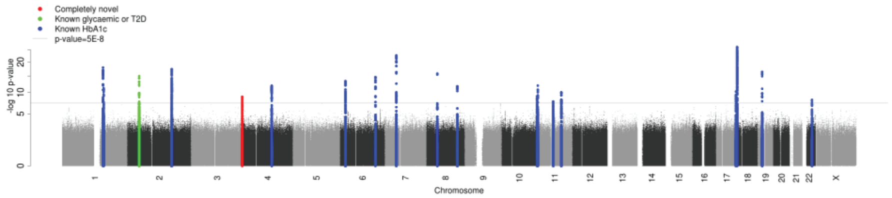


##### iii. Hispanic

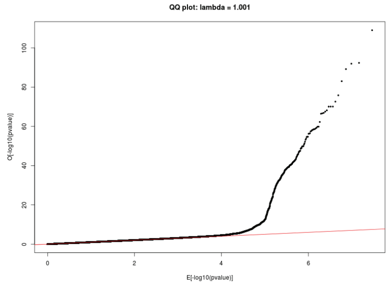


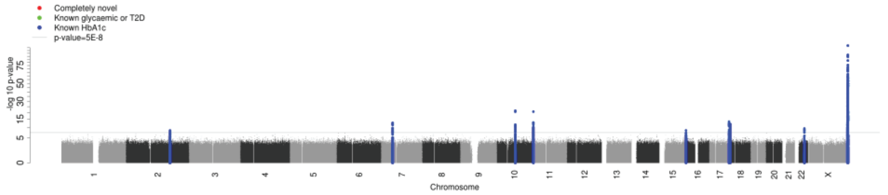


##### iv. African American

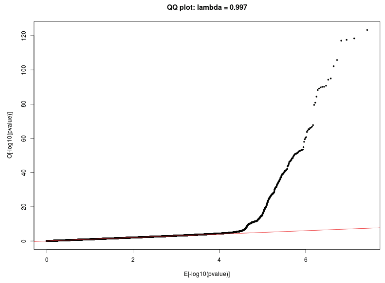


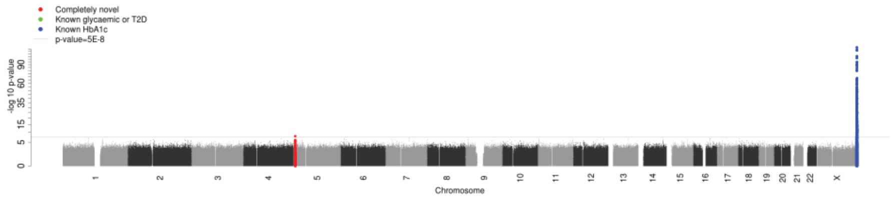


##### v. South Asian

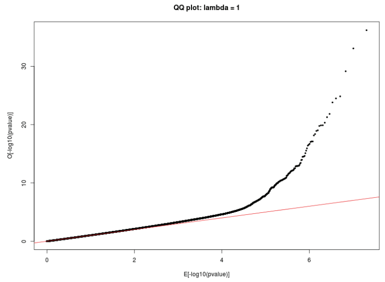


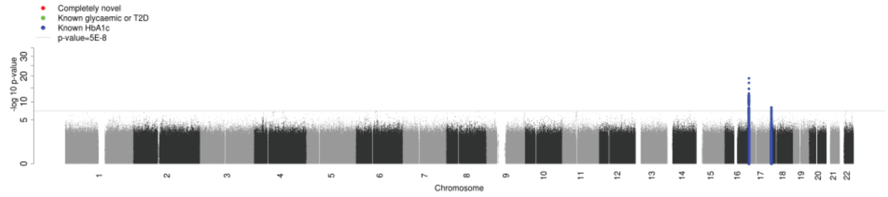

### C. HbA1c

#### vi. African

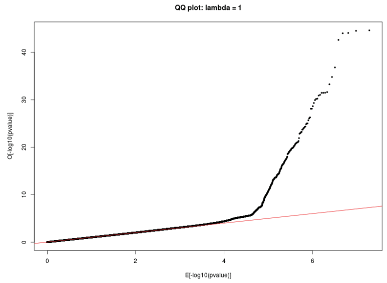


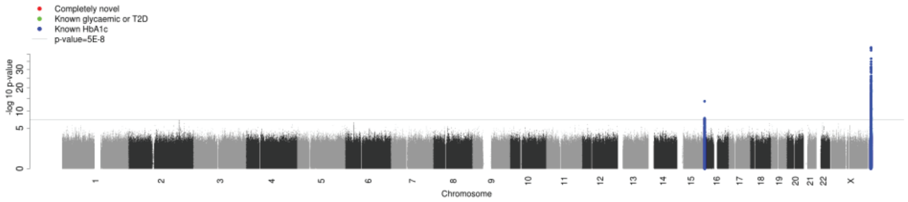

*Supplementary Figure 2, cont.*

#### D. 2 hour Glucose

##### i. European

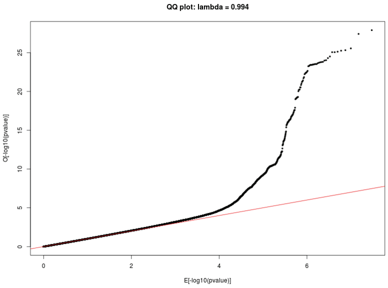


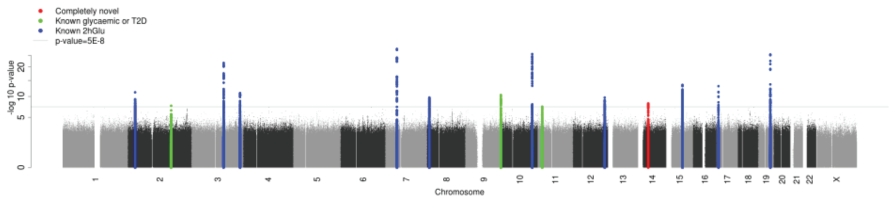


##### ii. East Asian

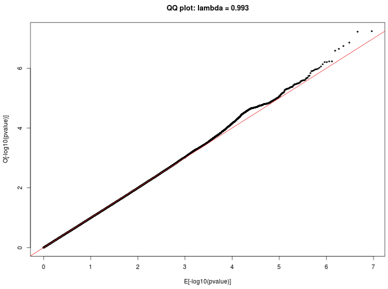


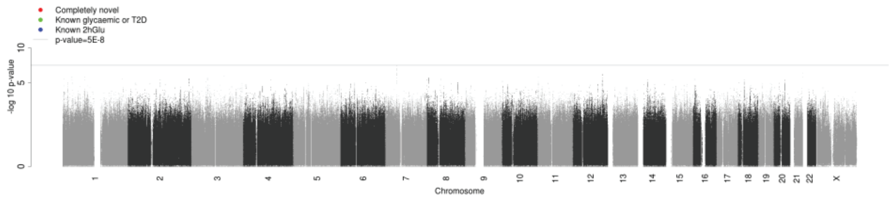


##### iii. Hispanic

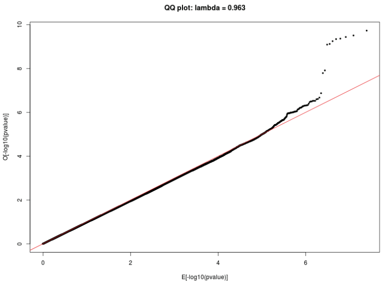


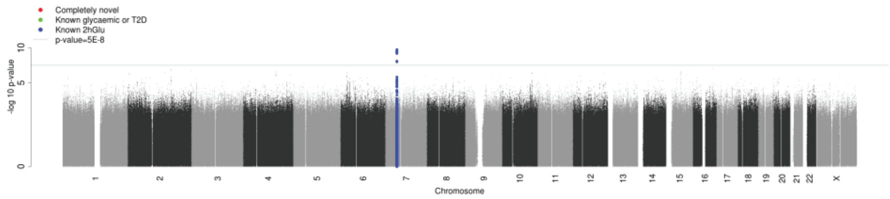


##### iv. African American

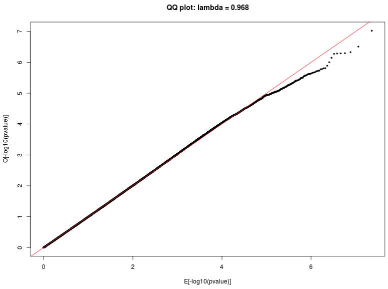


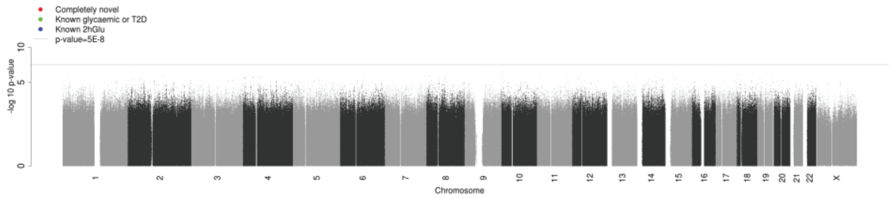

#### A. Fasting Glucose

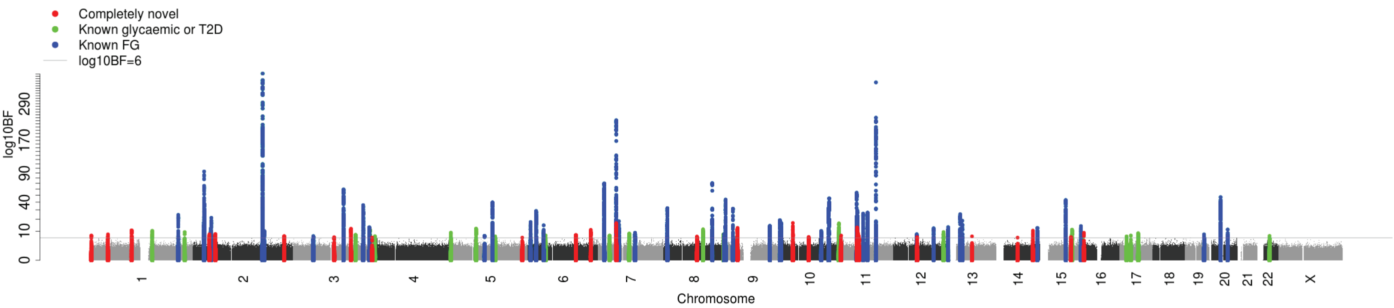


#### B. Fasting Insulin

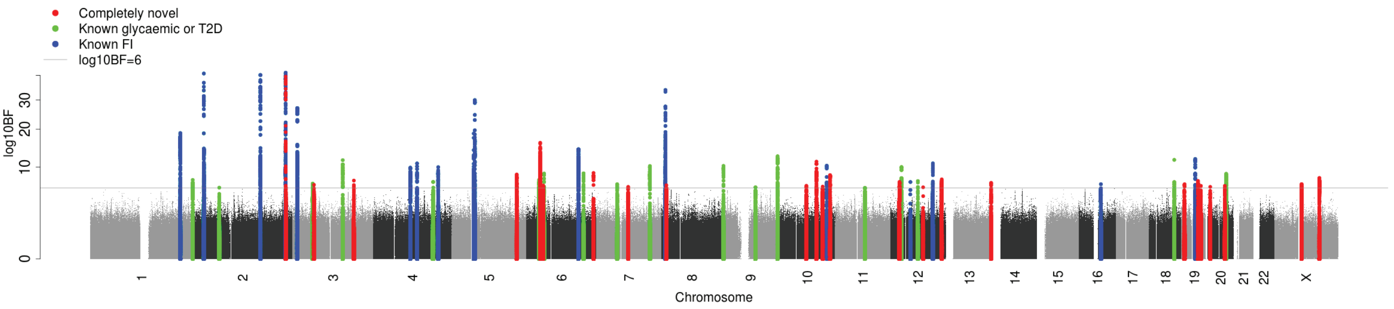


#### C. 2 hour Glucose

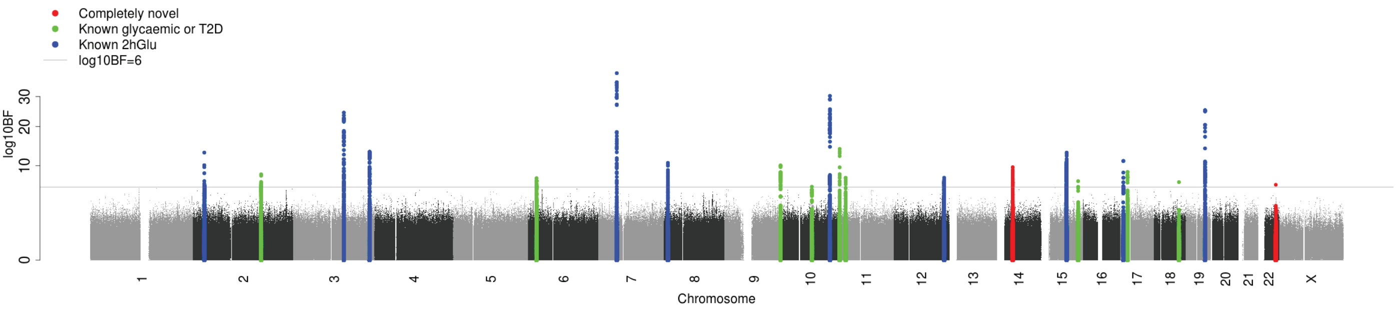


#### D. HbA1c

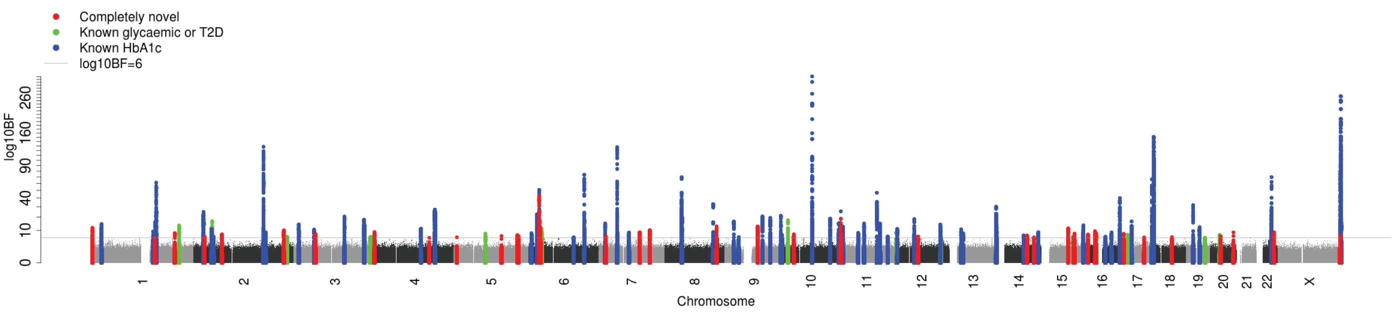

*Supplementary Figure 4*

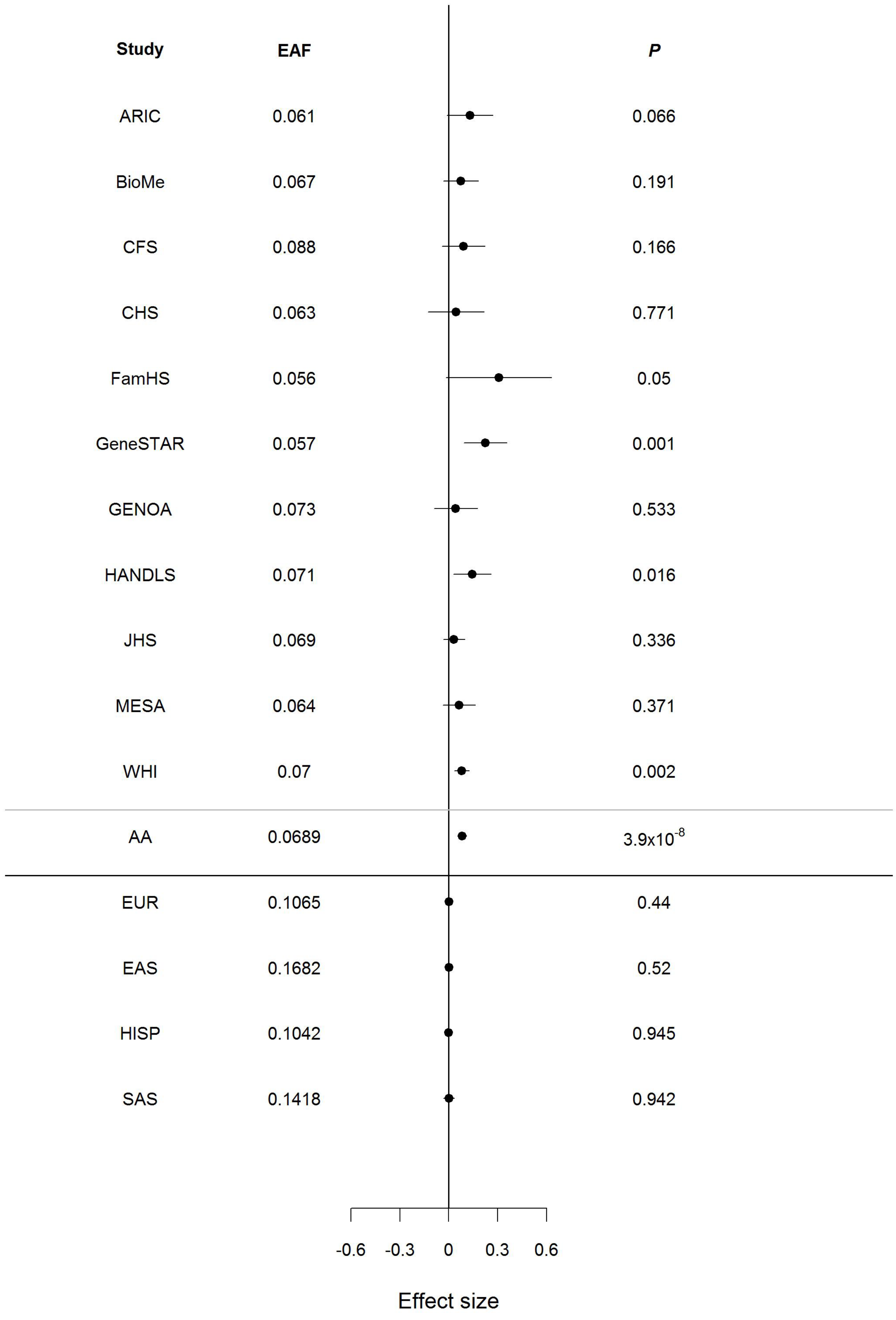

*Supplementary Figure 5*

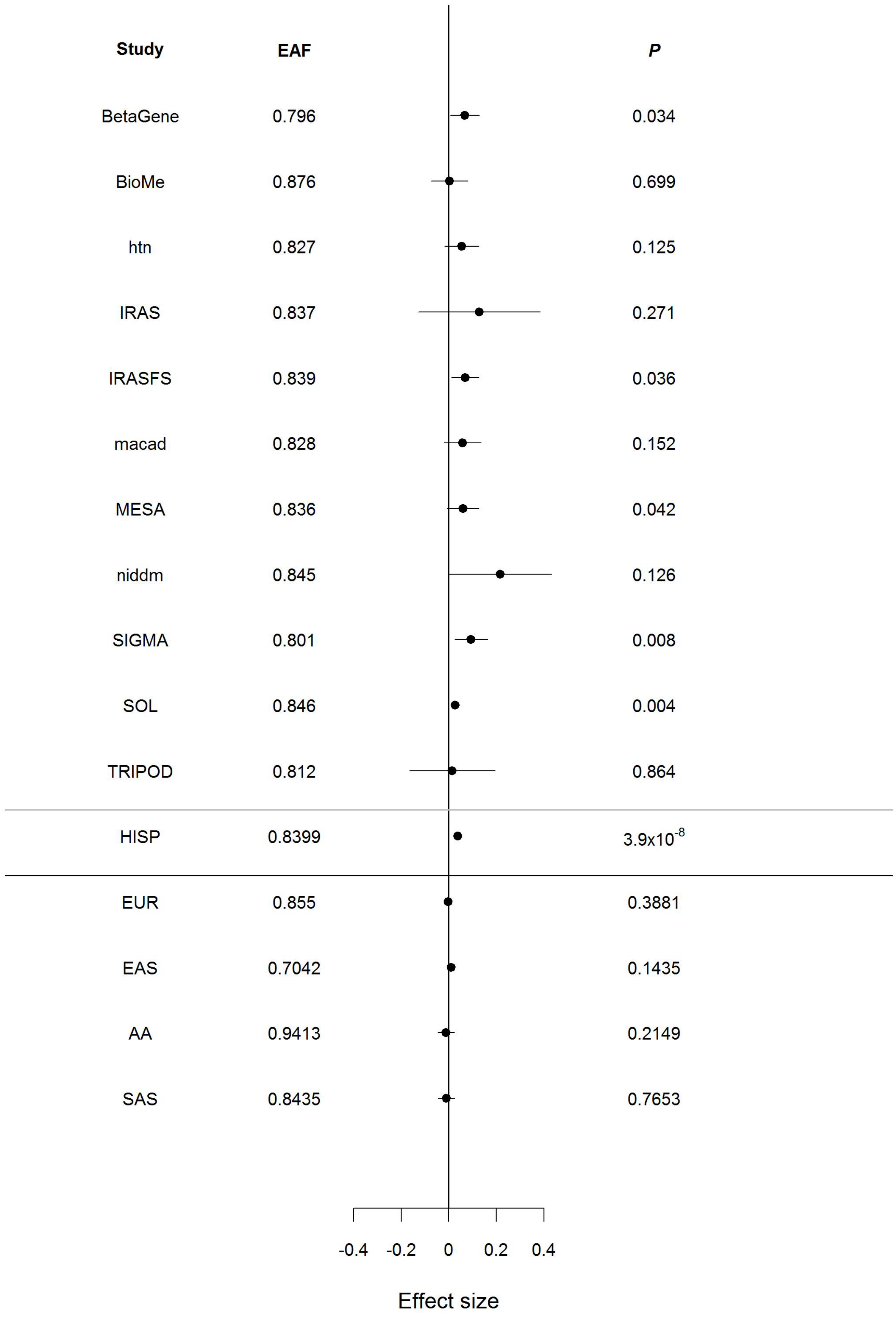

*Supplementary Figure 6*

| Study | EAF | *P* |
|---|---|---|
| AGES | 0.809 | 0.52 |
| ALSPACchildren | 0.751 | 0.13 |
| ALSPACmothers | 0.747 | 0.338 |
| AMISH | 0.847 | 0.367 |
| ARIC | 0.753 | 0.233 |
| BC1936 | 0.756 | 0.139 |
| CFS | 0.743 | 0.631 |
| CHS | 0.747 | 0.478 |
| CoLaus | 0.737 | 0.431 |
| CROATIA-Korcula | 0.736 | 0.26 |
| CROATIA-Vis | 0.784 | 0.184 |
| DGI | 0.768 | 0.929 |
| EGCUT | 0.774 | 0.089 |
| ERF4 | 0.781 | 0.045 |
| FamHS | 0.763 | 0.684 |
| Fenland | 0.748 | 0.541 |
| Fenland-OMICS | 0.766 | 0.244 |
| FHS | 0.748 | 0.59 |
| FrenchAdultControl | 0.747 | 0.692 |
| FrenchAdultObese | 0.759 | 0.527 |
| FrenchYoungObese | 0.727 | 0.3 |
| FUSION | 0.796 | 0.264 |
| GENOA | 0.759 | 0.288 |
| GOYA-Male-CASES | 0.744 | 0.928 |
| GOYA-Male-CONTROLS | 0.763 | 0.719 |
| HELICMANOLIS | 0.678 | 0.313 |
| HELICPomak | 0.716 | 0.169 |
| INCHIANTI | 0.721 | 0.611 |
| KORA | 0.76 | 0.017 |
| LEIPZIG-adults | 0.782 | 0.019 |
| LEIPZIG-kids | 0.761 | 0.079 |
| LLS | 0.776 | 0.614 |
| LOLIPOP-EW610 | 0.742 | 0.499 |
| LOLIPOP-EWA | 0.77 | 0.761 |
| LOLIPOP-EWP | 0.759 | 0.308 |
| LURIC | 0.751 | 0.491 |
| MESA | 0.745 | 0.219 |
| METSIM | 0.798 | 0.043 |
| NEO | 0.754 | 0.007 |
| NFBC1966 | 0.801 | 0.638 |
| NTR | 0.759 | 0.39 |
| ORCADES | 0.738 | 0.349 |
| PIVUS | 0.77 | 0.269 |
| PREVEND | 0.771 | 0.389 |
| PROCARDIS | 0.749 | 0.386 |
| PROSPER | 0.752 | 0.164 |
| Raine Study | 0.742 | 0.512 |
| Raine Study (20 year) | 0.742 | 0.237 |
| RISC | 0.753 | 0.123 |
| RS1 | 0.757 | 0.137 |
| RS2 | 0.749 | 0.002 |
| RS3 | 0.757 | 0.127 |
| SardiNIA | 0.812 | 0.225 |
| SORBS | 0.8 | 0.531 |
| TRAILSpop | 0.782 | 0.567 |
| TWINSUK | 0.756 | 0.135 |
| ULSAM | 0.769 | 0.568 |
| EUR | 0.7651 | $2.77 \times 10^{-8}$ |
| AA | 0.6862 | 0.01184 |
| EAS | 0.86 | 0.5845 |
| SAS | 0.8083 | 0.9288 |
| HISP | 0.8049 | 0.4673 |

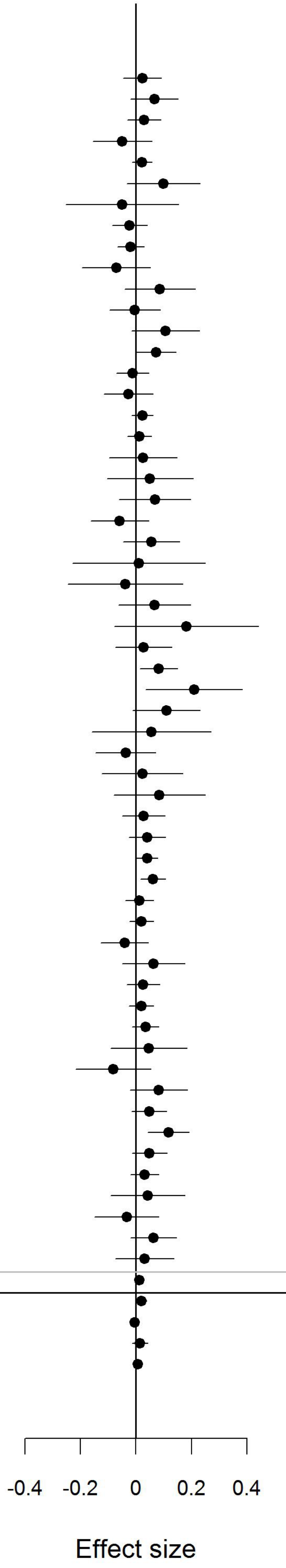

*Supplementary Figure 7*

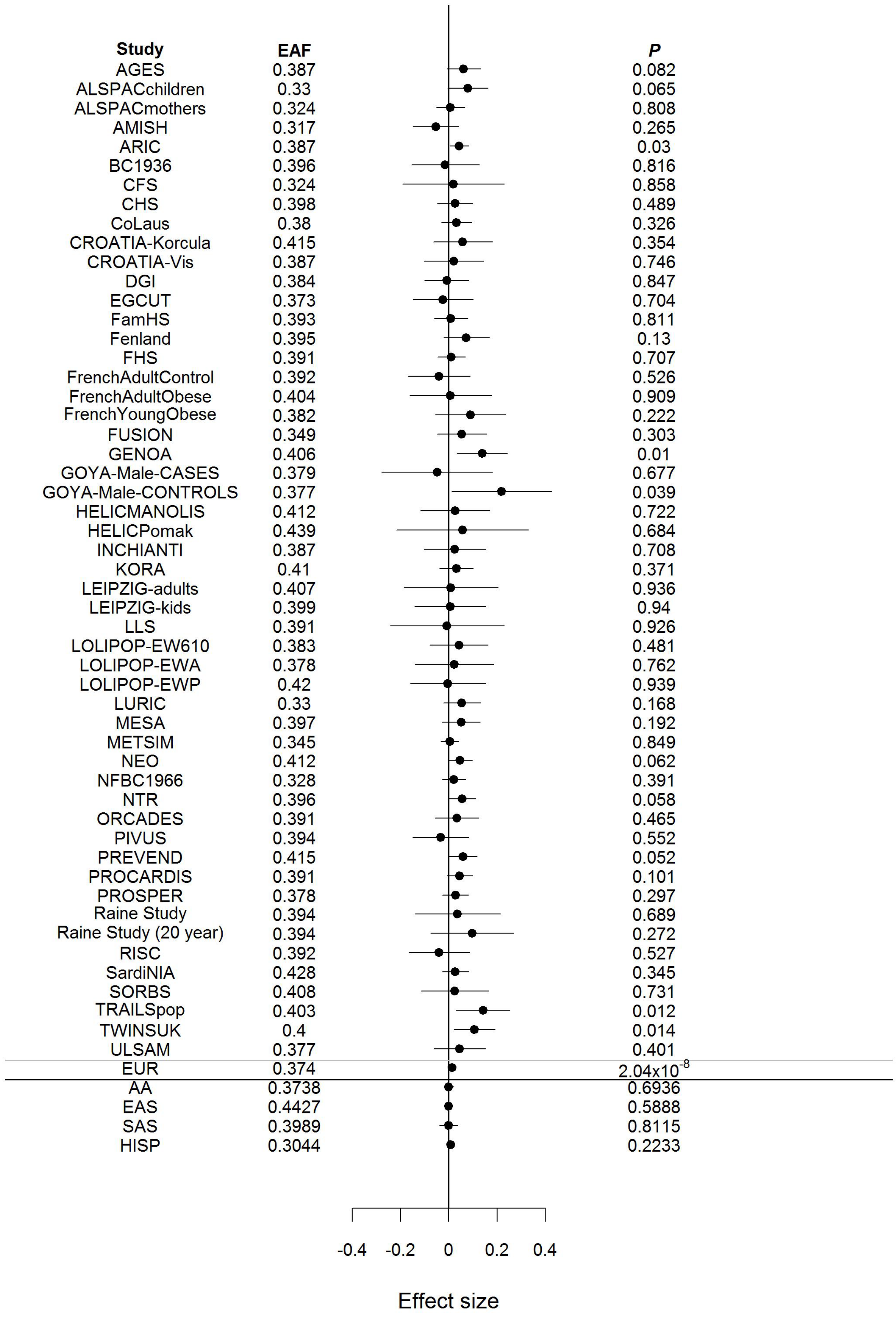

*Supplementary Figure 8*

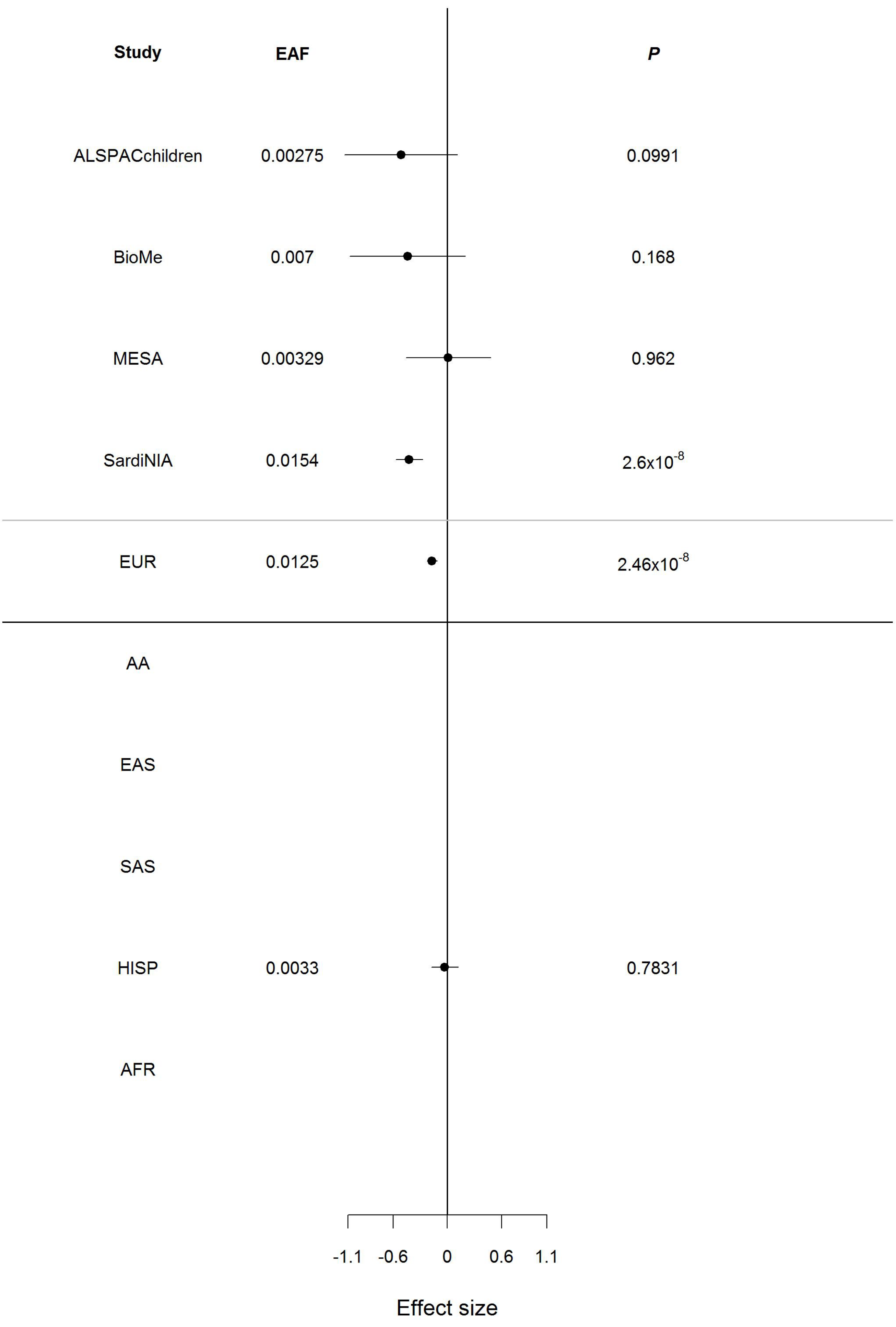

*Supplementary Figure 9*

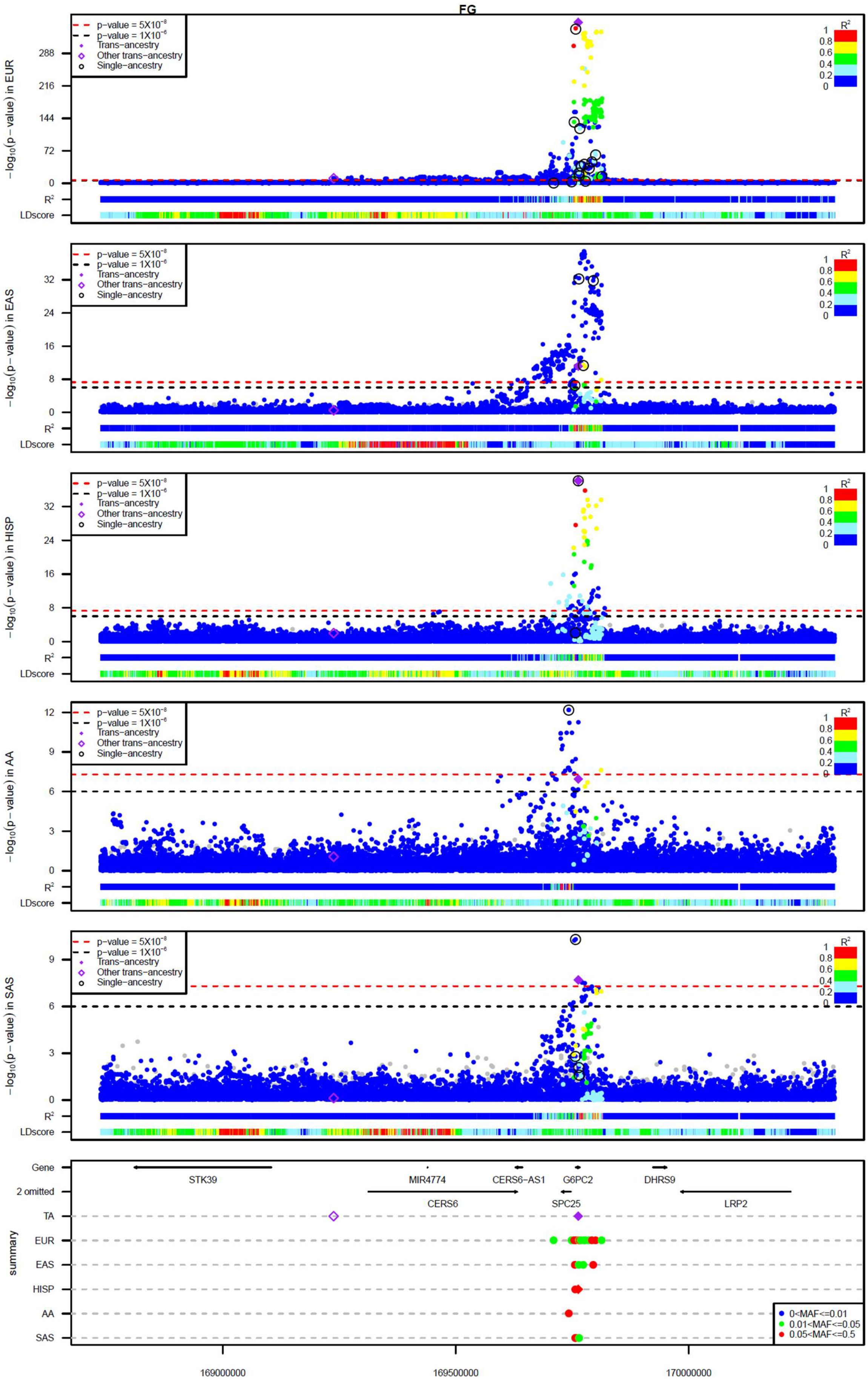

*Supplementary Figure 10*

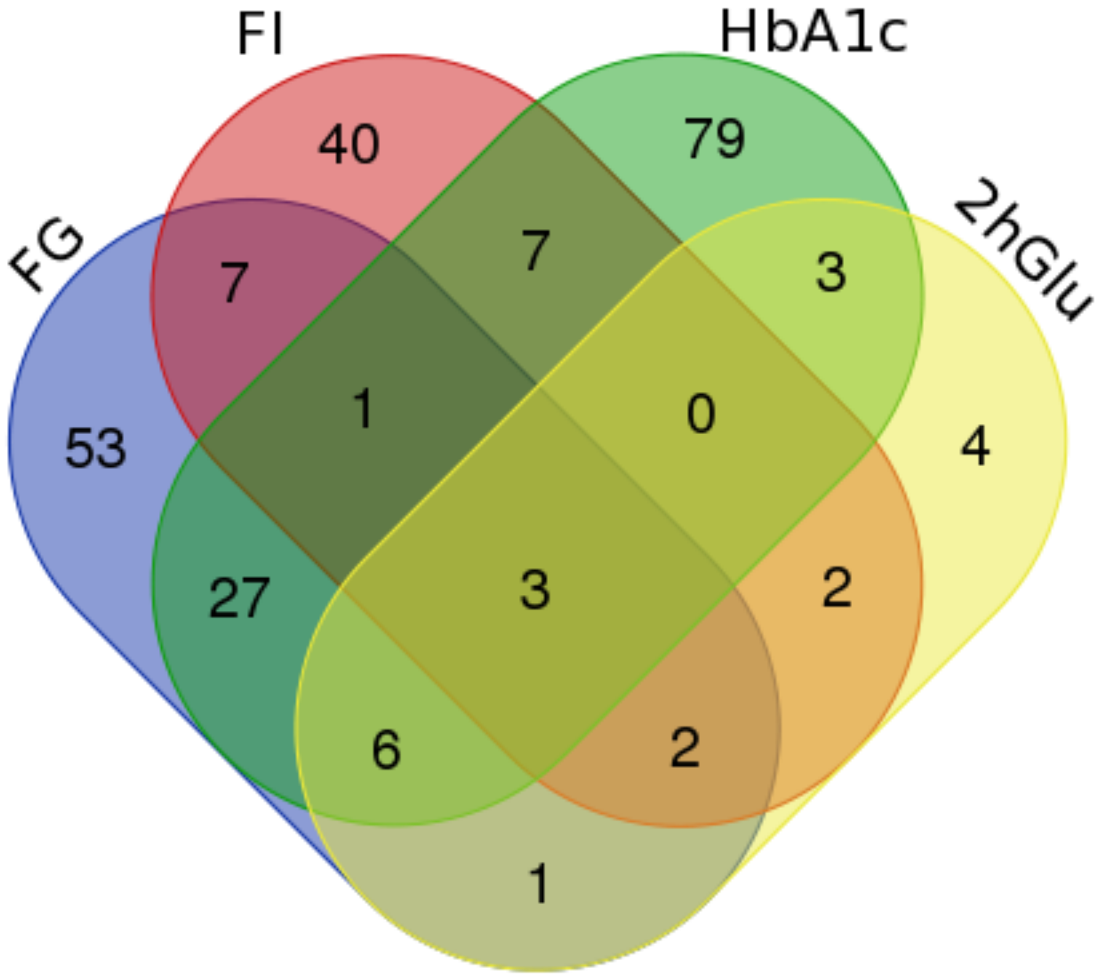

*Supplementary Figure 11*

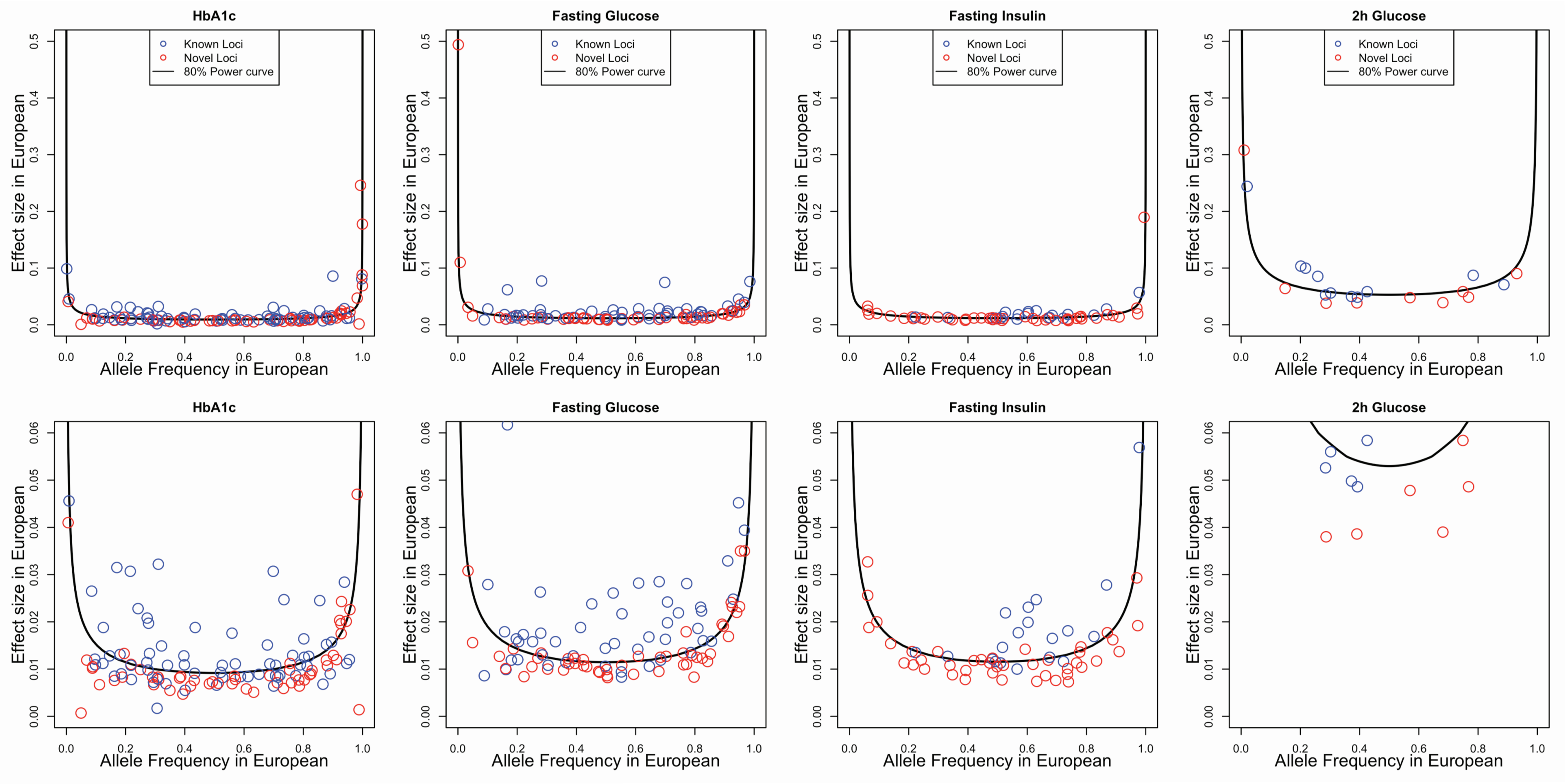

*Supplementary Figure 12*

| | | EUR | EAS | HISP | AA | SAS | AFR |
|---|---|---|---|---|---|---|---|
| **EUR** | | **FG** | $2.72X10^{-11}$ | 0.016 | $1.20X10^{-5}$ | 0.16 | |
| | | **HbA1c** | $1.55X10^{-15}$ | $1.98X10^{-7}$ | $<2.2X10^{-16}$ | 0.017 | $8.86X10^{-7}$ |
| | | **FI** | $1.13X10^{-6}$ | $1.7X10^{-4}$ | 0.352 | 0.3 | |
| | | **2hGlu** | 0.348 | 0.841 | 0.098 | | |
| **EAS** | | 0.36 | **FG** | $5.9X10^{-4}$ | 0.0262 | $7.5X10^{-4}$ | |
| | | 0.35 | **HbA1c** | 0.0099 | $9.24X10^{-8}$ | 0.057 | $1.01X10^{-5}$ |
| | | 0.34 | **FI** | 0.00103 | 0.224 | 0.014 | |
| | | 0.19 | **2hGlu** | 0.527 | 0.083 | | |
| **HISP** | | 0.79 | 0.58 | **FG** | 0.057 | 0.032 | |
| | | 0.88 | 0.55 | **HbA1c** | $1.98X10^{-6}$ | 0.531 | $<2.2X10^{-16}$ |
| | | 0.83 | 0.57 | **FI** | 0.044 | 0.623 | |
| | | 0.86 | 0.4 | **2hGlu** | 0.056 | | |
| **AA** | | 0.36 | 0.31 | 0.6 | **FG** | 0.084 | |
| | | 0.41 | 0.37 | 0.67 | **HbA1c** | $8.85X10^{-6}$ | $<2.2X10^{-16}$ |
| | | 0.53 | 0.09 | 0.59 | **FI** | 0.419 | |
| | | 0.21 | -0.03 | 0.37 | **2hGlu** | | |
| **SAS** | | 0.82 | 0.65 | 0.79 | 0.48 | **FG** | |
| | | 0.8 | 0.63 | 0.82 | 0.5 | **HbA1c** | 0.046 |
| | | 0.82 | 0.41 | 0.74 | 0.54 | **FI** | |
| | | | | | | **2hGlu** | |
| **AFR** | | | | | | | **FG** |
| | | 0.21 | 0.24 | 0.5 | 0.98 | 0.3 | **HbA1c** |
| | | | | | | | **FI** |
| | | | | | | | **2hGlu** |

| $R^2$ | | | | | | |
|---|---|---|---|---|---|---|
| | 0 | 0.2 | 0.4 | 0.6 | 0.8 | 1 |

### *Supplementary Figure 13*

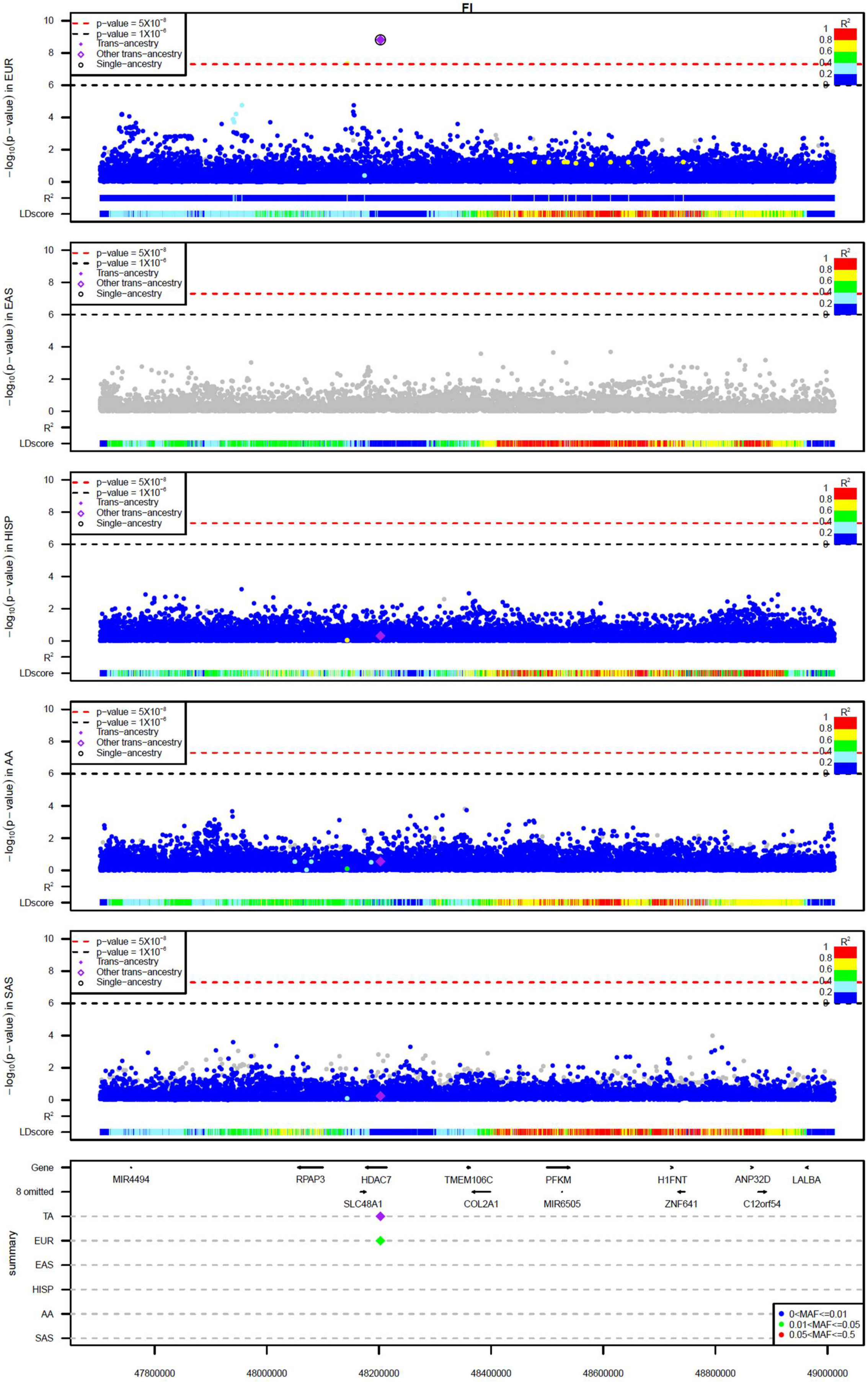

*Supplementary Figure 14*

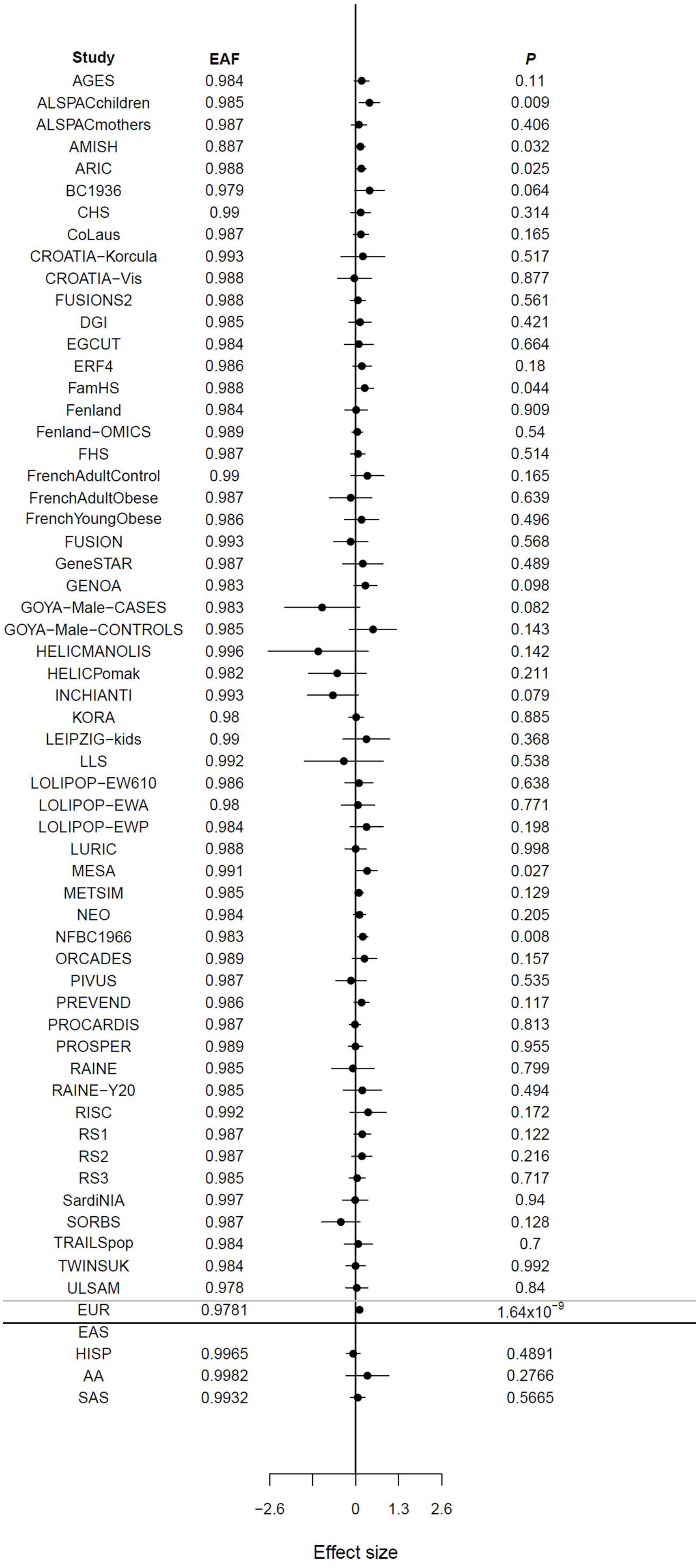

*Supplementary Figure 15*

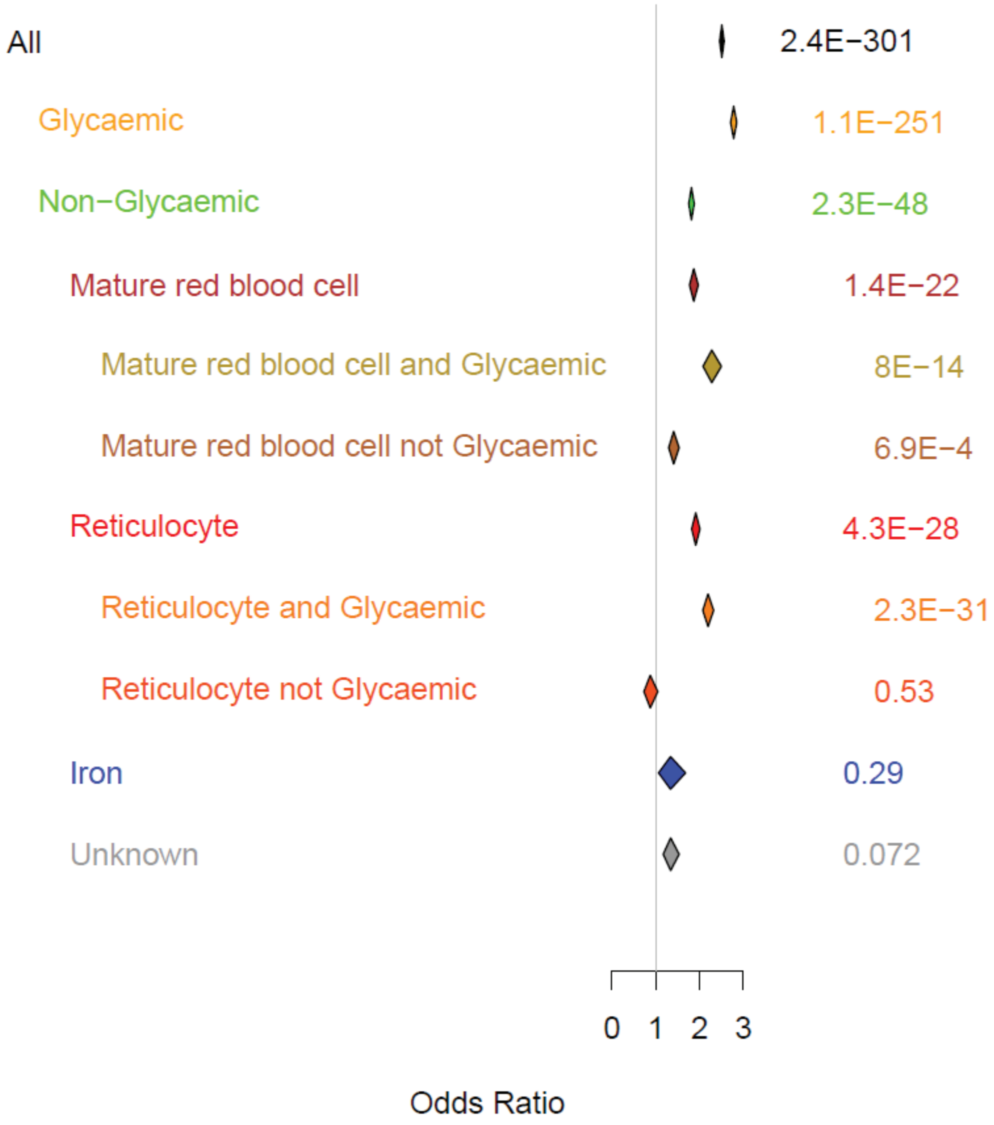

*Supplementary Figure 16*

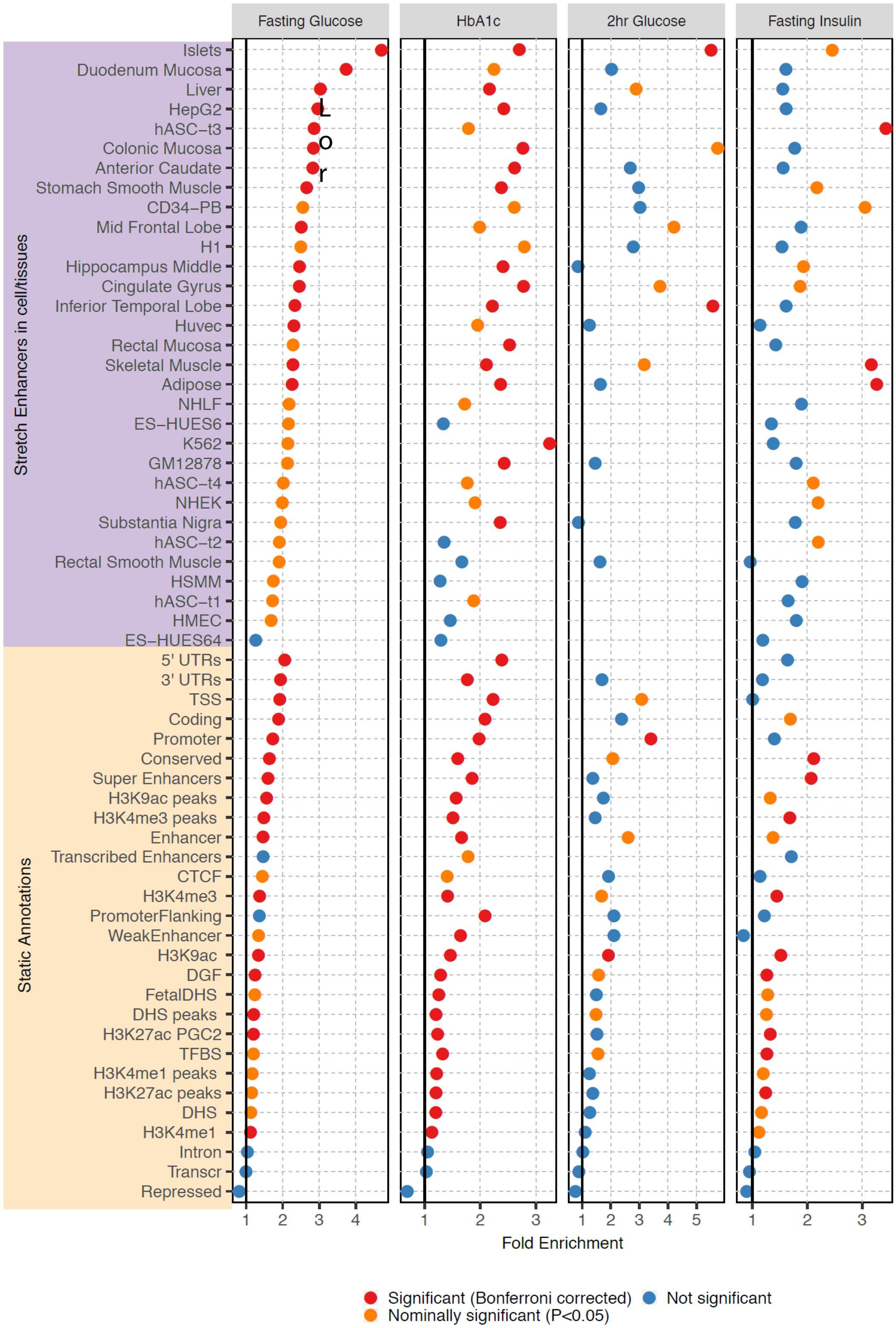

*Supplementary Figure 17*

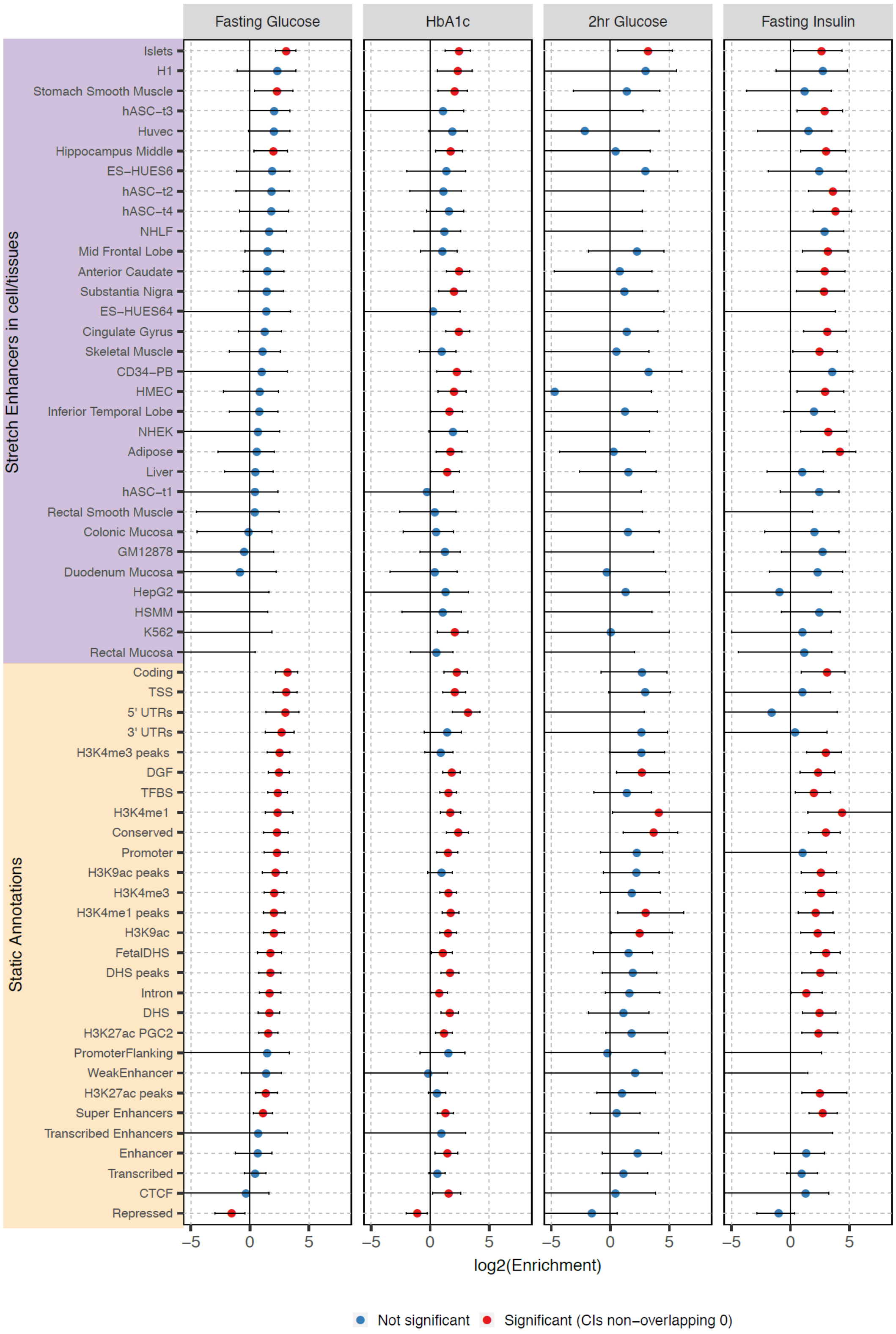

*Supplementary Figure 18*

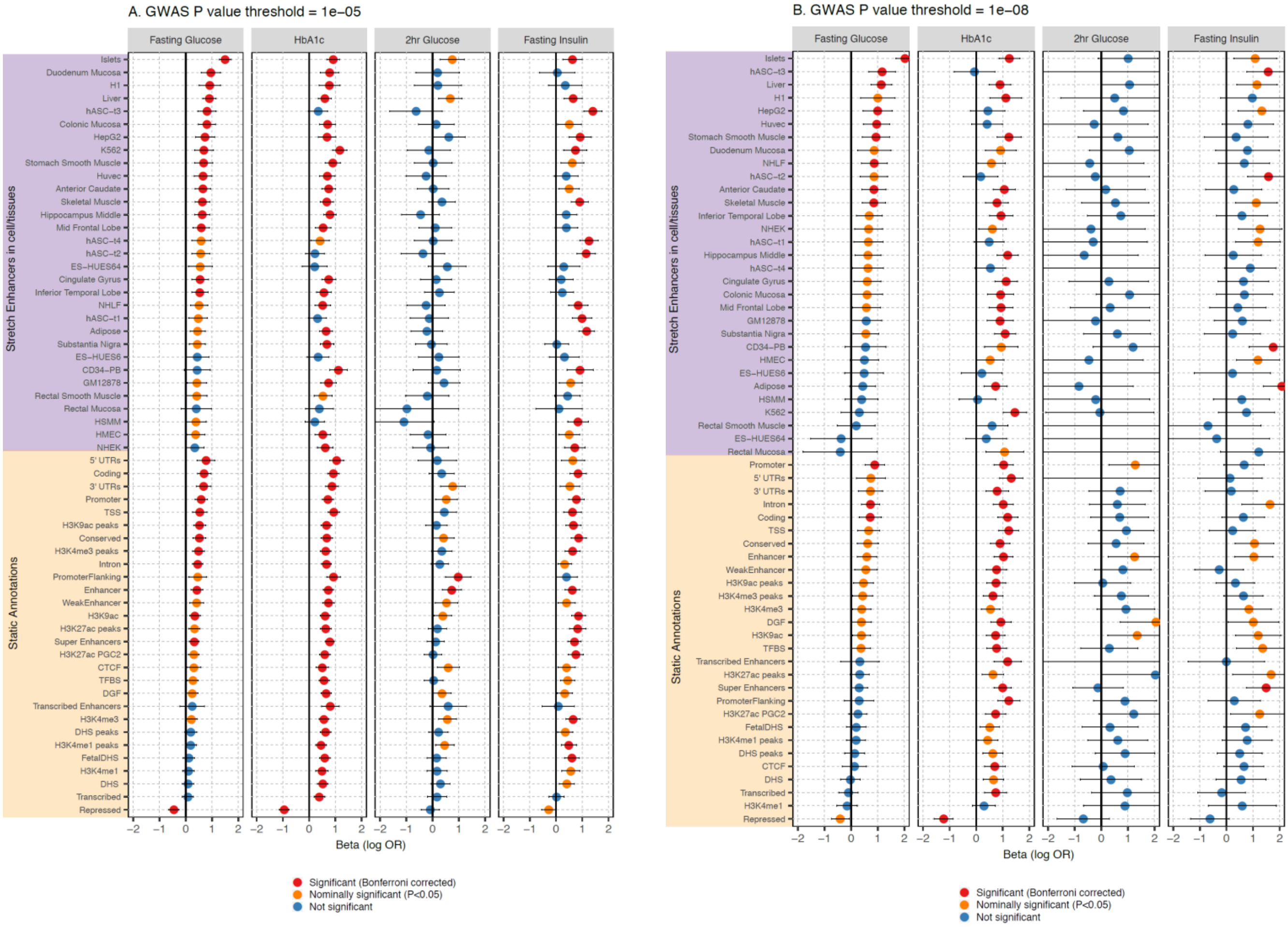
