## Supplementary note for "The Trans-Ancestral Genomic Architecture of Glycaemic Traits"

### 1. Glycaemic trait locus discovery

#### a. Manual curation of single-ancestry index and lead variants and trans-ancestry lead variants

To ensure single-ancestry index and lead variants were robust, we performed manual inspection of forest plots by at least two authors for any single-ancestry index and lead variants with QC flags. QC flags that led to manual inspection were : (i) ≤ 1 cohort with *P*-value < 0.05 & consistent direction of effect compared to single-ancestry METAL results; (ii) a single cohort within the ancestry provided data to the single-ancestry index and lead variant; (iii) single-ancestry meta-analysis heterogeneity *P* < $1x10^{-5}$ (rank inverse normal transformation); (iv) opposite direction of effect between the single-ancestry meta-analysis and the trans-ancestry meta-analysis in METAL (i.e. combining all the single-ancestry meta-analyses results - rank inverse normal transformation); (v) MAF < 1%; and (vi) sample size for single-ancestry index and lead variant < 1/3 maximum sample size for that ancestry. In total we detected 335 single-ancestry index and lead variants across all traits, of which 295 passed without inspection, 32 passed after manual inspection, and 8 failed the manual inspection.

Similarly to the single-ancestry analysis, we performed manual inspection of forest plots for TA lead variants meeting one of the following flags, indicating possible QC issues: (i) ≤ 1 cohort with *P* < 0.05 & consistent direction of effect with the trans-ancestry meta-analysis in METAL (i.e. combining all the single-ancestry meta-analyses results - rank inverse normal transformation); (ii) only one ancestry provided data to the trans-ancestry lead variant; (iii) ≤ 1 ancestry with single-ancestry meta-analysis *P* ≤ 0.05; (iv) all ancestries with single-ancestry meta-analysis *P* ≤ 0.05 have a single cohort providing data to the TA variant; (v) single-ancestry meta-analysis heterogeneity *P* (rank inverse normal transformation) < $1x10^{-5}$ for ≥ 1 ancestry; (vi) heterogeneity *P-value* for trans-ancestry meta-analysis in METAL combining the single-ancestry meta-analyses results (rank inverse normal transformation) < $1x10^{-5}$; (vii) heterogeneity $log_{10}$BF from MANTRA (rank inverse normal transformation) > 3.7 [=$log_{10}(0.05/1x10^{-5})$]; (viii) ≤ 4 variants with $log_{10}$BF > 6 in the trans-ancestry distance-based clump for that signal; (ix) opposite direction of effect between the raw and rank inverse normal meta-analysis results (trans-ancestry meta-analysis in METAL combining the single-ancestry meta-analyses results); (x) MAF < 1%; or (xi) sample size for the TA lead variant < 1/3 maximum sample size within an ancestry or overall. Of 463 trans-ancestry lead variants across all traits, 184 passed without inspection, 131 passed after inspection, and 148 failed the manual inspection.

#### b. Characterisation of loci

Based on discovery efforts across all four traits and ancestries, this effort led to the identification of 242 loci (235 trans-ancestry and seven single-ancestry) associated with at least one glycaemic trait (**Supplementary Table 2)**. The distribution of the length of each locus is shown in **Supplementary Figure N19** and encompasses between 763 Kb – 3.04 Mb and is < 1.5 Mb in 98% of the loci (237/242). In 94% of the loci (227/242), the maximum distance between two adjoining signals is ≤ 200 Kb (**Supplementary Figure N20**). The maximum distance between adjoining signals from different traits is also ≤ 200 Kb for 96% of loci (232/242 – **Supplementary Figure N21**).

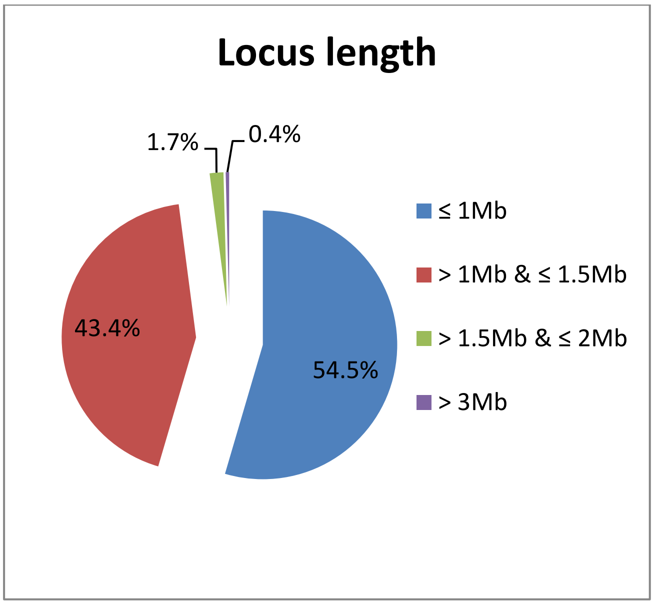

**Supplementary Figure N19 –** Distribution of the length (Mb) of each locus.

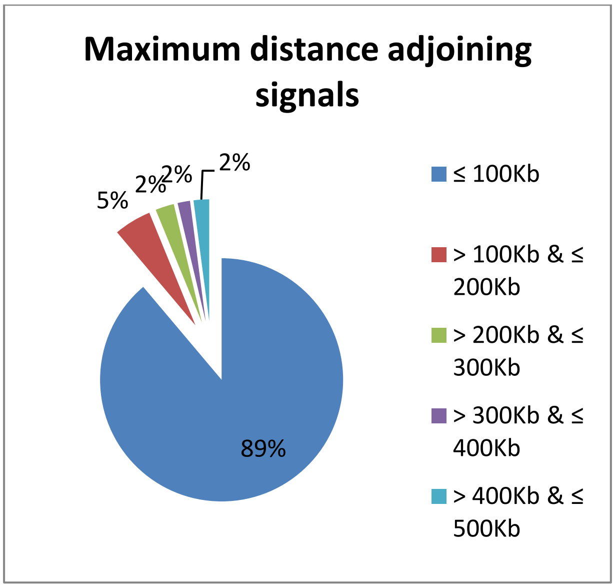

**Supplementary Figure N20 –** Distribution of the maximum distance (Kb) between two adjoining signals from the same trait.

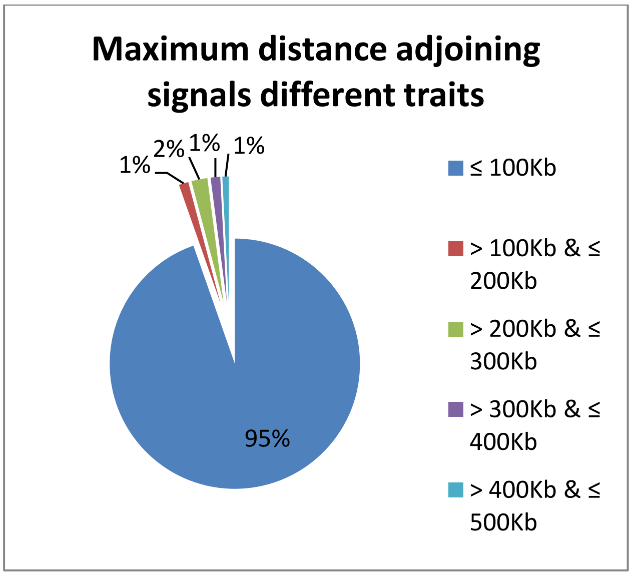

**Supplementary Figure N21** - Distribution of the maximum distance (Kb) between two adjoining signals from different traits.

The largest associated region is locus 242, spanning > 3 Mb on chromosome X, which includes 4 trans-ancestry lead variants and 12 single-ancestry lead variants from one trait (**Supplementary Table N23; Supplementary Figure N22**). Locus 149 has the longest maximum distance between adjoining signals from both the same and different traits (EUR FG rs10717442 and trans-ancestry rs34228231; > 479 Kb, EUR HbA1c rs10838696 and trans-ancestry FG rs34228231; nearly 457 Kb) with overall length of 1.5 Mb (**Supplementary Figure N23**). In these two extreme examples, the long length of the locus and distance between signals are due to the distance-based clumping (locus 242) and very strong association signals in the trans-ancestry analysis with wider LD blocks (locus 149), suggesting that overall this definition is correctly grouping signals together in relevant loci.

**Supplementary Table N23 –** Details of the two loci with the longest overall locus length (242) and the longest distance between adjoining signals from same trait or different traits (149).

| Locus ID | Chr. | Start position (bp) | End position (bp) | Length (bp) | Max. distance between adjacent signals (bp) | |
|---|---|---|---|---|---|---|
| | | | | | Same trait | Different traits |
| 149 | 11 | 46,778,502 | 48,320,241 | 1,541,740 | 479,561 | 456,956 |
| 242 | X | 152,362,433 | 155,405,080 | 3,042,648 | 319,200 | NA |

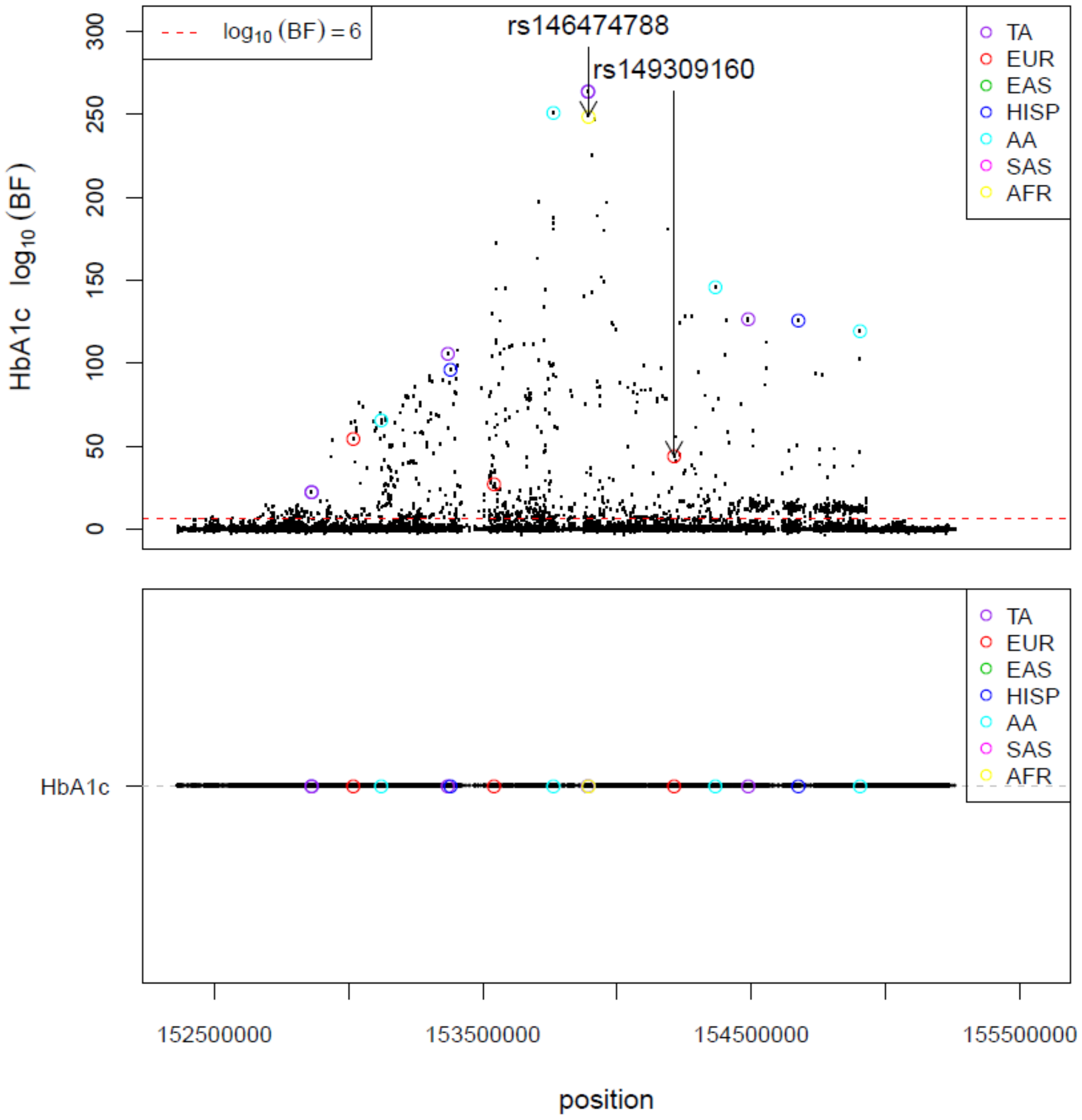

**Supplementary Figure N22** - Locus plot of the trans-ancestry lead variants and single-ancestry lead variants for HbA1c identified at locus 242, which spans over 3 Mb on Chromosome X. TA- trans-ancestry; EUR- European; EAS – East Asian; HISP – Hispanic; AA- African American; SAS- South Asian; AFR- African, specifically Ugandan.

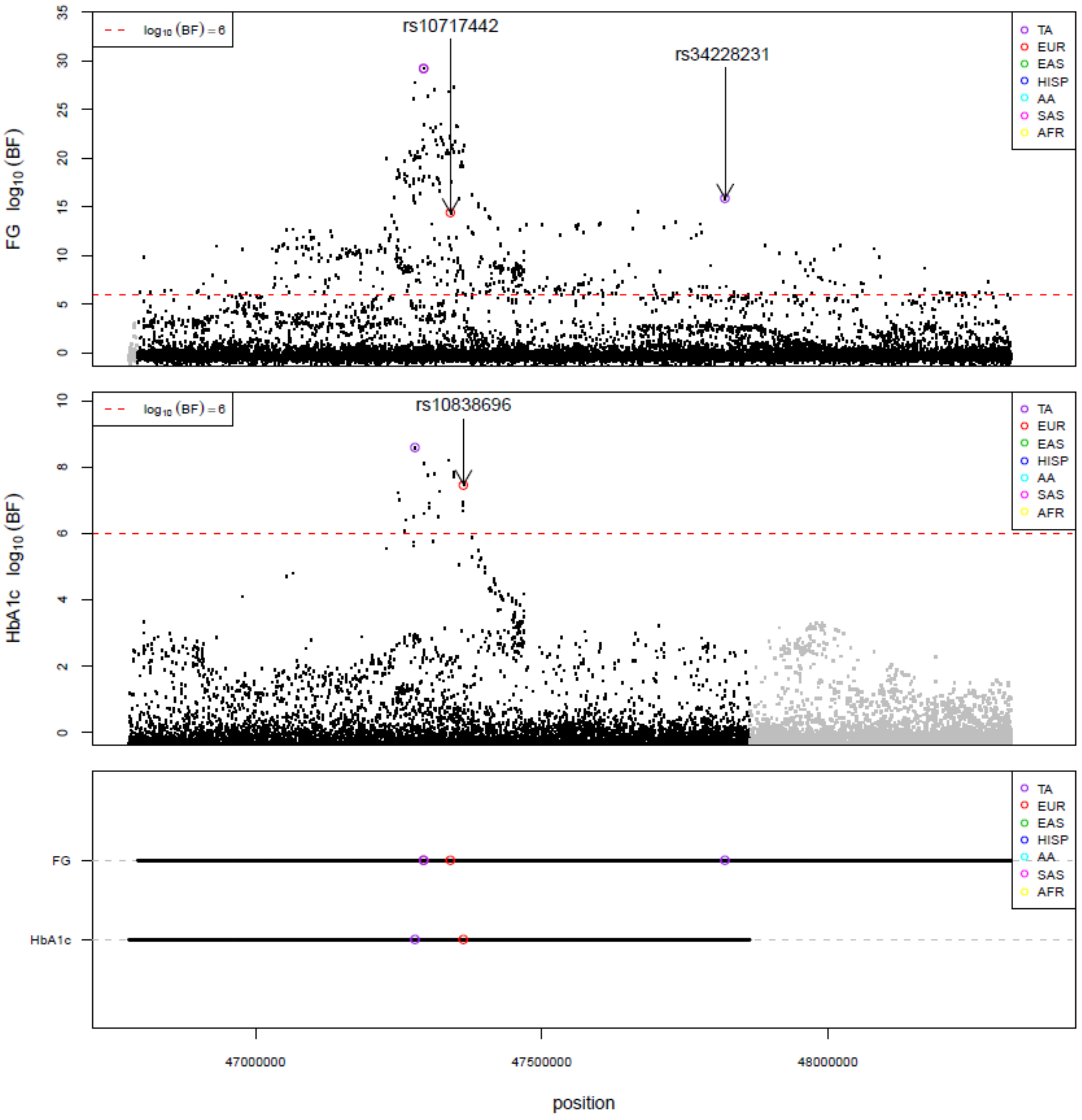

**Supplementary Figure N23 –** Locus plot of the trans-ancestry lead variants and single-ancestry index variants for FG (top) and HbA1c (bottom) identified at locus 149, which has the longest maximum distance between adjoining association variants of both the same and different traits with overall length of 1.5 Mb. TA- trans-ancestry; EUR- European; EAS – East Asian; HISP – Hispanic; AA- African American; SAS- South Asian; AFR- African, specifically Ugandan.

#### c. Definition of novel locus

Of the 242 identified loci, 99 had not been previously associated with any of the four glycaemic traits or type 2 diabetes at the time of first analysis (November 2017; **Supplementary Table 3**; lookups in more recent T2D association studies are reported in the main text and **Supplementary Table 4**). Loci were considered novel for a specific trait if no trait-associated signal within the locus mapped within 500 kb of a previously reported association for any glycaemic trait[1-3] or variants mapping to established type 2 diabetes[4,5] loci at the time of first analysis (November 2017). However, we acknowledge that some of these "novel" loci may in fact be due to established signals that map outside the 500Kb flanking regions.

##### d. Contribution of non-European ancestry data to locus discovery

Sixteen of the novel loci have *P* > $10^{-5}$ in the European-only meta-analyses which comprises the largest fraction of the data, suggesting their discovery was enabled by the power of the additional non-European samples. To test this hypothesis, we investigated each of the 242 loci assuming the sample size of the trans-ancestry analysis had been achieved in the European data alone. To do this, we scaled the standard error from the European analysis by multiplying the standard error by the square root of the ratio of the sample size from trans-ancestry analysis and the European analysis (**Supplementary Table N24**). We identified a total of 30 loci (21 novel; **Supplementary Table 3**) that were detected in the trans-ancestry meta-analyses that would not have achieved genome-wide significance in a similarly sized dataset of only European samples, highlighting the importance of diverse ancestries for novel locus discovery (**Supplementary Table N24).** For all 30 loci, their discovery in the trans-ancestry set is due to either higher EAFs or larger effect sizes in non-European populations **(Supplementary Table N25).**

**Supplementary Table N24 –** Summary of loci detected in trans-ancestry meta-analysis that would not have achieved genome-wide significance ($log_{10}BF > 6$) if the sample size had been comprised only of European ancestry individuals. Note that there is one overlapping locus (22) between FG and HbA1c so overall there are 30 loci detected due to contribution from non-European ancestry samples. "# TA loci" shows the number of loci that are associated with the trait in the trans-ancestry meta-analysis $log_{10}BF > 6$, "# TA loci with $log_{10}BF_{EUR} \leq 6.0$" shows the number of loci with $log_{10}BF > 6$ in trans-ancestry meta-analysis but $log_{10}BF \leq 6$ in European meta-analysis, "# TA loci with $log_{10}BF_{EUR} \leq 6.0$ when using TA sample size" shows the number of loci that are genome-wide significant in trans-ancestry analysis ($log_{10}BF > 6$) that would not have reached genome-wide significance $log_{10}BF > 6$ in European meta-analysis mimicking the same sample size used in trans-ancestry meta-analysis.

| Trait | # TA loci | # TA loci with $log_{10}BF_{EUR} \leq 6$ | TA loci with $log_{10}BF_{EUR} \leq 6.0$ when using TA sample size |
|---|---|---|---|
| FG | 100 | 18 | 8 |
| 2hGlu | 21 | 5 | 4 |
| FI | 62 | 18 | 8 |
| HbA1c | 126 | 32 | 11 |

**Supplementary Table N25 –** Results for 30 loci that were detected in the trans-ancestry meta-analyses that would not have achieved genome-wide significance in a similarly sized dataset of only European samples, highlighting the importance of diverse ancestries for novel locus discovery. Abbreviations: BF, Bayes factor; bp, base pair; EAF, effect allele frequency; TA, trans-ancestry.

| Trait | Locus ID | Lead TA variant | Closest Gene(s):Distance to closest gene | European | | Non-European Ancestry(s) with $log_{10}BF$ > 6.0 | | |
|---|---|---|---|---|---|---|---|---|
| | | | | **EAF** | **Effect size** | **Ancestry** | **EAF** | **Effect size** |
| *Fasting glucose* | | | | | | | | |
| | 1 | rs12142172 | *PRDM16*: 0 | 0.20 | -0.008 | AA | 0.46 | -0.021 |
| | 22 | rs12712928 | *SIX3*: 18,864 | 0.16 | 0.010 | EAS | 0.40 | 0.040 |
| | 32 | rs7572235 | *EPHA4*: 214,144 | 0.78 | -0.008 | SAS | 0.67 | -0.023 |
| | 47 | rs189651013 | *FGF12*: 0 | 0.007 | 0.110 | HISP | 0.002 | 0.150 |
| | 67 | rs3733977 | *FBLL1*: 0 | 0.16 | 0.010 | EAS | 0.49 | 0.007 |
| | 114 | rs60405463 | *KANK1*: 0 | 0.09 | 0.009 | EAS | 0.53 | 0.018 |
| | 182 | rs10781829 | NA | 0.95 | -0.016 | AA | 0.62 | -0.036 |
| | 187 | rs182584439 | *PTGDR*: 12,851 | NA | NA | AA | 0.006 | 0.233 |
| *2 hour glucose* | | | | | | | | |
| | 69 | rs34499031 | *CDKAL1*: 0 | 0.72 | -0.038 | AA<br>EAS<br>HISP | 0.41<br>0.53<br>0.69 | -0.081<br>-0.111<br>-0.072 |
| | 129 | rs35696875 | *HKDC1*: 0; *LOC101928994*: 0 | 0.32 | -0.039 | AA<br>HISP | 0.75<br>0.50 | -0.132<br>-0.066 |
| | 198 | rs115880135 | *PEX11A*: 0 | NA | NA | HISP | 0.9991 | 1.838 |
| | 237 | rs184389108 | *RRP7A*: 0; *SERHL:0* | 0.99 | -0.308 | HISP | 0.993 | -0.618 |
| *Fasting insulin* | | | | | | | | |
| | 25 | rs2252867 | *CEP68*: 0 | 0.64 | 0.007 | SAS | 0.58 | 0.015 |
| | 81 | rs5875762 | *FOXP4*: 0 | 0.31 | -0.008 | EAS | 0.31 | -0.011 |
| | 128 | rs10761762 | *JMJD1C*: 0 | 0.51 | 0.008 | AA | 0.69 | 0.023 |
| | 135 | rs7071062 | *MIR5694*: 123,574 | 0.97 | 0.019 | SAS | 0.96 | 0.084 |
| | 155 | rs3781926 | *PDE2A*: 0 | 0.36 | 0.009 | AA | 0.34 | 0.022 |
| | 163 | rs12369443 | *PDE3A*: 0 | 0.78 | 0.010 | SAS | 0.92 | 0.036 |
| | 170 | rs73343765 | *SYT1*: 100,219 | 0.006 | -0.190 | HISP | 0.003 | 0.327 |
| | 224 | rs339525 | *MAP3K10*: 1,137 | 0.26 | -0.007 | EAS | 0.31 | -0.022 |
| *HbA1c* | | | | | | | | |
| | 22 | rs12712928 | *SIX3*: 18,864 | 0.16 | 0.008 | EAS | 0.39 | 0.023 |
| | 48 | rs9846651 | *LINC00885*: 0 | 0.11 | 0.007 | EAS | 0.49 | 0.017 |
| | 53 | rs139577195 | *LOC101927282*: 267,587 | 0.0005 | -0.069 | HISP | 0.005 | -0.150 |
| | 57 | rs13164333 | *MIR4278*: 74,681 | 0.05 | 0.001 | AA | 0.14 | 0.069 |
| | 63 | rs144559191 | *AQPEP*: 0 | NA | NA | AA | 0.01 | -0.175 |
| | 91 | rs137954340 | *AGR2*: 5,617 | NA | NA | AFR | 0.008 | 0.237 |
| | 139 | rs73388897 | *OR51E2*: 0 | 0.0006 | -0.178 | HISP | 0.01 | -0.118 |
| | 140 | rs77121243 | *HBB*: 0 | 0.003 | -0.081 | AA<br>HISP | 0.07<br>0.02 | -0.108<br>-0.225 |
| | 141 | rs116006800 | *OR52N2*: 6,041 | NA | NA | AA | 0.02 | -0.145 |
| | 196 | rs114189680 | *ADAMTS7*: 0 | 0.006 | 0.041 | AA | 0.03 | -0.143 |
| | 228 | rs6113722 | *LINC00261*: 0 | 0.04 | -0.012 | EAS | 0.16 | -0.015 |

### 2. Allelic architecture of glycaemic traits

#### a. Complexity of association signals at a locus

We next examined the complexity of association signals within each locus and found that 29 loci harbour multiple, distinct single-ancestry index variants (**Supplementary Table N26**), with some having quite complex association patterns.

**Supplementary Table N26** - Table showing the distribution of single-ancestry index and lead variants per locus by trait and ancestry. Loci regions IDs map to those in Supplementary Table 2.

| Trait | Ancestry | # of loci | Minimum # of single-ancestry index and lead variants at a locus | Median # of single-ancestry index and lead variants at a locus | Maximum # of single-ancestry index and lead variants at a locus | Loci regions with > 1 signal |
|---|---|---|---|---|---|---|
| *Fasting glucose* | | | | | | |
| | EUR | 68 | 1 | 1 | 14 | 1, 25, 28, 68, 80, 90, 93, 103, 116, 133, 149, 156 |
| | EAS | 11 | 1 | 1 | 4 | 28, 90 |
| | HISP | 7 | 1 | 1 | 2 | 28 |
| | AA | 6 | 1 | 1 | 1 | NA |
| | SAS | 4 | 1 | 1 | 4 | 28 |
| | TA | 100 | 1 | 1 | 2 | 28, 149 |
| *2h glucose* | | | | | | |
| | EUR | 14 | 1 | 1 | 1 | NA |
| | HISP | 1 | 1 | 1 | 1 | NA |
| | TA | 21 | 1 | 1 | 1 | NA |
| *Fasting insulin* | | | | | | |
| | EUR | 36 | 1 | 1 | 3 | 27, 35, 59, 79, 82, 103, 172 |
| | HISP | 3 | 1 | 1 | 1 | NA |
| | AA | 1 | 1 | 1 | 1 | NA |
| | TA | 62 | 1 | 1 | 1 | NA |
| *HbA1c* | | | | | | |
| | EUR | 77 | 1 | 1 | 5 | 15, 28, 44, 54, 72, 93, 107, 129, 166, 184, 207, 216, 222, 242 |
| | EAS | 19 | 1 | 1 | 3 | 28 |
| | HISP | 9 | 1 | 1 | 4 | 242 |
| | SAS | 2 | 1 | 1 | 1 | NA |
| | AFR | 2 | 1 | 1 | 1 | NA |
| | TA | 126 | 1 | 1 | 4 | 54, 242 |

#### b. Detection of previously established loci/signals

We compared the current discovery effort against previously established glycaemic and type 2 diabetes associated signals for each trait. Loci were considered novel for a specific trait if no trait-associated signal within the locus mapped within 500 kb of a previously reported association for any glycaemic trait[1-3] or variants mapping to established type 2 diabetes[4,5] loci at the time of first analysis (November 2017). Overall, we identified novel loci for each trait in both the single-ancestry and trans-ancestry meta-analyses: 53 FG, 49 FI, 11 2hGlu, and 62 HbA1c (**Supplementary Table N27**), and identified 70-88% of previously established signals at genome-wide significance ($P < 5x10^{-8}$) (**Supplementary Table 22)**. However, there were 44 previously established signals that did not reach genome-wide significance in our analysis (i.e. did not reach BF > 6 in the trans-ancestry analysis or did not achieve $P < 5x10^{-8}$ threshold in any of the single-ancestry meta-analyses).

**Supplementary Table N27 -** Table summarising the number of known and novel loci and number of signals detected in this effort by trait.

| Trait | # of signals | # of loci | # of novel loci |
|---|---|---|---|
| FG | 182 | 102 | 53 |
| 2hGlu | 28 | 21 | 11 |
| FI | 95 | 66 | 49 |
| HbA1c | 218 | 127 | 62 |

To investigate why established signals were not observed in our analyses, we performed a lookup of all established signals in our current effort (**Supplementary Table 22**) and summarised the results in

**Supplementary Table N28** below. Overall, the vast majority of previously reported association signals are at least nominally significant ($P < 0.05$ or $\log_{10}BF > 0$) for the corresponding trait in our analysis (n = 290). The remaining seven established signals were not observed in our study for the following reasons: (i) rs6947345 was previously identified to be female-specific [FG[6]] and our analyses were limited to sex-combined models; (ii) two variants did not pass our QC stage (rs141203811, FI and rs1135071, HbA1c); (iii) rs7077836 was previously associated with FI in a smaller sample of 1,497 African American and African samples [7] but is not observed in our analysis with over 8,101 African American samples ($P$=0.25) suggesting this prior association could be a false-positive; (iv) rs1421085 was previously associated with FI without BMI adjustment, suggesting its association with FI is due to an effect on BMI (rs1421085 has been shown to be significantly associated with BMI, $P = 8.83 \times 10^{-151}$ [8]); (v) rs213676 was previously detected in a FI analysis of 14,043 African American participants [9] but not in our smaller analysis of 1,692 African American participants who contributed data for this variant (chr X), suggesting our analysis had less power to detect the association; and (vi) rs146779637 is a rare protein truncating variant in *G6PC2* (HbA1c [3]) for which we only had data on 22,617 European ancestry participants.

**Supplementary Table N28 –** Examination of the associations of established signals in our current analyses.

| Trait | Total | $P \le 5\times10^{-8}$ or $\log_{10}BF \ge 6$ | $5\times10^{-8} < P \le 5\times10^{-4}$ or $2 \le \log_{10}BF < 6$ | $5\times10^{-4} < P \le 5\times10^{-2}$ or $0 \le \log_{10}BF < 2$ | $P > 5\times10^{-2}$ or $\log_{10}BF < 0$ |
|---|---|---|---|---|---|
| FG | 102 | 90 | 8 | 3 | 1 |
| 2hGlu | 28 | 24 | 4 | 0 | 0 |
| FI | 43 | 30 | 6 | 3 | 4 |
| HbA1c | 124 | 109 | 10 | 3 | 2 |

#### c. Collider bias

There have been previous concerns regarding the possible effect of adjusting phenotypes for correlated heritable traits, leading to possible collider bias[10]. As we have conducted analyses of FG, FI, and 2hGlu adjusted for BMI, we investigated the possibility that our results were due to collider bias (i.e. that they were due to association with BMI only, and not with the trait in question). To evaluate this, we focused on all trans-ancestry lead variants and European index and lead variants (as these comprised the larger datasets and we had previous data available for traits adjusted and unadjusted for BMI) where the European meta-analysis results for FG adjusted for BMI (FGadjBMI), FI adjusted for BMI (FIadjBMI), and 2hGlu adjusted for BMI (2hGluadjBMI) reached $P \le 1\times10^{-5}$. We compared the effect size and significance of association between the variants and each of these traits with association results for the same variant and BMI. Where signals had evidence of potential collider bias [i.e. they were significantly associated with BMI ($P \le 0.05$ after Bonferroni correction) but in the opposite direction to that of their effect on the glycaemic trait], we assessed their association in analyses of glycaemic traits without adjustment for BMI. Results from unadjusted traits were available from previous MAGIC efforts, including results based on Metabochip analysis for FG, FI, and 2hGlu[11], GWAS data for FG and FI[12,13] and 2hGlu[14], and unpublished results from MAGIC (https://www.magicinvestigators.org/). BMI data was obtained from publicly available results in the UK Biobank (http://www.nealelab.is/uk-biobank/). If variants were available in both Metabochip and GWAS datasets of traits without BMI adjustment, we used results from the larger sample size to achieve greater power; if variants were missing in data for unadjusted traits or BMI, we used LD proxy variants (EUR LD $r^2 > 0.8$).

Results from these analyses demonstrated that the vast majority of the signals (85.6%) have no evidence of collider bias (**Supplementary Table N29)**. However, 36 signals associated with glycaemic traits adjusted for BMI were also significantly ($P \leq 2.1\times10^{-4}$) associated with BMI with opposite directions of effect, suggesting they may result from potential collider bias. Of the eight 2hGlu-associated signals with potential collider bias, all were nominally associated ($P \leq 0.05$) with 2hGlu unadjusted for BMI and six had $P \leq 0.00625$ (Bonferroni corrected $P = 0.05/8$). Of the 28 FG- and FI-associated signals with potential collider bias, 25 were nominally associated with FG or FI ($P \leq 0.05$) unadjusted for BMI. For these 25 variants, we tested the difference in effect size before and after BMI adjustment using the same data resource[13]. Effect sizes for 23 of 25 were not significantly different ($P > 0.002$, Bonferroni corrected $P = 0.05/25$) (**Supplementary Table N30 and Supplementary Figure N24**). These results suggested that of the 240 signals we were able to test for collider bias, at most four signals have some evidence of collider bias (two 2hGlu signals, one FG signal, and one FI signal).

**Supplementary Table N29 –** Of the 250 signals, 243 had available BMI association results. Table shows the total number of signals associated with 2hGlu, FG, and FI adjusted for BMI from this study (2nd column), the number of signals with missing data from the BMI analysis (3rd column), the number of signals with association results for BMI but where the association with BMI does not meet significance ($P > 2.1\times10^{-4}$, 4th column), the number of signals associated with BMI at $P \leq 2.1\times10^{-4}$ with the same direction of effect as the glycaemic trait (5th column), the number of signals associated with BMI at $P \leq 2.1\times10^{-4}$ but with opposite directions of effect as the glycaemic trait (6th column)

| **Trait** | **Total** | **Missing** | **BMI assoc $P > 2.1\times10^{-4}$** | **$P \leq 2.1\times10^{-4}$ with glycaemic trait and BMI; same effect direction** | **$P \leq 2.1\times10^{-4}$ with glycaemic trait and BMI; opposite effect direction** |
|---|---|---|---|---|---|
| **2hGluadjBMI** | 24 | 1 | 15 | 0 | 8 |
| **FGadjBMI** | 147 | 5 | 121 | 11 | 10 |
| **FIadjBMI** | 79 | 1 | 57 | 3 | 18 |

**Supplementary Table N30 – Comparison of effect sizes for variants with suspected collider bias in glycaemic traits adjusted and unadjusted for BMI.** Paired difference test was used to detect differences in the effect sizes. The genetic correlation between models with and without BMI adjustment is 0.9257 ($P$ = $3.3\times10^{-527}$) for FG and 0.7043 ($P$ = $2.8\times10^{-68}$) for FI based on the same data[13] using LD score regression[15,16]. Signals with difference test $P \leq 0.002$ are in bold.

| Trait | rsID | EA | OA | Proxy | $r^2$ | Without BMI adjustment | | | With BMI adjustment | | | $P_{eff\ diff}$ |
|---|---|---|---|---|---|---|---|---|---|---|---|---|
| | | | | | | Effect | SE | *P* | Effect | SE | *P* | |
| *Fasting glucose* | | | | | | | | | | | | |
| | rs1604038 | T | C | - | - | -0.020 | 0.003 | $2\times10^{-9}$ | -0.023 | 0.004 | $2.3\times10^{-11}$ | 0.024 |
| | rs1635852 | T | C | - | - | 0.005 | 0.003 | 0.13 | 0.008 | 0.003 | 0.012 | 0.0074 |
| | rs1820176 | T | C | rs7713317 | 0.96 | 0.016 | 0.003 | $1.8\times10^{-6}$ | 0.020 | 0.004 | $7.6\times10^{-9}$ | 0.0027 |
| | rs2238435 | C | G | rs879620 | 1.00 | -0.012 | 0.003 | 0.00079 | -0.013 | 0.003 | 0.00017 | 0.45 |
| | rs2657879 | A | G | - | - | -0.013 | 0.004 | 0.0017 | -0.017 | 0.004 | 0.000053 | 0.014 |
| | rs34872471 | T | C | rs7903146 | 0.99 | -0.021 | 0.004 | $1.1\times10^{-9}$ | -0.025 | 0.004 | $9.5\times10^{-13}$ | 0.0036 |
| | **rs3764400** | **T** | **C** | **-** | **-** | **0.010** | **0.005** | **0.039** | **0.017** | **0.005** | **0.00089** | **0.00023** |
| | rs6876986 | C | G | rs10476552 | 1.00 | -0.017 | 0.003 | $5.9\times10^{-7}$ | -0.020 | 0.003 | $6.4\times10^{-9}$ | 0.022 |
| | rs7903146 | T | C | - | - | 0.021 | 0.004 | $1.1\times10^{-9}$ | 0.025 | 0.004 | $9.5\times10^{-13}$ | 0.0036 |
| *Fasting insulin* | | | | | | | | | | | | |
| | rs1023667 | A | G | - | - | -0.003 | 0.004 | 0.00044 | -0.006 | 0.003 | 0.037 | 0.16 |
| | rs1128249 | T | G | - | - | -0.013 | 0.003 | 0.000032 | -0.018 | 0.003 | $7.1\times10^{-11}$ | 0.031 |
| | rs12454712 | T | C | - | - | 0.018 | 0.005 | 0.00069 | 0.020 | 0.005 | 0.000013 | 0.61 |
| | rs12541800 | A | G | - | - | 0.003 | 0.003 | 0.31 | 0.009 | 0.003 | 0.00091 | 0.012 |
| | rs13234269 | A | T | - | - | -0.011 | 0.003 | 0.00048 | -0.012 | 0.003 | 0.000015 | 0.67 |
| | rs13389219 | T | C | - | - | -0.013 | 0.003 | 0.000033 | -0.018 | 0.003 | $7.2\times10^{-11}$ | 0.031 |
| | rs330945 | T | C | rs330944 | 0.95 | -0.009 | 0.008 | 0.29 | -0.005 | 0.007 | 0.47 | 0.56 |
| | rs7133378 | A | G | - | - | -0.004 | 0.003 | 0.26 | -0.008 | 0.003 | 0.0089 | 0.12 |
| | rs75265117 | C | G | rs12328675 | 0.99 | 0.020 | 0.005 | 0.000034 | 0.029 | 0.004 | $1.5\times10^{-12}$ | 0.0098 |
| | rs7654571 | A | G | - | - | 0.002 | 0.004 | 0.59 | 0.008 | 0.003 | 0.023 | 0.054 |
| | **rs77935490** | **A** | **T** | **rs5017303** | **0.91** | **-0.012** | **0.004** | **0.0036** | **0.003** | **0.004** | **0.35** | **$2.7\times10^{-7}$** |
| | rs7975482 | A | G | - | - | 0.004 | 0.003 | 0.27 | 0.007 | 0.003 | 0.0096 | 0.12 |
| | rs848494 | A | G | - | - | 0.013 | 0.004 | 0.00041 | 0.011 | 0.003 | 0.00026 | 0.44 |
| | rs972283 | A | G | - | - | -0.012 | 0.003 | 0.00016 | -0.013 | 0.003 | 0.0000044 | 0.67 |
| | rs979012 | T | C | - | - | 0.003 | 0.003 | 0.42 | 0.008 | 0.003 | 0.0081 | 0.05 |
| | rs998584 | A | C | - | - | 0.004 | 0.004 | 0.34 | 0.006 | 0.003 | 0.043 | 0.28 |

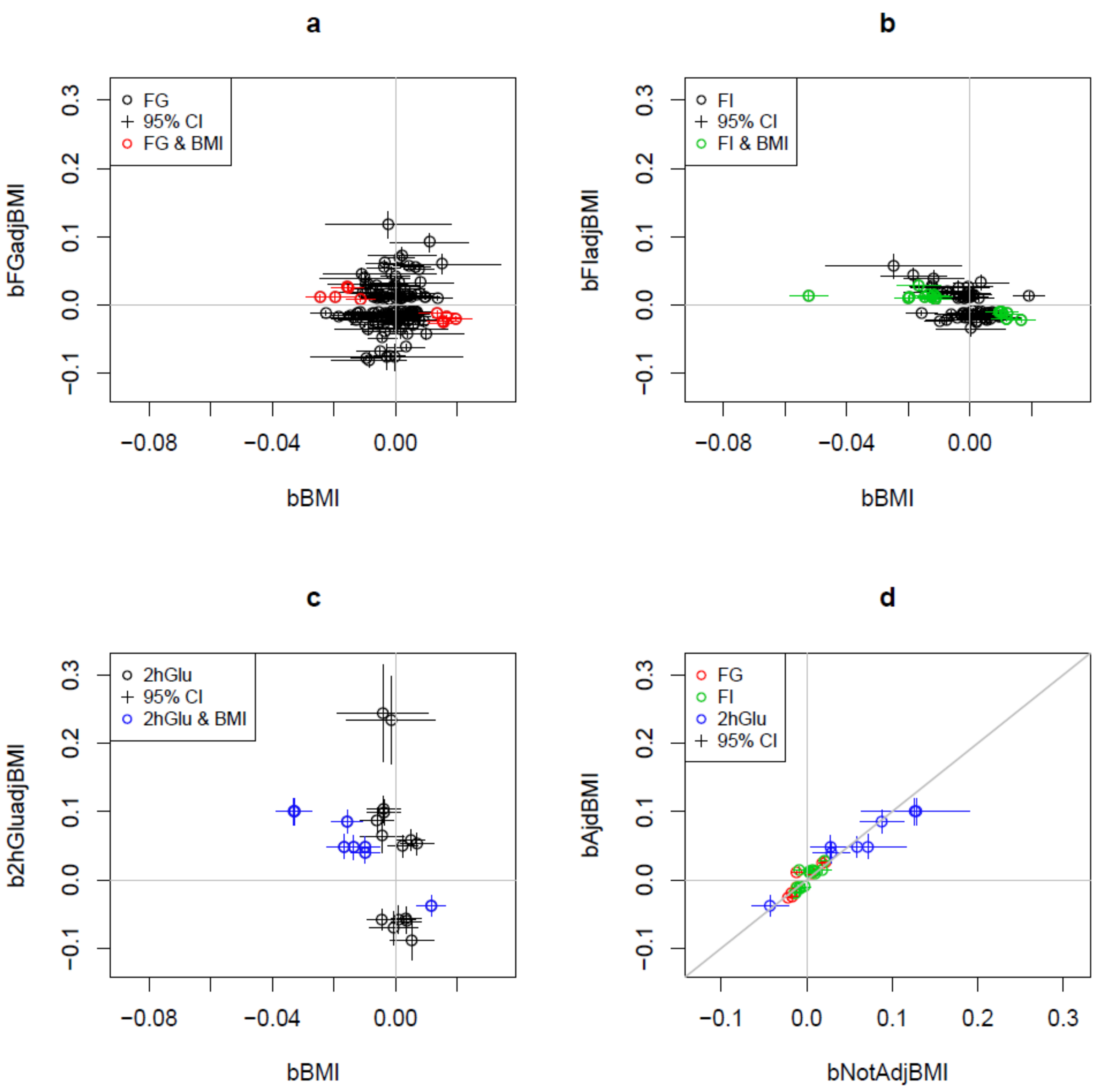

**Supplementary Figure N24 - Comparison of effect sizes between glycaemic traits and BMI for variants associated with each of the glycaemic traits.** Effect sizes are shown for signals associated with each glycaemic trait identified in trans-ancestry and European meta-analyses. Error bars are the 95% confidence intervals. **a**. Effect sizes for BMI (*x*-axis) and FG adjusted for BMI (*y*-axis). Signals associated with BMI in UK Biobank at $P \leq 2.1\times10^{-4}$ in the opposite direction are highlighted in red **b**. Effect sizes for BMI (*x*-axis) and FI adjusted for BMI (*y*-axis). Signals associated with BMI in UK Biobank at $P \leq 2.1\times10^{-4}$ in the opposite direction are highlighted in green. **c**. Effect sizes for BMI (*x*-axis) and 2hrGlu adjusted for BMI (*y*-axis). Signals associated with BMI in UK Biobank at $P \leq 2.1\times10^{-4}$ in the opposite direction are highlighted in blue. **d**. Effect sizes for the 28 glycaemic trait signals with suspected collider bias (BMI association $P \leq 2.1\times10^{-4}$ and opposite directions of effect; coloured red, green, and blue in panels a, b, and c) from analyses without BMI adjustment (*x*-axis) and with BMI adjustment (*y*-axis) are shown for highlighted signals on figures a, b and c.

#### 3. Characterisation of trans-ancestry lead variants and European index variants across ancestries

To compare association signals across all ancestries, we first took the trans-ancestry lead variant and evaluated the fraction of the times the same lead variant demonstrated at least nominal evidence of association *($P \leq 0.05$)* in all available ancestries. We found that between 0.8% (HbA1c) and 16% (FG) of the lead trans-ancestry variants had supportive evidence across all ancestries. This small percentage is likely due to differences in LD across the populations and/or because the trans-ancestry lead variant may not be the best representative of the signal within each ancestry (**Supplementary Table N31**). These analyses may also be hampered by the different sample sizes across ancestries, allelic heterogeneity, and/or stochastic variation. We therefore investigated the pairwise EAF correlation between ancestries (**Methods).** This demonstrated considerable EAF correlation ($r^2 > 0.7$) between Europeans and Hispanics, Europeans and South Asians, and Hispanics and South Asians consistent across all four traits, and between African Americans and Ugandans for HbA1c. We also investigated the pairwise summarised heterogeneity of effect sizes between ancestries[17] (**Methods, Supplementary Figure 12),** and found that, despite significant EAF correlation, there was strong evidence for effect size heterogeneity among some pairwise comparisons, which was more variable between traits (**Supplementary Figure 12).** For example, for HbA1c and FI, there is strong heterogeneity of effect sizes between Europeans and Hispanics ($P < 2.63\text{x}10^{-6}$), despite high EAF correlation ($r^2 > 0.8$). Overall, there are 41 pairwise trait-ancestry comparisons, 17 of which demonstrate evidence of significant heterogeneity [$P < 0.00122$ (Bonferroni correction = 0.05/41); **Supplementary Table N32**]. However, sensitivity analyses sequentially removing signals with evidence of between-ancestry heterogeneity (up to all with $P < 0.05$), demonstrated that a relatively small number of signals (range 7-23 per trait) were responsible for the heterogeneity (**Supplementary Table N32**).

**Supplementary Table N31** - Table showing number of trans-ancestry loci per trait, as well as the number where the TA lead variant is also the lead variant in that locus across all ancestries, or the number of loci where there is at least nominal evidence of association ($P < 0.05$) for the trans-ancestry lead variant in each ancestry even if it does not represent the lead variant in a particular ancestry.

| | **FG** | **2hGlu** | **FI** | **HbA1c** |
|---|---|---|---|---|
| # of TA loci | 100 | 21 | 62 | 126 |
| # loci where TA lead is also the lead variant across all ancestries | 1 | 0 | 0 | 0 |
| # loci where TA lead *P*<0.05 in other ancestries, but not necessarily the lead variant | 15 | 1 | 4 | 1 |

**Supplementary Table N32** – Results from a sensitivity analyses sequentially removing signals with evidence of between-ancestry heterogeneity

| Trait | Ancestry Comparison | | EAF Correlation | | Effect Correlation | | # of signals | Overall $P_{het}$ | # signals with $P_{het}$ ≤ $1\times10^{-6}$ | Overall $P_{het}$ after removing signals with $P_{het}$ ≤ $1\times10^{-6}$ | # signals with $P_{het}$ ≤ 0.05 | Overall $P_{het}$ excluding signals with $P_{het}$ ≤ 0.05 |
|---|---|---|---|---|---|---|---|---|---|---|---|---|
| | 1 | 2 | r | *P* | r | *P* | | | | | | |
| *Fasting glucose* | | | | | | | | | | | | |
| | EUR | AA | 0.36 | 0.00027 | 0.63 | $1.6\times10^{-12}$ | 100 | **0.000012** | 1 | 0.0086 | 13 | 0.97 |
| | EUR | EAS | 0.36 | 0.00025 | 0.42 | 0.00002 | 98 | **$2.7\times10^{-11}$** | 2 | 0.16 | 7 | 0.85 |
| | EUR | HISP | 0.79 | $1\times10^{-22}$ | 0.71 | $9.9\times10^{-17}$ | 101 | 0.016 | 0 | 0.016 | 10 | 0.71 |
| | EUR | SAS | 0.82 | $1\times10^{-25}$ | 0.75 | $9.6\times10^{-20}$ | 101 | 0.16 | 0 | 0.16 | 7 | 0.93 |
| | EAS | AA | 0.31 | 0.002 | 0.41 | 0.000028 | 98 | 0.026 | 0 | 0.026 | 6 | 0.77 |
| | EAS | HISP | 0.58 | $4\times10^{-10}$ | 0.44 | $4.5\times10^{-6}$ | 98 | **0.00059** | 0 | **0.00059** | 10 | 0.86 |
| | EAS | SAS | 0.65 | $6.8\times10^{-13}$ | 0.37 | 0.00017 | 98 | **0.00075** | 0 | **0.00075** | 14 | 0.97 |
| | HISP | AA | 0.60 | $4.7\times10^{-11}$ | 0.87 | $1.5\times10^{-31}$ | 101 | 0.057 | 0 | 0.057 | 8 | 0.98 |
| | HISP | SAS | 0.79 | $8.3\times10^{-23}$ | 0.55 | $2.8\times10^{-9}$ | 101 | 0.032 | 0 | 0.032 | 6 | 0.55 |
| | AA | SAS | 0.48 | $4.8\times10^{-7}$ | 0.23 | 0.021 | 100 | 0.085 | 0 | 0.085 | 9 | 0.91 |
| *2h glucose* | | | | | | | | | | | | |
| | EUR | AA | 0.21 | 0.36 | -0.51 | 0.023 | 20 | 0.098 | 0 | 0.098 | 3 | 0.86 |
| | EUR | EAS | 0.19 | 0.45 | 0.63 | 0.0054 | 18 | 0.35 | 0 | 0.35 | 1 | 0.6 |
| | EUR | HISP | 0.86 | $8.8\times10^{-7}$ | 0.95 | $3.1\times10^{-10}$ | 20 | 0.84 | 0 | 0.84 | 0 | 0.84 |
| | EAS | AA | -0.03 | 0.91 | 0.35 | 0.15 | 18 | 0.083 | 0 | 0.083 | 3 | 0.74 |
| | EAS | HISP | 0.40 | 0.097 | 0.56 | 0.015 | 18 | 0.53 | 0 | 0.53 | 2 | 0.97 |
| | HISP | AA | 0.37 | 0.095 | 0.54 | 0.012 | 21 | 0.056 | 0 | 0.056 | 3 | 0.62 |
| *Fasting insulin* | | | | | | | | | | | | |
| | EUR | AA | 0.53 | $7.6\times10^{-6}$ | 0.51 | 0.000028 | 62 | 0.35 | 0 | 0.35 | 3 | 0.88 |
| | EUR | EAS | 0.34 | 0.0076 | -0.08 | 0.54 | 60 | **$1.1\times10^{-6}$** | 0 | **$1.3\times10^{-6}$** | 11 | 0.51 |
| | EUR | HISP | 0.83 | $8.4\times10^{-17}$ | -0.6 | $2.4\times10^{-7}$ | 62 | **0.00017** | 1 | 0.13 | 7 | 0.97 |
| | EUR | SAS | 0.82 | $1.5\times10^{-15}$ | 0.8 | $3.7\times10^{-14}$ | 59 | 0.3 | 0 | 0.3 | 4 | 0.86 |
| | EAS | AA | 0.09 | 0.51 | -0.21 | 0.11 | 60 | 0.22 | 0 | 0.22 | 3 | 0.71 |
| | EAS | HISP | 0.57 | $2.5\times10^{-6}$ | 0.17 | 0.21 | 60 | **0.001** | 0 | **0.001** | 7 | 0.39 |
| | EAS | SAS | 0.41 | 0.0016 | -0.11 | 0.42 | 57 | 0.014 | 0 | 0.014 | 5 | 0.83 |
| | HISP | AA | 0.59 | $5.5\times10^{-7}$ | 0.08 | 0.53 | 62 | 0.044 | 0 | 0.044 | 4 | 0.81 |
| | HISP | SAS | 0.74 | $2.8\times10^{-11}$ | 0.64 | $4.2\times10^{-8}$ | 59 | 0.62 | 0 | 0.62 | 3 | 0.98 |
| | AA | SAS | 0.54 | 0.000011 | 0.66 | $1.3\times10^{-08}$ | 59 | 0.42 | 0 | 0.42 | 3 | 0.93 |
| *HbA1c* | | | | | | | | | | | | |
| | EUR | AA | 0.41 | $1.9\times10^{-6}$ | 0.46 | $9\times10^{-8}$ | 124 | **$<2.2\times10^{-16}$** | 2 | **$1\times10^{-13}$** | 23 | 0.87 |
| | EUR | AFR | 0.21 | 0.031 | 0.41 | 0.000012 | 107 | **$8.9\times10^{-07}$** | 0 | **$8.9\times10^{-7}$** | 14 | 0.9 |
| | EUR | EAS | 0.35 | 0.000097 | -0.13 | 0.15 | 119 | **$1.6\times10^{-15}$** | 2 | **0.0001** | 15 | 0.96 |
| | EUR | HISP | 0.88 | $3.6\times10^{-42}$ | 0.52 | $5.6\times10^{-10}$ | 125 | **$2\times10^{-7}$** | 2 | 0.025 | 12 | 0.93 |
| | EUR | SAS | 0.80 | $4.4\times10^{-28}$ | 0.44 | $4.3\times10^{-7}$ | 120 | 0.017 | 0 | 0.017 | 13 | 0.95 |
| | EAS | AA | 0.37 | 0.000035 | -0.11 | 0.23 | 118 | **$9.2\times10^{-8}$** | 0 | **$9.2\times10^{-8}$** | 15 | 0.62 |
| | EAS | AFR | 0.24 | 0.015 | -0.17 | 0.094 | 103 | **0.00001** | 0 | **0.00001** | 16 | 0.87 |
| | EAS | HISP | 0.55 | $7\times10^{-11}$ | 0.06 | 0.54 | 119 | 0.0099 | 0 | 0.0099 | 8 | 0.82 |
| | EAS | SAS | 0.63 | $2.1\times10^{-14}$ | 0.37 | 0.000056 | 116 | 0.057 | 0 | 0.057 | 8 | 0.96 |
| | HISP | AA | 0.67 | $7.8\times10^{-18}$ | 0.89 | $2.6\times10^{-46}$ | 129 | **0.000002** | 0 | **0.000002** | 15 | 0.89 |
| | HISP | AFR | 0.50 | $1.7\times10^{-8}$ | 0.56 | $1.1\times10^{-10}$ | 111 | **$<2.2\times10^{-16}$** | 5 | **0.000098** | 19 | 0.97 |
| | HISP | SAS | 0.82 | $2\times10^{-30}$ | 0.4 | $4.7\times10^{-6}$ | 120 | 0.53 | 0 | 0.53 | 5 | 0.99 |
| | AA | AFR | 0.98 | $4.9\times10^{-78}$ | 0.45 | $9.4\times10^{-7}$ | 111 | **$<2.2\times10_{-16}$** | 5 | **0.000026** | 16 | 0.78 |
| | AA | SAS | 0.50 | $5.5\times10^{-9}$ | -0.21 | 0.021 | 120 | **$8.9\times10^{-6}$** | 0 | **$8.9\times10^{-6}$** | 11 | 0.34 |
| | SAS | AFR | 0.30 | 0.0018 | 0.6 | $1.2\times10^{-11}$ | 104 | 0.046 | 0 | 0.046 | 9 | 0.97 |

Next, for each trait, we undertook concordance analyses to investigate whether we observed a greater proportion of independent variants with the same direction of effect than we would expect by chance (50%) between Europeans and each other ancestry. To ensure independence of association signals, variants reported in each ancestry were LD clumped in 1 Mb windows. The variants were then partitioned into five bins of *P*-values from the European meta-analysis. We calculated the number of variants within each bin, determined the proportion of those variants with the same direction of effect between ancestries, and used a binomial test (one-sided) of excess directional concordance over that expected by chance (**Supplementary Table 6**).

Among variants with the strongest association signals for FG and FI in Europeans ($P < 5\times10^{-4}$), there is strong concordance in the direction of effect between Europeans and all other ancestry groups. The concordance becomes weaker for less significant *P*-value bins. Among variants with the strongest association signals for 2hGlu ($P < 5\times10^{-4}$ in Europeans), there is also strong concordance in the direction of effect between European and all other ancestry groups, but the excess is not significant in the binomial test, reflecting the lower power of our analyses for this trait. Results for variants with the strongest association signals for HbA1c ($P < 5\times10^{-4}$ in Europeans) show a similar pattern of concordance as for FG and FI, except when considering the direction of effects into African Americans and Ugandans.

We hypothesised that the relatively low concordance of direction of effect observed between Europeans and African ancestry groups for HbA1c might be reflecting the different pathways (glycaemic and non-glycaemic) through which variants can affect HbA1c levels, particularly effects mediated through the red blood cell where balancing selection can lead to different associations in individuals of African ancestry[1]. To investigate this assertion, we classified European lead variants attaining genome-wide significance as acting through glycaemic and/or red blood cell pathways (**see Supplementary note section 6a**). In both African American and Ugandan ancestries, we observed greater concordance of the direction of effect with Europeans for signals classified as glycaemic, although the excess in concordance was not significant, likely due to the low numbers of variants in each stratum (**Supplementary Table N33**).

**Supplementary Table N33** - Concordance in the direction of effect of independent HbA1c-associated variants between Europeans and African Americans (AA) or Ugandans (AFR). Variants are stratified according to the classification of association signals acting through glycaemic and red blood cell pathways. For each stratum, the total number of variants attaining genome-wide significance in European is presented, together with the proportion of those variants with the same direction of effect. For example, there are 95 distance-clumped variants with $P < 5\times10^{-8}$ in EUR, of which 58 (61%) have the same effect direction between EUR and AA. We can find 77 of the 95 in LD (EUR $r^2 > 0.8$) with signals included in the HbA1c signal classification, of which 60 are in the red blood cell cluster (soft) and 17 are in the glycaemic pathway. The binomial test *P*-value for excess concordance is also presented for the glycaemic stratum of variants.

| **Ancestry** | **Distance-clumped signal** | **Classification proxy ($r^2 > 0.8$)** | **Red blood cell pathway** | **Glycaemic pathway** | ***P*-value of proportion test** |
|---|---|---|---|---|---|
| AA | 58/95 (0.61) | 48/77 (0.62) | 36/60 (0.6) | 12/17 (0.71) | 0.43 |
| AFR | 57/85 (0.67) | 45/67 (0.67) | 35/55 (0.64) | 10/12 (0.83) | 0.19 |

#### 4. Transferability of genetic scores (GSs) across ancestries

For each trait, we next assessed the utility of a genetic score (GS) by building them from the effect sizes at lead and index variants obtained from the EUR analysis and applying them to other ancestries. To provide a benchmark for the variance explained of the EUR GS, we performed two parallel analyses. For the first analysis, we created an effect-size weighted GS from the European meta-analysis and applied it to four cohorts of EUR ancestry that were not included in the meta-analysis and had individual-level phenotype and genotype data available: WHITEHALL II, Oxford Biobank, VIKING, and UKHLS. Prediction into these independent cohorts is a fairer comparison when assessing prediction into other non-European ancestry cohorts as it provides the baseline of how well these scores work in cohorts of the same ancestry that did not contribute to the initial discovery of GS variants. In each cohort, we obtained the trait variance explained ($R^2$) from a linear regression model of the trait on the GS. In the second analysis, we applied the same effect-size weighted GS from the EUR meta-analysis to EUR-ancestry cohorts that were included in the meta-analysis using available cohort-level association summary statistics. These analyses were restricted to cohorts with at least 1,000 individuals. To reduce over-fitting, we removed the contribution of the cohort from the EUR meta-analysis by generating adjusted effect sizes[18]. Cohort-level association summary statistics were then used to estimate the trait variance explained using the EUR GS weighted with adjusted effect sizes with the R package "gtx"[19]. We then tested the EUR GS in other ancestry groups by obtaining $R^2$ of the trait explained on the basis of single-ancestry meta-analysis summary statistics using the R package "gtx".

We considered different sets of variants contributing to the GS for each trait on the basis of their evidence for association in the European meta-analysis. For the first set, we considered an index variant for each distinct association signal (additional LD pruning of $r^2 < 0.1$, retaining the variant with the minimal *P*-value; "index variants"). For the second set, we expanded this list to include additional distance clumped variants with $P \leq 1\times10^{-5}$ ("$P \leq$ 1E-5"). In the final set, we extended the list to include additional distance clumped variants with $P \leq 0.05$ ("$P \leq 0.05$"). We also further evaluated each of these GS after removing variants with evidence for heterogeneity in effect sizes between ancestries: removing signals with strong evidence of heterogeneity ($P \leq 1\times10^{-6}$); and all signals even with modest evidence of heterogeneity ($P < 0.05$). **Supplementary Table 8** presents trait-specific summaries of the variance explained of each European GS into each ancestry group. In Europeans, the range of $R^2$ obtained across tested cohorts is presented, whilst in other ancestry groups, the $R^2$ obtained from the single-ancestry meta-analysis is presented.

Several key observations are shared across the four traits. First, even within European ancestry cohorts, the variance explained of the European GS is variable, most likely reflecting differences in sample size or LD differences even within European populations. Second, the variance explained of the European GS in European ancestry cohorts increases when variants modestly associated with the trait ($P \leq 0.05$) from the EUR meta-analysis are included, irrespective of the extent of heterogeneity. Third, for the GS based only on index variants from the European meta-analysis, the variance explained in non-European ancestries is generally within the range of $R^2$ observed in the European ancestry cohorts. However, GS expanded to signals with "$P \leq 1\times10^{-5}$" perform much worse into other ancestries than a GS with just the genome-wide significant signals (**Supplementary Table 8).**

#### a. Trait variance explained by associated loci

To determine how much of the phenotypic variance of each trait could be explained by the trait-associated loci identified in the GWAS, variants were combined in a weighted GS. The association between the GS and traits was tested in a linear regression framework both in cohorts included in the discovery GWAS and in a smaller number of independent cohorts. The cohorts that contributed to this analysis are identified in the **Supplementary Table 1**.

*Identification of variants to include in GS:* GS were generated using up to three different variant lists for each trait and ancestry: 1) Single-ancestry only - single-ancestry index and lead variants selected by the approximate conditional analysis in GCTA and LD-pruned (by ordering the variants from most to least significant and keeping each subsequent variant if the ancestry LD $r^2 < 0.1$); 2) Single-ancestry plus trans-ancestry - variants from the single-ancestry list plus trans-ancestry lead variants that achieved $P < 1x10^{-5}$ within that ancestry and LD-pruned (by ordering the trans-ancestry variants from most to least significant and keeping each subsequent variant if the LD $r^2 < 0.1$ between itself and any of the single-ancestry variants or trans-ancestry variants already included); 3) Complete list – all trans-ancestry lead variants based on the MANTRA results that have $P < 0.1$ in the given ancestry, plus all single-ancestry lead and index variants that are not in LD with the trans-ancestry variants (LD $r^2 < 0.1$). In all cases, *P*-values were taken from the inverse normal analysis and betas from the analysis of raw (or log transformed in the case of FI) trait values within the relevant ancestry. LD was estimated from the collected cohort pairwise LD information, where available, or from the European samples in 1000G Phase 3. A list of the variants used to make the GS can be found in **Supplementary Table 7**. Betas were extracted for these variants from the single-ancestry GWAS analysis of the raw traits (or log transformed in the case of FI).

*Adjustment of betas for non-independent cohorts:* To obtain unbiased estimates of the variance explained by the GS, the cohort for the variance explained analysis should be independent of the GWAS sample. However, this was not practical in this case since the majority of cohorts with the genotypic and phenotypic data required for the analysis were included in the discovery GWAS. Therefore, in the case of the European ancestry cohorts, we employed the method of Nolte *et al.*[18] to adjust the effect sizes (betas) from the GWAS for the contribution of each cohort, providing sets of cohort-specific effect sizes that were then used to generate the GS. This adjustment involves recalculating the variant's effect sizes and standard errors using inverse versions of the formula for an inverse-variance fixed-effects meta-analysis. We used a pre-release version of the R package MetaSubtract [https://cran.r-project.org/web/packages/MetaSubtract/index.html] to carry out this adjustment. All variants in the initial list for that ancestry and trait were retained, regardless of whether the recalculated *P*-value reached the significance threshold used in the GWAS; therefore, the list of variants contributing to the GS in each cohort were the same (subject to exclusions during cohort-level quality control). No such adjustment was made in the case of the non-European ancestries because each cohort contributed a relatively large proportion of samples to the single-ancestry GWAS (up to 75%) as opposed to the European cohorts each of whose contribution was generally less than 5%. This meant that in the case of the non-European ancestries, the adjusted betas would have been very imprecise. No adjustment of the effect sizes was required in the independent cohorts (i.e. those which did not contribute data to the discovery GWAS).

*Cohort-level quality control (QC) of variants:* Variant-level QC was applied to variants in the GS lists within each cohort and in-line with QC criteria applied in the GWAS. Imputed variants were excluded if the imputation quality was $r^2 < 0.4$ (for minimac/University of Michigan imputation) or INFO < 0.4 (for IMPUTE/Sanger imputation). Genotyped variants were excluded if they had a call rate < 95%.

*Calculation of GS:* GS were generated in Plink v1.9 using the --score function and dosage format genotype data[20]. The --sum and --double-dosage options were employed such that the betas were multiplied by diploid allele counts and summed across all loci. Betas used were either those taken directly from the GWAS results (in the case of non-European ancestry cohorts and all independent cohorts) or those adjusted as described above (for European ancestry cohorts included in the discovery meta-GWAS).

*Calculation of variance explained:* Phenotypes used were defined as described in the discovery GWAS (with the same sample exclusion criteria applied). A linear model was fit with the raw trait (or natural log transformed in the case of FI) as the dependent variable and the GS as the predictor (or independent variable). To determine the percentage of variance in the trait explained by the GS, we extracted the adjusted $R^2$ from the model. Where cohorts included related individuals, relatives were either excluded or appropriate adjustments made. Results are presented for between eight (2hGlu) to 27 (FG) cohorts (**Supplementary Tables 9-12).**

Our results showed the expected increase in variance explained relative to earlier estimates by the same methodology but with fewer contributing variants[18]. However, previously reported variance explained estimates by Scott et al[11] of 4.8%, 1.2%, and 1.7% for FG (36 variants), FI (19 variants), and 2hGlu (9 variants), respectively, are in excess of our estimates. We hypothesise that this is likely to be at least partly attributable to a difference in statistical approaches. Therefore, we also estimated the variance explained using the approach in which genotypes for the trait-associated loci are fit directly in the linear model (as opposed to generating a GS). This analysis was performed in two cohorts: ALSPACmothers and FENLAND-OMICS. The variance explained by this method tended to be higher than by the GS method with the unadjusted $R^2$ from this model, returning a variance explained estimate up to 80% higher (ALSPACmothers, FG) than when using the GS for FG. This may be the result of over-fitting when genotypes are fitted in the model directly.

#### 5. Fine-mapping

For all 237 autosomal loci identified in this study, we performed single-ancestry and trans-ancestry fine-mapping **Supplementary Figure N25 (Methods).** Trans-ancestry lead variants from MANTRA were taken forward for the trans-ancestry effort. For the single-ancestry fine-mapping, we incuded meta-analysis summary statistics from all relevant GWAS cohorts. Variants were kept in the analysis as long as they were present in at least 90% of the samples within any given ancestry or trans-ancestry lead ones.

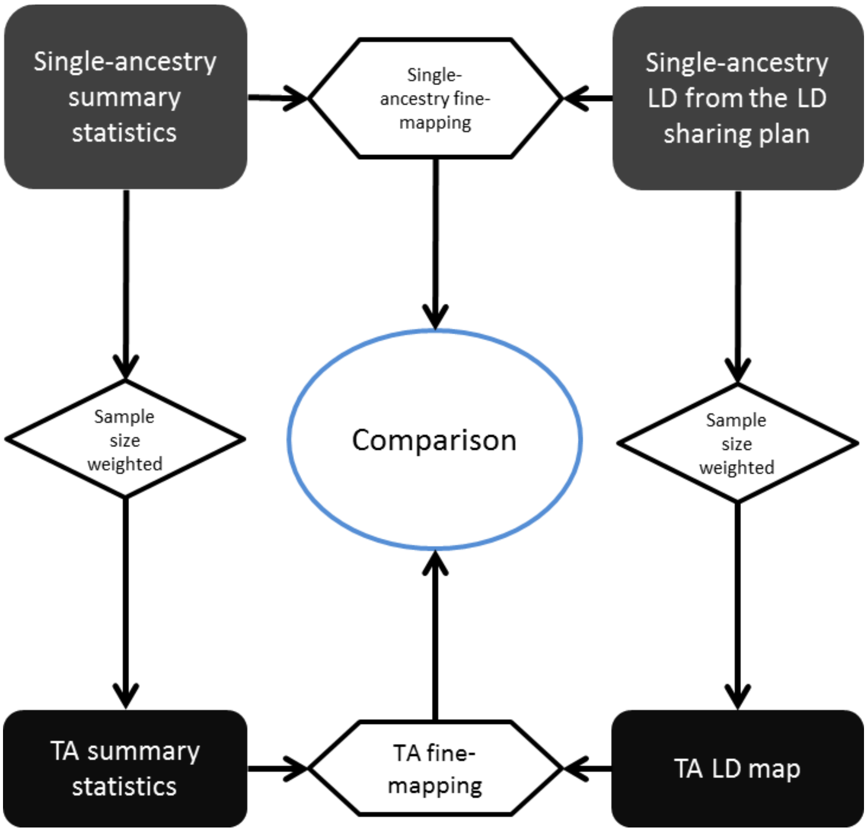

**Supplementary Figure N25 -** Flow chart depicting the pipeline used for single-ancestry and trans-ancestry fine mapping

We conducted single-ancestry and trans-ancestry fine-mapping at 237 autosome loci across 4 traits using FINEMAP to attempt to identify plausible causal variants within each locus (**Methods**). In each case, we used FINEMAP to construct 99% credible sets (99% CS), sets of variants that jointly account for 99% of the posterior probability of driving association at that locus. Credible sets containing fewer numbers of variants, and/or spanning a smaller chromosome region correspond to improved fine-mapping resolution.

To directly compare single-ancestry and trans-ancestry results, we focused on 98 loci with evidence for a single causal variant from both analyses, of which 8 had a single variant in the CS from both analyses and were excluded from the comparison. In 72/90 (80%) loci with a single causal variant, trans-ancestry fine-mapping yielded smaller 99% CS than the single-ancestry fine-mapping, reducing the size of the CS from a median of 36 variants to 20.5 variants (an average reduction of 43.1%, with a maximum reduction of 39 variants down to 1 variant, and minimum reduction of 169 variants to 167). In the remaining 18 loci (20%), fine-mapping resolution was not improved by trans-ancestry analyses (the median number of variants in the credible set was increased by 76.5%, from 8.5 variants to 15 variants). The poorer resolution was due to inconsistent directions of effect sizes across ancestries.

The improved resolution could be due to either the larger sample size available in the trans-ancestry effort, LD between ancestries, or both. To directly assess the contribution of LD differences between ancestries to improve fine-mapping resolution, we repeated the analysis emulating the same sample size in the trans-ancestry and single-ancestry fine-mapping (**Methods**). In this analysis, 47% of loci (34/72) yielded smaller credible sets in the trans-ancestry compared to the single-ancestry fine-mapping due to LD differences, reducing the size of the CS from a median of 24 to 15 variants (an average reduction of 37.5%, with a maximum reduction of 10 variants to 1, and minimum of 68 variants to 67), highlighting the importance of conducting genetic studies in diverse ancestries. In the remaining 38 loci, both the increased sample size and/or diverse LD structure contributed to the improvement.

There are several examples in which the trans-ancestry fine-mapping yielded known causal variants at established loci. At locus 22 (Chr2:44,690,056-45,692,080), the 99% CS is reduced from 5 variants (EAS) to a single variant, rs12712928 (**Supplementary Figure N26**), likely due to increased sample size in trans-ancestry fine-mapping. At locus 228 (Chr20: 22,057,099-23,067,608), the European 99% CS contained 27 variants and the East Asian 99% CS contained 23 variants. In the trans-ancestry fine-mapping, the 99% CS was reduced to 3 variants, including rs1974, a 3'UTR variant in the gene *FOXA2*. The improved resolution from the European 99% CS was due to the incorporation of other ancestries with different LD structures, including the East Asian samples. In contrast, the improved resolution from the East Asian 99% CS was driven by the increased sample size (**Supplementary Table N34**).

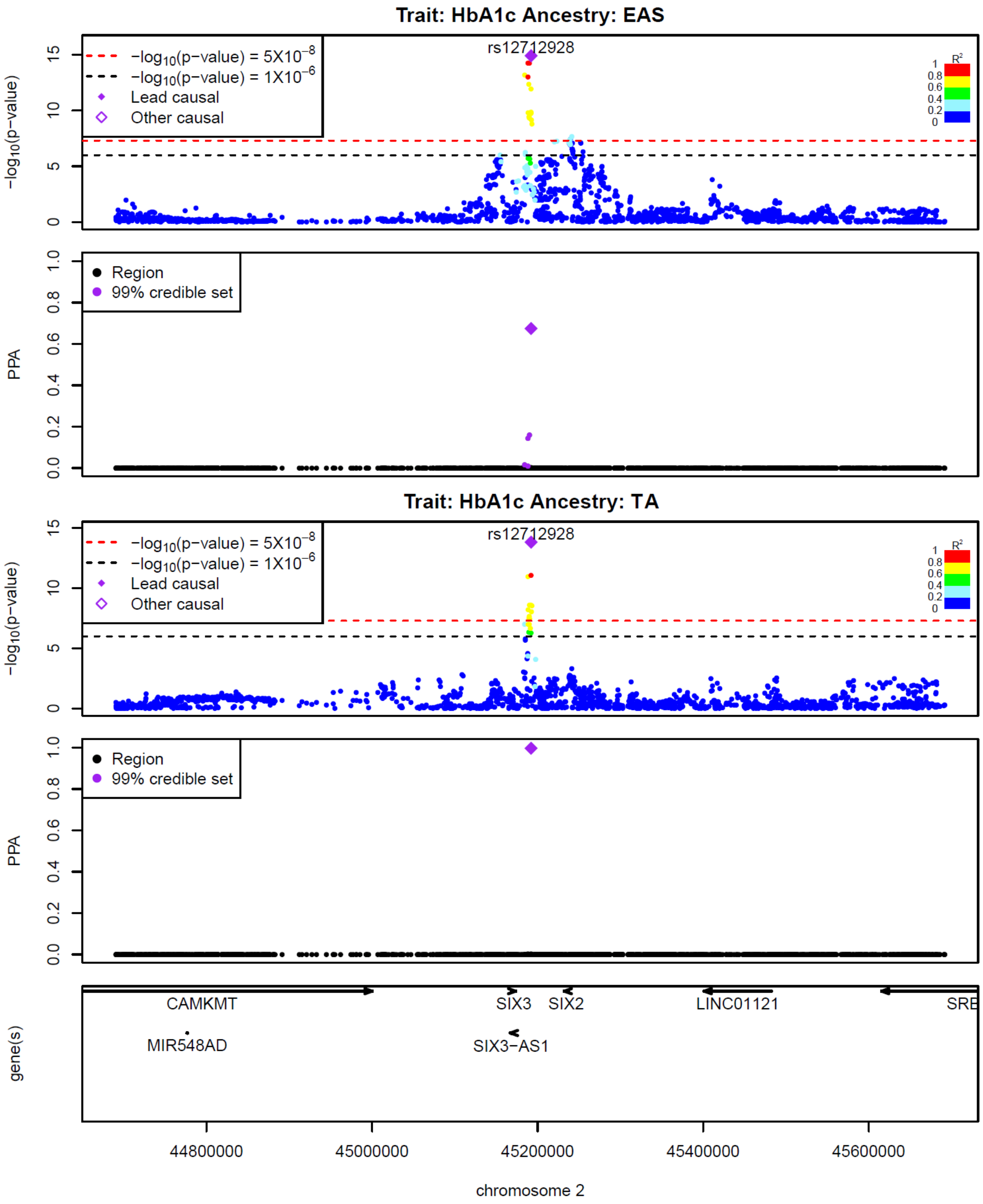

**Supplementary Figure N26 - On this locus zoom plot of locus 22, HbA1c association significance in EAS meta-analysis, PPA in EAS fine-mapping, HbA1c association significance in TA meta-analysis, PPA in TA fine-mapping and genes are present from the top to the bottom.**

**Supplementary Table N34** - Comparison of fine-mapping resolutions at FG-associated locus 228. The resolution of TA-fine-mapping (fifth row) is improved compared with the resolutions of EAS (first row) and EUR (third row) fine-mapping. On the second and fourth rows, the contribution of TA elements is investigated by mimicking the sample size in the TA fine-mapping to match the single-ancestry fine-mapping.

| Ancestry | Sample Size | Predicted Causal Variant | Posterior Probability | # variants in region | # variants in 99% CS |
|---|---|---|---|---|---|
| EAS | 31,669 | rs1337918 | 0.31 | 1,775 | 23 |
| TA | 31,669 | rs1974 | 0.12 | 1,775 | 82 |
| EUR | 165,515 | rs6036152 | 0.22 | 1,775 | 27 |
| TA | 165,515 | rs1974 | 0.81 | 1,775 | 6 |
| TA | 242,353 | rs1974 | 0.94 | 1,775 | 3 |

### 6. Biological signatures of glycaemic trait associated loci

#### a. HbA1c signal classification

Based on results from trans-ancestry and single-ancestry analyses, we had 202 autosomal HbA1c-associated signals. To classify these signals in terms of their likely mode of action, i.e., glycaemic, erythrocytic, or other[1], we made use of association summary statistics for a number of traits from other large European datasets from four broad groups: glycaemic (this effort), mature red blood cell (RBC) traits and reticulocyte traits from Astle et al. [21], and iron traits [22]. Lookups were available for 186 (177 direct, 9 proxies) of the 202 signals, leaving 16 signals with insufficient data for classification (**Methods**).

Before classifying our signals, we first confirmed that our glycaemic and additional traits would cluster together in a biologically meaningful way. For this, we used hierarchical clustering of the traits using the squared Pearson correlation of each trait based on the allele frequencies adjusted effect sizes of LD pruned signals. To avoid double counting across traits, the 186 signals were LD pruned, keeping 129 signals with low pairwise LD ($r^2 < 0.1$) in Europeans. This demonstrated that related sets of traits did cluster together. For example, the glycaemic traits formed a tight cluster, while the reticulocyte, mature red blood cell, and iron traits were grouped into distinct clusters (**Supplementary Figure N27**).

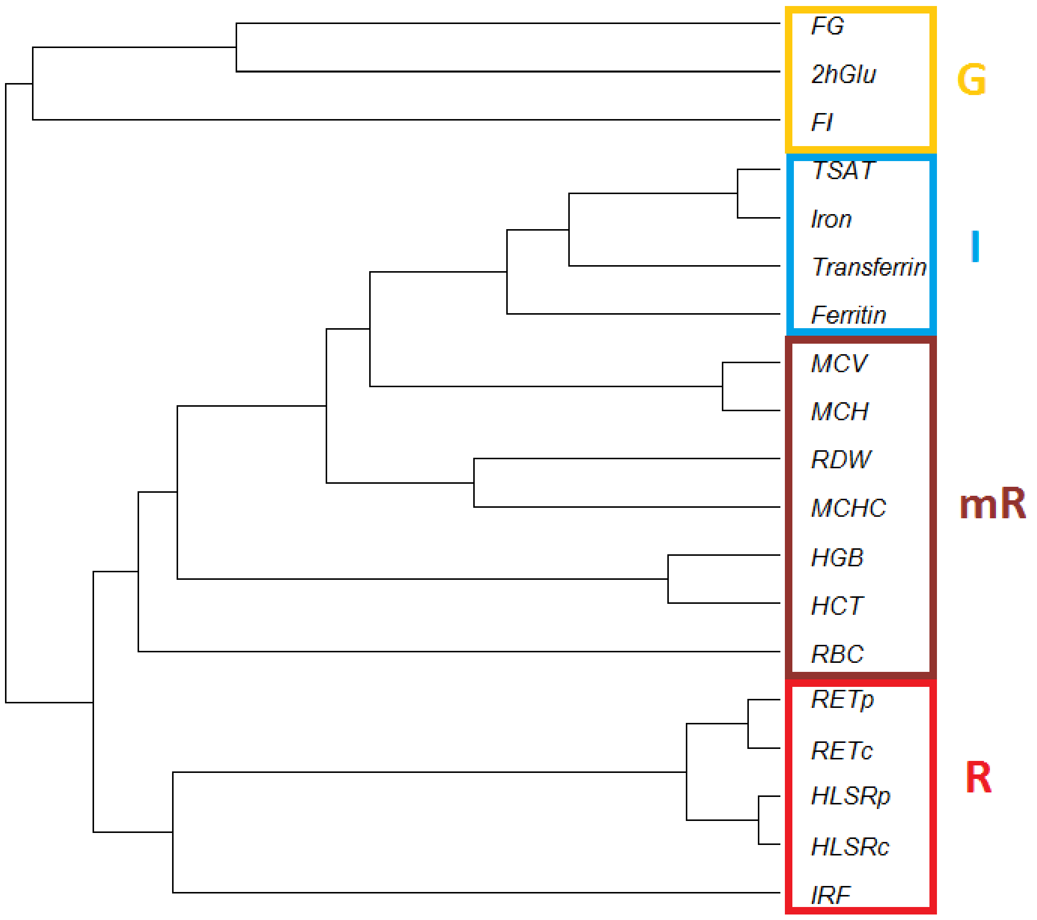

**Supplementary Figure N27 -** Hierarchical clustering of selected traits. Glycaemic traits (G) are in yellow, iron traits (I) are in blue, mature red blood cell traits (mR) are in dark red, and reticulocyte traits (R) are in red.

Next, we obtained uncorrelated trait estimates by conditioning each trait on the other traits, as this is a requirement of Pearson correlation used in the non-negative matrix factorization (NMF) [23] process used to cluster signals. Identification of the best number of clusters was determined by the unsupervised fuzzy evaluation criterion (UFEC) [23] , which suggested that our signals would be best clustered into 8 different clusters (**Supplementary Figure N28**). This clustering approach provides an estimate of the probability of each signal belonging to a given cluster.

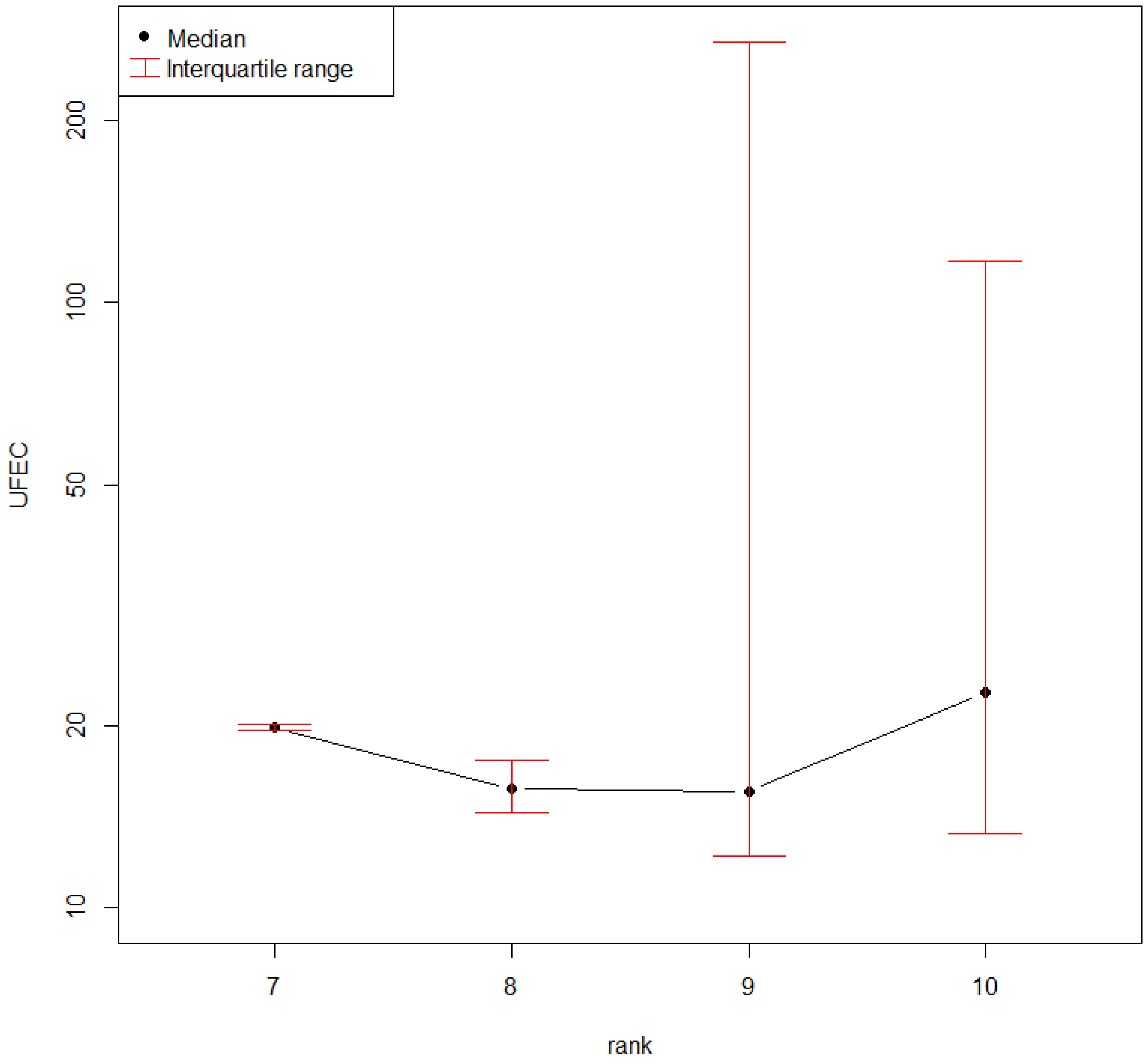

**Supplementary Figure N28 -** Medians and interquartile ranges of UFEC (*y*-axis; logarithmic scale) for different ranks.

To understand what each of these 8 clusters represented biologically, we next calculated a statistic corresponding to the sum of the MAF-adjusted effect sizes, weighted by the probability that a given signal belongs to a stated cluster. A cluster was named as glycaemic, reticulocyte, mature RBC, or iron related if the statistic for a given cluster-trait combination was nominally significant ($P<0.05$) and significantly larger than the mean compared to other traits. Bootstrap was used to evaluate the significance of the test. These results suggested that the eight clusters could in fact be combined into five clusters, each exerting their effects on HbA1c through a specific mechanism, namely clusters 1 and 4 were merged into a glycaemic cluster; 2 and 6 into a mature RBC cluster; 5 and 8 into a reticulocyte cluster; 3 is unknown; and 7 corresponded to an iron cluster (**Supplementary Table N35**).

**Supplementary Table N35** – *P*-value was obtained from Bootstrap, and the most significant cluster is highlighted in bold. G: glycaemic, mR: mature RBC; R: reticulocyte; I: iron; U:unknown

| **Trait** | **1: G** | **2: mR** | **3: U** | **4: G** | **5: R** | **6: mR** | **7: I** | **8: R** |
|---|---|---|---|---|---|---|---|---|
| 2hGlu | **0.00032** | 0.77 | 0.97 | 0.02 | 0.87 | 0.81 | 0.98 | 0.16 |
| FG | 0.18 | 0.97 | 0.96 | **$1\times10^{-18}$** | 0.98 | 0.91 | 0.98 | 0.76 |
| FI | 0.49 | 0.32 | 0.91 | 0.00046 | 0.87 | 0.63 | 0.76 | 0.59 |
| HLSRc | 1 | 0.41 | 0.45 | 0.96 | 0.02 | 0.69 | 0.67 | 0.00075 |
| HLSRp | 1 | 0.33 | 0.56 | 0.97 | 0.05 | 0.58 | 0.56 | 0.00074 |
| IRF | 1 | 0.44 | 0.71 | 0.9 | 0.44 | 0.49 | 0.75 | **$4.3\times10^{-7}$** |
| RETc | 1 | 0.45 | 0.37 | 0.98 | **0.0015** | 0.74 | 0.36 | 0.03 |
| RETp | 1 | 0.36 | 0.49 | 0.98 | 0.006 | 0.62 | 0.21 | 0.04 |
| HCT | 0.99 | 0.79 | 0.67 | 0.87 | 0.34 | 0.02 | 0.02 | 0.49 |
| HGB | 0.99 | 0.86 | 0.73 | 0.95 | 0.29 | 0.1 | 0.00011 | 0.38 |
| MCH | 0.99 | 0.57 | 0.96 | 0.97 | 0.96 | 0.02 | $9\times10^{-7}$ | 0.16 |
| MCHC | 1 | 0.71 | 0.93 | 0.98 | 0.0022 | 0.33 | 0.00049 | 0.39 |
| MCV | 1 | 0.51 | 0.97 | 0.98 | 0.5 | 0.04 | 0.000084 | 0.12 |
| RBC | 0.99 | 0.3 | 0.8 | 0.95 | 0.4 | **0.0018** | 0.65 | 0.03 |
| RDW | 1 | **0.0073** | 0.86 | 0.99 | 0.27 | 0.86 | 0.006 | 0.12 |
| Iron | 0.95 | 0.87 | 0.93 | 0.82 | 0.85 | 0.76 | $1.4\times10^{-13}$ | 0.64 |
| Ferritin | 0.97 | 0.79 | 0.91 | 0.77 | 0.76 | 0.82 | $3.1\times10^{-8}$ | 0.14 |
| Transferrin | 0.92 | 0.76 | 0.89 | 0.78 | 0.84 | 0.87 | $3\times10^{-12}$ | 0.71 |
| TSAT | 0.94 | 0.88 | 0.92 | 0.83 | 0.88 | 0.83 | **$1.9\times10^{-15}$** | 0.69 |

Finally, we classified signals as belonging to a given cluster. We performed hard clustering (a signal was only allowed in a single cluster) and soft clustering (a signal could belong to more than one cluster). Signals were classified into clusters if their probability of belonging to a given cluster was greater than the null expectation (1/number of clusters) or it was the largest probability (hard clustering, **Supplementary Table 20**). We used association results of HbA1c conditioned on FG, HbA1c conditioned on iron traits, and type 2 diabetes association results to verify the naming of each of the clusters. Clusters where the average effect size of signals in that cluster were significantly reduced when adjusted for FG or iron were confirmed as glycaemic and iron, respectively. The glycaemic cluster also had a high average risk of type 2 diabetes, as expected for variants affecting HbA1c through a glycaemic mechanism (**Supplementary Table N36; Supplementary Figure N29**).

**Supplementary Table N36 -** Verification of the labels for each cluster. “Is glycaemic” tests whether the impact on HbA1c (adjusted effect size) reduces more than 0 after adjusting for FG; “Is iron” tests whether the impact on HbA1c reduces more than 0 after adjusting for iron traits in cohorts InterAct and EPIC; “T2D” tests whether the impact on T2D association of signals in each cluster is greater than those not in that cluster.

| Cluster | Is glycaemic | Is iron (InterAct) | Is iron (EPIC) | T2D |
|---|---|---|---|---|
| G | **$1.0{\times}10^{-5}$** | 0.33 | 0.25 | **0.000041** |
| mR | 0.87 | 0.06 | 0.03 | 0.99 |
| R | 0.28 | 0.32 | 0.58 | 1.00 |
| I | 0.50 | **0.0048** | **0.0085** | 1.00 |
| U | 0.26 | 0.42 | 0.32 | 1.00 |

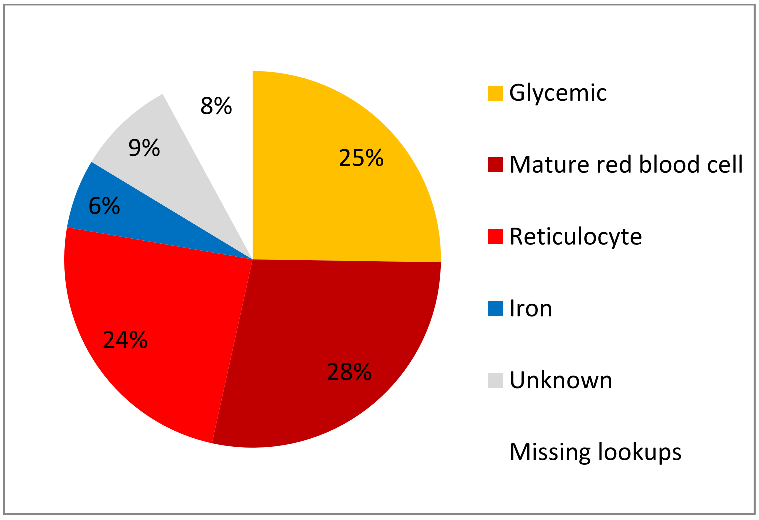

**Supplementary Figure N29 -** Percent of HbA1c signals that fall into each classification based on results from hard clustering.

Next, we compared the results from the current signal classification procedure to that done previously by Wheeler et al[1]. Previously, there were 60 established signals for HbA1c, of which we have data for 22 exact variant matches, 34 with proxies (LD $r^2 \geq 0.8$), and four with no data in this effort (three that did not have appropriate proxies and one on chromosome X we were unable to classify). Overall, there was strong consistency in the classification results between the two analyses with 84.6% of exact variants or proxy variants being in agreement *(P*=$7.1\text{x}10^{-5}$). Even when only poor proxies could be found based on distance ($0.4 < r^2 < 0.8$ or within 500 Kb), most of the results were consistent **(Supplementary Table N37**). Of the signals that shifted into a different classification, three previous RBC signals (rs1800562, rs198846 and rs4820268) have now moved into the iron cluster (which was not included in the previous effort), two previous RBC signals (rs12132919 and rs10774625) are unknown when using our hard clustering but fall into the unknown/reticulocyte/glycaemic and unknown/mature RBC clusters, respectively, in our soft clustering. Three previously-clustered glycaemic signals (rs13134327, rs174577 and rs11603334) are now classified as reticulocytes. Lastly, one variant in *G6PC2* previously classified as glycaemic is now unknown in the hard clustering but falls into the unknown/mature RBC/glycaemic clusters in our soft clustering (**Supplementary Table N37).**

Notably, we are now able to classify 18 of the 19 previously unknown signals, the majority (15/18) of which are now classified as either mature RBC or reticulocyte, one is classified as iron, and two are now classified as glycaemic according to our hard clustering (**Supplementary Table N37).**

**Supplementary Table N37 -** Comparison of HbA1c classifications between this project and prior classifications. Prior classification and updated variants come from Wheeler et al. [1]. 1KG refers to this meta-analysis. Chromosome and position based on hg19. '*' indicates variants classified as "probably RBC" and '#' indicates variants classified as "probably glycaemic" in Wheeler et al. Abbreviations: I, iron; mR, mature red blood cell; R, reticulocyte; RBC, red blood cell; U, unknown.

| Prior Classification | Updated Variant | Chr | Position | Nearest gene | 1KG Variant | LD (EUR $r^2$) or distance (bp) | 1KG soft cluster | 1KG hard cluster |
|---|---|---|---|---|---|---|---|---|
| *Erythrocytic / RBC* | | | | | | | | |
| | rs12132919 | 1 | 156,318,141 | *CCT3 (TMEM79)* | rs12127403 | 44,795 | U/R/G | U |
| | rs857691 | 1 | 158,626,378 | *SPTA1* | rs857725 | 0.78 | R/U | R |
| | rs7616006 | 3 | 12,267,648 | *SYN2* | rs12491937 | 0.99 | R | R |
| | rs1800562 | 6 | 26,093,141 | *HIST1H4A, HFE* | rs1800562 | 1.00 | I | I |
| | rs198846 | 6 | 26,107,463 | *HIST1H4A* | rs1799945 | 0.96 | I | I |
| | rs11964178 | 6 | 109,562,035 | *C6orf183* | rs13195517 | 174,218 | mR/R | mR |
| | rs9494142 | 6 | 135,431,640 | *HBS1L (MYB)* | rs9389268 | 0.78 | mR | mR |
| | rs592423 | 6 | 139,840,693 | *CITED2* | rs9389268 | 4,421,062 | mR | mR |
| | rs4737009 | 8 | 41,630,405 | *ANK1* | rs4737009 | 1.00 | R/U | R |
| | rs6980507 | 8 | 42,383,084 | *SLC20A2* | rs6980507 | 1.00 | mR/R | mR |
| | rs7040409* | 9 | 91,503,236 | *C9orf47* | rs61750929 | 0.90 | R/G | R |
| | rs4745982* | 10 | 71,089,843 | *HK1* | rs4745982 | 1.00 | mR/R | mR |
| | rs11224302* | 11 | 100,456,604 | *CNTN5* | rs11224302 | 1.00 | R/mR/U | R |
| | rs2408955* | 12 | 48,499,131 | *SENP1* | rs76261711 | 12,435 | mR/R | mR |
| | rs10774625 | 12 | 111,910,219 | *ATXN2* | rs10774624 | 0.86 | mR/U | U |
| | rs11248914 | 16 | 293,562 | *ITFG3* | rs11248914 | 1.00 | mR/U | mR |
| | rs4783565* | 16 | 68,750,190 | *CDH3* | rs7198799 | 0.88 | mR/U | mR |
| | rs837763 | 16 | 88,853,729 | *CDT1* | rs837763 | 1.00 | R | R |
| | rs9914988 | 17 | 27,183,104 | *ERAL1* | rs9914988 | 1.00 | R/mR | R |
| | rs17533903* | 19 | 17,256,523 | *MYO9B* | rs17533945 | 0.40 | R/mR | R |
| | rs4820268 | 22 | 37,469,591 | *TMPRSS6* | rs855791 | 0.77 | I | I |
| | rs1050828 | 23 | 153,533,569 | *G6PD* | - | - | - | - |
| *Glycaemic* | | | | | | | | |
| | rs13387347 | 2 | 169,754,846 | *G6PC2* | rs540524 | 0.56 | U/mR/G | U |
| | rs560887 | 2 | 169,763,148 | *G6PC2* | rs560887 | 1.00 | G | G |
| | rs11708067 | 3 | 123,065,778 | *ADCY5* | rs11719201 | 0.97 | G/mR | G |
| | rs8192675 | 3 | 170,724,883 | *SLC2A2* | rs1604038 | 0.97 | G/R | G |
| | rs13134327# | 4 | 144,659,795 | *FREM3* | rs13134327 | 1.00 | R | R |
| | rs7756992 | 6 | 20,679,709 | *CDKAL1* | rs34499031 | 0.98 | G | G |
| | rs2191349 | 7 | 15,064,309 | *DGKB* | rs2191349 | 1.00 | G | G |
| | rs4607517 | 7 | 44,235,668 | *YKT6 (GCK)* | rs2908286 | 0.99 | G | G |
| | rs3824065 | 7 | 44,247,258 | *YKT6 (GCK)* | rs3757840 | 0.73 | G | G |
| | rs11558471 | 8 | 118,185,733 | *SLC30A8* | rs11558471 | 1.00 | G | G |
| | rs2383208 | 9 | 22,132,076 | *MTAP* | rs10811661 | 0.95 | G | G |
| | rs17747324 | 10 | 114,752,503 | *TCF7L2* | rs7903146 | 0.69 | G/mR | G |
| | rs2237896# | 11 | 2,858,440 | *KCNQ1* | rs2237896 | 1.00 | G/R | G |
| | rs174577 | 11 | 61,604,814 | *FADS2* | rs174559 | 0.65 | R/U/mR | R |
| | rs11603334 | 11 | 72,432,985 | *ARAP1* | rs174584 | 10,822,235 | R/U/mR | R |
| | rs10830963 | 11 | 92,708,710 | *MTNR1B* | rs10830963 | 1.00 | G | G |
| | rs11619319 | 13 | 28,487,599 | *PDX1* | rs11619319 | 1.00 | G/mR/R | G |
| | rs576674 | 13 | 33,554,302 | *KL* | rs576674 | 1.00 | G/R | G |
| *Erythrocytic/RBC and Glycaemic* | | | | | | | | |
| | rs579459 | 9 | 136,154,168 | *ABO* | rs649129 | 1.00 | R/G | R |
| *Unknown* | | | | | | | | |
| | rs2375278 | 1 | 25,529,038 | *SYF2* | rs2375278 | 1.00 | U/mR/R | U |
| | rs267737 | 1 | 150,940,625 | *LASS2 (CERS2)* | rs267738 | 1.00 | G/R/mR | G |
| | rs17509001 | 2 | 24,021,231 | *ATAD2B* | rs12612492 | 0.91 | R | R |
| | rs12621844 | 2 | 48,414,735 | *FOXN2* | rs17037289 | 172,463 | mR/R/G | mR |
| | rs17256082 | 2 | 175,292,364 | *SCRN3* | rs17256082 | 1.00 | G/R/U | G |
| | rs9818758 | 3 | 49,382,925 | *USP4* | rs9818758 | 1.00 | R/I | R |
| | rs4874799 | 3 | 171,795,540 | *FNDC3B* | rs7632281 | 0.74 | I/U | I |
| | rs11954649 | 5 | 157,055,491 | *SOX30* | rs1948759 | 612,834 | R/U/mR | R |

| | | | | | | | | |
|---|---|---|---|---|---|---|---|---|
| | rs6474359 | 8 | 41,549,194 | *ANK1* | rs34664882 | 0.88 | R | R |
| | rs1467311 | 9 | 110,536,932 | *KLF4* | rs1467311 | 1.00 | mR/R/I | mR |
| | rs10823343 | 10 | 71,091,013 | *HK1* | rs150705486 | 2,203 | mR/R | mR |
| | rs3782123 | 11 | 205,198 | *BET1L* | rs4980325 | 29,253 | mR/U | mR |
| | rs2110073 | 12 | 7,075,882 | *PHB2* | rs2110073 | 1.00 | R | R |
| | rs282587 | 13 | 113,351,662 | *ATP11A* | rs76533333 | 0.63 | mR/U | mR |
| | rs9604573 | 13 | 114,542,858 | *GAS6* | rs7994900 | 0.92 | mR/I | mR |
| | rs1558902 | 16 | 53,803,574 | *FTO* | rs56137030 | 0.92 | R/I/G/mR | R |
| | rs20733285 | 17 | 76,117,361 | *TMC6* | rs2748427 | 4,503 | mR/R/I | mR |
| | rs1046896 | 17 | 80,685,533 | *FN3KRP* | rs9909940 | 1.00 | mR | mR |
| | rs11086054 | 19 | 17,246,737 | *MYO9B* | rs12982956 | 0.90 | R/U | R |

#### b. HbA1c clusters and T2D genetic risk score (GRS)

Next, we tested whether the GRS built with variants in each of the HbA1c clusters had different effects on T2D risk. To do this, we first performed LD pruning of the signals (**Methods**). This pruning left 126 autosomal signals associated with HbA1c to examine for association with T2D, of which 35 were glycaemic, 38 were mature red blood cell, 35 were reticulocyte, 7 were iron, and 11 were unknown. The GRS comprised of all 126 signals was strongly associated with increased odds for T2D (OR = 2.5, 95% CI 2.5-2.6; $P = 2.4\times10^{-301}$), which was primarily driven by signals in the glycaemic class (glycaemic class variants alone: OR = 2.8, 95% CI 2.7-2.9; $P = 1.1\times10^{-251}$). The GRS of the variants from the non-glycaemic classes was also associated with increased odds for T2D (OR = 1.8, 95% CI 1.8-1.9; $P = 2.3\times10^{-48}$) as well, which were mainly driven by signals in the mature RBC (OR = 1.9, 95% CI 1.8-2.0; $P = 1.4\times10^{-22}$) and reticulocyte (OR = 1.9, 95% CI 1.8-2.0; $P = 4.3\times10^{-28}$) classes. In sensitivity analysis, we found the RBC and reticulocyte GRS associations with T2D were mainly driven by 14 signals belonging to both mature RBC and glycaemic classes (OR = 2.3, 95% CI 2.1-2.5; $P = 8\times10^{-14}$) and 15 signals belonging to both reticulocyte and glycaemic class (OR = 2.2, 95% CI 2.1-2.3; $P = 2.3\times10^{-31}$). However, 24 signals belonging to mature RBC GRS and that were not glycaemic (i.e. $P > 0.05$ with FG, FI, and 2hGlu) were still associated with T2D risk (OR = 1.4, 95% CI 1.3-1.5; $P = 6.9\times10^{-4}$). These results could be partly driven by T2D cases being diagnosed based on HbA1c levels that may be influenced by the non-glycaemic signals, or by glycaemic effects not captured by FI, 2hGlu or FG measures. Unfortunately since T2D diabetes cases were derived from UK biobank (**Methods**) it is not possible to know what test was used to diagnose cases (**Supplemental Figure 15)**.

#### c. Epigenomic landscape of trait-associated variants

We included 'static' annotations, implying annotations that don't vary across cell types such as coding gene regions, intronic regions, or those created by merging epigenomic data such as histone modification peaks across cell types. We utilized 29 total static annotation bed files supplied by [24] (https://data.broadinstitute.org/alkesgroup/LDSCORE/baseline_bedfiles.tgz ). These annotations included: coding, un-translated regions (UTRs), promoter, and intronic regions obtained from UCSC [25]; marks indicating the monomethylation (H3K4me1) and trimethylation (H3K4me3) of histone H3 at lysine 4, acetylation of histone H3 at lysine 9 (H3K9ac) [25-27], and acetylation of histone H3 at lysine 27 (H3K27ac) [28,29]; open chromatin, as reflected by DNase I hypersensitivity sites (DHSs) [27,30]; combined chromHMM and Segway predictions [31], which partition the genome based on distinct and recurring patterns of histone marks into seven underlying chromatin states; regions that are conserved in mammals [32,33]; super-enhancers, which are large clusters of highly active enhancers [29]; and enhancers with balanced bidirectional capped transcripts identified using cap analysis of gene expression (CAGE) in the FANTOM5 panel of samples, which we call FANTOM5 enhancers [34]. Histone marks included in the static annotation set included merged histone mark data from different cell types into a single annotation.

We also included 'stretch' enhancer annotations defined previously in 31 individual cell or tissue types as enhancer chromatin states equal to or longer than 3 Kb [35]. The chromatin states were generated with chromHMM using ChIP-seq data for five histone modifications (H3K4me1, H3K4me3, H3K27ac, H3K36me3, H3K27me3) in each of the 31 cell types.

*GREGOR analysis*: GREGOR computes enrichment for GWAS loci to overlap genomic annotations by taking as input a pruned list of independent and significant GWAS variants. It then considers proxy variants for each lead input variant, since the causal variant(s) are not known. An overlap is reported if the feature overlaps any input lead variant or its LD proxies. For each input variant, GREGOR selects ~500 control variants matched for MAF, distance to the gene, and number of variants in LD with $r^2 \geq 0.8$. Fold enrichment is calculated as the number of unique overlaps over the mean number of loci at which the matched control variants (or their LD proxies) overlap the same feature. This process accounts for the length of the features, as longer features will have more overlap, by chance, with control variant sets.

*fGWAS analysis*: We utilized fGWAS [36] as an orthogonal approach of calculating enrichment of glycaemic trait loci in annotations. fGWAS uses summary level GWAS data in a Bayesian hierarchical model to determine shared properties of loci affecting a trait. The method divides the genome into windows generally larger than the expected LD patterns in the population, containing ~5,000 variants. The method assumes that there is either a single causal variant in a window or none. The model defines the prior probabilities that an association lies in a genomic window and that a variant within the genomic window is causal. These prior probabilities are allowed to depend on overlaps of variants with the user supplied genomic annotations, and are estimated using a Bayes approach based on enrichment patterns of annotations across the genome. We show the log2(max likelihood enrichment parameter estimate) (log2(enrichment)) of each individual annotation for each trait in **Supplementary Figure 17**. We observed consistent patterns of enrichment compared to GREGOR, which uses a pruned list of TA lead variants and EUR index and lead variants, and fGWAS, which uses summary statistics, in that Islet stretch enhancers were most enriched for FG loci (**Supplementary Figures 16-17; Supplementary Tables 15-16**). Coding regions were also enriched while repressed chromatin state regions across cell types were depleted (**Supplementary Figures 16-17; Supplementary Tables 15-16**). FI loci were also significantly enriched in adipose and skeletal muscle stretch enhancers across the two methods.

*GARFIELD analysis*: GARFIELD [37] is another approach to calculate enrichment of GWAS loci in annotations. It uses summary level GWAS data and selects independent variants based on user supplied P-value thresholds by LD pruning. For each independent signal, it then fetches proxy variants in high LD ($r^2 \geq 0.8$) and considers overlaps of user-supplied annotations with the selected set of variants. A generalized linear model (logistic regression) is then fitted that tests for enrichment while accounting for features such as variant distance to known transcription start sites (TSS) and number of proxy variants. Different GWAS significance thresholds can be used to calculate enrichment. We calculated enrichment at two GWAS P-value thresholds of $1x10^{-5}$ and $1x10^{-8}$ (**Supplementary Figure 18**).

While performing multiple testing correction across the different annotations tested for each trait, it is notable that a number of input annotations might be correlated. Therefore, taking the total N for multiple testing results in a stringent significance threshold. To address this, the method can estimate the effective number of independent tests performed or effective number of annotations (Neff). This is done by taking an independent subsample of variants and computing the eigenvalues of the correlation matrix between all considered annotations, and then the effective number of independent tests from the Galwey method [38]. We used the effective number of annotations for each trait to determine the enrichment significance thresholds after Bonferroni correction. We observed more significant enrichments at the lower GWAS threshold of 1x10-5, especially for the 2-hour glucose trait, likely because a more lenient threshold allows for higher power due to more signals. We again observed consistent enrichment patterns across all four traits with the three methods (**Supplementary Figures 16-18; Supplementary Tables 15-16, 18**).

#### 7. References

1 Wheeler, E. *et al.* Impact of common genetic determinants of Hemoglobin A1c on type 2 diabetes risk and diagnosis in ancestrally diverse populations: A transethnic genome-wide meta-analysis. *PLoS medicine* **14**, e1002383, doi:10.1371/journal.pmed.1002383 (2017).

2 Wheeler, E., Marenne, G. & Barroso, I. Genetic aetiology of glycaemic traits: approaches and insights. *Human molecular genetics* **26**, R172-r184, doi:10.1093/hmg/ddx293 (2017).

3 Ng, N. H. J. *et al.* Tissue-Specific Alteration of Metabolic Pathways Influences Glycemic Regulation. *bioRxiv*, 790618, doi:10.1101/790618 (2019).

4 Mahajan, A. *et al.* Fine-mapping type 2 diabetes loci to single-variant resolution using high-density imputation and islet-specific epigenome maps. *Nature genetics* **50**, 1505-1513, doi:10.1038/s41588-018-0241-6 (2018).

5 Xue, A. *et al.* Genome-wide association analyses identify 143 risk variants and putative regulatory mechanisms for type 2 diabetes. *Nature communications* **9**, 2941, doi:10.1038/s41467-018-04951-w (2018).

6 Horikoshi, M. *et al.* Discovery and Fine-Mapping of Glycaemic and Obesity-Related Trait Loci Using High-Density Imputation. *PLoS genetics* **11**, e1005230, doi:10.1371/journal.pgen.1005230 (2015).

7 Chen, G. *et al.* Genome-wide association study identifies novel loci association with fasting insulin and insulin resistance in African Americans. *Human molecular genetics* **21**, 4530-4536, doi:10.1093/hmg/dds282 (2012).

8 Locke, A. E. *et al.* Genetic studies of body mass index yield new insights for obesity biology. *Nature* **518**, 197-206, doi:10.1038/nature14177 (2015).

9 Liu, C. T. *et al.* Trans-ethnic Meta-analysis and Functional Annotation Illuminates the Genetic Architecture of Fasting Glucose and Insulin. *American journal of human genetics* **99**, 56-75, doi:10.1016/j.ajhg.2016.05.006 (2016).

10 Aschard, H., Vilhjalmsson, B. J., Joshi, A. D., Price, A. L. & Kraft, P. Adjusting for heritable covariates can bias effect estimates in genome-wide association studies. *American journal of human genetics* **96**, 329-339, doi:10.1016/j.ajhg.2014.12.021 (2015).

11 Scott, R. A. *et al.* Large-scale association analyses identify new loci influencing glycemic traits and provide insight into the underlying biological pathways. *Nature genetics* **44**, 991-1005, doi:10.1038/ng.2385 (2012).

12 Dupuis, J. *et al.* New genetic loci implicated in fasting glucose homeostasis and their impact on type 2 diabetes risk. *Nature genetics* **42**, 105-116, doi:10.1038/ng.520 (2010).

13 Manning, A. K. *et al.* A genome-wide approach accounting for body mass index identifies genetic variants influencing fasting glycemic traits and insulin resistance. *Nature genetics* **44**, 659-669, doi:10.1038/ng.2274 (2012).
14 Saxena, R. *et al.* Genetic variation in GIPR influences the glucose and insulin responses to an oral glucose challenge. *Nature genetics* **42**, 142-148, doi:10.1038/ng.521 (2010).
15 Bulik-Sullivan, B. K. *et al.* LD Score regression distinguishes confounding from polygenicity in genome-wide association studies. *Nature genetics* **47**, 291-295, doi:10.1038/ng.3211 (2015).
16 Bulik-Sullivan, B. *et al.* An atlas of genetic correlations across human diseases and traits. *Nature genetics* **47**, 1236-1241, doi:10.1038/ng.3406 (2015).
17 Lee, J. J. *et al.* Gene discovery and polygenic prediction from a genome-wide association study of educational attainment in 1.1 million individuals. *Nature genetics* **50**, 1112-1121, doi:10.1038/s41588-018-0147-3 (2018).
18 Nolte, I. M. *et al.* Missing heritability: is the gap closing? An analysis of 32 complex traits in the Lifelines Cohort Study. *European journal of human genetics : EJHG* **25**, 877-885, doi:10.1038/ejhg.2017.50 (2017).
19 Dastani, Z. *et al.* Novel loci for adiponectin levels and their influence on type 2 diabetes and metabolic traits: a multi-ethnic meta-analysis of 45,891 individuals. *PLoS genetics* **8**, e1002607, doi:10.1371/journal.pgen.1002607 (2012).
20 Chang, C. C. *et al.* Second-generation PLINK: rising to the challenge of larger and richer datasets. *GigaScience* **4**, 7, doi:10.1186/s13742-015-0047-8 (2015).
21 Astle, W. J. *et al.* The Allelic Landscape of Human Blood Cell Trait Variation and Links to Common Complex Disease. *Cell* **167**, 1415-1429.e1419, doi:10.1016/j.cell.2016.10.042 (2016).
22 Benyamin, B. *et al.* Novel loci affecting iron homeostasis and their effects in individuals at risk for hemochromatosis. *Nature communications* **5**, 4926, doi:10.1038/ncomms5926 (2014).
23 Binesh, N. & Rezghi, M. Fuzzy clustering in community detection based on nonnegative matrix factoriztion with two novel evaluation criteria. *Applied Soft Computing* **69**, 689-703 (2018).
24 Finucane, H. K. *et al.* Partitioning heritability by functional annotation using genome-wide association summary statistics. *Nature genetics* **47**, 1228-1235, doi:10.1038/ng.3404 (2015).
25 An integrated encyclopedia of DNA elements in the human genome. *Nature* **489**, 57-74, doi:10.1038/nature11247 (2012).
26 Kundaje, A. *et al.* Integrative analysis of 111 reference human epigenomes. *Nature* **518**, 317-330, doi:10.1038/nature14248 (2015).
27 Trynka, G. *et al.* Chromatin marks identify critical cell types for fine mapping complex trait variants. *Nature genetics* **45**, 124-130, doi:10.1038/ng.2504 (2013).
28 Biological insights from 108 schizophrenia-associated genetic loci. *Nature* **511**, 421-427, doi:10.1038/nature13595 (2014).
29 Hnisz, D. *et al.* Super-enhancers in the control of cell identity and disease. *Cell* **155**, 934-947, doi:10.1016/j.cell.2013.09.053 (2013).
30 Gusev, A. *et al.* Partitioning heritability of regulatory and cell-type-specific variants across 11 common diseases. *American journal of human genetics* **95**, 535-552, doi:10.1016/j.ajhg.2014.10.004 (2014).
31 Hoffman, M. M. *et al.* Integrative annotation of chromatin elements from ENCODE data. *Nucleic acids research* **41**, 827-841, doi:10.1093/nar/gks1284 (2013).
32 Lindblad-Toh, K. *et al.* A high-resolution map of human evolutionary constraint using 29 mammals. *Nature* **478**, 476-482, doi:10.1038/nature10530 (2011).
33 Ward, L. D. & Kellis, M. Evidence of abundant purifying selection in humans for recently acquired regulatory functions. *Science (New York, N.Y.)* **337**, 1675-1678, doi:10.1126/science.1225057 (2012).

34 Andersson, R. *et al.* An atlas of active enhancers across human cell types and tissues. *Nature* **507**, 455-461, doi:10.1038/nature12787 (2014).

35 Varshney, A. *et al.* Genetic regulatory signatures underlying islet gene expression and type 2 diabetes. *Proceedings of the National Academy of Sciences of the United States of America* **114**, 2301-2306, doi:10.1073/pnas.1621192114 (2017).

36 Pickrell, J. K. Joint analysis of functional genomic data and genome-wide association studies of 18 human traits. *American journal of human genetics* **94**, 559-573, doi:10.1016/j.ajhg.2014.03.004 (2014).

37 Iotchkova, V. *et al.* GARFIELD classifies disease-relevant genomic features through integration of functional annotations with association signals. *Nature genetics* **51**, 343-353, doi:10.1038/s41588-018-0322-6 (2019).

38 Galwey, N. W. A new measure of the effective number of tests, a practical tool for comparing families of non-independent significance tests. *Genetic epidemiology* **33**, 559-568, doi:10.1002/gepi.20408 (2009).

### The Trans-Ancestral Genomic Architecture of Glycaemic Traits

Ji Chen[1,2]*, Cassandra N. Spracklen[3,4]*, Gaëlle Marenne[1,5]*, Arushi Varshney[6]*, Laura J Corbin[7,8]*, Jian'an Luan[9], Sara Willems[9], Ying Wu[3], Xiaoshuai Zhang[9,10], Momoko Horikoshi[11,12,13], Thibaud S Boutin[14], Reedik Mägi[15], Johannes Waage[16,17], Achilleas Pitsilides[18], Ruifang Li-Gao[19], Kei Hang[20,21,22], Jie Yao[23], Mila D Anasanti[24], Audrey Y Chu[25], Annique Claringbould[26], Jani Heikkinen[24], Jaeyoung Hong[18], Jouke-Jan Hottenga[27,28], Shaofeng Huo[29], Marika A. Kaakinen[30,24], Tin Louie[31], Winfried März[32,33,34], Hortensia Moreno-Macias[35], Anne Ndungu[12], Sarah C. Nelson[31], Ilja M. Nolte[36], Kari North[37], Chelsea K. Raulerson[3], Debashree Ray[38], Rebecca Rohde[37], Denis Rybin[18], Claudia Schurmann[39,40], Xueling Sim[41,42,43], Loz Southam[1], Isobel Stewart[9], Carol A. Wang[44], Yujie Wang[37], Peitao Wu[18], Weihua Zhang[45,46], Tarunveer S. Ahluwalia[16,17], Emil VR Appel[47], Lawrence F. Bielak[48], Jennifer A. Brody[49], Noel P Burtt[50], Claudia P Cabrera[51,52], Brian E Cade[53,54], Jin Fang Chai[55], Xiaoran Chai[56,57], Li-Ching Chang[58], Chien-Hsiun Chen[58], Brian H Chen[59], Kumaraswamy N Chitrala[60], Yen-Feng Chiu[61], Hugoline G. de Haan[19], Graciela E Delgado[34], Ayse Demirkan[62], Qing Duan[3,63], Jorgen Engmann[64], Segun A Fatumo[65,66,67], Javier Gayán[68], Franco Giulianini[69], Jung Ho Gong[20], Stefan Gustafsson[70], Yang Hai[71], Fernando P Hartwig[72,73], Jing He[74], Yoriko Heianza[75], Tao Huang[76], Alicia Huerta-Chagoya[77,78], Mi Yeong Hwang[79], Richard A. Jensen[49], Takahisa Kawaguchi[80], Katherine A Kentistou[81,82], Young Jin Kim[79], Marcus E Kleber[34], Ishminder K Kooner[83], Shuiqing Lai[84], Leslie A Lange[85], Carl D Langefeld[86], Marie Lauzon[23], Man Li[87], Symen Ligthart[62], Jun Liu[62,88], Marie Loh[89,45], Jirong Long[74], Valeriya Lyssenko[90,91], Massimo Mangino[92,93], Carola Marzi[94,95], May E Montasser[96], Abhishek Nag[12], Masahiro Nakatochi[97], Damia Noce[98], Raymond Noordam[99], Giorgio Pistis[100], Michael Preuss[39,101], Laura Raffield[3], Laura J. Rasmussen-Torvik[102], Stephen S Rich[103,104], Neil R Robertson[12], Rico Rueedi[105,106], Kathleen Ryan[96], Serena Sanna[100,26], Richa Saxena[107,108,109], Katharina E Schraut[81,82], Bengt Sennblad[110], Kazuya Setoh[80], Albert V Smith[111,112], Lorraine Southam[113,114], Thomas Sparsø[47], Rona J Strawbridge[115,116], Fumihiko Takeuchi[117], Jingyi Tan[23], Stella Trompet[99,118], Erik van den Akker[119,120,121], Peter J Van der Most[36], Niek Verweij[122,123], Mandy Vogel[124], Heming Wang[53], Chaolong Wang[125,126], Nan Wang[127,128], Helen R Warren[51,52], Wanqing Wen[74], Tom Wilsgaard[129], Andrew Wong[130], Andrew R Wood[131], Tian Xie[36], Mohammad Hadi Zafarmand[132,133], Jing-Hua Zhao[134], Wei Zhao[48], Najaf Amin[62], Zorayr Arzumanyan[23], Arne Astrup[135], Stephan JL Bakker[136], Damiano Baldassarre[137,138], Marian Beekman[119], Richard N Bergman[139], Alain Bertoni[140], Matthias Blüher[141], Lori L. Bonnycastle[142], Stefan R Bornstein[143], Donald W Bowden[144], Qiuyin Cai[74], Archie Campbell[145,146], Harry Campbell[81], Yi Cheng Chang[147,148,149], Eco J.C. de Geus[27,28], Abbas Dehghan[62], Shufa Du[150], Gudny Eiriksdottir[112], Aliki Eleni Farmaki[151,152], Mattias Frånberg[153], Christian Fuchsberger[98], Yutang Gao[154], Anette P Gjesing[47], Anuj Goel[155,12], Sohee Han[79], Catharina A Hartman[156], Christian Herder[157,158,159], Andrew A. Hicks[98], Chang-Hsun Hsieh[160,161], Willa A. Hsueh[162], Sahoko Ichihara[163], Michiya Igase[164], M. Arfan Ikram[62], W. Craig Johnson[31], Marit E Jørgensen[17,165], Peter K Joshi[81], Rita R Kalyani[166], Fouad R. Kandeel[167], Tomohiro Katsuya[168,169], Chiea Chuen Khor[126], Wieland Kiess[124], Ivana Kolcic[170], Teemu Kuulasmaa[171], Johanna Kuusisto[172], Kristi Läll[15], Kelvin Lam[23], Deborah A Lawlor[7,8], Nanette R. Lee[173,174], Rozenn N. Lemaitre[49], Honglan Li[175], Lifelines Cohort Study[176], Shih-Yi Lin[177,178,179], Jaana Lindström[180], Allan Linneberg[181,182], Jianjun Liu[126,183], Carlos Lorenzo[184], Tatsuaki Matsubara[185], Fumihiko Matsuda[80], Geltrude Mingrone[186], Simon Mooijaart[99],

Sanghoon Moon[79], Toru Nabika[187], Girish N. Nadkarni[39], Jerry L. Nadler[188], Mari Nelis[15], Matthew J Neville[11], Jill M Norris[189], Yasumasa Ohyagi[190], Annette Peters[191,95,192], Patricia A. Peyser[48], Ozren Polasek[170], Qibin Qi[193], Dennis Raven[156], Dermot F Reilly[194], Alex Reiner[195], Fernando Rivideneira[196], Kathryn Roll[23], Igor Rudan[197], Charumathi Sabanayagam[56,198], Kevin Sandow[23], Naveed Sattar[199], Annette Schürmann[200,201], Jinxiu Shi[202], Heather M Stringham[43,203], Kent D. Taylor[23], Tanya M. Teslovich[204], Betina Thuesen[181], Paul RHJ Timmers[81], Elena Tremoli[138], Michael Y Tsai[205], Andre Uitterlinden[196], Rob M van Dam[41,183,206], Diana van Heemst[99], Astrid van Hylckama Vlieg[19], Jana V Van Vliet-Ostaptchouk[36], Jagadish Vangipurapu[207], Henrik Vestergaard[47,17], Tao Wang[193], Ko Willems van Dijk[208,209,210], Yongbing Xiang[175], Tatijana Zemunik[211], Goncalo R Abecasis[43], Linda S. Adair[150,212], Carlos Alberto Aguilar-Salinas[213,214,215], Marta E Alarcón-Riquelme[216,217], Ping An[218], Larissa Aviles-Santa[219], Diane M Becker[220], Lawrence J Beilin[221], Sven Bergmann[105,106,222], Hans Bisgaard[16,17], Corri Black[223], Michael Boehnke[224,225], Eric Boerwinkle[226,227], Bernhard O Böhm[228,229], Klaus Bønnelykke[16,17], D I. Boomsma[27,28], Erwin P. Bottinger[39,40,230], Thomas A Buchanan[231,232,128], Mickaël Canouil[233,234], Mark J Caulfield[51,52], John C. Chambers[89,45,83,235,236], Daniel I. Chasman[69,237], Yii-Der Ida Chen[23], Ching-Yu Cheng[56,198], Francis S. Collins[142], Adolfo Correa[238], Francesco Cucca[100], H. Janaka de Silva[239], George Dedoussis[240], Sölve Elmståhl[241], Michele K. Evans[60], Ele Ferranni[242], Luigi Ferruci[243], Jose C Florez[244,245,246], Paul Franks[247], Timothy M Frayling[131], Philippe Froguel[233,234,248], Bruna Gigante[249], Mark O. Goodarzi[250], Penny Gordon-Larsen[150,212], Harald Grallert[94,95], Niels Grarup[47], Sameline Grimsgaard[129], Leif Groop[251,252], Vilmundur Gudnason[112,253], Xiuqing Guo[23], Anders Hamsten[116], Torben Hansen[47], Caroline Hayward[254], Susan R. Heckbert[255], Bernardo L Horta[72], Wei Huang[202], Erik Ingelsson[256], Pankow S James[257], Jost B Jonas[258,259], J. Wouter Jukema[118,260], Pontiano Kaleebu[261], Robert Kaplan[193,195], Sharon L.R. Kardia[48], Norihiro Kato[117], Sirkka M. Keinanen-Kiukaanniemi[262,263], Bong-Jo Kim[79], Mika Kivimaki[264], Heikki A. Koistinen[265,266,267], Jaspal S. Kooner[83,235,236,268], Antje Körner[124], Peter Kovacs[141,269], Diana Kuh[130], Meena Kumari[270], Zoltan Kutalik[271,106], Markku Laakso[172], Timo A. Lakka[272,273,274], Lenore J Launer[275], Karin Leander[276], Huaixing Li[29], Xu Lin[29], Lars Lind[277], Cecilia Lindgren[12,278,279], Simin Liu[84], Ruth J.F. Loos[39,101], Patrik Magnusson[280], Anubha Mahajan[12], Andres Metspalu[15], Dennis O Mook-Kanamori[19,281], Trevor A Mori[221], Patricia B Munroe[51,52], Inger Njølstad[129], Jeffrey R O'Connell[96], Albertine J Oldehinkel[156], Ken K Ong[9], Sandosh Padmanabhan[282], Colin N.A. Palmer[283], Nicholette D Palmer[144], Oluf Pedersen[47], Craig E Pennell[44], David J Porteous[145,284], Peter P. Pramstaller[98], Michael A. Province[218], Bruce M. Psaty[49,255,285,286], Lu Qi[75], Leslie J. Raffel[287], Rainer Rauramaa[274], Susan Redline[53,54], Paul M Ridker[69,237], Frits R. Rosendaal[19], Timo E. Saaristo[288,289], Manjinder Sandhu[290], Jouko Saramies[291], Neil Schneiderman[292], Peter Schwarz[143,293,201], Laura J. Scott[224,225], Elizabeth Selvin[38], Peter Sever[268], Xiao-ou Shu[74], P Eline Slagboom[119], Kerrin S Small[92], Blair H Smith[294], Harold Snieder[36], Tamar Sofer[295,245], Thorkild I.A. Sørensen[296,297,7,8], Tim D Spector[92], Alice Stanton[298], Claire J Steves[92,299], Michael Stumvoll[141], Liang Sun[29], Yasuharu Tabara[80], E Shyong Tai[183,300,301], Nicholas J Timpson[7,8], Anke Tönjes[141], Jaakko Tuomilehto[302,303,304], Teresa Tusie[78,305], Matti Uusitupa[306], Pim van der Harst[122,26], Cornelia van Duijn[307,62], Veronique Vitart[254], Peter Vollenweider[308], Tanja GM Vrijkotte[132], Lynne E Wagenknecht[309], Mark Walker[310], Ya X Wang[259], Nick J Wareham[311], Richard M Watanabe[127,232,128], Hugh Watkins[155,12], Wen B Wei[312], Ananda R Wickremasinghe[313], Gonneke Willemsen[27,28], James F Wilson[81,254], Tien-Yin Wong[56,198], Jer-Yuarn Wu[58], Anny H Xiang[314], Lisa R Yanek[220], Loïc Yengo[233,234,315], Mitsuhiro Yokota[316], Eleftheria Zeggini[317,318,319], Wei Zheng[320], Alan B Zonderman[60], Jerome I Rotter[23], Anna L Gloyn[11,12,321,322], Mark I. McCarthy[11,323,324], Josée Dupuis[18], James B Meigs[325,245,326], Robert Scott[9], Inga Prokopenko[30,24], Aaron

Leong[327,328,329], Ching-Ti Liu[18], Stephen CJ Parker[6,330]*, Karen L. Mohlke[3]*, Claudia Langenberg[9]*, Eleanor Wheeler[9,1]*, Andrew P. Morris[331,332,12]*, Inês Barroso[1,333,2]*

[1]Department of Human Genetics, Wellcome Sanger Institute, Hinxton, Cambridge, UK, [2]Exeter Centre of Excellence in Diabetes (ExCEeD), Exeter Medical School, University of Exeter, Exeter, UK, [3]Department of Genetics, University of North Carolina, Chapel Hill, NC, USA, [4]Department of Biostatistics and Epidemiology, University of Massachusetts, Amherst, MA, USA, [5]Inserm, Univ Brest, EFS, UMR 1078, GGB, Brest, France, [6]Department of Computational Medicine and Bioinformatics, University of Michigan, Ann Arbor, MI, USA, [7]MRC Integrative Epidemiology Unit, University of Bristol, Bristol, Bristol, UK, [8]Department of Population Health Sciences, Bristol Medical School, University of Bristol, Bristol, Bristol, UK, [9]MRC Epidemiology Unit, Institute of Metabolic Science, University of Cambridge, Cambridge, UK, [10]Department of Biostatistics, School of Public Health, Shandong University, Jinan, Shandong, China, [11]Oxford Centre for Diabetes, Endocrinology and Metabolism, Radcliffe Department of Medicine, University of Oxford, Oxford, UK, [12]Wellcome Centre for Human Genetics, University of Oxford, Oxford, UK, [13]Laboratory for Genomics of Diabetes and Metabolism, RIKEN Centre for Integrative Medical Sciences, Yokohama, Japan, [14]Medical Research Council Human Genetics Unit, Institute for Genetics and Molecular Medicine, Edinburgh, UK, [15]Estonian Genome Center, Institute of Genomics, University of Tartu, Tartu, Estonia, [16]COPSAC, Copenhagen Prospective Studies on Asthma in Childhood, Herlev and Gentofte Hospital, University of Copenhagen, Copenhagen, Denmark, [17]Steno Diabetes Center Copenhagen, Gentofte, Denmark, [18]Department of Biostatistics, Boston University School of Public Health, Boston, MA, USA, [19]Department of Clinical Epidemiology, Leiden University Medical Center, Leiden, The Netherlands, [20]Department of Epidemiology, Brown University, Providence, RI, USA, [21]Department of Biomedical Sciences, City University of Hong Kong, Hong Kong SAR, China, [22]Department of Electrical Engineering, City University of Hong Kong, Hong Kong SAR, China, [23]The Institute for Translational Genomics and Population Sciences, `, The Lundquist Institute for Biomedical Innovation at Harbor-UCLA Medical Center, Torrance, CA, USA, [24]Department of Metabolism, Digestion, and Reproduction, Imperial College London, London, UK, [25]Division of Preventive Medicine, Brigham and Women's Hospital, Boston, MA, USA, [26]Department of Genetics, University Medical Center Groningen, University of Groningen, Groningen, The Netherlands, [27]Department of Biological Psychology, Faculty of Behaviour and Movement Sciences, Vrije Universiteit Amsterdam, Amsterdam, The Netherlands, [28]Amsterdam Public Health Research Institute, Amsterdam Universities Medical Center, Amsterdam, The Netherlands, [29]CAS Key Laboratory of Nutrition, Metabolism and Food Safety, Shanghai Institute of Nutrition and Health, University of Chinese Academy of Sciences, Chinese Academy of Sciences, Shanghai, China, [30]Department of Clinical and Experimental Medicine, Section of Statistical Multi-Omics, University of Surrey, Guildford, UK, [31]Department of Biostatistics, University of Washington, Seattle, WA, USA, [32]SYNLAB Academy, SYNLAB Holding Deutschland GmbH, Mannheim, Germany, [33]Clinical Institute of Medical and Chemical Laboratory Diagnostics, Medical University Graz, Graz, Austria, [34]Vth Department of Medicine (Nephrology, Hypertensiology, Rheumatology, Endocrinology, Diabetology), Medical Faculty Mannheim, Heidelberg University, Mannheim, Baden-Württemberg, Germany, [35]Department of Economics, Metropolitan Autonomous University, Mexico City, Mexico, [36]Department of Epidemiology, University Medical Center Groningen, University of Groningen, Groningen, The Netherlands, [37]CVD Genetic Epidemiology Computational Laboratory, Gillings School

of Global Public Health, University of North Carolina, Chapel Hill, NC, USA, [38]Department of Epidemiology, Johns Hopkins University, Baltimore, MD, USA, [39]The Charles Bronfman Institute for Personalized Medicine, Icahn School of Medicine at Mount Sinai, New York, NY, USA, [40]HPI Digital Health Center, Digital Health and Personalized Medicine, Hasso Plattner Institute, Potsdam, Germany, [41]Saw Swee Hock School of Public Health, National University of Singapore, Singapore, Singapore, [42]Center for Statistical Genetics, University of Michigan, Ann Arbor, MI, USA, [43]Department of Biostatistics, University of Michigan, Ann Arbor, MI, USA, [44]School of Medicine and Public Health, Faculty of Medicine and Health, The University of Newcastle, Newcastle, NSW, Australia, [45]Department of Epidemiology and Biostatistics, Imperial College London, London, UK, [46]Department of Cardiology, Ealing Hospital, London North West Healthcare NHS Trust, Middlesex, UK, [47]Novo Nordisk Foundation Center for Basic Metabolic Research, Faculty of Health and Medical Sciences, University of Copenhagen, Copenhagen, Denmark, [48]Department of Epidemiology, School of Public Health, University of Michigan, Ann Arbor, MI, USA, [49]Department of Medicine, Cardiovascular Health Research Unit, University of Washington, Seattle, WA, USA, [50]Metabolism Program, Program in Medical and Population Genetics, Broad Institute, Cambridge, MA, USA, [51]Department of Clinical Pharmacology, William Harvey Research Institute Barts, The London School of Medicine and Dentistry,, Queen Mary University of London, London, UK, [52]NIHR Barts Cardiovascular Biomedical Research Centre, Queen Mary University of London, London, UK, [53]Department of Medicine, Sleep and Circadian Disorders, Brigham and Women's Hospital, Boston, MA, USA, [54]Department of Medicine, Sleep Medicine, Harvard Medical School, Boston, MA, USA, [55]Saw Swee Hock School of Public Health, National Univeristy of Singapore and National University Health System, Singapore, Singapore, [56]Ocular Epidemiology, Singapore Eye Research Institute, Singapore National Eye Centre, Singapore, Singapore, [57]Department of Ophthalmology, National University of Singapore and National University Health System, Singapore, Singapore, [58]Institute of Biomedical Sciences, Academia Sinica, Taipei, Taiwan, Taiwan, [59]Department of Epidemiology, UCLA, Los Angeles, CA, USA, [60]Laboratory of Epidemiology and Population Sciences, National Institute on Aging Intramural Research Program, National Institutes of Health, Baltimore, MD, USA, [61]Institute of Population Health Sciences, National Health Research Institutes, Miaoli, Taiwan, [62]Department of Epidemiology, Erasmus Medical Center, Rotterdam, The Netherlands, [63]Department of Statistics, University of North Carolina at Chapel Hill, Chapel Hill, NC, USA, [64]Institute of Cardiovascular Science, UCL, London, UK, [65]Uganda Medical Informatics Centre (UMIC), MRC/UVRI and London School of Hygiene & Tropical Medicine (Uganda Research Unit), Entebbe, Uganda, [66]London School of Hygiene & Tropical Medicine, London, UK, [67]H3Africa Bioinformatics Network (H3ABioNet) Node, Centre for Genomics Research and Innovation, NABDA/FMST, Abuja, Nigeria, [68]Bioinfosol, Sevilla, Spain, [69]Division of Preventive Medicine, Brigham and Women's Hospital, Boston, MA, USA, [70]Department of Medical Sciences, Molecular Epidemiology and Science for Life Laboratory, Uppsala University, Uppsala, Sweden, [71]Department of Statistics, The University of Auckland, Science Center, Auckland, New Zealand, [72]Postgraduate Program in Epidemiology, Federal University of Pelotas, Pelotas, RS, Brazil, [73]MRC Integrative Epidemiology Unit, University of Bristol, Bristol, UK, [74]Department of Medicine, Epidemiology, Vanderbilt University Medical Center, Nashville, TN, USA, [75]Department of Epidemiology, Tulane University Obesity Research Center,, Tulane University, New Orleans, USA, [76]Department of Epidemiology and Biostatistics, School of Public Health, Peking University, Beijing, China, [77]Molecular Biology and Genomic Medicine Unit, National Council for Science and Technology, Mexico City, Mexico, [78]Molecular Biology and Genomic Medicine Unit, National Institute of Medical Sciences and Nutrition, Mexico City, Mexico, [79]Division of Genome Research, Center for Genome Research, Korea National Institute of Health, Cheongju-si, Chungcheongbuk-do, South Korea, [80]Center for Genomic Medicine, Kyoto Univsersity Graduate School of Medicine, Kyoto, Japan, [81]Centre for Global Health Research, Usher Institute of Population Health Sciences and Informatics, University of Edinburgh, Edinburgh, Scotland, [82]Centre for Cardiovascular Sciences, Queen's Medical Research Institute, University of Edinburgh, Edinburgh, Scotland, [83]Department of Cardiology, Ealing Hospital, London North West Healthcare NHS Trust, Middlesex, UK, [84]Department of Epidemiology, Brown University School of Public Health, Providence, RI, USA, [85]Department of Medicine, Divison of Biomedical Informatics and Personalized Medicine, University of Colorado Anschutz Medical Campus, Denver, CO, USA, [86]Department of Public Health Sciences, Biostatistics and Data Science, Wake Forest School of Medicine, Winston-Salem, NC, USA, [87]Department of Medicine, Division of Nephrology and Hypertension, University of Utah, Salt Lake City, UT, USA, [88]Nuffield Department of Population Health, University of Oxford, Oxford, UK, [89]Lee Kong Chian School of Medicine, Nanyang Technological University, Singapore, Singapore, [90]Department of Clinical Science, Center for Diabetes Research, University of Bergen, Bergen, Norway, [91]Department of Clinical Sciences, Lund University Diabetes Centre, Lund University, Malmo, Sweden, [92]Department of Twin Research and Genetic Epidemiology, School of Life Course Sciences, King's College London, London, UK, [93]NIHR Biomedical Research Centre, Guy’s and St Thomas’ Foundation Trust, London, UK, [94]Institute of Epidemiology, Research Unit of Molecular Epidemiology, Helmholtz Zentrum München Research Center for Environmental Health, Neuherberg, Bavaria, Germany, [95]German Center for Diabetes Research (DZD), Neuherberg, Bavaria, Germany, [96]Department of Medicine, Division of Endocrinology, Diabetes, and Nutrition, University of Maryland School of Medicine, Baltimore, MD, USA, [97]Public Health Informatics Unit, Department of Integrated Sciences, Nagoya University Graduate School of Medicine, Nagoya, Japan, [98]Institute for Biomedicine, Eurac Research, Bolzano, BZ, Italy, [99]Department of Internal Medicine, Section of Gerontology and Geriatrics, Leiden University Medical Center, Leiden, The Netherlands, [100]Istituto di Ricerca Genetica e Biomedica (IRGB), Consiglio Nazionale delle Ricerche (CNR), Monserrato, Italy, Italy, [101]The Mindich Child Health and Development Institute for Personalized Medicine, Icahn School of Medicine at Mount Sinai, New York, NY, USA, [102]Department of Preventive Medicine, Northwestern University Feinberg School of Medicine, Chicago, IL, USA, [103]Center for Public Health Genomics, University of Virginia, Charlottesville, VA, USA, [104]Department of Public Health Sciences, University of Virginia, Charlottesville, VA, USA, [105]Department of Computational Biology, University of Lausanne, Lausanne, Switzerland, [106]Swiss Institute of Bioinformatics, Lausanne, Switzerland, [107]Center for Genomic Medicine, Massachusetts General Hospital, Harvard Medical School, Boston, MA, USA, [108]Department of Anesthesia, Critical Care and Pain Medicine, Massachusetts General Hospital, Boston, MA, USA, [109]Program in Medical and Population Genetics,, Broad Institute, Cambridge, MA, USA, [110]Department of Cell and Molecular Biology., National Bioinformatics Infrastructure Sweden,, Science for Life Laboratory, Uppsala University, Uppsala, Sweden, [111]Department of Biostatistics, University of Mitchican, Ann Arbor, MI, USA, [112]Icelandic Heart Association, Kopavogur, Iceland, [113]Institute of Translational Genomics, Helmholtz Zentrum München – German Research Center for Environmental Health, Institute of Translational Genomics, Helmholtz Zentrum München – German Research Center for Environmental Health, Neuherberg, Germany, [114]Wellcome Sanger Institute, Hinxton, Cambridge, UK, [115]Institute of Health and Wellbeing, University of Glasgow, Glasgow, Glasgow, UK, [116]Department of Medicine Solna, Cardiovascular medicine, Karolinska Institutet,

Stockholm, Sweden, [117]National Center for Global Health and Medicine, Tokyo, Japan, [118]Department of Cardiology, Leiden University Medical Center, Leiden, The Netherlands, [119]Department of Biomedical Data Sciences, Molecular Epidemiology, Leiden University Medical Center, Leiden, The Netherlands, [120]Department of Pattern Recognition & Bioinformatics, Delft University of Technology, Delft, The Netherlands, [121]Department of Biomedical Data Sciences, Leiden Computational Biology Center, Leiden University Medical Center, Leiden, The Netherlands, [122]Department of Cardiology, University Medical Center Groningen, University of Groningen, Groningen, The Netherlands, [123]Genomics plc, Oxford, UK, [124]Center of Pediatric Research, University Children´s Hospital Leipzig, University of Leipzig Medical Center, Leipzig, Germany, [125]Department of Epidemiology and Biostatistics, School of Public Health, Tongji Medical College, Huazhong University of Science and Technology, Wuhan, China, [126]Genome Institute of Singapore, Agency for Science, Technology and Research, Singapore, Singapore, [127]Department of Preventive Medicine, Keck School of Medicine of USC, Los Angeles, CA, USA, [128]USC Diabetes and Obesity Research Institute, Keck School of Medicine of USC, Los Angeles, CA, USA, [129]Department of Community Medicine, Faculty of Health Sciences, UIT the Arctic University of Norway, Tromsø, Norway, [130]MRC Unit for Lifelong Health & Ageing at UCL, London, UK, [131]Department of Human Genetics, University of Exeter RILD Building Royal Devon and Exeter NHS trust Barrack Road, Exeter, UK, [132]Department of Public Health, Amsterdam Public Health Research Institute, Amsterdam Universities Medical Center, Amsterdam, The Netherlands, [133]Department of Clinical Epidemiology, Biostatistics, and Bioinformatics, Amsterdam Public Health Research Institute, Amsterdam Universities Medical Center, Amsterdam, The Netherlands, [134]Department of Public Health and Primary Care, School of Clinical Medicine, University of Cambridge, Cambridge, UK, [135]Department of Nutrition, Exercise, and Sports, Faculty of Science, University of Copenhagen, Copenhagen, Denmark, [136]Department of Internal Medicine, University Medical Center Groningen, University of Groningen, Groningen, The Netherlands, [137]Department of Medical Biotechnology and Translational Medicine, University of Milan, Milan, Milan, Italy, [138]Centro Cardiologico Monzino, IRCCS, Milan, Italy, [139]Department of Physiology and Biophysics, University of Southern California, Los Angeles, CA, USA, [140]Department of Epidemiology and Prevention, Division of Public Health Sciences, Wake Forest School of Medicine, Winston-Salem, NC, USA, [141]Medical Department III – Endocrinology, Nephrology, Rheumatology, University of Leipzig Medical Center, Leipzig, Germany, [142]Medical Genomics and Metabolic Genetics Branch, National Human Genome Research Institute, National Institues of Health, Bethesda, MD, USA, [143]Department for Prevention and Care of Diabetes, Faculty of Medicine Carl Gustav Carus, Technische Universität Dresden, Dresden, Germany, [144]Department of Biochemistry, Wake Forest School of Medicine, Winston-Salem, NC, USA, [145]Centre for Genomic and Experimental Medicine, Institute of Genetics & Molecular Medicine, University of Edinburgh, Western General Hospital, Edinburgh, UK, [146]Usher Institute for Population Health Sciences and Informatics, University of Edinburgh, Edinburgh, UK, [147]Internal Medicine, National Taiwan University Hospital, Taipei, Taiwan, [148]Medical Genomics and Proteomics, National Taiwan University, Taipei, Taiwan, [149]Biomedical Sciences, Institute of Biomedical Sciences, Academia Sinica, Taiepi, Taiwan, [150]Department of Nutrition, Gillings School of Global Public Health, University of North Carolina, Chapel Hill, NC, USA, [151]Department of Population Science and Experimental Medicine, Institute of Cardiovascular Science, University College London, London, UK, [152]Department of Nutrition and Dietetics, School of Health Science and Education, Harokopio University of Athens, Athens, Greece, [153]Department of Medicine Solna, Cardiovascular medicine, Stockholm, Sweden, [154]Department of Epidemiology, Shanghai Cancer Institute, Shanghai,

China, [155]Division of Cardiovascular Medicine, Radcliffe Department of Medicine, University of Oxford, Oxford, UK, [156]Department of Psychiatry, Interdisciplinary Center Psychopathology and Emotion Regulation, University Medical Center Groningen, University of Groningen, Groningen, The Netherlands, [157]Institute for Clinical Diabetology, German Diabetes Center, Leibniz Center for Diabetes Research at Heinrich Heine University Düsseldorf, Düsseldorf, Germany, [158]Division of Endocrinology and Diabetology, Medical Faculty, Heinrich Heine University Düsseldorf, Düsseldorf, Germany, [159]German Center for Diabetes Research (DZD), Düsseldorf, Germany, [160]Internal Medicine, Endocrine & Metabolism, Tri-Service General Hospital, Taipei, Taiwan, [161]School of Medicine, National Defense Medical Center, Taipei, Taiwan, [162]Internal Medicine, Endocrinology, Diabetes & Metabolism, Diabetes and Metabolism Research Center, The Ohio State University Wexner Medical Center, Columbus, OH, USA, [163]Department of Environmental and Preventive Medicine, Jichi Medical University School of Medicine, Shimotsuke, Japan, [164]Department of Anti-aging Medicine, Ehime University Graduate School of Medicine, Toon, Japan, [165]National Institute of Public Health, University of Southern Denmark, Odense, Denmark, [166]Department of Medicine, Endocrinology, Diabetes & Metabolism, Johns Hopkins University School of Medicine, Baltimore, MD, USA, [167]Clinical Diabetes, Endocrinology & Metabolism, Translational Research & Cellular Therapeutics, Beckman Research Institute of the City of Hope, Duarte, CA, USA, [168]Department of Clinical Gene Therapy, Osaka University Graduate School of Medicine, Suita, Japan, [169]Department of Geriatric and General Medicine, Osaka University Graduate School of Medicine, Suita, Japan, [170]Department of Public Health, University of Split School of Medicine, Split, Croatia, [171]Institute of Biomedicine, Bioinformatics Center, Univeristy of Eastern Finland, Kuopio, Finland, [172]Department of Medicine, University of Eastern Finland and Kuopio University Hospital, Kuopio, Finland, [173]USC-Office of Population Studies Foundation, University of San Carlos, Cebu City, Philippines, [174]Department of Anthropology, Sociology and History, University of San Carlos, Cebu City, Philippines, [175]State Key Laboratory of Oncogene and Related Genes & Department of Epidemiology, Shanghai Cancer Institute, Renji Hospital, Shanghai Jiaotong University School of Medicine, Shanghai, China, [176]Lifelines cohort, Groningen, The Netherlands, [177]Internal Medicine, Endocrine & Metabolism, Taichung Veterans General Hospital, Taichung, Taiwan, [178]Center for Geriatrics and Gerontology,, Taichung Veterans General Hospital, Taichung, Taiwan, [179]National Defense Medical Center, National Yang-Ming University, Taipei, Taiwan, [180]Diabetes Prevention Unit, National Institute for Health and Welfare, Helsinki, Finland, [181]Center for Clinical Research and Prevention, Bispebjerg and Frederiksberg Hospital, Copenhagen, Denmark, [182]Department of Clinical Medicine, Faculty of Health and Medical Sciences, University of Copenhagen, Copenhagen, Denmark, [183]Yong Loo Lin School of Medicine, National University of Singapore and National University Health System, Singapore, Singapore, [184]Department of Medicine, University of Texas Health Sciences Center, San Antonio, TX, USA, [185]Department of Internal Medicine, Aichi Gakuin University School of Dentistry, Nagoya, Japan, [186]Department of Diabetes, Diabetes, & Nutritional Sciences, James Black Centre, King's College London, London, UK, [187]Department of Functional Pathology, Shimane University School of Medicine, Izumo, Japan, [188]Department of Medicine and Pharmacology, New York Medical College School of Medicine, Valhalla, NY, USA, [189]Colorado School of Public Health, University of Colorado Anschutz Medical Campus, Aurora, CO, USA, [190]Department of Geriatric Medicine and Neurology, Ehime University Graduate School of Medicine, Toon, Japan, [191]Institute of Epidemiology, Helmholtz Zentrum München Research Center for Environmental Health, Neuherberg, Bavaria, Germany, [192]Institute for Medical Information Processing, Biometry, and Epidemiology, Ludwig-Maximilians

University Munich, Munich, Bavaria, Germany, [193]Department of Epidemiology and Population Health, Albert Einstein College of Medicine, Bronx, NY, USA, [194]Genetics and Pharmacogenomics, Merck Sharp & Dohme Corp., Kenilworth, NJ, USA, [195]Department of Public Health Sciences, Fred Hutchinson Cancer Research Center, Seattle, WA, USA, [196]Department of Internal Medicine, Erasmus Medical Center, Rotterdam, The Netherlands, [197]Usher Institute of Population Health Sciences and Informatics, University of Edinburgh, Edinburgh, UK, [198]Ophthalmology & Visual Sciences Academic Clinical Program (Eye ACP), Duke-NUS Medical School, Singapore, Singapore, [199]BHF Glasgow Cardiovascular Research Centre, Institute of Cardiovascular and Medical Sciences, University of Glasgow, Glasgow, UK, [200]Department of Experimental Diabetology, German Institute of Human Nutrition Potsdam-Rehbruecke, Nuthetal, Germany, [201]German Center for Diabetes Research (DZD), Neuherberg, Germany, [202]Department of Genetics, Shanghai-MOST Key Laboratory of Health and Disease Genomics, Chinese National Human Genome Center at Shanghai (CHGC) and Shanghai Academy of Science & Technology (SAST), Shanghai, China, [203]Center for Statistical Genetics, Ann Arbor, MI, USA, [204]Sarepta Therapeutics, Cambridge, Massachusetts, USA, [205]Department of Laboratory Medicine and Pathology, University of Minnesota, Minneapolis, MN, USA, [206]Department of Nutrition, Harvard T.H. Chan School of Public Health, Boston, MA, USA, [207]Institute of Clinical Medicine, Internal Medicine, University of Eastern Finland, Kuopio, Finland, [208]Department of Internal Medicine, Division of Endocrinology, Leiden University Medical Center, Leiden, The Netherlands, [209]Laboratory for Experimental Vascular Medicine, Leiden University Medical Center, Leiden, The Netherlands, [210]Department of Human Genetics, Leiden University Medical Center, Leiden, The Netherlands, [211]Department of Human Biology, University of Split School of Medicine, Split, Croatia, [212]Carolina Population Center, University of North Carolina, Chapel Hill, NC, USA, [213]Department of Endocrinology and Metabolism, Instituto Nacional de Ciencias Medicas y Nutricion, Mexico City, Mexico, [214]225Unidad de Investigacion de Enfermedades Metabolicas. Tec Salud, Mexico City, Mexico, [215]Instituto Tecnológico y de Estudios Superiores de Monterrey Tec Salud, Mexico City, Mexico, [216]Department of Medical Genomics, Pfizer/University of Granada/Andalusian Government Center for Genomics and Oncological Research (GENYO), Granada, Spain, [217]Institute for Environmental Medicine, Chronic Inflammatory Diseases, Karolinska Institutet, Solna, Sweden, [218]Department of Genetics, Division of Statistical Genomics, Washington University School of Medicine, St. Louis, MO, USA, [219]Clinical and Health Services Research, National Institute on Minority Health and Health Disparities, Bethesda, MD, USA, [220]Department of Medicine, General Internal Medicine, Johns Hopkins University School of Medicine, Baltimore, MD, USA, [221]School of Medicine and Pharmacology, Royal Perth Hospital Unit, University of Western Australia, Perth, WA, Australia, [222]Department of Integrative Biomedical Sciences, University of Cape Town, Cape Town, South Africa, [223]Aberdeen Centre for Health Data Science, 1:042 Polwarth Building,, School of Medicine, Medical, Science and Nutrition, University of Aberdeen, Foresterhill, Aberdeen, UK, [224]Department of Biostatistics, School of Public Health, University of Michigan, Ann Arbor, MI, USA, [225]Center for Statistical Genetics, University of Michigan, Ann Arbor, MI, USA, [226]Human Genetics Center, School of Public Health, The University of Texas Health Science Center at Houston, Houston, TX, USA, [227]Human Genome Sequencing Center, Baylor College of Medicine, Houston, TX, USA, [228]Division of Endocrinology and Diabetes, Graduate School of Molecular Endocrinology and Diabetes, University of Ulm, Ulm, Baden-Württemberg, Germany, [229]LKC School of Medicine, Nanyang Technological University, Singapore and Imperial College London, UK, Singapore, Singapore, [230]University of Potsdam, Berlin-Potsdam, Germany, [231]Department of Medicine, Keck School of Medicine of USC,

Los Angeles, CA, USA, [232]Department of Physiology and Neuroscience, Keck School of Medicine of USC, Los Angeles, CA, USA, [233]INSERM UMR 1283 / CNRS UMR 8199, European Institute for Diabetes (EGID), Université de Lille, Lille, France, [234]INSERM UMR 1283 / CNRS UMR 8199, European Institute for Diabetes (EGID), Institut Pasteur de Lille, Lille, France, [235]Imperial College Healthcare NHS Trust, Imperial College London, London, UK, [236]MRC-PHE Centre for Environment and Health, Imperial College London, London, UK, [237]Havard Medical School, Boston, MA, USA, [238]Department of Medicine, Jackson Heart Study, University of Mississippi Medical Center, Jackson, MS, USA, [239]Department of Medicine, Faculty of Medicine, University of Kelaniya, Ragama, Sri Lanka, [240]Department of Nutrition and Dietetics, School of Health Science and Education, Harokopio University of Athens, Kallithea, Greece, [241]Department of Clinical Sciences, Lund University, Malmö, Sweden, [242]CNR Institute of Clinical Physiology, Pisa, Italy, [243]Intramural Research Program, National Institute of Aging, Baltimore, MD, USA, [244]Diabetes Unit and Center for Genomic Medicine, Massachusetts General Hospital, Boston, MA, USA, [245]Department of Medicine, Harvard Medical School, Boston, MA, USA, [246]Programs in Metabolism and Medical & Population Genetics, Broad Institute, Cambridge, MA, USA, [247]Department of Clinical Sciences, Lund University Diabetes Centre, Malmö, Sweeden, [248]Department of Genomics of Common Disease, Imperial College London, London, UK, [249]Department of Medicine, Cardiovascular medicine, Karolinska Institutet, Stockholm, Sweden, [250]Department of Medicine, Division of Endocrinology, Diabetes & Metabolism, Cedars-Sinai Medical Center, Los Angeles, CA, USA, [251]Diabetes Centre, Lund University, Sweden, [252]Finnish Instituter of Molecular Medicine, Helsinki University, Helsinki, Finland, [253]Faculty of Medicine, School of health sciences, University of Iceland, Reykjavik, Iceland, [254]MRC Human Genetics Unit, Institute of Genetics and Molecular Medicine, University of Edinburgh, Edinburgh, Scotland, [255]Department of Epidemiology, Cardiovascular Health Research Unit, University of Washington, Seattle, WA, USA, [256]Department of Medicine, Division of Cardiovascular Medicine, Stanford University School of Medicine, Stanford University, Stanford, CA, USA, [257]Division of Epidemiology and Community Health, University of Minnesota, Minneapolis, MN, USA, [258]Department of Ophthalmology, Medical Faculty Mannheim, Heidelberg University, Mannheim, Germany, [259]Beijing Institute of Ophthalmology, Beijing Ophthalmology and Visual Science Key Lab, Beijing Tongren Eye Center, Beijing Tongren Hospital, Capital Medical University, Beijing, China, [260]Netherlands Heart Institute, Utrecht, The Netherlands, [261]MRC/UVRI and LSHTM (Uganda Research Unit), Entebbe, Uganda, [262]Faculty of Medicine, Institute of Health Sciences, University of Oulu, Oulu, Finland, [263]Unit of General Practice, Oulu University Hospital, Oulu, Finland, [264]Department of Epidemiology and Public Health, UCL, London, UK, [265]Department of Public Health Solutions, Finnish Institute for Health and Welfare, Helsinki, Finland, [266]Department of Medicine, University of Helsinki and Helsinki University Central Hospital, Helsinki, Finland, [267]Minerva Foundation Institute for Medical Research, Helsinki, Finland, [268]National Heart and Lung Institute, Imperial College London, London, UK, [269]IFB Adiposity Diseases, University of Leipzig Medical Center, Leipzig, Germany, [270]Institute for Social and Economic Research, University of Essex, Colchester, UK, [271]University Institute of Primary Care and Public Health, Division of Biostatistics, University of Lausanne, Lausanne, Switzerland, [272]Institute of Biomedicine, School of Medicine, University of Eastern Finland, Finland, [273]Department of Clinical Physiology and Nuclear Medicine, Kuopio University Hospital, Kuopio, Finland, [274]Foundation for Research in Health Exercise and Nutrition, Kuopio Research Institute of Exercise Medicine, Kuopio, Finland, [275]Laboratory of Epidemiology and Population Sciences, National Institute on Aging, National Institutes of Health, Baltimore, MD, USA, [276]Institute of Environmental Medicine, Cardiovascular and

Nutritional Epidemiology, Karolinska Institutet, Stockholm, Sweden, [277]Department of Medical Sciences, Uppsala, Sweden, [278]Big Data Institute, Nuffield Department of Medicine, University of Oxford, Oxford, UK, [279]Nuffield Department of Women's and Reproductive Health, University of Oxford, Oxford, UK, [280]Department of Medical Epidemiology and Biostatistics and the Swedish Twin Registry, Karolinska Institute, Stockholm, Sweeden, [281]Department of Public Health and Primary Care, Leiden University Medical Center, Leiden, The Netherlands, [282]Institute of Cardiovascular and Medical Sciences, University of Glasgow, Glasgow, UK, [283]Division of Population Health and Genomics, School of Medicine, University of Dundee, Ninewells Hospital and Medical School, Dundee, UK, [284]Centre for Cognitive Ageing and Cognitive Epidemiology, University of Edinburgh, Edinburgh, UK, [285]Department of Health Services, Cardiovascular Health Research Unit, University of Washington, Seattle, WA, USA, [286]Kaiser Permanente Washington Health Research Institute, Seattle, WA, USA, [287]Department of Pediatrics, Genetic and Genomic medicine, University of California, Irvine, Irvine, CA, USA, [288]Tampere, Finnish Diabetes Association, Tampere, Finland, [289]Pirkanmaa Hospital District, Tampere, Finland, [290]Department of Medicine, University of Cambridge, Cambridge, UK, [291]South Karelia Central Hospital, Lappeenranta, Finland, [292]Department of Psychology, University of Miami, Miami, FL, USA, [293]Paul Langerhans Institute Dresden of the Helmholtz Center Munich, University Hospital and Faculty of Medicine, Dresden, Germany, [294]Division of Population Health and Genomics, Ninewells Hospital and Medical School, University of Dundee, Dundee, UK, [295]Division of Sleep and Circadian Disorders, Brigham and Women's Hospital, Boston, MA, USA, [296]Novo Nordisk Foundation Center for Basic Metabolic Research, Section of Metabolic Genetics,, Faculty of Health and Medical Sciences, University of Copenhagen, Copenhagen, Denmark, [297]Department of Public Health, Section of Epidemiology, Faculty of Health and Medical Sciences, University of Copenhagen, Copenhagen, Denmark, [298]Departent of Molecular and Cellular Therapeutics, Royal College of Surgeons in Ireland, Dublin, Ireland, [299]Department of Aging and Health, Guy's and St Thomas' Foundation Trust, London, UK, [300]Saw Swee Hock School of Public Health, National University of Singapore and National University Health System, Singapore, Singapore, [301]Duke-NUS Medical School, Singapore, Singapore, [302]Department of Public Health Solutions, National Institute for Health and Welfare, Helsinki, Finland, [303]Department of Public Health, University of Helsinki, Helsinki, Finland, [304]Saudi Diabetes Research Group, King Abdulaziz University, Jeddah, Saudi Arabia, [305]Department of Genomic Medicine and Environmental Toxicology, Instituto de Investigaciones Biomedicas, Universidad Nacional Autonoma de Mexico, Mexico City, Mexico, [306]Department of Public Health and Clinical Nutrition, University of Eastern Finland, Finland, [307]Department of Epidemiology, University of Oxford, Oxford, UK, [308]Department of Medicine, Internal Medicine, Lausanne University Hospital (CHUV), Lausanne, Switzerland, [309]Department of Public Health Sciences, Wake Forest School of Medicine, Winston-Salem, NC, USA, [310]Faculty of Medical Sciences, Newcastle University, Newcastle upon Tyne, UK, [311]MRC Epidemiology Unit, University of Cambridge, Cambridge, United Kingdom, [312]Beijing Tongren Eye Center, Beijing Key Laboratory of Intraocular Tumor Diagnosis and Treatment, Beijing Ophthalmology & Visual Sciences Key Lab, Beijing Tongren Hospital, Capital Medical University, Beijing, China, China, [313]Department of Public Health, Faculty of Medicine, University of Kelaniya, Ragama, Sri Lanka, [314]Department of Research and Evaluation, Kaiser Permanente of Southern California, Pasadena, CA, USA, [315]Institute for Molecular Bioscience, The University of Queensland, Queensland, Australia, [316]Kurume University School of Medicine, Japan, [317]Institute of Translational Genomics, Helmholtz Zentrum München – German Research Center for Environmental Health,

Neuherberg, Germany, [318]Wellcome Sanger Institute, Hinxton, UK, [319]TUM School of Medicin, Technical University of Munich and Klinikum Rechts der Isar, Munich, Germany, [320]Department of Medicine, Division of Epidemiology, Vanderbilt Epidemiology Center, Vanderbilt University Medical Center, Nashville, TN, USA, [321]NIHR Oxford Biomedical Research Centre, Churchill Hospital, Oxford, UK, [322]Department of Pediatrics, Division of Endocrinology, Stanford School of Medicine, Stanford, CA, USA, [323]Wellcome Centre for Human Genetics, Nuffield Department of Medicine, University of Oxford, Oxford, UK, [324]Oxford NIHR Biomedical Research Centre, Oxford University Hospitals NHS Foundation Trust, Oxford, UK, [325]Department of Medicine, Division of General Internal Medicine, Massachusetts General Hospital, Boston, MA, USA, [326]Program in Medical and Population Genetics, Broad Institute, Cambridge, MA, USA, [327]Department of Medicine, General Internal Medicine, Massachusetts General Hospital, Boston, MA, USA, [328]Department of Medicine, Diabetes Unit and Endocrine Unit, Massachusetts General Hospital, Boston, MA, USA, [329]Harvard Medical School, Boston, MA, USA, [330]Department of Human Genetics, University of Michigan, Ann Arbor, MI, USA, [331]Department of Musculoskeletal and Dermatological Sciences, Faculty of Medicine, Biology and Health, University of Manchester, Manchester, UK, [332]Department of Biostatistics, University of Liverpool, Liverpool, UK, [333]MRC Epidemiology Unit, University of Cambridge, Cambridge, UK

* Denote shared authorship contributions

Lifelines Group Author and Affiliations

Behrooz Z Alizadeh (1), H Marike Boezen (1), Lude Franke (2), Pim van der Harst (3), Gerjan Navis (4), Marianne Rots (5), Harold Snieder (1), Morris Swertz (2), Bruce HR Wolffenbuttel (6), Cisca Wijmenga (2)

1. *Department of Epidemiology, University of Groningen, University Medical Center Groningen, The Netherlands*
2. *Department of Genetics, University of Groningen, University Medical Center Groningen, The Netherlands*
3. *Department of Cardiology, University of Groningen, University Medical Center Groningen, The Netherlands*
4. *Department of Internal Medicine, Division of Nephrology, University of Groningen, University Medical Center Groningen, The Netherlands*
5. *Department of Pathology and Medical Biology, University of Groningen, University Medical Center Groningen, The Netherlands*
6. *Department of Endocrinology, University of Groningen, University Medical Center Groningen, The Netherlands*

**Individual Author Conflicts of Interest and Other Disclosures**

| Author | Disclosure |
|---|---|
| Arne Astrup | Dr. Astrup is recipient of honoraria as speaker for a wide range of Danish and international concerns and receives royalties from textbooks, and from popular diet and cookery books. Dr. Astrup is also co-inventor of a number of patents, including Methods of inducing weight loss, treating obesity and preventing weight gain (licensee Gelesis, USA) and Biomarkers for predicting degree of weight loss (licensee Nestec SA, CH), owned by the University of Copenhagen, in accordance with Danish law. |
| Inês Barroso | IB and spouse own stock in GlaxoSmithKline and Incyte Corporation. |
| Mark Caulfield | MC is an NIHR Senior Investigator. MC is also seconded to Genomics England as Chief Scientist, but there is no conflict to declare. |
| Brian H. Chen | Current Employee of Life Epigenetics, Inc. All work was completed prior to employment at Life Epigenetics. |
| Audrey Y. Chu | Current Employee of Merck & Co. All work was completed prior to employment by Merck & Co. |
| Jose C Florez | Consulting honorarium from Janssen |
| Javier Gayan | He is currently an employee of F. Hoffmann-La Roche Ltd, and owns stock of Roche and GSK. |
| Anna L Gloyn | Consulting honorarium from Merck, speaker honorarium from Novo Nordisk |
| Christian Herder | CH has received honoraria from Sanofi and Lilly and research support from Sanofi unrelated to this manuscript**.** |
| Winfried März | Grants from Siemens Healthineers, grants and personal fees from Aegerion Pharmaceuticals, grants and personal fees from AMGEN, grants from Astrazeneca, grants and personal fees from Sanofi, grants and personal fees from Alexion Pharmaceuticals, grants and personal fees from BASF, grants and personal fees from Abbott Diagnostics, grants and personal fees from Numares AG, grants and personal fees from Berlin-Chemie, grants and personal fees from Akzea Therapeutics, grants from Bayer Vital GmbH , grants from bestbion dx GmbH, grants from Boehringer Ingelheim Pharma GmbH Co KG, grants from Immundiagnostik GmbH, grants from Merck Chemicals GmbH, grants from MSD Sharp and Dohme GmbH, grants from Novartis Pharma GmbH, grants from Olink Proteomics, other from Synlab Holding Deutschland GmbH, all outside the submitted work. |
| Mark I McCarthy | The views expressed in this article are those of the author(s) and not necessarily those of the NHS, the NIHR, or the Department of Health. He has served on advisory panels for Pfizer, NovoNordisk, Zoe Global and received honoraria from Merck, Pfizer, NovoNordisk and Eli Lilly. He holds stock options in Zoe Global and has received research funding from Abbvie, Astra Zeneca, Boehringer Ingelheim, Eli Lilly, Janssen, Merck, NovoNordisk, Pfizer, Roche, Sanofi Aventis, Servier, Takeda. As of June 2019, he is an employee of Roche-Genentech, and a holder of Roche stock. |
| James B Meigs | Consultant for Quest Diagnostics, Inc. They make an HbA1c assay. |
| May E Montasser | MEM reports grant from Regeneron Pharmaceuticals.MEM is an inventor on a patent that was published by the United States Patent and Trademark Office on December 6, 2018 under Publication Number US 2018-0346888, and international patent application that was published on December 13, 2018 |

| | |
|---|---|
| | under Publication Number WO-2018/226560. All work was completed before these COI arise, and all work in unrelated to these COI |
| Dennis O. Mook-Kanamori | Part-time clinical research consultant for Metabolon, Inc. |
| Colin N.A. Palmer | CP has received research support from GSK and AZ not related to this manuscript. |
| Bruce M Psaty | Psaty serves on the Steering Committee of the Yale Open Data Access Project funded by Johnson & Johnson. |
| Naveed Sattar | N.S. has consulted for Astrazeneca, Boehringer Ingelheim, Eli Lilly, Novo Nordisk, Napp and Sanofi and received grant support from Boehringer Ingelheim |
| Tim Spector | Zoe Global Ltd - Founder |

The views expressed in this manuscript are those of the authors and do not necessarily reflect on the official views of the National Institute on Minority Health and Health Disparities, the National Heart, Lung, and Blood Institute, the National Cancer Institute, the National Institutes of Health, or the U.S. federal government.

**Individual Funding and/or Other Acknowledgements**

| Individual | Acknowledgement(s) |
|---|---|
| Tarunveer S. Ahluwalia | Tarunveer S. Ahluwalia was supported by internal funding from Steno Diabetes Center Copenhagen, Gentofte, Denmark and from the Novo Nordisk Foundation (Steno Collaborative 2018) Grant NNF18OC0052457 and acknowledges the same |
| Mila D. Anasanti | MDA is supported by Indonesia Endowment Fund for Education (LPDP 20150822013694) |
| Inês Barroso | IB acknowledges funding from Wellcome (WT206194). This work was partly supported by "Expanding excellence in England" award from Research England. |
| Eric Boerwinkle | EB is supported by HHSN268201700001I |
| Dorret I Boomsma | DIB acknowledges the Royals Netherlands Academy of Science (KNAW) Academy Professor Award (PAH/6635) t |
| Brian E. Cade | BEC is supported by funds from the NIH (K01-HL135405) |
| John C Chambers | JCC is supported by the Singapore Ministry of Health's National Medical Research Council under its Singapore Translational Research Investigator (STaR) Award (NMRC/STaR/0028/2017). |
| Brian H. Chen | BHC was supported by Burroughs Wellcome Fund Inter-school Training Program in Metabolic Diseases and UCLA Genomic Analysis Training Program (NHGRI T32-HG002536) |
| Laura J Corbin | LJC is supported by a Wellcome Trust Investigator grant (PI: N.J. Timpson, 202802/Z/16/Z) ) and works within the Medical Research Council Integrative Epidemiology Unit [MC_UU_12013/3, MC_UU_00011]. |
| Josée Dupuis | JD was supported by NIDDK 5U01DK078616 |
| Segun Fatumo | SF was supported by funds from H3Africa Bioinformatics Network (H3ABioNet) Node CGRI/NABDA Abuja, Cambridge-Africa alborada research fund and NIH U01MH115485 |
| Timothy Frayling | TMF is supported by the European Research Council grant: 323195:SZ-245 50371-GLUCOSEGENES-FP7-IDEAS-ERC. |
| Anna L Gloyn | ALG is a Wellcome Trust Senior Fellow Basic Biomedical Science. This work was funded in Oxford by the Wellcome Trust (095101 [ALG], 200837 [A.L.G.], 06130 [A.L.G.], 203141 ], Medical Research Council (MR/L020149/1), European Union Horizon 2020 Programme (T2D Systems), and NIH (U01-DK105535; U01-DK085545) The research was funded by the National Institute for Health Research (NIHR) Oxford Biomedical Research Centre (BRC). The views expressed are those of the |

| | |
|---|---|
| | author(s) and not necessarily those of the NHS, the NIHR or the Department of Health. |
| Niels Grarup | The Novo Nordisk Foundation Center for Basic Metabolic Research is an independent Research Center at the University of Copenhagen partially funded by an unrestricted donation from the Novo Nordisk Foundation (www.metabol.ku.dk). |
| Fernando P. Hartwig | FPH was supported by the Brazilian National Council for Scientific and Technological Development (CNPq) [postdoctoral fellowship, process number: 153134/2018]. |
| Caroline Hayward | CH is supported by an MRC University Unit Programme Grant MC_PC_U127592696. |
| Wei Huang | HW is supported by funds from Shanghai Municipal Science and Technology Commission(18JC1420100) , The Shanghai Municipal Finance Bureau(Leading talents in Shanghai (2017). |
| Erik Ingelsson | EI is now an employee of GSK. |
| J. Wouter Jukema | Prof. Dr. J. W. Jukema is an Established Clinical Investigator of the Netherlands Heart Foundation (grant 2001 D 032). |
| Marika A. Kaakinen | MAK is supported by the European Commission under the Marie Curie Intra-European Fellowship (project MARVEL, PIEF-GA-2013-626461), the European Foundation for the Study of Diabetes (EFSD) Albert Renold Travel Fellowship, and the European Union’s Horizon 2020 research and innovation programme (LongITools, H2020-SC1-2019-874739). |
| Katherine A Kentistou | KAK is suuported by the MRC Doctoral Training Programme in Precision Medicine |
| Mika Kivimaki | M Kivimaki is supported by the MRC, UK (K013351, R024227, S011676), NIA, US (R01AG056477) and the Academy of Finland (311492). |
| Antje Körner | AK is supported by the Deutsche Forschungsgemeinschaft (DFG, German Research Foundation)–Project No 209933838 – CRC 1052“ for the Clinical Research Center “Obesity Mechanisms” CRC1052 C05 |
| Man Li | M.L. was supported by the NHLBI Cardiovascular Epidemiology Training grant T32HL007024 |
| Cecilia Lindgren | C.M.L is supported by the Li Ka Shing Foundation, WT-SSI/John Fell funds, Oxford, NIHR Oxford Biomedical Research Centre, Oxford, Widenlife and NIH (5P50HD028138-27) |
| Ching-Ti Liu | CTL was supported by NIDDK 5U01DK078616 |
| Jianjun Liu | Agency for Science, Technology and Research |
| Ruth J.F. Loos | RJFL is supported by funds from the NIH (R01DK110113; R01DK107786; R01HL142302; R01DK101855; R56HG010297) |
| Massimo Mangino | MM is supported by the National Institute for Health Research (NIHR)-funded BioResource, Clinical Research Facility and Biomedical Research Centre based at Guy’s and St Thomas’ NHS Foundation Trust in partnership with King’s College London. |
| Mark I McCarthy | MMcC is a Wellcome Investigator (212259/Z/18/Z) and an NIHR Senior Investigator. |
| James B Meigs | NIDDK R01DK078616, U01DK078616, K24DK080140 and an ADA Mentored Career Development Award |
| Andres Metspalu | AM was supported by Estonian Research Council IUT20-60 and European Union through the European Regional Development Fund [2014-2020.4.01.15-0012 GENTRANSMED]. |

| | |
|---|---|
| May E Montasser | MEM is supported by funds from AHA 17GRNT33661168, Regeneron 19112531, NIH U01HL137181, NIH R01DK118942, ADA 1-16-ICTS-112 |
| Dennis O. Mook-Kanamori | Dennis Mook-Kanamori is supported by Dutch Science Organization (ZonMW-VENI Grant 916.14.023). |
| Trevor A Mori | TAM is a recipient of an Australian National Health and Medical Research Council Research Fellowship |
| Jeffrey R O'Connell | JRO is supported by NIH U01HL137181 |
| Colin NA Palmer | CP acknowledges funding from National Institute for Health Research using Official Development Assistance (ODA) [INSPIRED 16/136/102]. |
| Inga Prokopenko | IP is funded by the World Cancer Research Fund (WCRF UK) and World Cancer Research Fund International (2017/1641), the Wellcome Trust (WT205915/Z/17/Z), and the European Union's Horizon 2020 research, and innovation programmes (DYNAhealth, H2020-PHC-2014-633595, LongITools, H2020-SC1-2019-874739). |
| Chelsea K. Raulerson | CKR was supported by NIH T32GM067553 |
| Dermot Reilly | DFR is a current employee of Janssen Research and Development. All work was completed while an employee of Merck, Sharp, and Dohme Corp. |
| Stephen S. Rich | SSR is supported by funds from the NIH (U01HL120393; DP3DK111906;P01HL136275) |
| Naveed Sattar | NS acknowledges funding support from the British Heart Foundation Research Excellence Award (RE/18/6/34217) |
| Bengt Sennblad | BS is financially supported by the Knut and Alice Wallenberg Foundation as part of the National Bioinformatics Infrastructure Sweden at SciLifeLab. |
| Cassandra N. Spracklen | CNS was supported by American Heart Association Postdoctoral Fellowship 15POST24470131 and 17POST33650016 |
| Rona J Strawbridge | RJS is supported by a UKRI Innovation–HDR-UK Fellowship (MR/S003061/1) |
| Paul RHJ Timmers | PRHJT is supported by the MRC Doctoral Training Programme in Precision Medicine |
| Nicholas J. Timpson | NJT is a Wellcome Trust Investigator (202802/Z/16/Z), is the PI of the Avon Longitudinal Study of Parents and Children (MRC & WT 217065/Z/19/Z), is supported by the University of Bristol NIHR Biomedical Research Centre (BRC-1215-20011), the MRC Integrative Epidemiology Unit (MC_UU_12013/3, MC_UU_00011) and works within the CRUK Integrative Cancer Epidemiology Programme (C18281/A19169). |
| Niek Verweij | Niek Verweij is supported by Dutch Science Organization (ZonMW-VENI Grant 016.186.125) |
| Jana V. van Vliet-Ostaptchouk | JVvVO was supported by a Diabetes Funds Junior Fellowship from the Dutch Diabetes Research Foundation (project no. 2013.81.1673) |
| Veronique Vitart | VV is supported by an MRC University Unit Programme Grant MC_PC_U127592696. |
| TGM Vrijkotte | Dr. T.G.M. Vrijkotte was supported by ZonMW (TOP 40–00812–98–11010) |
| Hugh Watkins | HW has received support from the National Institute for Health Research Oxford Biomedical Research Centre, the Oxford British Heart Foundation Centre of Research Excellence and National Heart, Lung, and Blood Institute grant U01HL117006-01A1. |

| | |
|---|---|
| Andrew Wood | ARW is supported by the European Research Council grant: 323195:SZ-245 50371-GLUCOSEGENES-FP7-IDEAS-ERC. |
| MH Zafarmand | Dr M.H. Zafarmand was supported by BBMRI-NL (CP2013-50) |
| Ele Zeggini | EZ acknowledges funding from Wellcome Trust (098051) |
| Xiaoshuai Zhang | XSZ was supported by China Scholarship Council (201406220101) |

**Cohort Funding and/or Other Acknowledgements**

| Cohort | Acknowledgements |
|---|---|
| AASC | None |
| ABCD | The ABCD study has been supported by grants from The Netherlands Organisation for Health Research and Development (ZonMW) and The Netherlands Heart Foundation. We thank all participating hospitals, obstetric clinics, general practitioners and primary schools for their assistance in implementing the ABCD study. We also gratefully acknowledge all the women and children who participated in this study for their cooperation. |
| AGES | The AGES study was funded by the National Institute on Aging (NIA) (N01-AG-12100) and HHSN27120120022C, Hjartavernd (the Icelandic Heart Association), and the Althingi (the Icelandic Parliament), with contributions from the Intramural Research Programs at the NIA, the National Heart, Lung, and Blood Institute (NHLBI), The authors are grateful to the study participants and the IHA staff. |
| ALSPAC | We are extremely grateful to all the families who took part in this study, the midwives for their help in recruiting them, and the whole ALSPAC team, which includes interviewers, computer and laboratory technicians, clerical workers, research scientists, volunteers, managers, receptionists and nurses. The UK Medical Research Council and Wellcome (Grant ref: 217065/Z/19/Z) and the University of Bristol provide core support for ALSPAC. This publication is the work of the authors and NJT and LJC will serve as guarantors for the contents of this paper. A comprehensive list of grants funding is available on the ALSPAC website (http://www.bristol.ac.uk/alspac/external/documents/grant-acknowledgements.pdf). This research was specifically funded by: National Institutes of Health (Grant ref: R01 DK077659, PI: D.Lawlor) (ALSPACchildren FI and FG); British Heart Foundation (Grant ref: SP/07/008/24066, PI: Debbie Lawlor) (ALSPACmothers FI and FG); and Wellcome Trust (Grant ref: WT088806) (ALSPACmothers genotype data). ALSPACchildren genotype data was generated by Sample Logistics and Genotyping Facilities at Wellcome Sanger Institute and LabCorp (Laboratory Corporation of America) using support from 23andMe. |
| AMISH | We gratefully thank our Amish community and research volunteers for their long-standing partnership in research, and acknowledge the dedication of our Amish liaisons, field workers and the Amish Research Clinic staff, without which these studies would not have been possible.The Amish studies are supported by grants and contracts from the NIH, including R01 AG18728, R01 HL088119, U01 |

| | |
|---|---|
| | GM074518, U01 HL072515, U01 HL84756, U01 HL137181, R01 DK54261,the University of Maryland General Clinical Research Center, grant M01 RR 16500, and the Mid-Atlantic Nutrition Obesity Research Center grant P30 DK72488, the Baltimore Diabetes Research and Training Center grant P60DK79637. |
| ARIC | The Atherosclerosis Risk in Communities study has been funded in whole or in part with Federal funds from the National Heart, Lung, and Blood Institute, National Institutes of Health, Department of Health and Human Services (contract numbers HHSN268201700001I, HHSN268201700002I, HHSN268201700003I, HHSN268201700004I and HHSN268201700005I), R01HL087641, R01HL059367 and R01HL086694; National Human Genome Research Institute contract U01HG004402; and National Institutes of Health contract HHSN268200625226C. The authors thank the staff and participants of the ARIC study for their important contributions. Infrastructure was partly supported by Grant Number UL1RR025005, a component of the National Institutes of Health and NIH Roadmap for Medical Research. |
| ASCOT | This work was funded by the National Institutes for Health Research (NIHR) as part of the portfolio of translational research of the NIHR Barts Biomedical Research Centre and the NIHR Biomedical Research Centre at Imperial College, the International Centre for Circulatory Health Charity and the Medical Research Council through G952010. We thank all ASCOT trial participants, physicians, nurses, and practices in the participating countries for their important contribution to the study |
| BC1936 | None |
| BES | The Beijing Eye Study (BES) was supported by National Natural Science Foundation of China (grant 81570835). |
| BetaGene | The BetaGene study were supported by National Institutes of Health grants [R01-DK-061628 and UL1-RR-031986] |
| BioMe | The Mount Sinai BioMe Biobank is supported by The Andrea and Charles Bronfman Philanthropies. |
| CAGE-GWAS1 | The CAGE Network studies were supported by grants for the Core Research for Evolutional Science and Technology (CREST) from the Japan Science Technology Agency; the Program for Promotion of Fundamental Studies in Health Sciences, National Institute of Biomedical Innovation Organization (NIBIO); and the Grant of National Center for Global Health and Medicine (NCGM). We thank the participants who made this work possible and who gave it value. We also thank Drs. Toshio Ogihara, Yukio Yamori, Akihiro Fujioka, Chikanori Makibayashi, Sekiharu Katsuya, Ken Sugimoto, Kei Kamide, and Ryuichi Morishita and the many physicians of the participating hospitals and medical institutions in Amagasaki Medical Association for their assistance in collecting the DNA samples and accompanying clinical information. |
| CAGE-KING | The KING Study was supported in part by Grants-in-Aid from MEXT (nos. 24390169, 16H05250, 25293144, 15K19242, 16H06277, and 19K19434) as well as by a grant from the Funding Program for Next-Generation World-Leading Researchers (NEXT Program, no. LS056). |
| CLHNS | We thank the Office of Population Studies Foundation research and data collection teams and the study participants who generously provided their time for this study. This work was supported by National Institutes of Health grants DK078150, TW005596 and HL085144; pilot funds from RR020649, ES010126, and DK056350; and the Office of Population Studies Foundation |

| | |
|---|---|
| CHNS | We thank the National Institute for Nutrition and Health, Chinese Center for Disease Control and Prevention, the Chinese National Human Genome Center at Shanghai, the Carolina Population Center, the University of North Carolina at Chapel Hill, and all of the participants and study investigators involved in the China Health and Nutrition Survey. Data collection and analysis was supported by the Carolina Population Center (P2C HD050924, T32 HD007168), the NIH (R01HD30880, R01 DK056350,R01DK104371, R24 HD050924, R01 HD38700, R01 DK072193 and U01 DK105561), the NIH Fogarty International Center (D43 TW009077, D43 TW007709), the China Ministry of Health, the Chinese National Human Genome Center at Shanghai, the China-Japan Friendship Hospital, and the Beijing Municipal Center for Disease Prevention and Control. |
| CHS | Cardiovascular Health Study: This CHS research was supported by NHLBI contracts HHSN268201200036C, HHSN268200800007C, HHSN268201800001C, N01HC55222, N01HC85079, N01HC85080, N01HC85081, N01HC85082, N01HC85083, N01HC85086; and NHLBI grants U01HL080295, R01HL087652, R01HL105756, R01HL103612, R01HL120393, and U01HL130114 with additional contribution from the National Institute of Neurological Disorders and Stroke (NINDS). Additional support was provided through R01AG023629 from the National Institute on Aging (NIA). A full list of principal CHS investigators and institutions can be found at CHS-NHLBI.org. The provision of genotyping data was supported in part by the National Center for Advancing Translational Sciences, CTSI grant UL1TR001881, and the National Institute of Diabetes and Digestive and Kidney Disease Diabetes Research Center (DRC) grant DK063491 to the Southern California Diabetes Endocrinology Research Center. The content is solely the responsibility of the authors and does not necessarily represent the official views of the National Institutes of Health. |
| CFS | The Cleveland Family Study has been supported by National Institutes of Health grants [R01-HL046380, KL2-RR024990, R35-HL135818, and R01-HL113338]. |
| CoLaus | The CoLaus study was and is supported by research grants from GlaxoSmithKline, the Faculty of Biology and Medicine of Lausanne, and the Swiss National Science Foundation (grants 33CSCO-122661, 33CS30-139468, 33CS30-148401, and 32473B-182210). The authors would like to thank all the people who participated in the recruitment of the participants, data collection and validation, particularly Nicole Bonvin, Yolande Barreau, Mathieu Firmann, François Bastardot, Julien Vaucher, Panagiotis Antiochos and Cédric Gubelmann. |
| COPSAC2000 | All funding received by COPSAC is listed on www.copsac.com. The Lundbeck Foundation (Grant no R16-A1694); The Ministry of Health (Grant no 903516); Danish Council for Strategic Research (Grant no 0603-00280B) and The Capital Region Research Foundation have provided core support to the COPSAC research center.We express our deepest gratitude to the children and families of the COPSAC2000 cohort study for all their support and commitment. We acknowledge and appreciate the unique efforts of the COPSAC research team. |
| CROATIA | The CROATIA_Vis ,CROATIA_Korcula and CROATIA_Split studies were funded by grants from the Medical Research Council (UK), European Commission Framework 6 project EUROSPAN (Contract No. LSHG-CT-2006-018947) and Republic of Croatia Ministry of Science, Education and Sports research grants. (108-1080315-0302). We would like to acknowledge the staff of several institutions in Croatia that supported the field work, including but not limited to The University of Split and Zagreb Medical Schools, Institute for Anthropological Research in Zagreb and Croatian Institute for Public Health. |

| | |
|---|---|
| DPS | The DPS has been financially supported by grants from the Academy of Finland (117844 and 40758, 211497, and 118590; The EVO funding of the Kuopio University Hospital from Ministry of Health and Social Affairs |
| DRSEXTRA | The DR's EXTRA Study was supported by grants to R. Rauramaa by the Ministry of Education and Culture of Finland (627;2004-2011), Academy of Finland (102318; 123885), Kuopio University Hospital , Finnish Diabetes Association, Finnish Heart Association, Päivikki and Sakari Sohlberg Foundation and by grants from European Commission FP6 Integrated Project (EXGENESIS); LSHM-CT-2004-005272, City of Kuopio and Social Insurance Institution of Finland (4/26/2010). |
| DGI | None |
| DIAGEN | The DIAGEN study was supported by the Commission of the European Communities, Directorate C - Public Health and Risk Assessment, Health & Consumer Protection, Grant Agreement number - 2004310 and by the Dresden University of Technology Funding Grant, Med Drive. We are grateful to all of the patients who cooperated in this study and to their referring physicians and diabetologists in Saxony. |
| DRECA | The DRECA studies were supported by the Andalusian Health Council and planned and contributed by primary care physicians of Seville who contributed for many years with the clinical information and laboratory tests used for the DRECA-1 and DRECA-2 studies. |
| EGCUT | EGCUT analysis were funded by EU H2020 grant 692145, Estonian Research Council Grant IUT20-60, IUT24-6, and European Union through the European Regional Development Fund Project No. 2014-2020.4.01.15-0012 GENTRANSMED and 2014-2020.4.01.16-0125. Data analyzes were carried out in part in the High-Performance Computing Center of University of Tartu |
| Ely | The Ely study was supported by the Medical Research Council (MC_UU_12015/1) and NHS Research and Development. |
| EPIC | The EPIC-Norfolk study (DOI 10.22025/2019.10.105.00004) has received funding from the Medical Research Council (MR/N003284/1 and MC-UU_12015/1) and Cancer Research UK (C864/A14136). The genetics work in the EPIC-Norfolk study was funded by the Medical Research Council ((MC_PC_13048)).We are grateful to all the participants who have been part of the project and to the many members of the study teams at the University of Cambridge who have enabled this research. |
| EPIHEALTH | Funded by the SFO Epihealth. |
| ERF | ERF study is grateful to all study participants and their relatives, general practitioners and neurologists for their contributions and to P. Veraart for her help in genealogy, J. Vergeer for the supervision of the laboratory work and P. Snijders for his help in data collection.Erasmus Rucphen Family (ERF) was supported by the Consortium for Systems Biology (NCSB), both within the framework of the Netherlands Genomics Initiative (NGI)/Netherlands Organisation for Scientific Research (NWO). ERF study as a part of EUROSPAN (European Special Populations Research Network) was supported by European Commission FP6 STRP grant number 018947 (LSHG-CT-2006-01947) and also received funding from the European Community's Seventh Framework Programme (FP7/2007-2013)/grant agreement HEALTH-F4-2007-201413 by the European Commission under the programme “Quality of Life and Management of the Living Resources” of 5th Framework Programme (no. QLG2-CT-2002-01254) as well as FP7 project EUROHEADPAIN (nr 602633). The ERF study was further supported by ENGAGE consortium and CMSB. High-throughput analysis of the |

| | |
|---|---|
| | ERF data was supported by joint grant from Netherlands Organization for Scientific Research and the Russian Foundation for Basic Research (NWO-RFBR 047.017.043)The exome-chip measurements have been funded by the Netherlands Organization for Scientific Research (NWO; project number 184021007) and by the Rainbow Project (RP10; Netherlands Exome Chip Project) of the Biobanking and Biomolecular Research Infrastructure Netherlands (BBMRI-NL; www.bbmri.nl (http://www.bbmri.nl) ). Ayse Demirkan is supported by a Veni grant (2015) from ZonMw. Ayse Demirkan, Jun Liu and Cornelia van Duijn have used exchange grants from PRECEDI. The funders had no role in study design, data collection and analysis, decision to publish, or preparation of the manuscripts. |
| FamHS | The Family Heart Study (FamHS) was supported by NIH/NHLBI grants RO1-HL-087700 and RO1-HL-088215. |
| Fenland | The Fenland Study (10.22025/2017.10.101.00001) is funded by the Medical Research Council (MC_UU_12015/1). We are grateful to all the volunteers and to the General Practitioners and practice staff for assistance with recruitment. We thank the Fenland Study Investigators, Fenland Study Co-ordination team and the Epidemiology Field, Data and Laboratory teams. We further acknowledge support for genomics and metabolomics from the Medical Research Council (MC_PC_13046). |
| FIN-D2D2007 | The FIN-D2D study has been financially supported by the hospital districts of Pirkanmaa, South Ostrobothnia, and Central Finland, the Finnish National Public Health Institute (current National Institute for Health and Welfare), the Finnish Diabetes Association, the Ministry of Social Affairs and Health in Finland, the Academy of Finland (grant number 129293),Commission of the European Communities, Directorate C-Public Health (grant agreement no. 2004310) and Finland’s Slottery Machine Association. |
| FHS | The Framingham Heart Study (FHS) was supported by the National Heart, Lung and Blood Institute's Framingham Heart Study Contract Nos. N01-HC-25195 and HHSN268201500001I and 75N92019D00031) and its contract with Affymetrix, Inc for genotyping services (Contract No. N02-HL-6-4278), and by NIDDK R01DK078616 and U01DK078616. |
| French Adult Control | The patients were recruited by the laboratory "Integrated Genomics and Metabolic Diseases Modeling" (UMR 8199 CNRS / Université de Lille 2 / Institut Pasteur de Lille) of Pr. Philippe Froguel. |
| French Adult Obese | The patients were recruited by the laboratory "Integrated Genomics and Metabolic Diseases Modeling" (UMR 8199 CNRS / Université de Lille 2 / Institut Pasteur de Lille) of Pr. Philippe Froguel. |
| French Young Control | The patients were recruited by the laboratory "Integrated Genomics and Metabolic Diseases Modeling" (UMR 8199 CNRS / Université de Lille 2 / Institut Pasteur de Lille) of Pr. Philippe Froguel. |
| French Young Obese | The patients were recruited by the laboratory "Integrated Genomics and Metabolic Diseases Modeling" (UMR 8199 CNRS / Université de Lille 2 / Institut Pasteur de Lille) of Pr. Philippe Froguel. |
| FUSION | Support for FUSION was provided by NIH grants R01-DK062370 (to M.B.), R01-DK072193 (to K.L.M.), and intramural project number 1Z01-HG000024 (to F.S.C.). Genome-wide genotyping was conducted by the Johns Hopkins University Genetic Resources Core Facility SNP Center at the Center for Inherited Disease Research (CIDR), with support from CIDR NIH contract no. N01-HG-65403. |

| | |
|---|---|
| Generation Scotland | Generation Scotland received core funding from the Chief Scientist Office of the Scottish Government Health Directorate CZD/16/6 , the Scottish Funding Council HR03006 and the Wellcome Trust through a Strategic Award (reference 104036/Z/14/Z) for Stratifying Resilience and Depression Longitudinally (STRADL). Genotyping of the GS:SFHS samples was carried out by staff at the Genetics Core Laboratory at the Clinical Research Facility, University of Edinburgh, Scotland and was funded by the UK's Medical Research Council. |
| GeneSTAR | GeneSTAR was supported by NIH grants through the National Heart, Lung, and Blood Institute (HL58625, HL59684, HL49762, HL071025, U01HL72518, and HL087698) and the National Institute of Nursing Research (NR0224103, NR008153) and by M01-RR000052 to the Johns Hopkins General Clinical Research Center. |
| GENOA | Support for GENOA was provided by the National Heart, Lung and Blood Institute (HL119443, HL118305, HL054464, HL054457, HL054481, HL071917 and HL087660) of the National Institutes of Health. Genotyping was performed at the Mayo Clinic (Stephen T. Turner, MD, Mariza de Andrade PhD, Julie Cunningham, PhD). We thank Eric Boerwinkle, PhD and Megan L. Grove from the Human Genetics Center and Institute of Molecular Medicine and Division of Epidemiology, University of Texas Health Science Center, Houston, Texas, USA for their help with genotyping. We would also like to thank the families that participated in the GENOA study. |
| GLACIER | None |
| GoDARTS | We are grateful to all the participants in this study, the general practitioners, the Scottish School of Primary Care for their help in recruiting the participants, and to the whole team, which includes interviewers, computer and laboratory technicians, clerical workers, research scientists, volunteers, managers, receptionists, and nurses. The study complies with the Declaration of Helsinki. We acknowledge the support of the Health Informatics Centre, University of Dundee for managing and supplying the anonymised data and NHS Tayside, the original data owner. The Wellcome Trust United Kingdom Type 2 Diabetes Case Control Collection (GoDARTS) was funded by The Wellcome Trust (072960/Z/03/Z, 084726/Z/08/Z, 084727/Z/08/Z, 085475/Z/08/Z, 085475/B/08/Z) |
| GOYA (Male) | We thank all the participants of the study the MRC centre for Causal Analyses in Translational Epidemiology (MRC CAiTE) |
| HANDLS | The Healthy Aging in Neighborhoods of Diversity across the Life Span (HANDLS) study was supported by the Intramural Research Program of the NIH, National Institute on Aging and the National Center on Minority Health and Health Disparities (project # Z01-AG000513 and human subjects protocol number 09-AG-N248). |
| Health2006 | The Health 2006 was financially supported by grants from the Velux Foundation; The Danish Medical Research Council, Danish Agency for Science, Technology and Innovation; The Aase and Ejner Danielsens Foundation; ALK-Abello A/S, Hørsholm, Denmark, and Research Centre for Prevention and Health, the Capital Region of Denmark. |
| HELIC MANOLIS and Helic Pomak | This work was funded by the Wellcome Trust (098051) and the European Research Council (ERC-2011-StG 280559-SEPI). The MANOLIS cohort is named in honour of Manolis Giannakakis, 1978-2010. We thank the residents of the Mylopotamos villages, and of the Pomak villages, for taking part. |
| HTN | None |

| | |
|---|---|
| IMPROVE | The IMPROVE study was supported by the European Commission (Contract number: QLG1-CT-2002-00896), the Swedish Heart-Lung Foundation, the Swedish Research Council (projects 8691 and 0593), the Stockholm County Council (project 562183), Academy of Finland (Grant #110413) the British Heart Foundation (RG2008/014) and the Italian Ministry of Health (Ricerca Corrente) |
| INCHIANTI | None |
| Inter99 | The Inter99 was initiated by Torben Jørgensen (PI), Knut Borch-Johnsen (co-PI), Hans Ibsen and Troels F. Thomsen. The steering committee comprises the former two and Charlotta Pisinger. The study was financially supported by research grants from the Danish Research Council, the Danish Centre for Health Technology Assessment, Novo Nordisk Inc., Research Foundation of Copenhagen County, Ministry of Internal Affairs and Health, the Danish Heart Foundation, the Danish Pharmaceutical Association, the Augustinus Foundation, the Ib Henriksen Foundation, the Becket Foundation, and the Danish Diabetes Association. |
| InterAct | The EPIC-InterAct Study: We thank all EPIC participants and staff and the InterAct Consortium members for their contributions to the study. The InterAct project received funding from the European Union (Integrated Project LSHM-CT-2006-037197 in the Framework Programme 6 of the European Community). We thank staff from the technical, field epidemiology and data teams of the Medical Research Council Epidemiology Unit in Cambridge, UK, for carrying out sample preparation, DNA provision and quality control, genotyping and data handling work. |
| IRAS | The Insulin Resistance Atherosclerosis Study (IRAS) was supported by NIH grants HL047887, HL047889, HL047890, HL47902, DK085175 and DK118062. |
| IRASFS | The Insulin Resistance Atherosclerosis Family Study (IRASFS) was supported by NIH grants HL060944, HL061019, HL060919, DK085175 and DK118062. |
| JHS | The Jackson Heart Study (JHS) is supported and conducted in collaboration with Jackson State University (HHSN268201800013I), Tougaloo College (HHSN268201800014I), the Mississippi State Department of Health (HHSN268201800015I) and the University of Mississippi Medical Center (HHSN268201800010I, HHSN268201800011I and HHSN268201800012I) contracts from the National Heart, Lung, and Blood Institute (NHLBI) and the National Institute on Minority Health and Health Disparities (NIMHD). The authors also wish to thank the staff and participants of the JHS. The views expressed in this manuscript are those of the authors and do not necessarily represent the views of the National Heart, Lung, and Blood Institute; the National Institutes of Health; or the U.S. Department of Health and Human Services. |
| KARE | This work was supported by an intramural grant from the Korea National Institute of Health (2019-NG-053-00). This study was performed with bioresources from National Biobank of Korea, the Centers for Disease Control and Prevention, Republic of Korea. |
| KORA F4 | The KORA study was initiated and financed by the Helmholtz Zentrum München – German Research Center for Environmental Health, which is funded by the German Federal Ministry of Education and Research (BMBF) and by the State of Bavaria. Furthermore, KORA research was financed by a grant from the BMBF to the German Center for Diabetes Research (DZD) It was also supported within the Munich Center of Health Sciences (MC-Health), Ludwig-Maximilians-Universität, as part of LMUinnovativ. The German Diabetes Center was supported by the Ministry of Culture and Science of the state of North Rhine-Westphalia (Düsseldorf, Germany) and the German Federal Ministry of Health (Berlin, |

| | |
|---|---|
| | Germany). This study was supported in part by a grant from the German Federal Ministry of Education and Research to the German Center for Diabetes Research (DZD). |
| Leiden Longevity Study | The Leiden Longevity Study has received funding from the European Union's Seventh Framework Programme (FP7/2007-2011) under grant agreement number 259679. This study was financially supported by the Innovation-Oriented Research Program on Genomics (SenterNovem IGE05007), the Centre for Medical Systems Biology and the Netherlands Consortium for Healthy Ageing (grant 050-060-810), all in the framework of the Netherlands Genomics Initiative, Netherlands Organization for Scientific Research (NWO), and by BBMRI-NL, a Research Infrastructure financed by the Dutch government (NWO 184.021.007). |
| LEIPZIG-adults | This work was supported by grants from the Federal Ministry of Education and Research (BMBF), Germany (the Kompetenznetz Adipositas - Competence network for Obesity - German Obesity Biomaterial Bank; FKZ 01GI1128 and FKZ: 01EO1501; AD2-060E, AD2-6E95, K7-117), and from the Deutsche Forschungsgemeinschaft (DFG, German Research Foundation) – Projektnummer 209933838 – SFB 1052 (C01, B01, B03). |
| LEIPZIG-kids | LEIPZIG Kids were supported by the Federal Ministry of Education and Research (BMBF), Germany, Integrated Research and Treatment Centre (IFB) Adiposity Diseases FKZ: 01EO1001, by the by the LIFE (Leipzig Research Center for Civilization Diseases, Universität Leipzig), funded by the European Union, by the European Regional Development Fund (ERFD) by means of the Free State of Saxony within the framework of the excellence initiative |
| Lifelines | The Lifelines Cohort Study, and generation and management of GWAS genotype data for the Lifelines Cohort Study is supported by the Netherlands Organization of Scientific Research NWO (grant 175.010.2007.006), the Economic Structure Enhancing Fund (FES) of the Dutch government, the Ministry of Economic Affairs, the Ministry of Education, Culture and Science, the Ministry for Health, Welfare and Sports, the Northern Netherlands Collaboration of Provinces (SNN), the Province of Groningen, University Medical Center Groningen, the University of Groningen, Dutch Kidney Foundation and Dutch Diabetes Research Foundation. The authors wish to acknowledge the services of the Lifelines Cohort Study, the contributing research centers delivering data to Lifelines, and all the study participants. |
| Living Biobank | The Living Biobank was supported by grants from National Medical Research Council (NMRC), Biomedical Research Council (BMRC), National University of Singapore and National University Health System, and Ministry of Health, Singapore, and Merck Sharp & Dohme Corp. |
| LOLIPOP (EW610, EWA, EWP, IA317, IA610, IAP, OmniEE) | The LOLIPOP study is supported by the National Institute for Health Research (NIHR) Comprehensive Biomedical Research Centre Imperial College Healthcare NHS Trust, the British Heart Foundation (SP/04/002), the Medical Research Council (G0601966, G0700931), the Wellcome Trust (084723/Z/08/Z, 090532 & 098381) the NIHR (RP-PG-0407-10371), the NIHR Official Development Assistance (ODA, award 16/136/68), the European Union FP7 (EpiMigrant, 279143) and H2020 programs (iHealth-T2D, 643774). We acknowledge support of the MRC-PHE Centre for Environment and Health, and the NIHR Health Protection Research Unit on Health Impact of Environmental Hazards. The work was carried out in part at the NIHR/Wellcome Trust Imperial Clinical Research Facility. The views expressed are those of the author(s) and not necessarily those of the Imperial College Healthcare NHS Trust, the NHS, the NIHR or the Department of |

| | |
|---|---|
| | Health. We thank the participants and research staff who made the study possible. |
| LURIC | LURIC has received funding from the 6th Framework Program (integrated project Bloodomics, grant LSHM-CT-2004-503485) and from the 7th Framework Programs Atheroremo (grant agreement number 201668) and RiskyCAD (grant agreement number 305739) of the European Union. |
| MACAD | This research was supported by the GUARDIAN Study DK085175 from the (MACAD), HL0697974 (HTN-IR), HL055798 (NIDDM-Atherosclerosis), and DK079888 (work related to insulin clearance in HTN-IR, MACAD and NIDDM-Atherosclerosis). The provision of genotyping data was supported in part by the National Center for Advancing Translational Sciences, CTSI grant UL1TR001881, and the National Institute of Diabetes and Digestive and Kidney Disease Diabetes Research (DRC) grant DK063491. |
| MEGA | We thank the directors of the Anticoagulation Clinics of Amersfoort (M.H.H. Kramer), Amsterdam (M. Remkes), Leiden (F.J.M.van der Meer), The Hague (E. van Meegen), Rotterdam(A.A.H. Kasbergen), and Utrecht (J. de Vries-Goldschmeding) who made the recruitment of patients possible. The interviewers (J.C.M. van den Berg, B. Berbee, S. van der Leden, M. Roosen, and E.C. Willems of Brilman) performed the blood draws. We also thank I. de Jonge, R. Roelofsen, M. Streevelaar, L.M.J. Timmers, and J.J.Schreijer for their secretarial and administrative support and data management. C.M. Cobbaert, C.J.M. van Dijk, R. van Eck, J. van der Meijden, P.J. Noordijk, L. Mahic and T. Visser performed the laboratory measurements. This research was supported by The Netherlands Heart Foundation (NHS 98.113), the Dutch Cancer Foundation (RUL 99/1992), and The Netherlands Organization for Scientific Research (912-03-033 2003). |
| MESA | Support for the Multi-Ethnic Study of Atherosclerosis (MESA) projects are conducted and supported by the National Heart, Lung, and Blood Institute (NHLBI) in collaboration with MESA investigators. Support for MESA is provided by contracts 75N92020D00001, HHSN268201500003I, N01-HC-95159, 75N92020D00005, N01-HC-95160, 75N92020D00002, N01-HC-95161, 75N92020D00003, N01-HC-95162, 75N92020D00006, N01-HC-95163, 75N92020D00004, N01-HC-95164, 75N92020D00007, N01-HC-95165, N01-HC-95166, N01-HC-95167, N01-HC-95168, N01-HC-95169, UL1-TR-000040, UL1-TR-001079, and UL1-TR-001420. Also supported in part by the National Center for Advancing Translational Sciences, CTSI grant UL1TR001881, and the National Institute of Diabetes and Digestive and Kidney Disease Diabetes Research Center (DRC) grant DK063491 to the Southern California Diabetes Endocrinology Research Center. |
| METSIM | The METSIM study was funded by the Academy of Finland (grants no. 77299 and 124243). Additional support was provided by US National Institutes of Health R01 DK062370, R01 DK093757 and Z01 HG000024. |
| MICROS | We thank all study participants, all primary care practitioners, and the personnel of the Hospital of Silandro (Department of Laboratory Medicine) for their participation and collaboration in the research project. The study was supported by the Ministry of Health and Department of Educational Assistance, University and Research of the Autonomous Province of Bolzano, the South Tyrolean Sparkasse Foundation, and the European Union framework program 6 EUROSPAN project (contract no. LSHG-CT2006-018947) |

| | |
|---|---|
| Nagahama | We are grateful to the Nagahama City Office and the nonprofit organization Zeroji Club for their assistance in performing the Nagahama study. |
| NEO | The authors of the NEO study thank all individuals who participated in the Netherlands Epidemiology of Obesity study, all participating general practitioners for inviting eligible participants and all research nurses for collection of the data. We thank the NEO study group,Pat van Beelen,Petra Noordijk and Ingeborg de Jonge for the coordination, lab and data management of the NEO study. The genotyping in the NEO study wassupported by the Centre National de Génotypage (Paris, France), headed by Jean-Francois Deleuze. The NEO study is supported by the participating Departments, the Division and the Board of Directors of the Leiden University Medical Center, and by the Leiden University, Research Profile Area Vascular and Regenerative Medicine. |
| NFBC1966 | We thank all cohort members and researchers who participated in the 31 yrs study. We also wish to acknowledge the work of the NFBC project center. NFBC1966 received financial support from University of Oulu Grant no. 65354, Oulu University Hospital Grant no. 2/97, 8/97, Ministry of Health and Social Affairs Grant no. 23/251/97, 160/97, 190/97, National Institute for Health and Welfare, Helsinki Grant no. 54121, Regional Institute of Occupational Health, Oulu, Finland Grant no. 50621, 54231. |
| NFBC1986 | We thank all cohort members and researchers who have participated in the study. We also wish to acknowledge the work of the NFBC project center. EU QLG1-CT-2000-01643 (EUROBLCS) Grant no. E51560, NorFA Grant no. 731, 20056, 30167, USA / NIHH 2000 G DF682 Grant no. 50945. |
| NHAPC | This study is supported by the Major Project of the Ministry of Science and Technology of China (2017YFC0909700, 2016YFC1304903), the National Natural Science Foundation of China (81700700), and the Chinese Academy of Sciences (ZDBS-SSW-DQC-02, ZDRW-ZS-2016-8). |
| NIDDM | None |
| NSHD | This work was funded by the Medical Research Council (MC_UU_12019/1). We are very grateful to the members of this birth cohort for their continuing interest and participation in the study. We would like to acknowledge the Swallow group, UCL, who performed the DNA extractions (Rousseau, et al 2006). DOI: 10.1111/j.1469-1809.2006.00250.x |
| NTR | Funding was obtained from the Netherlands Organization for Scientific Research (NWO) and The Netherlands Organisation for Health Research and Development (ZonMW) grants 904-61-090, 985-10-002, 912-10-020, 904-61-193,480-04-004, 463-06-001, 451-04-034, 400-05-717, Addiction-31160008, 016-115-035, 481-08-011, 056-32-010, Middelgroot-911-09-032, OCW_NWO Gravity program – 024.001.003, NWO-Groot 480-15-001/674, Center for Medical Systems Biology (CSMB, NWO Genomics), NBIC/BioAssist/RK(2008.024), Biobanking and Biomolecular Resources Research Infrastructure (BBMRI –NL, 184.021.007 and 184.033.111); Spinozapremie (NWO- 56-464-14192), KNAW Academy Professor Award (PAH/6635) and University Research Fellow grant (URF) to DIB; Amsterdam Public Health research institute (former EMGO+) , Neuroscience Amsterdam research institute (former NCA) ; the European Science Foundation (ESF, EU/QLRT-2001-01254), the European Community's Seventh Framework Program (FP7- HEALTH-F4-2007-2013, grant 01413: ENGAGE and grant 602768: ACTION); the European Research Council (ERC Starting 284167, ERC Consolidator 771057, ERC Advanced 230374), Rutgers University Cell and DNA Repository (NIMH U24 MH068457-06), the National Institutes of Health (NIH, R01D0042157-01A1, R01MH58799-03, MH081802, DA018673, R01 DK092127-04, Grand |

| | |
|---|---|
| | Opportunity grants 1RC2 MH089951, and 1RC2 MH089995); the Avera Institute for Human Genetics, Sioux Falls, South Dakota (USA). Part of the genotyping and analyses were funded by the Genetic Association Information Network (GAIN) of the Foundation for the National Institutes of Health. Computing was supported by NWO through grant 2018/EW/00408559, BiG Grid, the Dutch e-Science Grid and SURFSARA. |
| ORCADES | The Orkney Complex Disease Study (ORCADES) was supported by the Chief Scientist Office of the Scottish Government (CZB/4/276, CZB/4/710), a Royal Society URF to J.F.W., the MRC Human Genetics Unit quinquennial programme “QTL in Health and Disease”, Arthritis Research UK and the European Union framework program 6 EUROSPAN project (contract no. LSHG-CT-2006-018947). DNA extractions were performed at the Wellcome Trust Clinical Research Facility in Edinburgh. We would like to acknowledge the invaluable contributions of the research nurses in Orkney, the administrative team in Edinburgh and the people of Orkney. |
| Oxford Biobank | Supported by the NIHR Oxford Biomedical Research Centre, Obesity Diet and Nutrition Theme |
| PELOTAS | The 1982 Pelotas Birth Cohort Study is conducted by the Postgraduate Program in Epidemiology at Universidade Federal de Pelotas with the collaboration of the Brazilian Public Health Association (ABRASCO). From 2004 to 2013, the Wellcome Trust supported the study. The International Development Research Center, World Health Organization, Overseas Development Administration, European Union, National Support Program for Centers of Excellence (PRONEX), the Brazilian National Research Council (CNPq), and the Brazilian Ministry of Health supported previous phases of the study.<br>Genotyping of 1982 Pelotas Birth Cohort Study participants was supported by the Department of Science and Technology (DECIT, Ministry of Health) and National Fund for Scientific and Technological Development (FNDCT, Ministry of Science and Technology), Funding of Studies and Projects (FINEP, Ministry of Science and Technology, Brazil), Coordination of Improvement of Higher Education Personnel (CAPES, Ministry of Education, Brazil). |
| PIVUS/ULSAM | The PIVUS/ULSAM studies were supported by Wellcome Trust Grants WT098017, WT064890, WT090532, Uppsala University, Uppsala University Hospital, the Swedish Research Council and the Swedish Heart-Lung Foundation. |
| Prevend | PREVEND genetics is supported by the Dutch Kidney Foundation (Grant E033), the EU project grant GENECURE (FP-6 LSHM CT 2006 037697), the National Institutes of Health (Grant 2R01LM010098), The Netherlands Organization for Health Research and Development (NWO-Groot Grant 175.010.2007.006, NWO VENI Grant 916.761.70, ZonMw Grant 90.700.441) and the Dutch Inter-University Cardiology Institute Netherlands (ICIN). |
| Procardis | PROCARDIS was supported by the BHF, European Community Sixth Framework Program (LSHM-CT- 2007-037273), AstraZeneca, the Swedish Research Council (8691), the Knut and Alice Wallenberg Foundation, the Swedish Heart-Lung Foundation, the Torsten and Ragnar Söderberg Foundation, the Strategic Cardiovascular Program of Karolinska Institutet and Stockholm County Council, the Foundation for Strategic Research and the Stockholm County Council (560283), HEALTH-F2-2013-601456 (CVGenes@Target). |
| PROSPER | The PROSPER study was supported by an investigator initiated grant obtained from Bristol-Myers Squibb.Support for genotyping was provided by the seventh framework program of the European commission (grant 223004) and by the |

| | |
|---|---|
| | Netherlands Genomics Initiative (Netherlands Consortium for Healthy Aging grant 050-060-810). |
| RHS | The Ragama Health Study (RHS) was supported by the International Medical Centre of Japan |
| RISC | The RISC Study was supported by European Union Grant QLG1-CT-2001-01252 |
| Rotterdam Study | The Rotterdam Study is funded by Erasmus Medical Center and Erasmus University, Rotterdam, Netherlands Organization for the Health Research and Development (ZonMw), the Research Institute for Diseases in the Elderly (RIDE), the Ministry of Education, Culture and Science, the Ministry for Health, Welfare and Sports, the European Commission (DG XII), and the Municipality of Rotterdam. The authors are grateful to the study participants, the staff from the Rotterdam Study and the participating general practitioners and pharmacists. |
| SardiNIA | The SardiNIA study was supported by the Intramural Research Program of the US National Institutes of Health, National Institute on Aging, contracts N01-AG-1-2109 and HHSN271201100005C, and by Sardinian Autonomous Region (L.R. 7/2009) grant cRP3-154. |
| SCARFSHEEP | The SCARF-SHEEP study was supported by grants from the Swedish Reserach Council and the Swedish Heart and Lung Foundation, the Cardiovascular Programme at Karolinska Institutet, the Strategic Support for Epidemiological Research at Karolinska Institutet, the Stockholm County Council |
| SIGMA | The SIGMA Cohort is partially supported by the National Council of Science and Technology (CONACyT) (grant 262070). This work was partially supported by the Carlos Slim Health Institute. We thank Saúl Cano-Colín for technical assistance. |
| SEED (SCES, SiMES, SINDI) | The Singapore Chinese Eye Study (SCES), Singapore Malay Eye Study (SiMES) and Singapore Indian Eye Study (SINDI) are supported by the National Medical Research Council (NMRC), Singapore (grants 0796/2003, 1176/2008, 1149/2008, STaR/0003/2008, 1249/2010, CG/SERI/2010, CIRG/1371/2013, and CIRG/1417/2015), and Biomedical Research Council (BMRC), Singapore (08/1/35/19/550 and 09/1/35/19/616). |
| SP2 | The Singapore Prospective Study Program (SP2) were supported by the individual research grant and clinician scientist award schemes from the National Medical Research Council (NMRC) and the Biomedical Research Council (BMRC) of Singapore. |
| HCHS/SOL | The Hispanic Community Health Study/Study of Latinos is a collaborative study supported by contracts from the National Heart, Lung, and Blood Institute (NHLBI) to the University of North Carolina (HHSN268201300001I / N01-HC-65233), University of Miami (HHSN268201300004I / N01-HC-65234), Albert Einstein College of Medicine (HHSN268201300002I / N01-HC-65235), University of Illinois at Chicago – HHSN268201300003I / N01-HC-65236 Northwestern Univ), and San Diego State University (HHSN268201300005I / N01-HC-65237). The following Institutes/Centers/Offices have contributed to the HCHS/SOL through a transfer of funds to the NHLBI: National Institute on Minority Health and Health Disparities, National Institute on Deafness and Other Communication Disorders, National Institute of Dental and Craniofacial Research, National Institute of Diabetes and Digestive and Kidney Diseases, National Institute of Neurological Disorders and Stroke, NIH Institution-Office of Dietary Supplements. The Genetic Analysis Center at the University of Washington was supported by NHLBI and NIDCR contracts (HHSN268201300005C AM03 and MOD03). |
| SORBS | This work was supported by grants from the Federal Ministry of Education and Research (BMBF), Germany - FKZ: 01EO1501 (AD2-060E, AD2-6E95, AD2-7123); |

| | |
|---|---|
| | the Deutsche Forschungsgemeinschaft (DFG, German Research Foundation) – Projektnummer 209933838 – SFB 1052 (C01, B01, B03), SPP 1629 (TO 718-2 ) and the German Diabetes Association. We thank all those who participated in the study. Sincere thanks are given to Dr. Knut Krohn (University of Leipzig) for the genotyping support. |
| TAICHI | The TAICHI-G study was supported by grants from the National Health Research Institutes, Taiwan (PH-099-PP-03, PH-100-PP-03, and PH-101-PP-03); the National Science Council, Taiwan (NSC 101-2314-B-075A-006-MY3, MOST 104-2314-B-075A-006-MY3, MOST 104-2314-B-075A-007, and MOST 105-2314-B-075A-003); and the Taichung Veterans General Hospital, Taiwan (TCVGH-1020101C, TCVGH-1020102D, TCVGH-1023102B, TCVGH-1023107D, TCVGH-1030101C, TCVGH-1030105D, TCVGH-1033503C, TCVGH-1033102B, TCVGH-1033108D, TCVGH-1040101C, TCVGH-1040102D, TCVGH-1043504C, and TCVGH-1043104B). The provision of genotyping data was supported in part by the National Center for Advancing Translational Sciences, CTSI grant UL1TR001881, and the National Institute of Diabetes and Digestive and Kidney Disease Diabetes Research (DRC) grant DK063491. |
| The CRC Study | None |
| TRAILS | Participating centers of TRAILS (TRacking Adolescents' Individual Lives Survey) include the University Medical Center and University of Participating centers of TRAILS (TRacking Adolescents' Individual Lives Survey) include the University Medical Center and University of Groningen, the University of Utrecht, the Radboud Medical Center Nijmegen, and the Parnassia Bavo group, all in the Netherlands. TRAILS has been financially supported by various grants from the Netherlands Organization for Scientific Research NWO (Medical Research Council program grant GB-MW 940-38-011; ZonMW Brainpower grant 100-001-004; ZonMw Risk Behavior and Dependence grants 60-60600-97-118; ZonMw Culture and Health grant 261-98-710; Social Sciences Council medium-sized investment grants GB-MaGW 480-01-006 and GB-MaGW 480-07-001; Social Sciences Council project grants GB-MaGW 452-04-314 and GB-MaGW 452-06-004; NWO large-sized investment grant 175.010.2003.005; NWO Longitudinal Survey and Panel Funding 481-08-013 and 481-11-00; NWO Vici 016.130.002 and 453-16-007/2735; NWO Gravitation 024.001.003), the Dutch Ministry of Justice (WODC), the European Science Foundation (EuroSTRESS project FP-006), the European Research Council (ERC-2017-STG-757364 en ERC-CoG-2015-681466), Biobanking and Biomolecular Resources Research Infrastructure BBMRI-NL (CP 32), and the participating universities. Statistical analyses were carried out on the Genetic Cluster Computer (http://www.geneticcluster.org), which is financially supported by the Netherlands Scientific Organization (NWO 480-05-003) along with a supplement from the Dutch Brain Foundation. We are grateful to everyone who participated in this research or worked on this project to make it possible. |
| TRIPOD | None |
| TROMSO | The Tromsø study was funded by UiT the Arctic University of Norway |
| TWINGENE | This project was supported by grants the US National Institutes of Health (AG028555, AG08724, AG04563, AG10175, AG08861), the Swedish Heart-Lung Foundation, the Swedish Foundation for Strategic Research, the Royal Swedish Academy of Science, and ENGAGE (within the European Union Seventh Framework Programme, HEALTH-F4-2007-201413), the Ministry for Higher Education, the Swedish Research Council (M-2005-1112 and 2009-2298), GenomEUtwin (EU/QLRT-2001-01254; QLG2-CT-2002-01254), NIH grant DK U01- |

| | |
|---|---|
| | 066134, Knut and Alice Wallenberg Foundation (Wallenberg Academy Fellow), European Research Council (ERC Starting Grant), Swedish Diabetes Foundation (grant no. 2013-024), Swedish Research Council (grant no. 2012-1397), and Swedish Heart-Lung Foundation (20120197). The SNP Technology Platform is supported by Uppsala University, Uppsala University Hospital and the Swedish Research Council for Infrastructures. RJS is funded by the SRP diabetes programme at Karolinska Institutet. We thank Tomas Axelsson, Ann-Christine Wiman and Caisa Pöntinen at the SNP&SEQ Technology Platform in Uppsala (www.genotyping.se) for their excellent assistance with genotyping. The computations were performed on resources provided by SNIC through Uppsala Multidisciplinary Center for Advanced Computational Science (UPPMAX) under Project b2011036. |
| TWINSUK | TwinsUK is funded by the Wellcome Trust, Medical Research Council, European Union, the National Institute for Health Research (NIHR)-funded BioResource, Clinical Research Facility and Biomedical Research Centre based at Guy's and St Thomas' NHS Foundation Trust in partnership with King's College London. |
| TWSC | This work was supported by Academia Sinica GMM project. |
| Uganda | This work was funded by the Wellcome Trust, The Wellcome Sanger Institute (WT098051), the UK Medical Research Council (G0901213-92157, G0801566, and MR/K013491/1), and the Medical Research Council/Uganda Virus Research Institute Uganda Research Unit on AIDS core funding. |
| UKBB | This research has been conducted using the UK Biobank Resource under Application # 3913. |
| UKHLS | These data are from Understanding Society: The UK Household Longitudinal Study, which is led by the Institute for Social and Economic Research at the University of Essex and funded by the Economic and Social Research Council (Grant Number: ES/M008592/1). The data were collected by NatCen and the genome wide scan data were analysed by the Wellcome Trust Sanger Institute. Information on how to access the data can be found on the Understanding Society website https://www.understandingsociety.ac.uk/<br>Data governance was provided by the METADAC data access committee, funded by ESRC, Wellcome, and MRC. (2015-2018: Grant Number MR/N01104X/1 2018-2020: Grant Number ES/S008349/1) |
| Vanderbilt | This work is partially supported by NIH grants R01CA092585, R01CA064277, R01CA124558, UM1CA173640, and UM1CA182910. |
| VIKING | The Viking Health Study – Shetland (VIKING) was supported by the MRC Human Genetics Unit quinquennial programme grant "QTL in Health and Disease". DNA extractions and genotyping were performed at the Edinburgh Clinical Research Facility, University of Edinburgh. We would like to acknowledge the invaluable contributions of the research nurses in Shetland, the administrative team in Edinburgh and the people of Shetland. |
| The Raine Study | The Raine Study acknowledges the National Health and Medical Research Council (NHMRC) for their long-term contribution to funding the study over the last 30 years. Core Management of the Raine Study has been funded by the University of Western Australia (UWA), Curtin University, the Raine Medical Research Foundation, the Telethon Kids Institute, the Women's and Infants Research Foundation, Edith Cowan University, Murdoch University, and the University of Notre Dame. This study was supported by the National Health and Medical Research Council of Australia [grant numbers 572613, 403981 and 003209] and the Canadian Institutes of Health Research [grant number MOP-82893]. We also |

| | |
|---|---|
| | like to acknowledge The University of Western Australia (Division of Obstetrics and Gynaecology, King Edward Memorial Hospital and Medical School, Royal Perth Hospital), and Telethon Kids Institute for providing in-kind support for the storage and curation of biological samples. All analytic work was supported by resources provided by the Pawsey Supercomputing Centre with funding from the Australian Government and the Government of Western Australia. |
| WHI | The Women's Health Initiative (WHI) program is funded by the National Heart, Lung, and Blood Institute, National Institutes of Health, and the United States Department of Health and Human Services. |
| Whitehall II | The Whitehall II study has been supported by grants from the UK Medical Research Council (MRC K013351, R024227, S011676); the British Heart Foundation (RG/16/11/32334, PG/11/63/29011 and RG/13/2/30098); the British Health and Safety Executive; the British Department of Health; the National Heart, Lung, and Blood Institute (R01HL036310); the National Institute on Aging, National Institute of Health (NIA R01AG056477, R01AG034454); the Economic and Social Research Council (ES/J023299/1). |
| WGHS | The WGHS has been supported supported by the National Heart, Lung, and Blood Institute (HL043851 and HL080467) and the National Cancer Institute (CA047988 and UM1CA182913), with funding for genotyping provided by Amgen. |
